# *MEG3* Enhances Survival of Developing Human Neurons with *CLCN4*-Linked Autophagy Impairment

**DOI:** 10.1101/2025.07.16.665078

**Authors:** Dayeon Kim, Yongjun Koh, Hyeon Seong Jeong, Jaehyung Kim, Hyunsu Do, Geurim Son, Yeni Kim, Donghyuk Kim, Hyun-Ho Lim, Jong-Eun Park, Jinju Han

**Affiliations:** Graduate School of Medical Science and Engineering, Korea Advanced Institute of Science and Technology (KAIST), Daejeon, 34051, Republic of Korea; Neurovascular Unit Research Group, Korea Brain Research Institute (KBRI), Daegu, 41062, Republic of Korea; Department of Brain Sciences, Daegu Gyeongbuk Institute of Science & Technology (DGIST), Daegu, 42988, Republic of Korea; School of Energy and Chemical Engineering, Ulsan National Institute of Science and Technology (UNIST), Ulsan 44919, Republic of Korea; Department of Neuropsychiatry, Dongguk University, School of Medicine, Seoul 04620, Republic of Korea; Dongguk University International Hospital, Institute of Clinical Psychopharmacology, Goyang 10326, Republic of Korea; BioMedical Research Center, KAIST, Daejeon, 34051, Republic of Korea; KAIST Stem Cell Center, KAIST, Daejeon, 34141, Republic of Korea

## Abstract

Genetic variations in *CLCN4*, encoding the H^+^/Cl^-^ exchanger CLC-4, are associated with human neurodevelopmental disorders with highly variable phenotypes. A lack of physiologically relevant models has hampered molecular understanding of pathogenic mechanisms. We now establish engineered brain organoid and neuronal cell systems to examine impacts of patient-relevant *CLCN4* genetic variations. We find that *CLCN4* variants reduced excitatory neuron numbers due to early-stage cell death, accompanied by altered endo-lysosomal dynamics and disrupted autophagic flux. Transcriptomic profiling showed significant downregulation of long non-coding RNA *MEG3* in *CLCN4*-variant neurons. Restoring *MEG3* expression is sufficient to rescue cellular defects and improve survival of *CLCN4*-variant neurons. These findings link *CLCN4* dysfunction with impaired autophagy and neuronal cell death, highlighting *MEG3* as a potential therapeutic target for neurodevelopmental disorders involving autophagic dysfunction.

## Introduction

Genetic variations in X-linked gene *CLCN4* have been identified among the individuals with intellectual disabilities and epileptic encephalopathies ^1–7^. Recent extensive studies have revealed that *CLCN4* variants not only contribute to neurodevelopmental and neurological symptoms, but also psychiatric and non-neurological features including gastrointestinal and growth-related defects ^2–4^. However, the molecular mechanisms by which these symptoms in patients arise due to *CLCN4* variations remain unknown.

*CLCN4* encodes CLC-4 protein, a member of the CLC protein family, functions as a chloride ion (Cl^-^) and proton (H^+^) antiporter in intracellular membranes ^8–12^. The CLC Cl-/H+ exchangers are known to cause the influx of Cl^-^ to facilitate the maintenance of vesicular acidic condition ^13,14^. Consistent with other Cl^-^/H^+^ exchangers in the family (CLC-3, 5-7) ^15–18^, CLC-4 is detected in endosomes and lysosomes ^19–22^. In addition to the endo-lysosomal vesicles, CLC-4 is also localized in the endoplasmic reticulum (ER) ^21–23^. In the ER, CLC-4 exists in an unstable homodimer form but forms a stable heterodimer with CLC-3 when it moves to the endo-lysosomal pathway ^21^. Given that CLC-3 is detected in a subset of synaptic vesicles in the mouse brain ^22^, a similar localization pattern can be inferred for CLC-4.

The phenotypic manifestations associated with *CLCN4*-related patients, coupled with its highest expression level in the brain compared to other tissues, underscore the vital roles of CLC-4 in brain function and development. This is further supported by observations of reduced dendritic length and synapse formation in CLC-4-depleted neurons, neurodegeneration in *Clcn4* knockout (KO) mice with a proton-uncoupled CLC-3 background, and autistic-like behaviors in *Clcn4* KO mice ^5,22,24^. Despite these insights, the molecular mechanisms linking CLC-4 to the endo-lysosomal pathway and its impact on neuronal development and degeneration remain poorly defined, largely due to the absence of physiologically relevant disease models incorporating patient-specific *CLCN4* variants.

In this study, we explored the profound effects of *CLCN4* variants on human brain development using brain organoids and neural cells derived from pluripotent stem cells (PSCs), genetically engineered to harbor the variants. We found that *CLCN4* variants lead to premature cell death in excitatory neurons during development. *CLCN4* variants alter vesicle dynamics within the endo-lysosomal pathway and significantly reduce autophagic flux in developing neurons. Crucially, we found that the long non-coding RNA *MEG3* is downregulated in neurons with *CLCN4*-variants, and that restoration of its expression effectively reverses the cell-death phenotypes associated with reduced autophagic flux. These findings enhance our understanding of the roles of CLC-4 in human brain development by elucidating the initial effects of single-gene variations in *CLCN4*. Moreover, our results underscore the potential of targeting *MEG3* as a therapeutic strategy for conditions linked to *CLCN4*-related neurodevelopmental disorders.

## Results

### Reduced number of excitatory neurons in *CLCN4*-variant brain organoids

To explore the effects of *CLCN4* variants on brain development and related diseases, we aimed to establish a model system using hPSCs by applying the CRISPR/Cas9 genome editing technique. We chose to edit the CLC-4 A555, which is predicted to be located in the transmembrane domain, near the chloride-transporting helices. Among the diverse *CLCN4* variants identified in patients, c.1664C>T (p.Ala555Val) is a recurrent *de novo* variation observed in more than seven unrelated families, and it is exclusively identified in females. Three distinct single guide RNAs (sgRNAs) targeting c.1664C were designed and co-transfected into hPSCs with a dCas9-deaminase base editor (Fig. 1a). Each transfection with a different sgRNA resulted in a PSC clone harboring a specific *CLCN4* variant, collectively yielding three PSC lines. Two of these lines are heterozygous (A555L and A556V) and the third line is homozygous (V557I) (Fig. 1b). PSC clones carrying the A555L, A556V, and V557I variants were designated as Variant A, B, and C, respectively.

**Figure 1.**
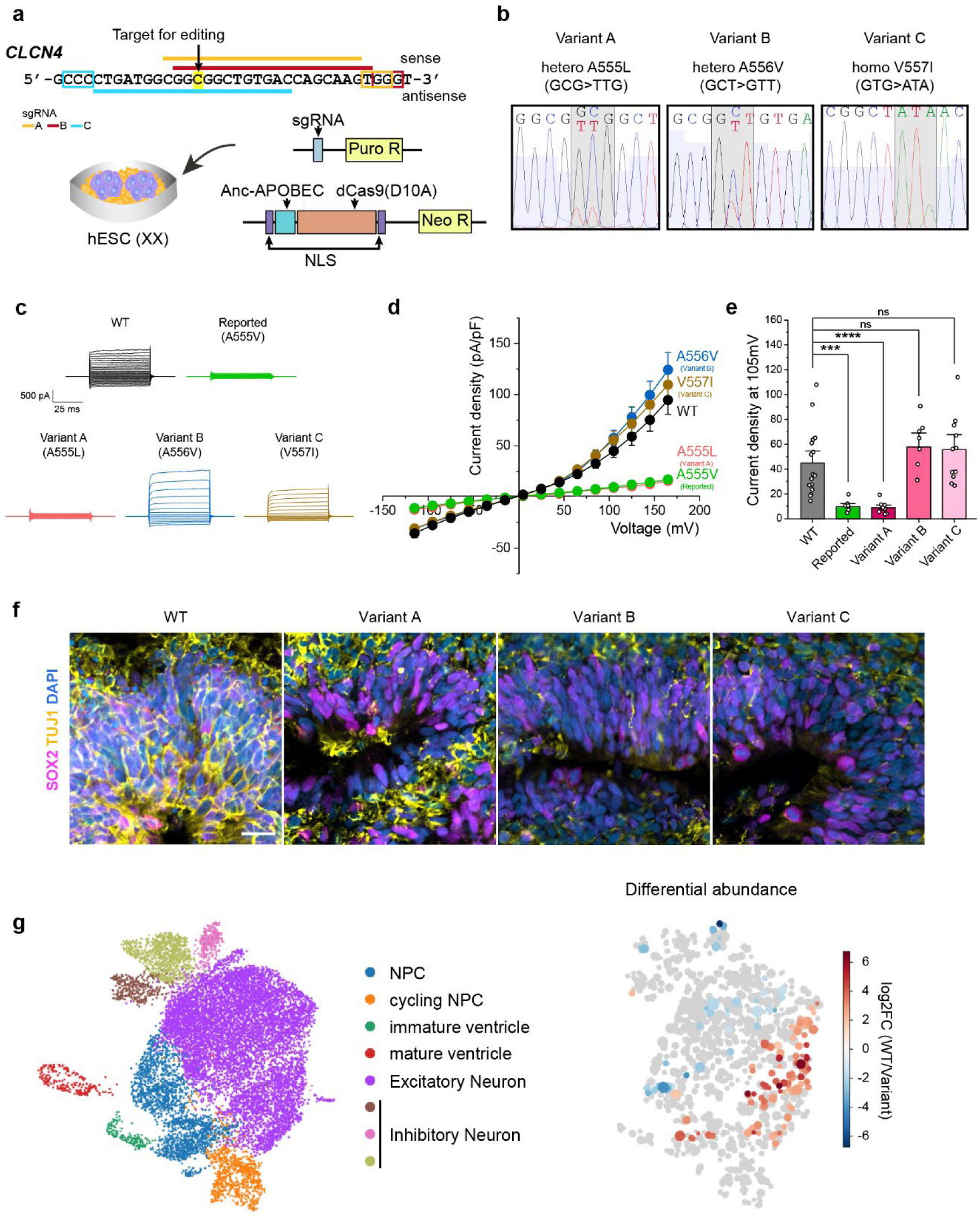
Decrease of neurons in *CLCN4*-variant brain organoids. (a) Scheme of genome editing strategies used in this study. Cytidine deaminase Anc-APOBEC linked to dCas9 with *CLCN4*-targeting sgRNAs was applied to hESCs (HUES6, 46XX). A target nucleotide sequence is highlighted in yellow. PAM sequence for each three sgRNA is marked in color-matched rectangles. (b) Allelic modification in *CLCN4* verified by Sanger sequencing. Each sgRNA results in specific changes: sgRNA_A, CCG to TTG (A555L) in one allele; sgRNA_B, GCT to CTT (A556V) in one allele; and sgRNA_C, GTG to ATA (V557I) in both alleles. The hESC clones generated by each sgRNA were named Variants A, B, and C. (c) Representative whole-cell current traces from HEK293T cells expressing WT (black), reported mutant A555V (green), variant A (A555L, red), variant B (A556V, blue), and variant C (V557I, brown) hCLCN4 channels. Current traces are shown in 20-mV voltage increments for clarity. (d) Current-voltage (I-V) relationships of WT (black), A555V (green), A555L (red), A556V (blue), and V557I (brown) hCLCN4 channels. Current amplitudes were normalized to current densities. Each symbol represents mean ± SEM (WT, n=17; A555V, n=8; A555L, n=10; A556V, n=8; V557I, n=11). (e) Current density at voltage 150 mV. (One-way ANOVA, \*\*\**p*=0.0003019 for A555V, \*\*\*\**p*=0.00008188 for A555L, *p*=0.16391 for A5556, and *p*=0.1923 for A557I) (f) Representative images of day 50 forebrain organoids. SOX2 (magenta) and TUJ1 (yellow) were detected as makers for NPCs and neurons, respectively. Blue represents DAPI. Images are from a single plane of confocal images. Scale bar = 20 μm. (g) scRNA-seq results of day 50 forebrain organoids. The cell clusters are presented based on the gene expression pattern (left) and the differential abundance of cell population (right).

To assess the electrophysiological impact of each *CLCN4* variant on CLC-4 function, each variant was heterologously expressed in HEK293T cells, and ionic currents were measured using whole-cell patch-clamp recordings (Fig. 1c-e). As previously reported, sufficient CLC-4 protein was targeted to the plasma membrane to allow whole-cell recordings, although the majority of CLC-4 resides in intracellular organelles ^25,26^. Although all variants displayed similar expression levels and membrane localization as wild-type (WT), the current of CLC-4 A555L (Variant A) was markedly reduced, similar to the current of A555V variant identified in patients. Variants B and C showed a slight, though not statistically significant, increase in outward rectification. Structural modeling showed that all three variant residues are located on helix Q (hQ), which mediates protomer–protomer interactions at the dimer interface and is connected to helix R (hR), where the inner gate-forming residue Y572 resides (Supplementary Fig. 1b). Although the overall electrostatic properties of the variants were predicted to be similar to WT, substitutions at A555 (to valine or leucine) are likely to introduce steric clashes with neighboring residues, including F335, A523, and P551. Such physical clashes may destabilize dimer interactions or disrupt inner gate regulation, ultimately impairing the electrophysiological function of CLC-4 (Supplementary Fig. 1c). However, the A556V and V557I substitutions may be accommodated within the surrounding space without generating steric hindrance.

Next, we investigated the impact of CLCN4 variants on brain development. Given that affected individuals display early-onset symptoms and cortical atrophy, prominent indicators of disrupted early neurodevelopment ^2,6,26,27^, we differentiated the *CLCN4*-variant PSCs into forebrain organoids to model early cortical development. After 50 days of differentiation, the forebrain organoids were immunostained with TUJ1 antibody to determine the presence of neurons within the organoids. We observed a drastic decrease in TUJ1-positive cells in all three *CLCN4* variant organoids compared to the WT (Fig. 1f), regardless of electrophysiological characteristics of each variant. No significant differences were found in the distribution of neural progenitor cells (NPCs), identified as SOX2-positive cells.

Single-cell RNA sequencing (scRNA-seq) revealed that brain organoids were mainly composed of NPCs, excitatory, and inhibitory neurons, with fewer excitatory neurons in *CLCN4*-variant organoids (Fig. 1g). Pseudotime trajectory analysis based on gene expression showed high NPC marker expression at early stages that declined over time as neuron-specific gene expression increased (Supplementary Fig. 2). *CLCN4*-variant organoids exhibited reduced cell density at early pseudotime points, suggesting that *CLCN4* variations impaired neuronal development during these initial stages.

### *CLCN4* variant-induced cell death during early neurogenesis

To determine whether *CLCN4* variants impair the initiation of neuronal development or induce neuronal cell death, we utilized a two-dimensional cell culture protocol to generate NPCs from PSCs, which primarily differentiate into excitatory neurons ^28^. *CLCN4* mRNA levels were elevated in NPCs relative to PSCs and further increased upon neuronal differentiation (Fig. 2a), indicating a pronounced impact of *CLCN4* on excitatory neuronal development. This aligns with the reduced excitatory neuron population observed in *CLCN4*-variant brain organoids. In contrast, *CLCN4* expression remained low in astrocytes, suggesting a minimal contribution in astrocytes.

**Figure 2.**
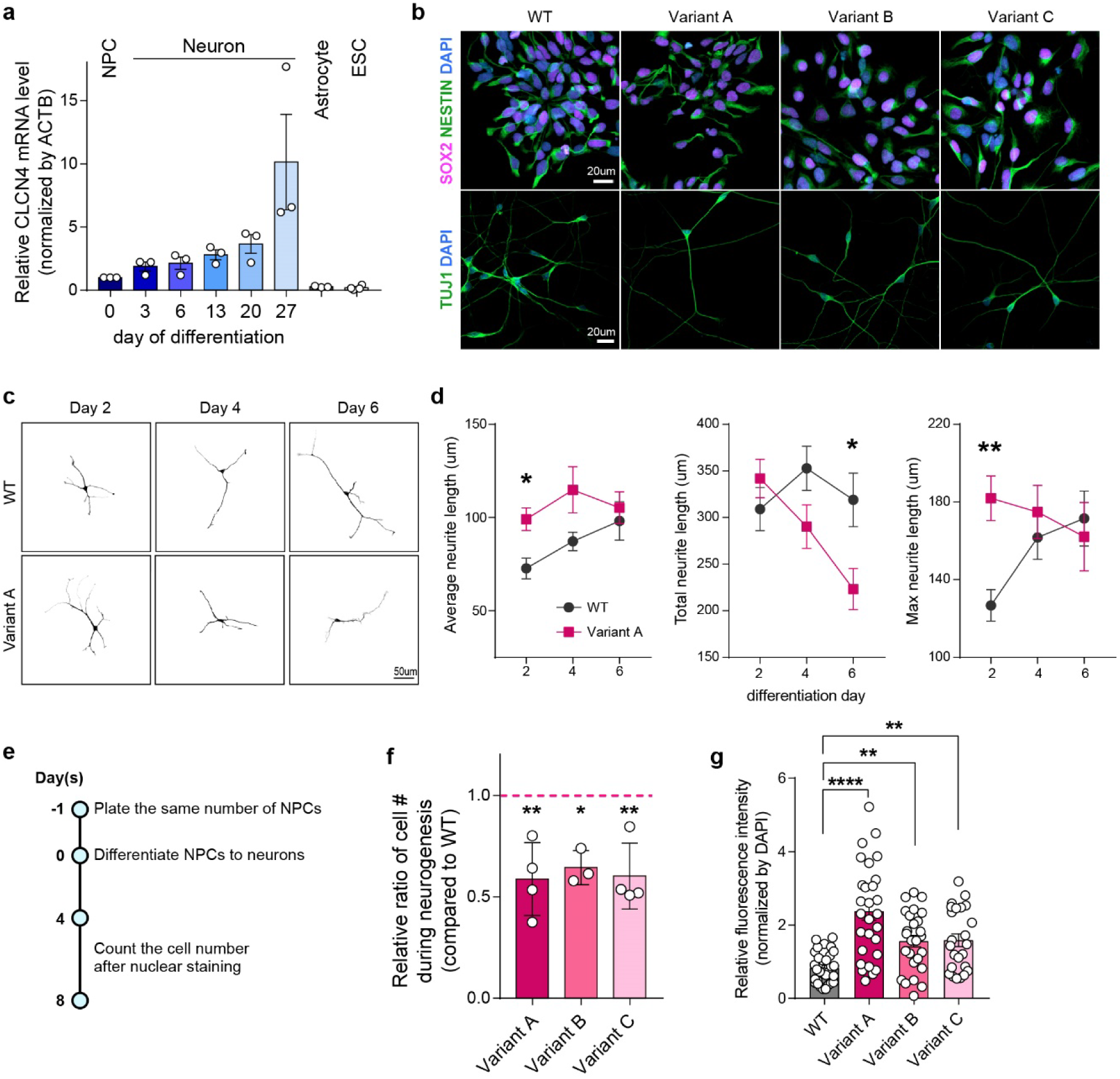
Aberrant neuronal cell death in early stage of neurogenesis. (a) Relative expression levels of CLCN4 mRNA during neurogenesis and in non-neuronal cells, normalized to ACTB expression. (n=3 or 4, mean ± SEM, one-way ANOVA, \**p*=0.0172) (b) NPCs (top) and day 4 neurons (bottom) derived from WT and *CLCN4*-variant hESCs. NPCs were stained with SOX2 (magenta) and NESTIN (green), neurons with TUJ1 (green). DAPI is shown in blue. Scale bar = 20 μm. (c) Representative images of developing neurons used for neurite length analysis. Scale bar = 50 μm. (d) Quantification of average (left), total (middle), and maximum (right) neurite length. Average neurite length was calculated as total neurite length divided by the number of branches. The following numbers of neurons from three or more biological replicates were analyzed: day 2 (WT = 35; variant A = 38), day 4 (WT = 26; variant A = 25), and day 6 (WT = 28; variant A = 25). (mean ± SEM, two-way ANOVA with Sidak’s multiple comparisons test; \**p*=0.0288 at average; \**p*=0.0228 at total; \*\**p*=0.0021 at maximum) (e) Schematic timeline for quantifying changes in neuronal cell numbers. (f) Quantification of relative neuronal cell numbers during neurogenesis. (n = 3 or 4, mean ± SEM, one-way ANOVA with Dunnett’s multiple comparisons test; Variant A, *\*\*p=0.0076*; Variant B, *\*p=0.0259*; Variant C, *\*\*p=0.0095*). The magenta dashed line indicates the WT value. Each dot represents the relative ratio between the variant and WT from one biological replicate. (g) Relative TUNEL intensity in day 6 neurons. (n=3, mean ± SEM, one-way ANOVA with Dunnett’s multiple comparisons test; \*\*\*\**p*<0.0001 for variant A, \*\**p*=0.0054 for variant B, and \*\**p*=0.0048 for variant C)

The *CLCN4* variants did not affect the differentiation of PSCs into NPCs, as both WT and *CLCN4* variant cells consistently expressed the NPC markers, SOX2 and NESTIN, with no detectable differences among groups (Fig. 2b). When cultured under differentiation conditions for four days, NPCs developed into TUJ1-positive neurons, indicating successful initiation of neuronal differentiation

However, developing neurons with *CLCN4* variants displayed aberrant morphogenesis (Fig. 2c, d). At the early stage of neuronal development (day 2), total neurite length was comparable between WT and *CLCN4*-variant neurons, but the average neurite length was significantly longer in *CLCN4*-variant neurons, indicating fewer but more elongated neurites. In addition, maximum neurite (the longest neurite) length was substantially longer in *CLCN4*-variant neurons than in WT neurons. Upon further differentiation, WT neurons extended both average and maximum neurite lengths with minimal changes in total neurite length, suggesting pruning of shorter neurites and extension of selected neurites. In contrast, *CLCN4*-variant neurons showed a progressive decline in total and maximum neurite lengths, with total neurite lengths significantly reduced by day 6.

During continuous neuronal differentiation, we observed an unexpected reduction in the number of *CLCN4*-variant neurons compared to the control (Fig. 2e-g). As neurons are post-mitotic cells, this reduction likely reflects increased cell death. We compared the number of cells on days 4 and 8 of differentiation and found that the relative number of remaining *CLCN4*-variant neurons on day 8 compared to day 4 was about 50% that of the control, indicating a significant reduction in the viability of *CLCN4* variant neurons (Fig. 2f). TUNEL assay, which detects DNA fragments during programmed cell death, showed increased signals in *CLCN4*-variant neurons (Fig. 2g), confirming elevated neuronal cell death associated with *CLCN4* variants.

### Endo-lysosomal and autophagic defects in *CLCN4*-variant neurons

To understand the mechanisms of *CLCN4* variant-induced neuronal death, we investigated changes in intracellular vesicles within the endo-lysosomal pathway before the onset of neuronal cell death. As CLC-4 localizes to vesicle membranes within this pathway, it is expected to play a crucial role in endo-lysosomal dynamics ^19,21–23,29,30^. LysoTracker staining, which detects acidic organelles such as endosomes and lysosomes, showed an increased signal in *CLCN4*-variant neurons (Fig. 3a, b). However, LysoSensor, a pH indicator for vesicles, revealed no significant change in vesicular acidity (Fig. 3c).

**Figure 3.**
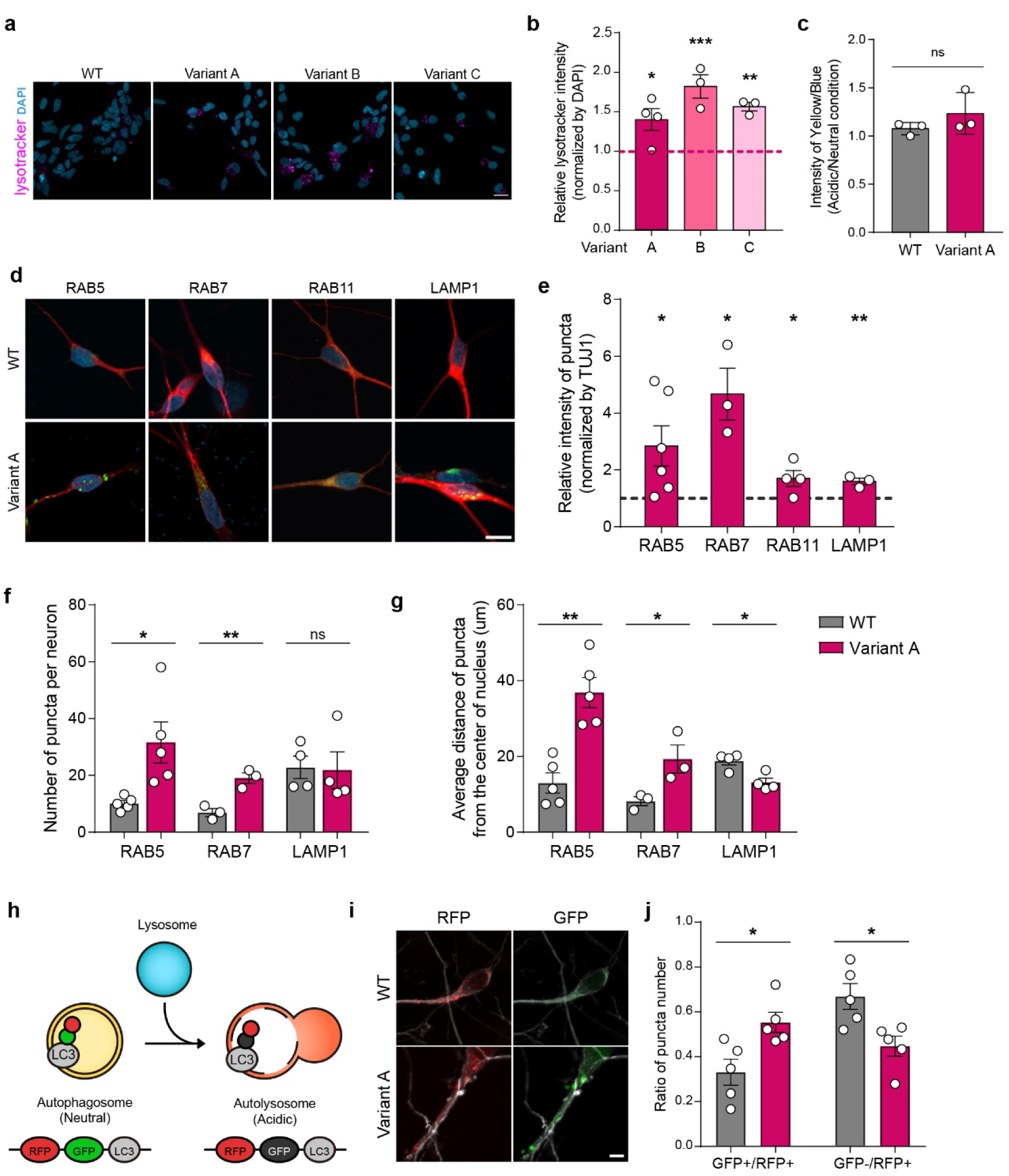
Disrupted endo-lysosomal pathway in *CLCN4*-variant neurons. (a) Representative images of day 6 neurons after Lysotracker treatment. Lysotracker is shown in magenta, and DAPI in blue. Scale bar = 20 μm. (b) Quantification of Lysotracker signal in day 6 neurons. The relative Lysotracker intensity of *CLCN4*-variant neurons compared to the WT was presented. Each dot represents one biological replicate. (n=3-4, mean ± SEM, one-way ANOVA with Dunnett’s multiple comparisons test \**p*=0.039 for variant A; \*\*\**p*=0.0007 for variant B; \*\**p*=0.0092 for variant C) The magenta dashed line indicates the value for WT. (c) Relative acidity in acidic subcellular organelles. Each dot represents one biological replicate. (n=3, mean ± SEM, unpaired Student’s t-test; *p*=0.2894) (d-g) Dynamics of endo-lysosomal vesicles. RAB5, RAB7, RAB11 and LAMP1 indicates early endosome, late endosome, recycling endosome and lysosome markers, respectively. (RAB5, n=6; RAB7, n=3; RAB11, n=4; and LAMP1, n=3 or 4; mean ± SEM, unpaired Student’s t-test) (d) Representative images of a neuron with GFP-labeled vesicular organelles. TUJ1 and DAPI are shown in red and blue, respectively. Scale bar = 10 μm. (e) Relative GFP intensity expressed in vesicular organelles (\**p*=0.0265 for RAB5, \**p*=0.0158 for RAB7, \**p*=0.0475 for RAB11, and \*\**p*=0.0067 for LAMP1) (f) Number of GFP-expressing vesicles in neurons. (\**p*=0.0186 for RAB5, \**\*p*=0.0066 for RAB7, and *p*=0.9031 for LAMP1) (g) Average distance of GFP-expressing vesicles from the center of nucleus, representing the mean linear distance of puncta from the center of the nucleus in each cell. (\*\**p*=0.0011 for RAB5, \**p*=0.0463 for RAB7, and \**p*=0.0103 for LAMP1) (h) Scheme of autophagic flux analysis using tandem fluorescent protein fused to LC3. (i) Representative images of day 6 neurons expressing tandem fluorescent-tagged LC3. Red: RFP, green: GFP, white: TUJ1. Scale bar = 5 μm. (j) Quantification of the GFP-/RFP+ puncta. (n=5, mean ± SEM, unpaired Student’s t-test; \**p*=0.0164)

We further examined the impact of *CLCN4* variants on specific subpopulations of acidic vesicles within the endo-lysosomal pathway. To visualize these organelles, neurons were infected with lentiviruses expressing fluorescent protein-tagged markers: RAB5 for early endosomes, RAB7 for late endosomes, RAB11 for recycling endosomes, and LAMP1 for lysosomes (Fig. 3d). Consistent with Lysotracker staining results, fluorescence intensity for all markers increased in *CLCN4*-variant neurons (Fig. 3e). Notably, RAB5- and RAB7-positive puncta counts were significantly elevated, while quantification of RAB11-positive puncta was complicated by their irregular morphology (Fig. 3f). The increased number of endosomal puncta corresponds well with the elevated fluorescence intensity observed in *CLCN4*-variant neurons relative to WT neurons. Interestingly, despite the increased LAMP1 signal intensity, the number of LAMP1-positive puncta remained unchanged in *CLCN4*-variant neurons, suggesting that lysosomes likely increase in size rather than number (Fig. 3e, f). In neurons, endosomes originate near the plasma membrane and migrate toward the nucleus, while lysosomes form near the nucleus and move outward ^31^. Acidic vesicles showed abnormal positioning in *CLCN4*-variant neurons (Fig. 3e). To assess vesicle localization, we measured their distance from the nuclear center. In *CLCN4*-variant neurons, RAB5- and RAB7-positive vesicles were more peripheral, while LAMP1-positive puncta were closer to the nucleus, suggesting that the CLC-4 variants likely disrupt endo-lysosomal trafficking dynamics.

Lysosomes degrade cargo molecules by fusing with autophagosomes to form acidic autolysosomes. To evaluate whether *CLCN4* variants interfere autophagic flux, we employed an LC3 reporter, tagged with RFP and GFP in tandem, which localizes to both autophagosomes and autolysosomes. In acidic environments like autolysosomes, the GFP signal is quenched while RFP remains stable, allowing distinction between autophagosomes and autolysosome (Fig. 3h). While the total number of RFP-positive puncta per neuron was similar between groups, the number of GFP-positive puncta differed between WT and *CLCN4*-variant neurons (Fig. 3i, j). In WT neurons, approximately 70% of RFP-positive LC3 puncta were GFP-negative, whereas in *CLCN4*-variant neurons, only about 45% were GFP-negative, indicating impaired autolysosome formation.

To explore potential treatments for rescuing *CLCN4*-variant neurons from early cell death, we tested autophagy-promoting compounds, including rapamycin, LiCl, and calpain inhibitor I (Supplementary Fig. 3a) ^32–34^. Despite their known effects on autophagy, none of these agents prevented cell death in *CLCN4*-variant neurons. Similarly, neurodevelopmental disorder medications such as risperidone and clozapine failed to restore neuronal viability, highlighting the complexity of *CLCN4* variant-induced neuronal vulnerability and the need for targeted therapeutic strategies (Supplementary Fig. 3b).

### *MEG3* downregulation in *CLCN4*-variant neurons

To identify early molecular changes preceding neuronal cell death and uncover potential therapeutic targets in *CLCN4*-variant neurons, we performed total RNA sequencing (total RNA-seq) on Variant A and WT neurons at differentiation days 2 and 6 (Fig. 4a, Supplementary Fig. 4a). Total RNA-seq revealed reduced *CLCN4* mRNA levels in variant A neurons (Fig. 4b), which was further validated in neurons carrying other variants using quantitative PCR (qPCR) (Fig. 4c). At both time points, extensive transcriptional changes were observed, with 956 and 1,184 differentially expressed genes (DEGs, |log2FC| ≥ 1, padj < 0.05) identified, respectively. Gene ontology (GO) analysis indicated that DEGs are enriched in neurodevelopmental processes (Supplementary Fig. 4b), suggesting that *CLCN4* variation impairs neurodevelopment prior to the onset of cell death.

**Figure 4.**
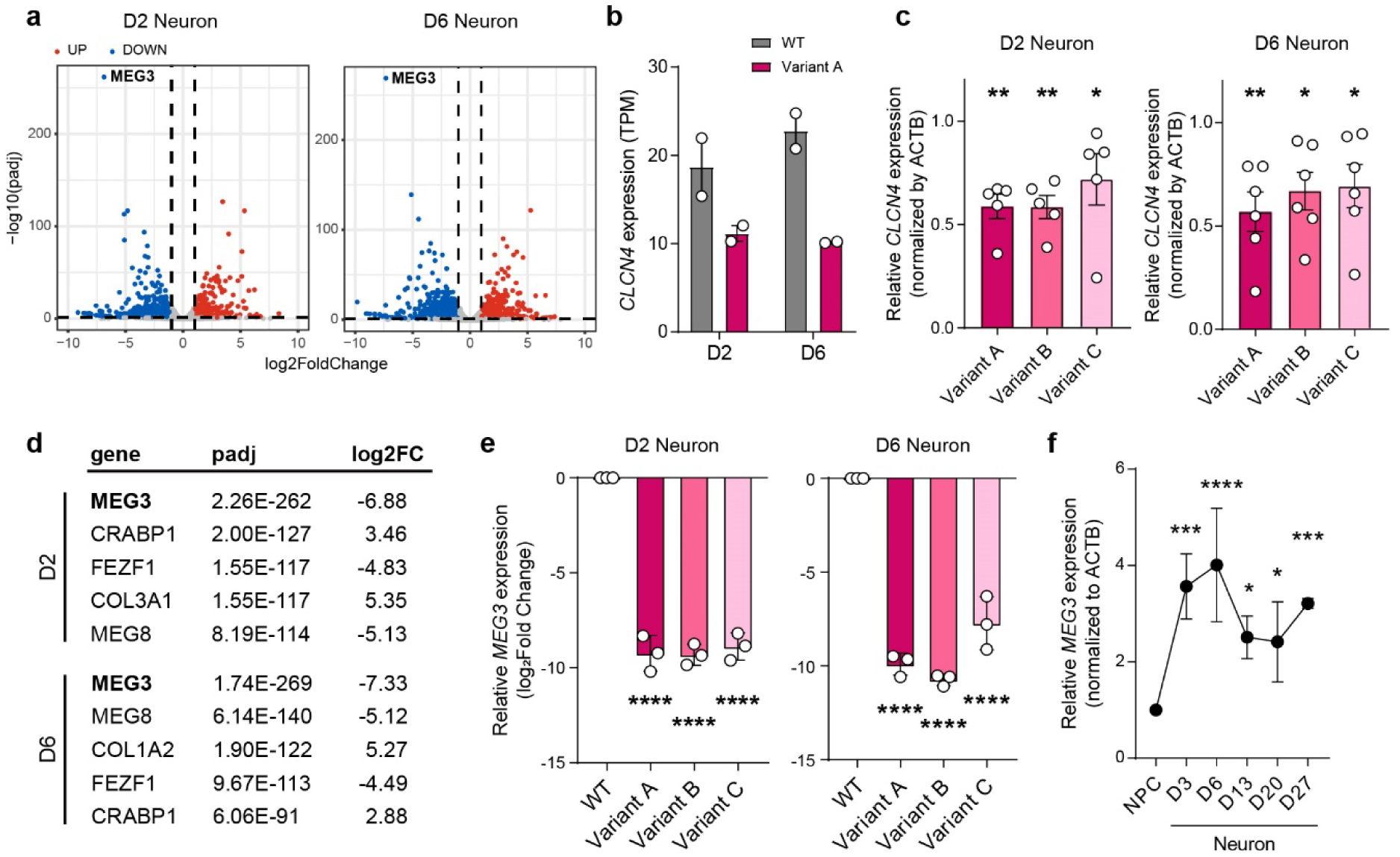
RNA expression profiling in *CLCN4*-variant neurons. (a) Volcano plots in day 2 and day 6 neurons total RNA seq. Horizontal dashed lines: threshold of adjust p-value, 0.05; dashed lines: threshold of |log2FoldChange|, 1. Blue and red dots indicate significantly down- and up-regulated genes, respectively. (b) TPM of CLCN4 in total RNA seq. (c) The confirmed expression of CLCN4 in *CLCN4*-variant day 2 (left) and day 6 (right) neurons by real-time PCR. (n=5, mean ± SEM, one-way ANOVA with Dunnett’s multiple comparisons test, D2 neuron; \*\**p*=0.0033 for variant A, \*\**p*=0.0031 for variant B, and \**p*=0.0424 for variant C, D6 neuron; \*\**p*=0.0047 for variant A, \**p*=0.0301 for variant B, and \**p*=0.0462 for variant C) (d) Top 5 significant gene list from total RNA seq. (e) The confirmed expression of MEG3 in *CLCN4*-variant day 2 (left) and day 6 (right) neurons by real-time PCR. (n=3, mean ± SEM, one-way ANOVA with Dunnett’s multiple comparisons test, \*\*\*\**p* < 0.0001) (f) Changes in expression of MEG3 during neurogenesis. (n=3, mean ± SEM, two-way ANOVA with Dunnett’s multiple comparisons test; \*\*\**p*=0.0003 at D3, \*\*\*\**p* < 0.0001 at D6, \**p*=0.0114 at D13, \**p*=0.0169 at D20, and \*\*\**p*=0.0008 at D27)

Among the DEGs, *MEG3* was the most significantly downregulated gene at both time points (Fig. 4d). *MEG3* is a long non-coding RNA (lncRNA) known to produce multiple transcript isoforms. To determine whether different *MEG3* isoforms show distinct expression patterns, we analyzed their expression at the isoform level (Supplementary Fig. 4c). The top 10 most abundant isoforms, accounting for over 95% of total *MEG3* expression, were uniformly reduced in *CLCN4*-variant neurons. qPCR using primers targeting the majority of *MEG3* isoforms, including both the canonical and most abundant isoforms, confirmed reduced *MEG3* expression across all three *CLCN4*-variant neurons (Fig. 4e). Notably, *MEG3* expression increased dramatically during the early stages of neuronal differentiation from NPCs (Fig. 4f), suggesting its potential role in neurodevelopment, particularly in the NPC-to-neuron transition.

### *MEG3* rescues the phenotypes of *CLCN4*-variant neurons

To investigate whether reduced *MEG3* contributes to neuronal death in *CLCN4*-variant neurons, we suppressed *MEG3* expression in WT NPCs using a lentiviral system expressing dCas9-KRAB and sgRNAs targeting the *MEG3* promoter (sgMEG3-i) and examined its effect on cell death (Fig. 5a). *MEG3* expression was successfully reduced by approximately 50% in transduced WT NPCs (Fig. 5b), which were subsequently differentiated into neurons. Upon differentiation, the number of surviving neurons on day 8 relative to day 4 was reduced by 50% in the *MEG3*-suppressed group compared to the control, indicating that *MEG3* downregulation promotes neuronal cell death (Fig. 5c).

**Figure 5.**
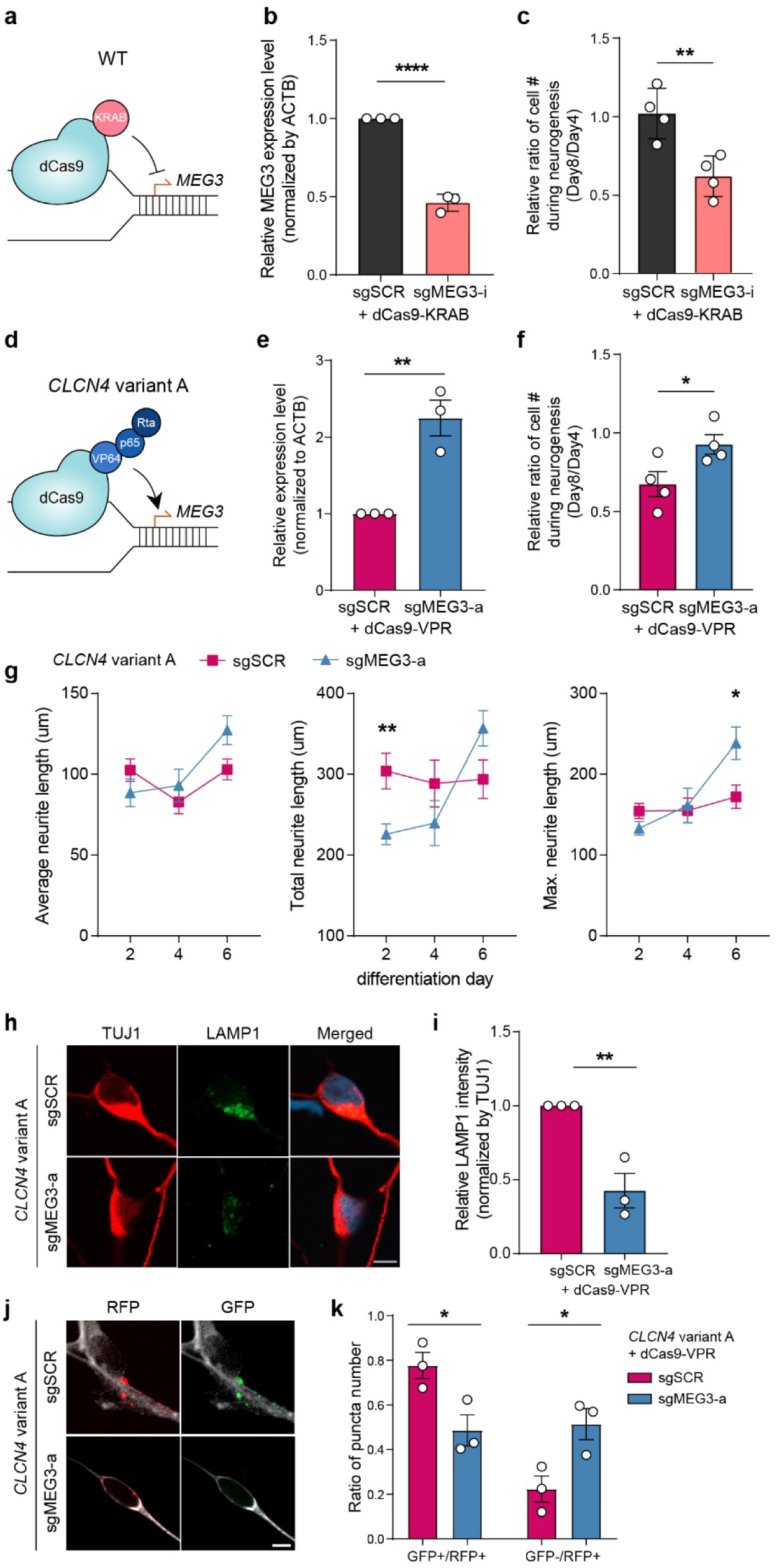
*MEG3* in restoring phenotypes in *CLCN4* variant neurons. (a) Diagram of CRISPR/Cas9-based *MEG3* inhibition system applied to WT NPC and neuron. (b) Relative expression level of *MEG3* in WT NPCs after *MEG3* inhibition. (n=3, mean ± SEM, unpaired Student’s t-test, \*\*\*\**p*<0.0001) (c) Quantification of cell number ratio in WT neurons during neurodevelopment after *MEG3* inhibition. (n=4, mean ± SEM, unpaired Student’s t-test; \*\**p*=0.0083) (d) Diagram of CRISPR/Cas9-based *MEG3* activation system applied in *CLCN4*-variant A cells. (e) Relative expression of *MEG3* in *CLCN4*-variant NPCs after *MEG3* activation. (n=3, mean ± SEM, unpaired Student’s t-test, \*\**p*=0.0058) (f) Changes in cell number was compared in *CLCN4*-variant neurons after *MEG3* activation. (n=4, mean ± SEM, unpaired Student’s t-test, \**p*=0.0475) (g) Quantification of average (left), total (middle), and maximum (right) neurite length in *CLCN4*-variant neurons. The following numbers of neurons were analyzed: day 2 (sgSCR = 34; sgMEG3-a = 36), day 4 (sgSCR = 11; sgMEG3-a = 10), and day 6 (sgSCR = 32; sgMEG3-a = 37). (n=3, mean ± SEM, two-way ANOVA with Sidak’s multiple comparisons test; \**p*=0.0372 at total; \*\**p*=0.0028 at maximum) (h) Representative images of LAMP1 in day 6 neuron. LAMP1 (green), TUJ1 (red), and DAPI (blue) were shown. Scale bar = 5 μm. (i) Relative GFP intensity expressed in LAMP1-positive vesicles. (n=3, mean ± SEM, unpaired Student’s t-test, \*\**p*=0.0078) (j) Representative images of day 6 neurons expressing tandem fluorescent-tagged LC3. Red: RFP, green: GFP, white: TUJ1. Scale bar = 5 μm. (k) Quantification of the GFP-/RFP+ puncta in CLCN4-variant neurons after MEG3 activation. (n=3, mean ± SEM, unpaired Student’s t-test, \**p*=0.0327)

Next, to determine whether restoring *MEG3* expression could rescue the phenotypes observed in *CLCN4*-variant neurons, we overexpressed *MEG3* using a lentiviral system encoding dCas9-VPR and a sgRNA targeting the *MEG3* promoter (sgMEG3-a) (Fig. 5d, e). *MEG3* overexpression in *CLCN4*-variant neurons significantly improved neuronal survival compared to controls expressing a non-targeting sgRNA (Fig. 5f). It also rescued abnormal neurite outgrowth, restoring neurite length during early neurodevelopment (Fig. 5g). Furthermore, *MEG3* activation reduced the elevated levels of LAMP1-positive vesicles (Fig. 5h, i) and restored the impaired autophagic flux. (Fig. 5j, k). These findings demonstrate that modulating *MEG3* expression can reverse neurodevelopmental deficits in *CLCN4*-variant neurons and highlight *MEG3* as a potential therapeutic target for autophagy-related neurodevelopmental disorders.

## Discussion

In this study, we investigated how single amino acid alterations in CLC-4 affect human brain development using PSC-derived brain organoids and neurons. We found that *CLCN4* variants disrupt endo-lysosomal dynamics and impair autophagic flux, ultimately leading to premature neuronal cell death. Through transcriptomic analysis, we identified *MEG3* as a critical regulator that counteracts the deleterious effects of *CLCN4* variants. Notably, *MEG3* inhibition recapitulated the early neuronal cell death observed in *CLCN4*-variant neurons, whereas *MEG3* upregulation effectively restored autophagic flux and improved neuronal survival, suggesting that *MEG3* may serve as a potential therapeutic target for *CLCN4*-associated neurodevelopmental disorders.

Patch-clamp recordings revealed a dramatic current reduction in variant A, but not in variants B or C (Fig. 1c-e). Based on this, we expected variant A neurons to display distinct cellular phenotypes, potentially due to impaired vesicle acidification, as observed in CLC-4 null fibroblasts with elevated endosomal pH ^35^. However, lysosensor-based pH measurements revealed no detectable differences in vesicular acidity between *CLCN4* variant A and WT neurons (Fig. 3c), suggesting vesicular acidity remains largely unaffected. While subtle changes in luminal pH cannot be entirely excluded, it is notable that disruption of other chloride/proton exchangers often does not affect vesicle pH ^36^. Supporting this, all three *CLCN4* variants presented similar cellular phenotypes, including neuronal death, despite their distinct electrophysiological properties. These results indicate that altered chloride conductance or vesicle acidification is unlikely to be the primary cause of the observed cellular phenotypes in *CLCN4*-variant neurons. Given the marked alterations in endosome and lysosome levels and distribution in *CLCN4*-variant neurons (Fig. 3a-g), further investigation is required to explore alternative mechanisms, such as changes in other ion concentrations affecting vesicular osmosis ^37,38^.

Lysosomes are normally distributed throughout the cytoplasm, but their positioning is tightly regulated. Perinuclear lysosomes are more acidic and actively involved in autophagosome fusion, whereas distal lysosomes tend to be less acidic ^39^. In *CLCN4*-variant neurons, lysosomes were redistributed toward the perinuclear region, despite no detectable changes in luminal pH (Fig. 3g), suggesting that lysosomal repositioning occurs independently of acidification status. Inhibition in distal transport of lysosomes induces abnormal accumulation of autophagosomes and aggregates ^40^. This accumulation may reflect impaired fusion of lysosomes with autophagosomes during autophagy. Supporting this, *CLCN4*-variant neurons displayed reduced autophagic flux, as evidenced by defective autolysosome formation (Fig. 3h-j). Autolysosomes emerge during the neuronal differentiation of neural stem cells through the fusion of lysosomes and autophagosomes ^41,42^ and play essential roles in neurite outgrowth, pruning, and synaptogenesis. In addition to these developmental roles, autophagy protects neurons by clearing toxic cellular waste, including misfolded proteins and aggregates, thereby preventing degeneration and cell death ^43–45^. Consistent with these functions, *CLCN4*-variant neurons with impaired autolysosome formation display aberrant neurite growth during early differentiation and ultimately undergo premature cell death (Fig. 2c-g). Notably, similar defects in lysosomal positioning and autophagy have been reported in models of Alzheimer’s, Huntington’s and Parkinson’s diseases ^46–48^ highlighting that perinuclear lysosome accumulation and impaired autophagic flux as pathological hallmarks commonly associated with neurodegeneration.

Autophagy-related genes are typically upregulated during neuronal differentiation, and their dysregulation has been implicated in a range of neurodevelopmental disorders ^49,50^. However, in *CLCN4*-variant neurons, the expression of canonical autophagy genes, including ATG family genes, remained unchanged (Supplementary Table 1), despite significant alterations in genes associated with neurodevelopment-related GO terms (Supplementary Fig. 4c). Instead, we identified *MEG3* as a key regulator of autophagy in neurons harboring *CLCN4*-variants (Fig. 4). Activating *MEG3* in *CLCN4*-variant neurons restored autophagic flux and rescued premature cell death (Fig. 5f, k), highlighting its essential role in maintaining autophagic function during early neuronal differentiation. Although *MEG3* has been primarily studied in cancers ^51,52^, it is known to regulate autophagy in a context-dependent manner, acting as either an enhancer or inhibitor ^51,53,54^. While the precise mechanisms by which *MEG3* modulates autophagy in developing neurons remain unclear, previous studies have shown that *MEG3* can regulate autophagy-related genes such as GRB2, PTEN and ATG3 at the post-transcriptional level, suggesting it may influence autophagy indirectly through downstream targets ^51,53,55^. Investigating microRNA networks involved in post-transcriptional gene regulation, along with proteomic alterations and post-translational modifications in *CLCN4*-variant or *MEG3*-modulated neurons, may offer important insights into how *MEG3* contributes to impaired autophagy early neurodegeneration ^56^.

A recent study reported that *MEG3* contributes to neurodegeneration in mature neurons ^57^. Specifically, *MEG3* was upregulated exclusively in human neurons modeling Alzheimer’s disease, where it induced necroptosis, whereas such induction was not observed in mouse neurons. This effect of *MEG3* on neuronal death contrasts with our findings, which show that *MEG3* suppression in developing neurons induces premature cell death (Fig. 5c). Considering the rapid upregulation of *MEG3* during early neuronal differentiation and its continued expression in mature neurons (Fig. 4f), these findings suggest that disruption of its normal expression pattern may lead to neuronal cell death. Interestingly, *MEG3* expression did not change in the mouse NPCs following *Clcn4* knockdown ^24^. Although the effects of knockdown may not fully recapitulate those of disease-associated variants, it is noteworthy that *MEG3* downregulation occurred exclusively in human NPCs and neurons (Fig. 4e, Supplementary Fig. 5). This suggests that *MEG3* may be involved in human-specific regulatory pathways governing neuronal cell death. Understanding the mechanisms that regulate *MEG3* expression may provide broader insights into the pathways underlying neuronal degeneration and offer potential avenues for therapeutic intervention in neurodegenerative diseases. Although the mechanisms by which *CLCN4* variants regulate *MEG3* expression remain unknown, the consistent downregulation of all *MEG3* isoforms in *CLCN4*-variant neurons (Supplementary Fig. 4b) suggests the potential involvement of transcriptional or epigenetic regulation.

To date, over 70 distinct *CLCN4* variants have been identified across the gene, and the clinical spectrum of *CLCN4*-related disorders ranges from asymptomatic presentation to severe neurodevelopmental disabilities. Functional studies in *Xenopus* oocytes have also shown that *CLCN4* variants display diverse electrophysiological phenotypes ^2,3,26^. This genetic and phenotypic heterogeneity suggests that individual variants may differentially affect neurodevelopmental processes. In this study, we focused on the *CLCN4* A555V variant by introducing the structurally similar A555L substitution into human ESCs and identified autophagy-related neuronal cell death as a key phenotype. However, whether this phenotype is shared across other disease-associated variants remains to be determined. Conventional knockout or knockdown models often fail to recapitulate the precise molecular disturbances caused by patient-specific point mutations ^58–60^, highlighting the need for more accurate and clinically relevant disease models. Notably, even among individuals carrying the same variant, clinical variability can arise depending on genetic background ^61–63^, underscoring the importance of using patient-derived iPSC models to investigate both shared and variant-specific mechanisms influenced by genetic background. Such models will be critical for determining whether the defects in the *CLCN4*-*MEG3*-autophagy axis observed in this study are generalizable across different variants and genetic backgrounds, and for uncovering additional phenotypes associated with *CLCN4* variations. Furthermore, elucidating the *CLCN4*-*MEG3*-autophagy axis may not only enhance our understanding of *CLCN4*-related neurodevelopmental disorders but also provide broader insights into autophagy-dependent mechanisms underlying other forms of neurodevelopmental pathology.

## Methods

### Cell culture

Mouse embryonic fibroblasts (MEFs) were isolated from E13.5 pregnant CF-1 mice embryos (KRIBB) and cultured in MEF media comprising DMEM (Welgene, LM001-07), 15% fetal bovine serum (Gibco, 12483-020), 2-mercaptoethanol. HUES6 obtained from HSCI iPS Core was maintained on the mitomycin c-treated MEF in human embryonic stem cell (hESC) medium containing DMEM/F-12 (Gibco, 12400-024), 20% Serum Replacement (Gibco, 10828028), and 10 ng/ml bFGF (R&D systems, 4114-TC-01M). hESCs were passed using Collagenase type IV (Gibco, 17104019) in following a previously reported protocol. The generation and usage of MEF and hESC was approved in accordance with the Institutional Animal Care and Use Committees of KAIST (KA2020-37) and the ethical requirements and regulations of the Institutional Review Board of KAIST (KH2021-069).

### Plasmid constructs for *CLCN4* editing

A neomycin resistance gene (NeoR) from pcDNA3.1(+) (Invitrogen, V79020) was subcloned into pCMV_AncBE4max (Addgene plasmid #112094) to produce pCMV-AncBE4max-NeoR. Three sgRNAs targeting CLCN4 were designed using the Broad Institute GPP sgRNA designer. Complementary oligos for sgRNA constructs were synthesized by Macrogen. Oligos were annealed by decreasing temperature from 95℃ to room temperature (RT) and subcloned into lentiGuide-puro (Addgene plasmid #52963). The sequence of generated plasmids was confirmed by Sanger sequencing. The oligos for sgRNA were as follows: sgCLCN4-01: 5’- CACCGGGCGGCTGTGACCAGCAAGT-3’ (forward) and 5’- AAACACTTGT GGTCACAGCCGCCC-3’ (reverse); sgCLCN4-02: 5’- CACCGGTCACAGCCGCCGCCATCA G-3’ (forward) and 5’- AAACCTGATGGCGGCGGCTGTGACC-3’ (reverse); sgCLCN4-03: 5’- CACCGCGGCGGCTGTGACCAGCAAG-3’ (forward) and 5’- AAACCTTGCTGGTCACAGCC GCCGC-3’ (reverse).

### Generation of *CLCN4*-variant hPSCs

HUES6 ESCs were maintained on Matrigel (Corning, 354230)–coated dishes using mTeSR medium (Stem cell technologies, ST85850) with 10 ng/ml bFGF. A day before transfection, ESCs were dissociated with Accutase (Innovative cell technologies, AT104) and 75,000 cells were seeded into a single well of a Matrigel-coated 4-well plate. Transfection was performed using lipofectamine stem transfection reagent (ThermoFisher, STEM00003) with total 1 ug of plasmids. lentiGuide-puro with sgRNA and pCMV_AncBE4max-NeoR were transfected with 1:1 molecular ratio. Twenty-four hours post-transfection, 1 ug/ml of puromycin (Gibco, A1113803) and 250 ug/ml of G418 (Gibco, 10131035) were added to the culture medium to select for cells successfully transfected with both plasmids. Puromycin and G418 were treated for 24 hours and 3 days, respectively. To verify *CLCN4* mutation, genomic DNA was extracted using G-DEX IIc Genomic DNA Extraction Kit (Intron biotechnology, 17231) and the target region was amplified by PCR using primers (5’-GCCCAGTGTAGGAAGCATGT -3’ and 5’-AAAGGATGCCATTCGGCTCT-3’) and the PCR products were sequenced by Sanger sequencing. PCR products were subcloned into pGEM-T easy (Promega, A1360) and total 10 clones were sequenced. From the ratio of the edited allele and wild type allele, homozygote and heterozygote mutations were determined.

### Whole-cell patch-clamp recording

For electrophysiological recording of hCLC-4 channel currents, the WT or mutant open reading frame (ORF) of human *CLCN4* (NCBI accession no. NM001830) was cloned into the enhanced episome vector (EEV600A-1; Systems Biosciences) with an in-frame C-terminal fusion to GFP and a Twin-Strep-tag. The full coding sequences of all constructs were verified by DNA sequencing. HEK293T cells (5X10^5^ cells / 35mm plate) were transiently transfected with 1.0 μ g of WT or mutant CLCN4-GFP-Twin-Strep plasmid DNA using polyethylenimine (PEI) transfection reagent (Polysciences) ^64^. Approximately 24 hours after transfection, the cells were gently transferred onto poly-L-lysine (PLL)-coated coverslips and allowed to attach for at least 6 hours prior to recording. Whole-cell patch-clamp recordings were conducted 30 to 48 hours post-transfection. Recording pipettes were fabricated from borosilicate glass (World Precision Instruments) using a P-1000 puller (Sutter Instruments) and fire-polished using an MF-900 microforge (Narishige). The pipettes had a resistance of 2 to 4 MΩ when filled with the internal solution contained (in mM): 145 NaCl, 15 HEPES, 4 K-gluconate, 2 CaCl₂, and 1 MgCl₂ (pH 7.4). The external bath solution contained (in mM): 120 NaCl, 15 HEPES, 5 MgCl₂, 5 EGTA, and 5 Na-ATP (pH 7.4). Recordings were conducted using an Axopatch 200B amplifier (Molecular Devices), with data acquisition controlled by pCLAMP 11.2 software via a Digidata 1550B interface (Molecular Devices). Whole-cell currents were elicited by applying 50-ms voltage steps from -115 mV to +175 mV in 10 mV increments from a holding potential of 0 mV. Data were sampled at 2 kHz and low-pass filtered at 1 kHz using a four-pole Bessel filter. All recordings were conducted at room temperature (20∼25 °C). Macroscopic currents were analyzed using Clampfit 11.2 (Molecular Devices) and Origin 9.1 (OriginLab Corporation).

### Prediction of electrostatic change

To analyze the electrostatic (ES) potential changes between CLC-4 and its variants, the structures were predicted using RoseTTAFold ^65^. Among the five predicted models, the structure with the highest predicted Local Distance Difference Test (pLDDT) score was selected and preprocessed with PDB2PQR. The electrostatic potential maps were then calculated using APBS software ^66^. The calculations were conducted under conditions of pH 7, 298.15 K, using the AMBER force field, with protonation states assigned by the PropKa method, and no additional ions were added to maintain zero net ionic concentration. The global electrostatic potential of each model was visualized using PyMOL, with the biomolecular surface colored from red (−5 kT/e) to blue (+5 kT/e) according to the electrostatic potentials. The electrostatic potential of each residue was calculated as the arithmetic mean of the electrostatic potentials of all non-hydrogen atoms within the residue.

### Structural visualization of the AlphaFold-predicted CLC-4 protein model

The dimeric structural model of human CLC-4 protein was predicted using AlphaFold 3 via the AlphaFold server (https://alphafoldserver.com/), with the UniProt ID P51793 as the input sequence ^67^. The predicted structure was visualized, and atom-to-atom distance measurements were performed using UCSF ChimeraX version 1.9 ^68^.

### Brain organoid differentiation

The brain organoid differentiated as described in a previous report (Do et al.). Briefly, WT and *CLCN4*-variant ESC colonies were detached to be differentiated into the embryoid bodies. The embryoid bodies were cultured in floating system for a week with the first medium: DMEM/F-12+Glutamax (Gibco, 10565042), 20% serum replacement, MEM-NEAA (Gibco, 11140050), 2-Mercaptoethanol (Merck, M3148), 1% penicillin/streptomycin (Gibco, 14140122), 2 uM Dorsomorphin (Stem cell technologies, 72102), 2 uM A-83 (Stem cell technologies, 72022). Then the embryoid bodies were embedded into Matrigel and cultured in the second medium: DMEM/F-12+Glutamax, N2 supplement (Gibco, 17502001), MEM-NEAA, 1% penicillin/streptomycin, 1 uM CHIR-99021 (Stem cell technologies, 72052), 1 uM SB-431542 (Cayman, 13031). The organoids were mechanically dissociated from Matrigel on day 14 and cultured in the shaking condition with the third medium: DMEM/F-12+Glutamax, N2 supplement, B27 supplement (Gibco, 12587001), MEM-NEAA, 2-Merceptoethanol, 1% penicillin/streptomycin, and 2.5 ug/ml insulin (Sigma, I9278).

### scRNA-seq

Day 50 brain organoid samples were first divided into two pieces and dissociated into single cells using Accutase with 100 ug/ml of DNase I (Roche, 10104159001). More than 10,000 cells from each sample and each batch were used for sequencing. The sequencing library was generated using Chromium Next GEM Single Cell 3’ Kit v3.1 (10x Genomics) and run using the Illumina NovaSeq 6000 platform. The sequencing result was analyzed by python toolkit, Scanpy 1.8.1. Cells with total counts less than 1000 and number of genes less than 500 were filtered and further analysis was performed. GEO accession number: GSE298461.

### Immunochemistry

Organoid samples were rinsed 1-2 times with PBS and subsequently fixed with 4% cold paraformaldehyde for 1 hour at RT. Organoids were then cryoprotected in 30% sucrose at 4°C until fully saturated, embedded in a block mold using a tissue freezing medium (Leica, 14020108926), and rapidly frozen on dry ice. Frozen samples were sectioned at 40 μm thickness using a cryostat (CM1850, Leica Biosystems). Sections were permeabilized and blocked in PBS containing 0.1% Triton X-100 (Promega, H5141) and 5% donkey serum (Abcam, Ab7475) for 1 hour at RT.

Cells cultured on microscope cover glass (Marienfeld) were fixed with 4% paraformaldehyde for 30 minutes at RT, permeabilized with 0.1% Triton X-100 in PBS containing 0.1% tween-20 (PBS-T) for 30 minutes, and blocked with 5% donkey serum in PBS-T for 1 hour at RT.

After blocking, both organoid sections and cultured cells were incubated with primary antibodies overnight at 4°C. The following day, samples were incubated with fluorescent dye--conjugated secondary antibodies for 1 hour at RT. DAPI (1 ug/ml; Sigma, D9542) was used to stain nuclei for 10 minutes at RT. Samples were mounted using mounting solution (Dako, S3023) and imaged using a confocal microscope (LSM800, Zeiss). Primary antibodies used for immunofluorescence were rabbit anti-SOX2 (Cell signaling, 3579S; 1:1000), mouse anti-NESTIN (Milipore, MAB5326; 1:1000), mouse anti-TUJ1 (R&D systems, mab1195; 1:1000), and rabbit anti-TUJ1 (abcam, ab52623; 1:1000). Secondary antibodies used were Alexa Fluor 488 anti-mouse (abcam, ab150105; 1:1000), Alexa Fluor 594 anti-mouse (abcam, ab150108 1:1000), Alexa Fluor 647 anti-mouse (abcam, ab150107; 1:1000), Alexa Fluor 488 anti-rabbit (abcam, ab150073; 1:1000), Alexa Fluor 594 anti-rabbit (abcam, ab150068; 1:1000), and Alexa Fluor 647 anti-rabbit (abcam, ab150075; 1:1000). All images within each figure panel were scaled consistently.

### Differentiation of hESCs into NPCs and neurons

hESCs were differentiated into neurons following a previously described protocol. Cultured on MEFs, the hESCs were treated with 10 μg/ml of collagenase type IV and incubated at 37°C, 5% CO2 incubator for 1 hour until the colonies detached completely from the dish. The detached colonies were then transferred to a non-adherent dish (SPL, 10090). Embryoid bodies (EBs) were cultured under floating condition in EB media comprising DMEM/F-12+Glutamax, N2 supplement, B27 supplement, 10 μM SB-431542, and 0.2 μM LDN193189 (Selleckchem, S2618). After 7 days, the EBs were transferred to a matrigel-coated plate with EB media supplemented with 1 μg/ml laminin (Gibco, 23017015). After 3-4 days, neural rosettes were manually picked, dissociated into NPCs using Accutase and plated on poly-L-ornithine (Sigma, P3655)/laminin-coated plate. NPCs were then cultured in DMEM/F-12+Glutamax, supplemented with N2, B27, and 20 ng/ml bFGF. For neuronal differentiation, NPCs were plated at a density of 2.5-3×10^6^ cells/cm^2^ in DMEM/F-12+Glutamax, supplemented with N2, B27, 20 ng/ml BDNF (Peprotech, 450-02), 20 ng/ml GDNF (peprotech, 450-10), 500ng/ml dcAMP (Selleckchem, S7858), 200 nM Ascorbic acid (Sigma, A4544), and 1 ug/ml laminin.

### Generation of lentivirus

Lentivirus was packaged in HEK293T cells cultured in DMEM with 10% FBS. HEK293T cells were transfected with Polyethylenimine (PEI) (Polysciences, 23966). For a 150 mm dish, the following solution was prepared: 12.2 μg lentiviral DNA, 8.1 μg MDL-gagpol, 3.1 μg Rev-RSV, 4.1 μg CMV-VSVG, 1 ml of serum-free DMEM and 110 μg of PEI. After gently mixing, the solution was incubated at RT for 5 minutes and then added drop-wise to the dishes. The culture medium was changed four hours later, and the virus was harvested at 72 hours post-transfection. To collect viruses, the culture medium was centrifuged at 19,400 rpm for two hours. The viral pellet was resuspended in DPBS. The titer of the harvested lentivirus was determined by serial dilution and infection.

### RNA extraction and real-time PCR

Total RNA from cultured cells was extracted using TRIzol (Invitrogen, 15596-018) and synthesized into cDNA using RevertAid First Strand cDNA Synthesis Kit (ThermoFisher, EP0442) according to the manufacturer’s protocol. Real-time PCR was conducted by SYBR green method with SYBR Green real-time PCR master mix (Applied biosystems, 4368702). Quantification was performed using the relative standard curve method and normalized to ACTB. The primer sequences used in this study were as follows: *CLCN4*, 5’-GCTGGAGTCTC TGTTGCCTT-3’ (forward) and 5’- TCCACAAGGTCTTCAGGGGA-3’ (reverse); *MEG3*, 5’- CC TGCTGCCCATCTACACCTC-3’ (forward) and 5’- CCAGGATGGCAAAGGATGAAGAGG-3’ (reverse); ACTB, 5’- GCCAACCGCGAGAAGATGAC-3’ and 5’- GAGGCGTACAGGGATAGC ACAG-3’ (reverse).

### TUNEL assay

The TUNEL assay was performed on day 6 differentiated neurons using In Situ Cell Death Detection Kit, Fluorescein (Roche, 11684795910) according to the manufacturer’s protocol. After the TUNEL reaction, the cells were stained with 1 μg/ml DAPI. The prepared samples were imaged using an LSM800 confocal microscope (Zeiss) with a 20x objectives. The intensity of the TUNEL reaction was measured using ImageJ and normalized to the DAPI intensity.

### Quantification of neuronal cell number

To compare changes in neuronal cell numbers, an equal number of NPCs were plated onto the poly-L-ornithine/laminin-coated 4-well cell culture plate. Twenty-four hours after plating, the NPC medium was replaced with neuronal medium to induce differentiation. On day 4 of differentiation, cells were treated with 1 μg/ml of Hoechst 33342 (Invitrogen, H3570) and incubated at 37℃, in 5% CO2 incubator for 15 minutes. Before imaging, the samples were washed twice with PBS to remove debris. Images were captured using an inverted fluorescence microscope (Olympus) with 10x objectives. This process was repeated for neurons differentiated for 8 days. The number of Hoechst-stained cells were manually counted using the ImageJ plugin, Cell Counter. Each experiment included WT as a control and the variants as experimental groups. Values from the variants were normalized to those of WT in each biological replicates. Results were expressed as relative values compared to the WT.

### Subcellular organelle visualization

Neurons differentiated from NPCs for 6 days were stained with 50 nM LysoTracker Red DND-99 (Invitrogen, L7528) and 1 μM Lysosensor Yellow/Blue DND-160 (Invitrogen, L7545) to visualize acidic organelles and measure their relative pH, respectively. The neurons were incubated with LysoTracker for 1 hour and with Lysosensor for 5 minutes at 37°C, followed by fixation with 4% paraformaldehyde. LysoTracker treated samples were stained with DAPI for 10 minutes at RT, and LysoSenser treated samples underwent immunocytochemistry using TUJ1 antibody and Alexa Fluor 647-conjugated secondary antibody. For specific subcellular vesicle visualization, lentiviral constructs encoding GFP-tagged RAB5 (Addgene #134858), RAB7 (Addgene #133027), RAB11 (Addgene #134860), and LAMP1 (Addgene #134868) were utilized. Neurons, differentiated from NPCs for three days, were infected with these GFP-fused markers using lentivirus at an MOI of 0.5-1. Three days post-infection, the neurons were prepared for immunocytochemistry. To monitor autophagic flux, 500 ng of ptfLC3 (Addgene plasmid #21074) was transfected into 20,000 neurons using 1 µl of Lipofectamine stem transfection reagent. Imaging was carried out with a Zeiss LSM800 and LSM980 confocal microscope. The intensity and spatial relationships of the signals were quantified using ImageJ. All images within the same figure were presented at the same scale. The values from the variants were normalized to those of the WT in each biological replicate, and the results were expressed as relative values compared to the WT. In each figure, all representative images within a panel are scaled consistently.

### Neurite length analysis

The NPCs were plated onto the poly-L-ornithine/laminin-coated cover glass. A day after plating, NPCs were differentiated into neurons by infection with GFP-expressing lentiviruses at MOI of 0.1-0.2. On the desired day of neuronal differentiation day, the cells were collected for immunocytochemistry using TUJ1 antibodies and Alexa Fluor 594-conjugated secondary antibodies. Imaging was performed on an LSM800 confocal microscope using a 20x objectives. Neurite length was measured using NeuronJ plugin in ImageJ. In each figure, all representative images within a panel are scaled consistently.

### Drug and chemical treatment

Neurons were treated with the following drugs and chemicals from differentiation day 4 to day 8: risperidone (Sigma-Aldrich, R3030; 10 μM), clozapine (Cayman-Aldrich, 12059; 10 μg/mL), LiCl (Sigma-Aldrich, 203637; 1 mM), rapamycin (Calbiochem, 553210; 1 μM), and calpain inhibitor I (Sigma-Aldrich, A6185; 500 nM). After 4 days of treatment, cells were stained with Hoechst, and total cell numbers were quantified. Cell counts were normalized to those measured on differentiation day 4.

### Total RNA-seq

RNA was extracted from the cells using TRIzol, ribosomal RNAs were depleted, and RNA-Seq libraries were generated using KAPA RNA HyperPrep Kit (Roche 08098107702). Total RNA-Seq libraries were sequenced paired-end 2 x 100 base pairs (bps) using the Illumina NovaSeq 6000 platform. The sequencing reads were mapped to the human genome GRCh38 (hg38) using STAR, version 2.7.9. The mapped reads were annotated and quantified using RSEM. Differential gene expression was analysis using R package, DESeq2. DAVID (https://david.ncifcrf.gov/) was used to perform the functional annotation analysis. GEO accession number: GSE296114.

### *MEG3* regulation via dCas9

*MEG3* activation in *CLCN4*-variant NPCs was conducted using CRISPR/Cas9-activation system. *MEG3*-targeting sgRNAs were designed based on Broad Institute GPP sgRNA designer. Oligonucleotides for sgRNA constructs (sgMEG3-a-F: 5’-CACCGGCAATTTG TCATAGAATCTG-3’ and sgMEG3-a-R: 5’-AAACCAGATTCTATGACAAATTGCC-3’) were synthesized by Bionics (Republic of Korea). Forward and reverse oligos were mixed and annealed by decreasing temperature from 95℃ to room temperature. The annealed sgRNAs were inserted into lentiGuide-puro (Addgene #52963). The sequence of generated plasmids was confirmed by Sanger sequencing. Lentiviruses expressing sgRNA with puromycin resistance gene or dCas9-VPR with blasticidin resistance gene were generated with the previously described method. Both types of lentiviruses were co-infected to *CLCN4*-variant NPCs at an MOI of 0.5 -1 for each. Twenty-four hours post infection, puromycin (1 μg/ml) and blasticidin (10 μg/ml, Invivogen, ant-bl-05) were added to the NPC media for selection. *MEG3* inhibition in WT NPCs was conducted using CRISPR/Cas9-inhibition system. The process replicated that used for generating MEG3-activated NPCs. Oligonucleotides for the inhibition sgRNA constructs (sgMEG3-i-F: 5’-CACCGGCGGCTCCTCAGGAGAGCTG-3’ and sgMEG3-R: 5’-AAACCAGCTCTCCTGAGGAGCCGCC-3’) were synthesized by Bionics. Lentiviruses expressing sgRNA with puromycin resistance gene and dCas9-KRAB with hygromycin resistance gene were generated. Both types of lentiviruses were co-infected to WT NPCs at an MOI of 0.5-1 for each. Twenty-four hours post infection, puromycin (1 μg/ml) and hygromycin (50 μg/ml, ThermoFisher, 10687010) were added to the NPC media for selection.

### Statistical analysis

All error bars are presented as the mean ± standard error of the mean (SEM). P values were determined by the unpaired Student’s t-test, one-way ANOVA with Dunnett’s multiple comparisons, and two-way ANOVA using GraphPad Prism (GraphPad Software). Statistical significance was indicated as follows: *p < 0.05, **p < 0.01, ***p < 0.001, and ****p<0.0001. All experiments were performed in technical and biological replicates. Detailed number of biological replicates and statistical information for each experiment are described in figure legends. For imaging analysis, region of interest (ROI) was randomly selected.

## Acknowledgments

We thank all members of the laboratory for their helpful discussions. ChatGPT (OpenAI) was used for grammar editing and improving language clarity during manuscript preparation

## Funding

Ministry of Health & Welfare, Republic of Korea: HI18C1077 (Yeni Kim, JH)

National Research Foundation of Korea (NRF) grant funded by the Korea government (MSIT): 2021R1A2C1004884 (HHL), RS-2024-00398786 (JH), RS-2024-00335144 (JH) and RS-2024-00407383 (JEP and JH)

KBRI basic research program through Korea Brain Research Institute funded by the Ministry of Science and ICT: 25-BR-01-02 and 25-BR-05-07 (HHL)

Korea Environment Industry & Technology Institute (KEITI) through Technology Development Project for Safety Management of Household Chemical Products funded by Korea Ministry of Environment (MOE): RS-2025-02223058 (JH)

Institute for Basic Science: IBS-R002-A1 (JH)

Fostering the Next Generation of Researchers Program of the NRF of Korea: NRF-2022R1A6A3A13073152 (Dayeon Kim)

KAIST Jang Young Sil Fellow Program (GS)

## Author contributions

Conceptualization: YK (Yeni Kim), JH

Study Design: DK (Dayeon Kim), JH

Experimental Investigation:

- scRNA-seq: YK (Yongjun Koh) under supervision of JEP
- *CLCN4* variants cloning and patch-clamp electrophysiology: HSJ under supervision of HHL
- AlphaFold protein structure prediction: HHL
- RosettaFold-based electrostatic potential prediction: JK under supervision of DK (Donghyuk Kim)
- PSC-derived neural cells and organoid culture: DK (Dayeon Kim), HD, GS under supervision of JH
- Most of all other experiments: DK (Dayeon Kim) under supervision of JH

Data Interpretation: DK (Dayeon Kim), YK (Yongjun Koh), HSJ, DK (Donghyuk Kim), HHL, JEP, JH

Funding acquisition: YK (Yeni Kim), DK (Donghyuk Kim), HHL, JEP, JH

Project administration: JH

Writing – original draft: DK (Dayeon Kim), JH Writing – review & editing: All authors

