## Supplementary Fig for "*MEG3* Enhances Survival of Developing Human Neurons with *CLCN4*-Linked Autophagy Impairment"

#### Supplementary Figures

### Kim et al. Supplementary Fig. 1

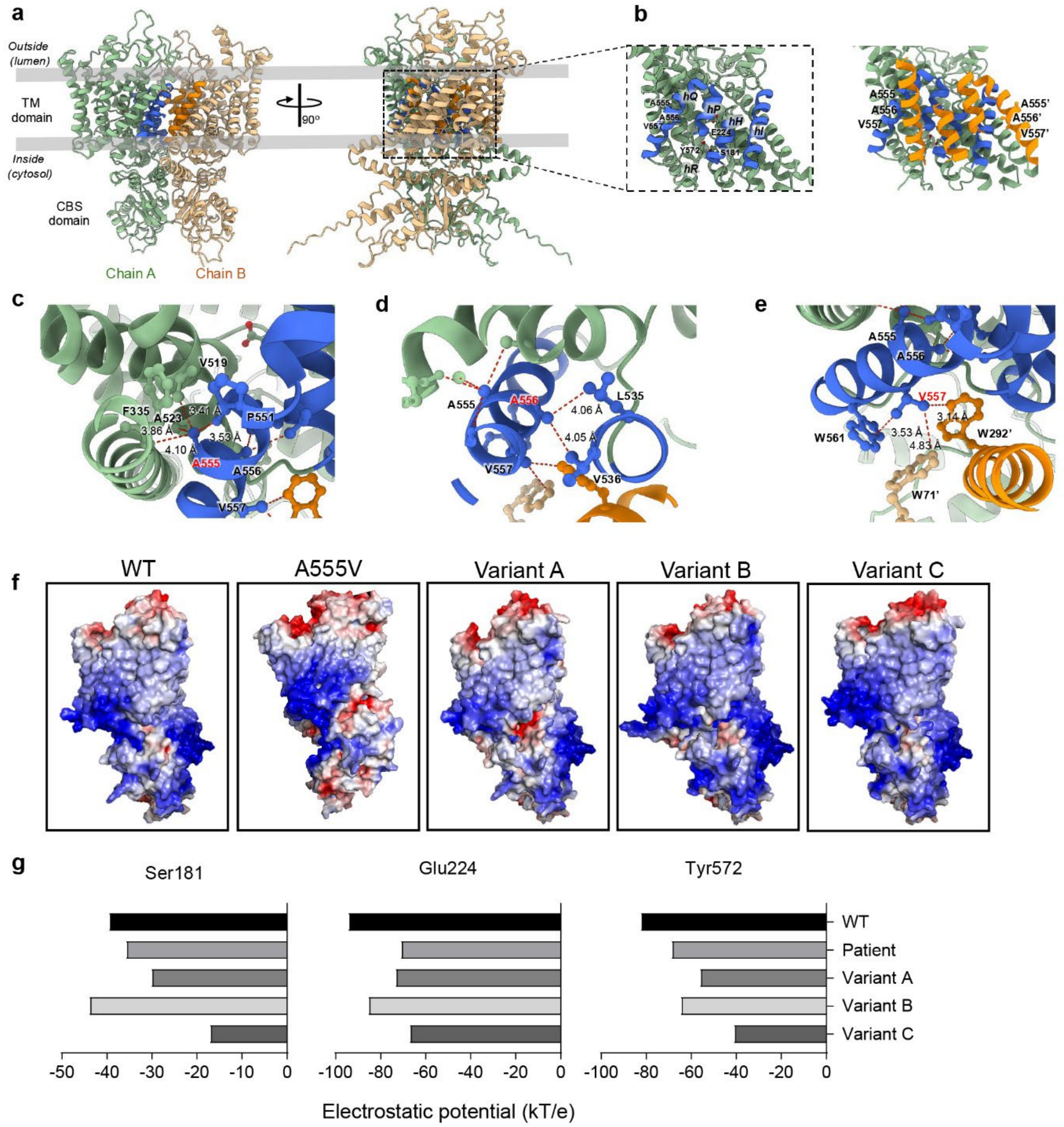


**Supplementary Figure 1. scRNA-seq in WT and *CLCN4*-variant brain organoids**

(a) Cell trajectories from pseudotime analysis. Each panel shows cells colored by sample (left), cell type (middle), and peusotime (right). Arrows represent the direction of cell trajectory.

(b) Density distribution of cell population depending on pseudotime.

#### Kim et al. Supplementary Fig. 2

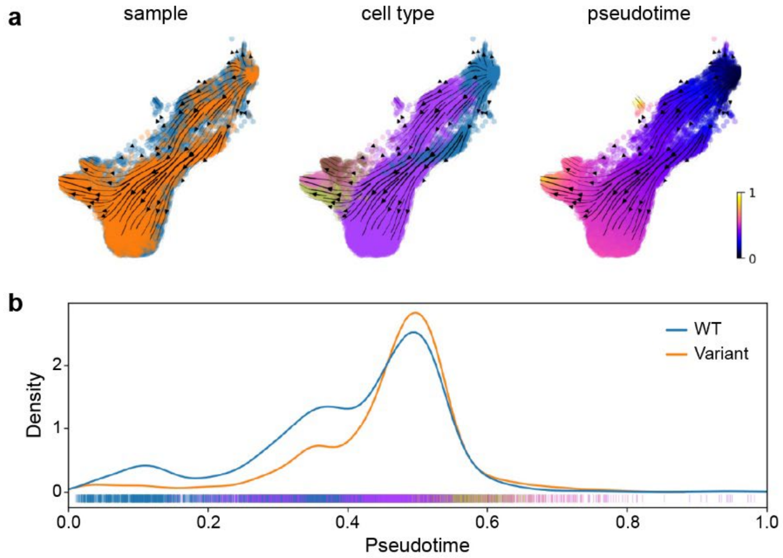


**Supplementary Figure 2. Structural analysis of deep learning–based predicted models of CLC-4 in WT and variant forms**

(a) AlphaFold-predicted structural model of human CLC-4. The two subunits are shown in light green (chain A) and sand (chain B), with dimer-interfacing helices highlighted in blue and orange, respectively.

(b) Helical organization at the dimer interface. Helices hI, hH, hP, and hQ constitute the dimer-interfacing helices. Key residues involved in the inner and outer gates of ion transport (S181, E224, and Y572), as well as the variant residues examined in this study (A555, A556, and V557), are shown as stick model. For clarity, only chain A is shown (left). The inter-subunit contacts formed between dimer-interfacing helices of two subunits are illustrated (right). The residues from chain B are labeled with apostrophes.

(c-e) Atomic distance between each variant residue and its neighboring residues, measured using UCSF ChimeraX: (c) A555 and neighboring residues (side chains of V519 and A523; backbone carbonyls of F335 and P551), (d) A556 and neighboring residues (side chains of L535 and V536), and (e) V557 and neighboring residues (side chains of W561, W71’, and W292’).

(f) RosettaFold-based electrostatic potential prediction of CLC-4 WT and its variants. Surface electrostatic potential was colored from red (-5 kT/e) to blue (+5 kT/e).

(g) Predicted electrostatic potentials at three key residues involved in ion transport of CLC-4.

#### Kim et al. Supplementary Fig. 3

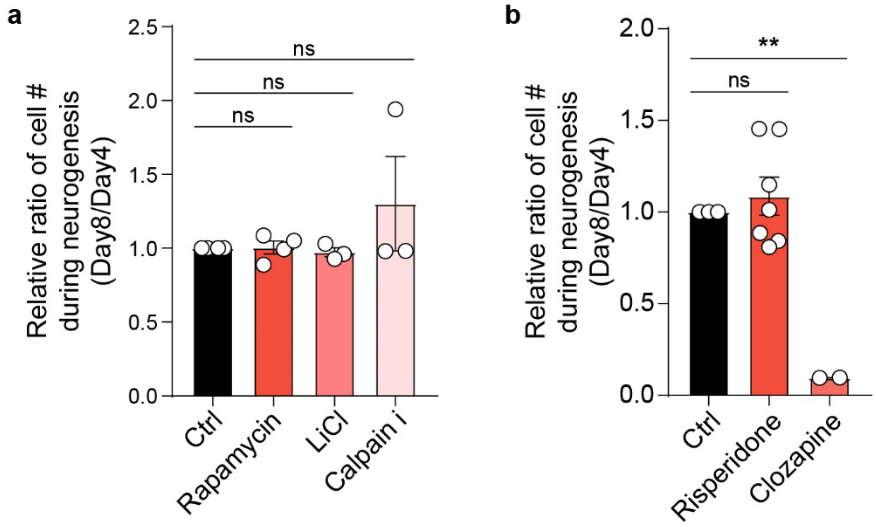


**Supplementary Figure 3. Impact of drug treatment on *CLCN4*-variant neuronal viability**

(a) Changes in cell number ratio in *CLCN4*-variant neurons after treatment of chemicals including rapamycin, LiCl, and calpain inhibitor I, which can enhance autophagy (n>3, mean ± SEM, one-way ANOVA with Dunnett's multiple comparisons test).

(b) Changes in cell number ratio in *CLCN4*-variant neurons after treatment of drugs including risperidone and clozapine (n=7 for risperidone, n=2 for clozapine, mean ± SEM, one-way ANOVA with Dunnett's multiple comparisons test; ***p*=0.0032).

### Kim et al. Supplementary Fig. 4

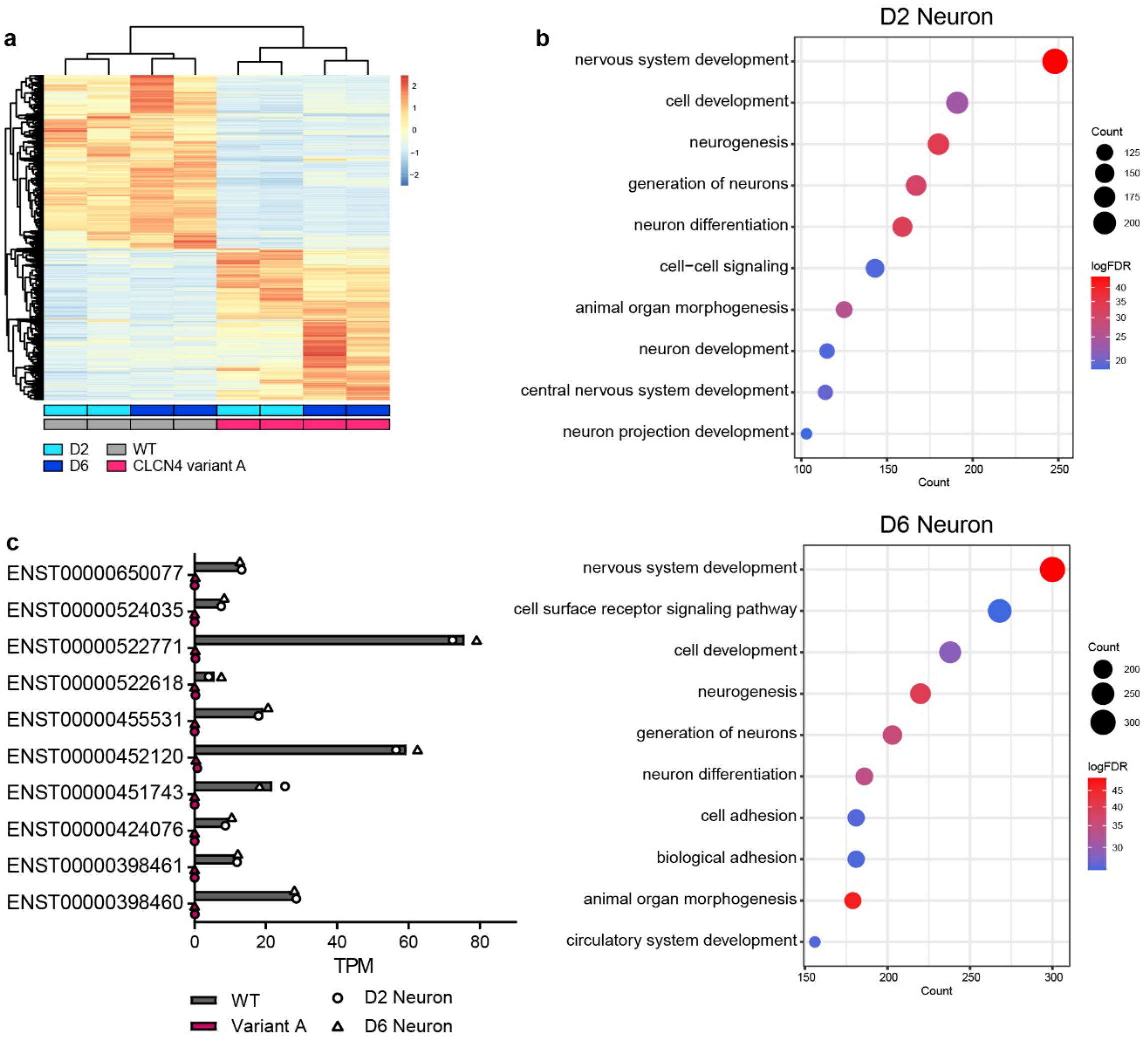


**Supplementary Figure 4. Total RNA sequencing in WT and *CLCN4*-variant neurons at day 2 and 6**

(a) Heatmap and clustering from total RNA sequencing.

(b) GO term analysis using DEGs in day 2 neurons (upper) and day 6 neurons (lower).

(c) Expression level (TPM) of top 10 highly expressed MEG3 transcripts in neurons.

### Kim et al. Supplementary Fig. 5

a

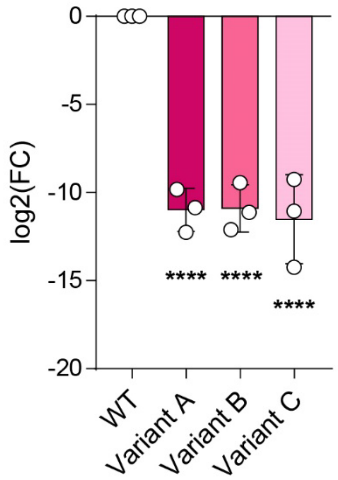


**Supplementary Figure 5. *MEG3* expression in WT and *CLCN4*-variant human NPCs**

(a) The expression of *MEG3* in WT and *CLCN4*-variant NPCs quantified by real-time PCR. (n=3, mean ± SEM, one-way ANOVA with Dunnett's multiple comparisons test, ****$p$ < 0.0001)
