## Supplementary Table 1 for "*MEG3* Enhances Survival of Developing Human Neurons with *CLCN4*-Linked Autophagy Impairment"

| | ensembl_gene_id | baseMean | log2FoldCl | lfcSE | stat | pvalue | padj | D2_WT_1 | D2_varA_1 | D2_WT_2 | D2_varA_2 | hgnc_symb | chromosome_name |
|---|---|---|---|---|---|---|---|---|---|---|---|---|---|
| 1 | ENSG00000002746 | 201.8273 | -1.13897 | 0.288784 | -3.94403 | 8.01E-05 | 0.001779 | 6.724171 | 3.117029 | 5.491689 | 2.208402 | HECW1 | 7 |
| 2 | ENSG00000003137 | 71.3559 | -2.07478 | 0.54861 | -3.78188 | 0.000156 | 0.003183 | 2.398271 | 0.547426 | 5.764047 | 1.302077 | CYP26B1 | 2 |
| 3 | ENSG00000003147 | 61.42863 | 2.447884 | 0.506143 | 4.836347 | 1.32E-06 | 4.30E-05 | 1.228017 | 9.924642 | 2.402694 | 8.724936 | ICA1 | 7 |
| 4 | ENSG00000003402 | 477.2924 | 1.546844 | 0.274746 | 5.630079 | 1.80E-08 | 8.39E-07 | 4.248467 | 19.3866 | 9.157499 | 17.47109 | CFLAR | 2 |
| 5 | ENSG00000004848 | 182.8713 | -5.53515 | 0.472407 | -11.7169 | 1.04E-31 | 3.41E-29 | 17.28234 | 0.477044 | 12.03748 | 0.161121 | ARX | X |
| 6 | ENSG00000005020 | 49.68086 | 1.725308 | 0.551033 | 3.131045 | 0.001742 | 0.024488 | 0.736767 | 3.468446 | 1.640428 | 3.945725 | SKAP2 | 7 |
| 7 | ENSG00000005249 | 555.0624 | -1.29474 | 0.185483 | -6.98037 | 2.94E-12 | 2.41E-10 | 27.31721 | 10.59993 | 25.4601 | 9.914476 | PRKAR2B | 7 |
| 8 | ENSG00000005379 | 246.7286 | -1.22232 | 0.247717 | -4.93434 | 8.04E-07 | 2.73E-05 | 14.9418 | 6.382732 | 14.71392 | 5.741436 | TSPOAP1 | 17 |
| 9 | ENSG00000006210 | 94.0444 | -1.47381 | 0.4087 | -3.6061 | 0.000311 | 0.005769 | 3.594318 | 1.438961 | 5.848267 | 1.771319 | CX3CL1 | 16 |
| 10 | ENSG00000006283 | 238.347 | 1.102685 | 0.267325 | 4.124881 | 3.71E-05 | 0.000895 | 3.052925 | 6.11506 | 3.693418 | 7.640392 | CACNA1G | 17 |
| 11 | ENSG00000006377 | 27.55624 | -8.18839 | 1.583485 | -5.17112 | 2.33E-07 | 8.70E-06 | 4.567116 | 0 | 4.170553 | 0 | DLX6 | 7 |
| 12 | ENSG00000007174 | 104.6943 | -1.70411 | 0.379892 | -4.48578 | 7.26E-06 | 0.000201 | 8.999209 | 2.120969 | 6.661953 | 2.447472 | DNAH9 | 17 |
| 13 | ENSG00000007237 | 240.3777 | -1.00334 | 0.272829 | -3.67756 | 0.000235 | 0.00455 | 4.878784 | 2.457378 | 6.715892 | 3.044259 | GAS7 | 17 |
| 14 | ENSG00000007350 | 55.83542 | -2.20921 | 0.578405 | -3.81949 | 0.000134 | 0.002804 | 7.165915 | 1.264918 | 2.981319 | 0.883263 | TKTL1 | X |
| 15 | ENSG00000007372 | 3442.483 | -2.40373 | 0.185302 | -12.972 | 1.76E-38 | 7.72E-36 | 295.6995 | 59.18303 | 236.0404 | 37.22931 | PAX6 | 11 |
| 16 | ENSG00000008196 | 486.8157 | 3.694221 | 0.264599 | 13.96158 | 2.67E-44 | 1.61E-41 | 2.065481 | 24.00424 | 1.232603 | 16.74365 | TFAP2B | 6 |
| 17 | ENSG00000008283 | 218.7459 | -1.18795 | 0.264124 | -4.49771 | 6.87E-06 | 0.000191 | 13.18167 | 5.838991 | 15.59076 | 6.159831 | CYB561 | 17 |
| 18 | ENSG00000008394 | 352.5082 | 2.620395 | 0.282639 | 9.271162 | 1.84E-20 | 3.37E-18 | 10.36975 | 88.52319 | 21.41128 | 96.60567 | MGST1 | 12 |
| 19 | ENSG00000009709 | 321.2594 | 1.89755 | 0.232947 | 8.145849 | 3.77E-16 | 4.98E-14 | 2.783218 | 11.03502 | 2.887225 | 9.132176 | PAX7 | 1 |
| 20 | ENSG00000009950 | 38.88141 | 4.232562 | 0.715987 | 5.911508 | 3.39E-09 | 1.76E-07 | 0.123571 | 3.833505 | 0.271688 | 3.290103 | MLXIPL | 7 |
| 21 | ENSG00000011201 | 2938.033 | -3.30628 | 0.184804 | -17.8907 | 1.39E-71 | 2.44E-68 | 95.65078 | 13.01474 | 122.906 | 8.154355 | ANOS1 | X |
| 22 | ENSG00000011677 | 111.8262 | -1.39757 | 0.353787 | -3.95032 | 7.80E-05 | 0.001741 | 5.555902 | 1.975451 | 4.766602 | 1.766438 | GABRA3 | X |
| 23 | ENSG00000013293 | 44.54103 | -2.80818 | 0.612155 | -4.58736 | 4.49E-06 | 0.000129 | 1.257344 | 0.104458 | 0.683932 | 0.154387 | SLC7A14 | 3 |
| 24 | ENSG00000015592 | 512.9163 | -1.2293 | 0.19593 | -6.27417 | 3.52E-10 | 2.16E-08 | 70.39315 | 31.81779 | 69.41108 | 25.19933 | STMN4 | 8 |
| 25 | ENSG00000018408 | 558.2768 | 1.220859 | 0.212222 | 5.752749 | 8.78E-09 | 4.29E-07 | 8.97873 | 25.98539 | 13.50203 | 23.85819 | WWTR1 | 3 |
| 26 | ENSG00000021645 | 534.7105 | -2.25596 | 0.192386 | -11.7262 | 9.36E-32 | 3.16E-29 | 24.91215 | 5.443214 | 26.0499 | 4.744813 | NRXN3 | 14 |
| 27 | ENSG00000021826 | 783.5559 | 1.135566 | 0.197479 | 5.75031 | 8.91E-09 | 4.34E-07 | 9.087257 | 21.79715 | 13.15677 | 24.56634 | CPS1 | 2 |
| 28 | ENSG00000024422 | 52.00023 | 2.03775 | 0.552542 | 3.687952 | 0.000226 | 0.004393 | 0.712399 | 2.722446 | 1.1082 | 4.323742 | EHD2 | 19 |
| 29 | ENSG00000026508 | 69.84521 | 1.438465 | 0.463981 | 3.100263 | 0.001933 | 0.02658 | 1.951284 | 7.438743 | 2.836667 | 5.04308 | CD44 | 11 |
| 30 | ENSG00000026559 | 98.6753 | 2.462258 | 0.475665 | 5.176449 | 2.26E-07 | 8.48E-06 | 2.328188 | 7.498843 | 2.557442 | 17.91376 | KCNG1 | 20 |
| 31 | ENSG00000028116 | 62.93285 | 2.010911 | 0.474019 | 4.242262 | 2.21E-05 | 0.000561 | 1.632198 | 7.311125 | 1.7821 | 5.839786 | VRK2 | 2 |
| 32 | ENSG00000033122 | 238.9304 | -1.5012 | 0.273884 | -5.48114 | 4.23E-08 | 1.85E-06 | 3.971088 | 1.765389 | 5.510958 | 1.428213 | LRRC7 | 1 |
| 33 | ENSG00000033170 | 881.7679 | -1.12873 | 0.160261 | -7.04303 | 1.88E-12 | 1.59E-10 | 51.60466 | 21.97338 | 49.34384 | 21.98837 | FUT8 | 14 |
| 34 | ENSG00000035862 | 1530.718 | -1.24643 | 0.17139 | -7.27244 | 3.53E-13 | 3.30E-11 | 62.08834 | 30.3994 | 88.24664 | 29.82767 | TIMP2 | 17 |
| 35 | ENSG00000036565 | 72.19848 | -2.69751 | 0.459404 | -5.87177 | 4.31E-09 | 2.19E-07 | 7.987126 | 0.965784 | 6.262265 | 1.121592 | SLC18A1 | 8 |
| 36 | ENSG00000039139 | 99.52527 | -1.66017 | 0.433769 | -3.82732 | 0.00013 | 0.002728 | 1.561385 | 0.758276 | 1.861077 | 0.310747 | DNAH5 | 5 |
| 37 | ENSG00000040731 | 209.9688 | -1.47455 | 0.281105 | -5.24553 | 1.56E-07 | 6.04E-06 | 9.305669 | 4.136084 | 12.27141 | 3.282913 | CDH10 | 5 |
| 38 | ENSG00000041353 | 29.03 | -2.21909 | 0.745899 | -2.97506 | 0.002929 | 0.037336 | 1.52332 | 0.203925 | 0.525425 | 0.201119 | RAB27B | 18 |
| 39 | ENSG00000041982 | 413.8253 | -1.12765 | 0.25224 | -4.47053 | 7.80E-06 | 0.000215 | 13.29071 | 4.60402 | 8.21849 | 4.782678 | TNC | 9 |
| 40 | ENSG00000046889 | 1860.063 | -1.40606 | 0.162881 | -8.63245 | 6.01E-18 | 9.47E-16 | 47.5294 | 14.9423 | 36.39416 | 15.24016 | PREX2 | 8 |
| 41 | ENSG00000047644 | 1011.488 | -1.06053 | 0.160714 | -6.59887 | 4.14E-11 | 2.90E-09 | 22.70483 | 11.10551 | 26.858 | 11.4766 | WWC3 | X |
| 42 | ENSG00000047648 | 69.13652 | 1.387546 | 0.455015 | 3.04945 | 0.002293 | 0.030578 | 0.847343 | 3.038615 | 1.402984 | 2.588413 | ARHGAP6 | X |
| 43 | ENSG00000048540 | 747.2966 | -1.071 | 0.187879 | -5.7005 | 1.19E-08 | 5.69E-07 | 42.33624 | 20.58137 | 36.82545 | 15.49056 | LMO3 | 12 |
| 44 | ENSG00000050030 | 363.9831 | -1.15626 | 0.222058 | -5.20702 | 1.92E-07 | 7.26E-06 | 4.951403 | 2.225805 | 4.403211 | 1.790602 | NEXMIF | X |
| 45 | ENSG00000052126 | 2141.821 | -1.46273 | 0.149375 | -9.79231 | 1.21E-22 | 2.46E-20 | 122.891 | 46.31787 | 154.6063 | 49.31011 | PLEKHA5 | 12 |
| 46 | ENSG00000053438 | 3147.583 | -2.18969 | 0.160028 | -13.6831 | 1.28E-42 | 7.04E-40 | 614.7466 | 137.7648 | 520.206 | 100.3829 | NNAT | 20 |
| 47 | ENSG00000059804 | 835.4304 | 1.13995 | 0.212093 | 5.374755 | 7.67E-08 | 3.18E-06 | 14.79563 | 36.96655 | 23.69574 | 43.46541 | SLC2A3 | 12 |
| 48 | ENSG00000064300 | 182.7318 | 1.165312 | 0.295604 | 3.942143 | 8.08E-05 | 0.001791 | 3.247296 | 9.000589 | 4.634785 | 7.841015 | NGFR | 17 |
| 49 | ENSG00000064692 | 240.4686 | -1.23549 | 0.316589 | -3.9025 | 9.52E-05 | 0.002056 | 18.89483 | 7.924685 | 11.80394 | 4.650367 | SNCAIP | 5 |
| 50 | ENSG00000066382 | 184.5257 | -2.72273 | 0.327366 | -8.31708 | 9.02E-17 | 1.32E-14 | 26.87445 | 3.485731 | 17.86316 | 2.987693 | MPPED2 | 11 |
| 51 | ENSG00000066468 | 814.0978 | 1.727475 | 0.195666 | 8.828708 | 1.06E-18 | 1.73E-16 | 15.6068 | 50.79695 | 12.39128 | 37.81817 | FGFR2 | 10 |
| 52 | ENSG00000066735 | 188.0719 | -1.64913 | 0.28546 | -5.77709 | 7.60E-09 | 3.75E-07 | 4.490201 | 1.615617 | 5.026121 | 1.287055 | KIF26A | 14 |
| 53 | ENSG00000067182 | 163.9488 | 1.595867 | 0.334354 | 4.77299 | 1.82E-06 | 5.74E-05 | 4.220279 | 16.86434 | 7.792664 | 17.67884 | TNFRSF1A | 12 |
| 54 | ENSG00000067798 | 435.7608 | -1.40852 | 0.207576 | -6.78556 | 1.16E-11 | 8.73E-10 | 17.50415 | 7.224539 | 21.37363 | 6.720674 | NAV3 | 12 |
| 55 | ENSG00000069431 | 366.1971 | 1.531561 | 0.229179 | 6.682804 | 2.34E-11 | 1.70E-09 | 4.322811 | 15.07522 | 5.427003 | 11.85718 | ABCC9 | 12 |
| 56 | ENSG00000069696 | 51.65617 | -2.62658 | 0.604469 | -4.34526 | 1.39E-05 | 0.000368 | 10.64098 | 1.990007 | 6.074952 | 0.671918 | DRD4 | 11 |
| 57 | ENSG00000070193 | 214.1543 | 4.367714 | 0.392063 | 11.14033 | 7.98E-29 | 2.36E-26 | 0.398169 | 5.999231 | 0.472789 | 11.12134 | FGF10 | 5 |
| 58 | ENSG00000070729 | 49.72343 | -3.88678 | 0.622606 | -6.24276 | 4.30E-10 | 2.59E-08 | 3.011317 | 0.330766 | 3.302119 | 0.12566 | CNGB1 | 16 |
| 59 | ENSG00000070808 | 140.5388 | 1.551103 | 0.335231 | 4.626968 | 3.71E-06 | 0.000109 | 2.160465 | 8.239055 | 3.017032 | 6.279083 | CAMK2A | 5 |
| 60 | ENSG00000071967 | 94.22658 | 1.214042 | 0.406024 | 2.990078 | 0.002789 | 0.035927 | 1.286814 | 3.413578 | 2.190714 | 4.228597 | CYBRD1 | 2 |
| 61 | ENSG00000072133 | 614.1942 | -1.01475 | 0.1768 | -5.73954 | 9.49E-09 | 4.59E-07 | 12.83206 | 6.026677 | 12.49987 | 5.91134 | RPS6KA6 | X |
| 62 | ENSG00000072315 | 110.9141 | -1.03823 | 0.351299 | -2.95539 | 0.003123 | 0.0395 | 1.225215 | 0.627415 | 1.501143 | 0.63464 | TRPC5 | X |
| 63 | ENSG00000072657 | 54.38704 | 2.244375 | 0.513695 | 4.369078 | 1.25E-05 | 0.000331 | 0.365475 | 2.346955 | 0.597131 | 2.005136 | TRHDE | 12 |
| 64 | ENSG00000072682 | 93.83723 | 1.313501 | 0.397867 | 3.301361 | 0.000962 | 0.01495 | 2.281303 | 6.148015 | 3.457802 | 7.451419 | P4HA2 | 5 |
| 65 | ENSG00000075035 | 87.05654 | 3.01095 | 0.449815 | 6.693757 | 2.18E-11 | 1.59E-09 | 1.062164 | 5.895283 | 0.606958 | 6.23428 | WSCD2 | 12 |
| 66 | ENSG00000076706 | 459.2245 | 1.054219 | 0.200643 | 5.254213 | 1.49E-07 | 5.79E-06 | 10.57646 | 20.05166 | 11.02574 | 22.55788 | MCAM | 11 |
| 67 | ENSG00000076770 | 746.0879 | 1.1762 | 0.190387 | 6.177945 | 6.49E-10 | 3.86E-08 | 6.05443 | 11.88516 | 4.525914 | 10.93149 | MBNL3 | X |
| 68 | ENSG00000077264 | 4754.713 | -1.02475 | 0.125364 | -8.17415 | 2.98E-16 | 4.04E-14 | 95.01464 | 45.36143 | 105.7486 | 48.46388 | PAK3 | X |
| 69 | ENSG00000077274 | 57.65704 | -2.22362 | 0.607085 | -3.66279 | 0.000249 | 0.004792 | 1.4203 | 0.803562 | 4.897951 | 0.492528 | CAPN6 | X |
| 70 | ENSG00000077943 | 14.97483 | -3.11406 | 1.070683 | -2.90848 | 0.003632 | 0.044047 | 1.127109 | 0.05467 | 0.659926 | 0.171617 | ITGA8 | 10 |
| 71 | ENSG00000078295 | 269.2041 | -2.32536 | 0.327977 | -7.09002 | 1.34E-12 | 1.16E-10 | 8.03713 | 2.811537 | 16.01436 | 1.734286 | ADCY2 | 5 |
| 72 | ENSG00000078549 | 396.8982 | 1.959339 | 0.238676 | 8.209198 | 2.23E-16 | 3.08E-14 | 2.734306 | 11.33514 | 3.989763 | 13.46105 | ADCYAP1F | 7 |
| 73 | ENSG00000079156 | 335.9206 | -1.11563 | 0.227308 | -4.90802 | 9.20E-07 | 3.08E-05 | 8.806044 | 3.603044 | 9.352364 | 4.338971 | OSBPL6 | 2 |
| 74 | ENSG00000079841 | 185.9829 | -1.44834 | 0.283855 | -5.1024 | 3.35E-07 | 1.21E-05 | 7.967052 | 3.197488 | 8.402583 | 2.528717 | RIMS1 | 6 |
| 75 | ENSG00000079931 | 144.933 | -2.41256 | 0.348357 | -6.92555 | 4.34E-12 | 3.39E-10 | 11.15319 | 1.457237 | 10.01162 | 2.307996 | MOXD1 | 6 |
| 76 | ENSG00000080166 | 119.0056 | -3.20764 | 0.454404 | -7.05901 | 1.68E-12 | 1.43E-10 | 3.230085 | 0.401414 | 7.239765 | 0.657526 | DCT | 13 |
| 77 | ENSG00000080493 | 64.68697 | -1.66696 | 0.530082 | -3.14473 | 0.001662 | 0.02356 | 1.362705 | 0.839272 | 3.352289 | 0.588461 | SLC4A4 | 4 |

| | | | | | | | | | | | | | | |
|---|---|---|---|---|---|---|---|---|---|---|---|---|---|---|
| 78 | ENSG00000080573 | 93.45793 | 2.511338 | 0.488011 | 5.146071 | 2.66E-07 | 9.81E-06 | 0.547864 | 4.04514 | 0.46533 | 1.525145 | COL5A3 | | 19 |
| 79 | ENSG00000080947 | 231.1932 | 1.104045 | 0.323927 | 3.408309 | 0.000654 | 0.010771 | 5.956862 | 18.27577 | 6.911742 | 8.589167 | CROCCP3 | | 1 |
| 80 | ENSG00000081138 | 413.8096 | -3.7421 | 0.238431 | -15.6946 | 1.65E-55 | 1.86E-52 | 21.43638 | 1.437272 | 19.24912 | 1.454574 | CDH7 | | 18 |
| 81 | ENSG00000081665 | 435.2573 | -1.19982 | 0.215923 | -5.55668 | 2.75E-08 | 1.25E-06 | 24.54223 | 8.727457 | 23.8554 | 11.26268 | ZNF506 | | 19 |
| 82 | ENSG00000081803 | 170.0358 | -2.56083 | 0.368591 | -6.94761 | 3.72E-12 | 2.97E-10 | 5.193749 | 1.256575 | 10.53299 | 1.267747 | CADPS2 | | 7 |
| 83 | ENSG00000082684 | 517.2972 | 1.43155 | 0.205957 | 6.950711 | 3.63E-12 | 2.94E-10 | 12.63379 | 32.05834 | 10.1025 | 26.54965 | SEMA5B | | 3 |
| 84 | ENSG00000082781 | 338.8128 | -1.03023 | 0.225852 | -4.56152 | 5.08E-06 | 0.000145 | 20.47342 | 8.62909 | 20.36577 | 10.32446 | ITGB5 | | 3 |
| 85 | ENSG00000083123 | 1615.74 | 1.025617 | 0.173673 | 5.905451 | 3.52E-09 | 1.82E-07 | 41.74877 | 74.4116 | 48.26661 | 99.39333 | BCKDHB | | 6 |
| 86 | ENSG00000084710 | 789.8739 | -1.28243 | 0.182837 | -7.01407 | 2.31E-12 | 1.92E-10 | 17.61137 | 7.883723 | 17.06845 | 5.776491 | EFR3B | | 2 |
| 87 | ENSG00000085276 | 398.7697 | -1.5551 | 0.21418 | -7.26072 | 3.85E-13 | 3.58E-11 | 16.49049 | 6.320696 | 19.5359 | 5.378381 | MECOM | | 3 |
| 88 | ENSG00000086696 | 59.95449 | 3.759563 | 0.573684 | 6.553365 | 5.63E-11 | 3.87E-09 | 0.2018 | 5.994311 | 0.62514 | 4.355467 | HSD17B2 | | 16 |
| 89 | ENSG00000087495 | 104.2219 | -2.17844 | 0.39411 | -5.52749 | 3.25E-08 | 1.46E-06 | 7.06344 | 2.15123 | 9.54491 | 1.388377 | PHACTR3 | | 20 |
| 90 | ENSG00000087510 | 89.01844 | -3.91505 | 0.480291 | -8.15141 | 3.60E-16 | 4.84E-14 | 7.360741 | 0.570457 | 6.493322 | 0.326088 | TFAP2C | | 20 |
| 91 | ENSG00000089116 | 226.6662 | -3.20465 | 0.348898 | -9.18508 | 4.11E-20 | 7.40E-18 | 21.74439 | 2.901748 | 16.30032 | 1.198039 | LHX5 | | 12 |
| 92 | ENSG00000089472 | 691.6095 | 1.063983 | 0.209508 | 5.078494 | 3.80E-07 | 1.36E-05 | 8.815531 | 21.43056 | 13.68039 | 23.18667 | HEPH | X | |
| 93 | ENSG00000091129 | 2459.891 | -1.87354 | 0.141898 | -13.2034 | 8.39E-40 | 4.04E-37 | 98.26278 | 23.26929 | 95.74783 | 27.04359 | NRCAM | | 7 |
| 94 | ENSG00000091409 | 1177.235 | -1.18586 | 0.148298 | -7.99649 | 1.28E-15 | 1.62E-13 | 45.04471 | 20.0684 | 45.73362 | 17.99166 | ITGA6 | | 2 |
| 95 | ENSG00000091513 | 25.81937 | -3.75769 | 0.78862 | -4.76489 | 1.89E-06 | 5.92E-05 | 0.463063 | 0.034591 | 0.882147 | 0.057833 | TF | | 3 |
| 96 | ENSG00000092421 | 3620.035 | -1.44873 | 0.137956 | -10.5013 | 8.52E-26 | 2.16E-23 | 151.3107 | 62.20659 | 188.4796 | 56.29983 | SEMA6A | | 5 |
| 97 | ENSG00000095596 | 222.3114 | 6.221296 | 0.507157 | 12.26701 | 1.36E-34 | 5.14E-32 | 1.308554 | 46.47128 | 0.373424 | 44.15317 | CYP26A1 | | 10 |
| 98 | ENSG00000096060 | 851.9949 | 1.339762 | 0.180158 | 7.43658 | 1.03E-13 | 1.01E-11 | 12.05753 | 32.56357 | 15.96903 | 34.85203 | FKBP5 | | 6 |
| 99 | ENSG00000096696 | 89.62723 | 1.333447 | 0.3929 | 3.393861 | 0.000689 | 0.01125 | 0.608269 | 1.767204 | 0.720533 | 1.438451 | DSP | | 6 |
| 100 | ENSG00000099250 | 1246.641 | -1.4551 | 0.169482 | -8.58559 | 9.04E-18 | 1.40E-15 | 54.1292 | 22.48501 | 73.19719 | 21.65642 | NRP1 | | 10 |
| 101 | ENSG00000099994 | 39.475 | -2.42182 | 0.603702 | -4.01162 | 6.03E-05 | 0.001388 | 2.599838 | 0.412228 | 2.079669 | 0.418978 | SUSD2 | | 22 |
| 102 | ENSG00000100060 | 43.23451 | -1.6631 | 0.552487 | -3.0102 | 0.002611 | 0.034112 | 5.301758 | 1.473063 | 5.831039 | 1.839679 | MFNG | | 22 |
| 103 | ENSG00000100095 | 196.0604 | -1.68978 | 0.312418 | -5.40872 | 6.35E-08 | 2.69E-06 | 4.373541 | 1.622637 | 7.27714 | 1.797482 | SEZ6L | | 22 |
| 104 | ENSG00000100311 | 187.3629 | 1.183398 | 0.310673 | 3.809143 | 0.000139 | 0.002898 | 5.033505 | 7.935405 | 3.506356 | 10.37245 | PDGFB | | 22 |
| 105 | ENSG00000100678 | 171.4944 | 1.400365 | 0.31308 | 4.472861 | 7.72E-06 | 0.000213 | 2.083414 | 7.367235 | 3.020983 | 5.511617 | SLC8A3 | | 14 |
| 106 | ENSG00000100918 | 108.1414 | -1.64988 | 0.356097 | -4.63322 | 3.60E-06 | 0.000107 | 10.3923 | 3.133764 | 11.01755 | 3.35596 | REC8 | | 14 |
| 107 | ENSG00000101204 | 258.4473 | 1.235387 | 0.26632 | 4.638732 | 3.51E-06 | 0.000104 | 5.743257 | 10.85774 | 5.760184 | 14.85671 | CHRNA4 | | 20 |
| 108 | ENSG00000101335 | 188.9256 | 2.36028 | 0.339957 | 6.942886 | 3.84E-12 | 3.06E-10 | 3.482312 | 29.4142 | 6.66473 | 20.10302 | MYL9 | | 20 |
| 109 | ENSG00000101349 | 126.4906 | -1.30563 | 0.344677 | -3.78799 | 0.000152 | 0.003119 | 4.859902 | 1.915163 | 3.831809 | 1.456556 | PAK5 | | 20 |
| 110 | ENSG00000101438 | 17.99354 | -5.20253 | 1.286316 | -4.04452 | 5.24E-05 | 0.001222 | 1.621068 | 0.04982 | 1.686805 | 0.03706 | SLC32A1 | | 20 |
| 111 | ENSG00000101489 | 208.3435 | -1.76059 | 0.362471 | -4.85719 | 1.19E-06 | 3.91E-05 | 15.7137 | 6.99609 | 17.57316 | 2.594524 | CELF4 | | 18 |
| 112 | ENSG00000101542 | 440.406 | -4.4123 | 0.273035 | -16.1602 | 9.62E-59 | 1.23E-55 | 44.54524 | 1.590105 | 39.86829 | 2.0861 | CDH20 | | 18 |
| 113 | ENSG00000101638 | 41.98059 | -2.00485 | 0.588294 | -3.4079 | 0.000655 | 0.010778 | 0.954815 | 0.188614 | 0.844862 | 0.233558 | ST8SIA5 | | 18 |
| 114 | ENSG00000101825 | 270.4807 | 1.288447 | 0.242753 | 5.307634 | 1.11E-07 | 4.47E-06 | 1.622734 | 4.205026 | 1.949853 | 4.10218 | MXRA5 | X | |
| 115 | ENSG00000102174 | 129.7872 | -1.10236 | 0.34251 | -3.21848 | 0.001289 | 0.019028 | 4.588599 | 2.346338 | 4.096433 | 1.549365 | PHEX | X | |
| 116 | ENSG00000102265 | 63.38134 | 1.431531 | 0.471718 | 3.034716 | 0.002408 | 0.031781 | 5.845521 | 18.57233 | 9.443356 | 20.483 | TIMP1 | X | |
| 117 | ENSG00000102290 | 470.0442 | 2.054157 | 0.201588 | 10.18988 | 2.20E-24 | 5.16E-22 | 3.476053 | 15.43441 | 3.979231 | 14.03848 | PCDH11X | X | |
| 118 | ENSG00000102760 | 26.54304 | 2.668519 | 0.77677 | 3.435404 | 0.000592 | 0.009928 | 0.657367 | 5.313695 | 1.564002 | 8.060481 | RGCC | | 13 |
| 119 | ENSG00000102924 | 269.1018 | 1.22264 | 0.329948 | 3.70556 | 0.000211 | 0.004162 | 4.890048 | 18.56454 | 12.0857 | 19.00503 | CBLN1 | | 16 |
| 120 | ENSG00000103056 | 195.0122 | -1.36009 | 0.278993 | -4.875 | 1.09E-06 | 3.60E-05 | 6.911268 | 3.047858 | 8.137568 | 2.558355 | SMPD3 | | 16 |
| 121 | ENSG00000103154 | 183.4358 | 1.117506 | 0.29457 | 3.793685 | 0.000148 | 0.003065 | 14.45494 | 27.18037 | 16.10318 | 35.65029 | NECAB2 | | 16 |
| 122 | ENSG00000103449 | 803.3479 | -2.69548 | 0.197744 | -13.6312 | 2.61E-42 | 1.40E-39 | 33.83201 | 6.072669 | 33.57922 | 3.968919 | SALL1 | | 16 |
| 123 | ENSG00000103489 | 1285.202 | -1.69346 | 0.191656 | -8.83594 | 9.93E-19 | 1.63E-16 | 35.62253 | 10.14508 | 25.25199 | 7.885256 | XYLT1 | | 16 |
| 124 | ENSG00000104327 | 1982.401 | -3.32379 | 0.549011 | -6.05415 | 1.41E-09 | 7.83E-08 | 132.7792 | 23.15071 | 284.7924 | 16.53125 | CALB1 | | 8 |
| 125 | ENSG00000104332 | 675.1783 | -1.87148 | 0.204548 | -9.14934 | 5.73E-20 | 1.02E-17 | 33.1642 | 6.964827 | 25.27749 | 8.212172 | SFRP1 | | 8 |
| 126 | ENSG00000105281 | 419.1533 | 1.236942 | 0.230228 | 5.372675 | 7.76E-08 | 3.21E-06 | 11.3906 | 22.30093 | 12.35156 | 30.72289 | SLC1A5 | | 19 |
| 127 | ENSG00000105392 | 15.58575 | -3.7709 | 0.995258 | -3.78886 | 0.000151 | 0.003112 | 1.134479 | 0.087265 | 1.688882 | 0.107805 | CRX | | 19 |
| 128 | ENSG00000105792 | 79.01859 | -1.60148 | 0.458523 | -3.49269 | 0.000478 | 0.008336 | 7.980326 | 2.95341 | 5.737856 | 1.470532 | CFAP69 | | 7 |
| 129 | ENSG00000105810 | 3630.039 | 1.226147 | 0.121825 | 10.06483 | 7.90E-24 | 1.73E-21 | 22.35613 | 49.77606 | 22.36582 | 49.84246 | CDK6 | | 7 |
| 130 | ENSG00000105825 | 2048.794 | 2.403638 | 0.743892 | 3.231165 | 0.001233 | 0.018308 | 26.47436 | 204.3739 | 75.30434 | 303.5564 | TFPI2 | | 7 |
| 131 | ENSG00000105877 | 214.3462 | -2.40636 | 0.37016 | -6.50087 | 7.99E-11 | 5.41E-09 | 4.621726 | 1.410544 | 11.04448 | 1.38583 | DNAH11 | | 7 |
| 132 | ENSG00000105880 | 38.90753 | -8.64769 | 1.548358 | -5.58507 | 2.34E-08 | 1.08E-06 | 9.505473 | 0 | 6.899285 | 0 | DLX5 | | 7 |
| 133 | ENSG00000105939 | 213.4648 | 1.502645 | 0.322291 | 4.662389 | 3.13E-06 | 9.38E-05 | 1.436409 | 4.464111 | 2.572887 | 6.303593 | ZC3HAV1 | | 7 |
| 134 | ENSG00000105996 | 62.27511 | 2.083181 | 0.509766 | 4.086543 | 4.38E-05 | 0.001037 | 1.110763 | 8.069583 | 2.392271 | 6.098035 | HOXA2 | | 7 |
| 135 | ENSG00000106018 | 99.10735 | -1.6909 | 0.376294 | -4.49356 | 7.00E-06 | 0.000195 | 4.879066 | 1.673081 | 5.113497 | 1.297844 | VIPR2 | | 7 |
| 136 | ENSG00000106069 | 442.3357 | -1.28399 | 0.198029 | -6.48386 | 8.94E-11 | 6.04E-09 | 30.68783 | 12.67173 | 33.21728 | 12.31303 | CHN2 | | 7 |
| 137 | ENSG00000106236 | 302.4926 | -1.88543 | 0.258308 | -7.29914 | 2.90E-13 | 2.73E-11 | 23.01478 | 7.017889 | 21.04449 | 4.473961 | NPTX2 | | 7 |
| 138 | ENSG00000106538 | 136.3013 | 2.653944 | 0.382008 | 6.947352 | 3.72E-12 | 2.97E-10 | 7.941803 | 38.05311 | 9.177509 | 63.74113 | RARRES2 | | 7 |
| 139 | ENSG00000106689 | 148.4799 | -2.90892 | 0.371156 | -7.83746 | 4.60E-15 | 5.33E-13 | 21.26286 | 3.524645 | 19.18772 | 1.612923 | LHX2 | | 9 |
| 140 | ENSG00000106772 | 687.0313 | -1.1281 | 0.182775 | -6.17207 | 6.74E-10 | 3.99E-08 | 19.96437 | 8.11844 | 21.14592 | 9.739903 | PRUNE2 | | 9 |
| 141 | ENSG00000106853 | 34.09751 | -2.16734 | 0.655051 | -3.30866 | 0.000937 | 0.014621 | 5.462575 | 0.958978 | 7.491086 | 1.771472 | PTGR1 | | 9 |
| 142 | ENSG00000107242 | 178.0864 | 1.216706 | 0.388149 | 3.134635 | 0.001721 | 0.024273 | 3.234586 | 7.867472 | 6.988399 | 14.62345 | PIP5K1B | | 9 |
| 143 | ENSG00000107249 | 2220.561 | -2.03996 | 0.16507 | -12.3581 | 4.40E-35 | 1.70E-32 | 123.7446 | 32.64584 | 113.6993 | 22.67432 | GLIS3 | | 9 |
| 144 | ENSG00000107295 | 119.0012 | -1.06861 | 0.362769 | -2.94571 | 0.003222 | 0.0404 | 8.466749 | 4.018297 | 6.275334 | 2.736364 | SH3GL2 | | 9 |
| 145 | ENSG00000107338 | 146.8004 | -2.13827 | 0.333698 | -6.40779 | 1.48E-10 | 9.73E-09 | 4.875861 | 1.150417 | 4.082465 | 0.80732 | SHB | | 9 |
| 146 | ENSG00000107719 | 113.7834 | 1.135699 | 0.37049 | 3.065396 | 0.002174 | 0.02936 | 1.443484 | 3.183469 | 2.110627 | 4.210261 | PALD1 | | 10 |
| 147 | ENSG00000107731 | 178.2757 | -1.74129 | 0.293901 | -5.92475 | 3.13E-09 | 1.64E-07 | 4.49142 | 1.225591 | 5.011295 | 1.464554 | UNC5B | | 10 |
| 148 | ENSG00000107796 | 447.2371 | 2.086333 | 0.285306 | 7.31261 | 2.62E-13 | 2.48E-11 | 14.24692 | 98.61223 | 23.05905 | 53.09311 | ACTA2 | | 10 |
| 149 | ENSG00000108018 | 1062.848 | -3.04759 | 0.171991 | -17.7195 | 2.97E-70 | 4.76E-67 | 46.13076 | 5.757484 | 42.39417 | 4.508651 | SORCS1 | | 10 |
| 150 | ENSG00000108379 | 368.8278 | 4.083101 | 0.280619 | 14.55034 | 5.81E-48 | 4.47E-45 | 1.573195 | 23.67771 | 1.938712 | 32.57884 | WNT3 | | 17 |
| 151 | ENSG00000108439 | 14.50996 | -6.96696 | 1.700438 | -4.09716 | 4.18E-05 | 0.000995 | 1.272553 | 0 | 1.89237 | 0 | PNPO | | 17 |
| 152 | ENSG00000109099 | 106.6506 | 1.316235 | 0.35761 | 3.680641 | 0.000233 | 0.004509 | 4.456638 | 10.57041 | 4.458897 | 10.56146 | PMP22 | | 17 |
| 153 | ENSG00000109339 | 8485.596 | 1.728301 | 0.13353 | 12.94317 | 2.57E-38 | 1.10E-35 | 179.903 | 574.2196 | 211.3535 | 657.2995 | MAPK10 | | 4 |
| 154 | ENSG00000109436 | 869.2587 | -1.20581 | 0.16971 | -7.10516 | 1.20E-12 | 1.04E-10 | 25.23955 | 12.34342 | 29.78512 | 10.40866 | TBC1D9 | | 4 |
| 155 | ENSG00000109819 | 339.1997 | 1.211876 | 0.295322 | 4.103575 | 4.07E-05 | 0.000969 | 4.744957 | 12.68547 | 9.177797 | 17.68527 | PPARGC1A | | 4 |

| | | | | | | | | | | | | | |
|---|---|---|---|---|---|---|---|---|---|---|---|---|---|
| 156 | ENSG00000109911 | 827.5678 | -1.06258 | 0.16655 | -6.37998 | 1.77E-10 | 1.15E-08 | 33.24804 | 16.68618 | 33.66381 | 13.93276 | ELP4 | 11 |
| 157 | ENSG00000110060 | 117.6818 | 1.139937 | 0.347674 | 3.278752 | 0.001043 | 0.016015 | 5.740866 | 10.85274 | 5.96811 | 13.65883 | PUS3 | 11 |
| 158 | ENSG00000110148 | 64.75887 | -1.82563 | 0.569011 | -3.20842 | 0.001335 | 0.019549 | 2.927677 | 1.488804 | 9.953263 | 1.931047 | CCKBR | 11 |
| 159 | ENSG00000110675 | 85.0614 | 4.789084 | 0.588429 | 8.13876 | 3.99E-16 | 5.23E-14 | 0.134102 | 6.585261 | 0.553702 | 10.62765 | ELMOD1 | 11 |
| 160 | ENSG00000110693 | 823.2604 | -2.03306 | 0.171453 | -11.8578 | 1.96E-32 | 6.86E-30 | 33.38974 | 8.381065 | 32.32633 | 6.952617 | SOX6 | 11 |
| 161 | ENSG00000111057 | 57.15962 | 2.049294 | 0.522978 | 3.918511 | 8.91E-05 | 0.001953 | 1.301615 | 8.408492 | 2.858043 | 7.988378 | KRT18 | 12 |
| 162 | ENSG00000111186 | 134.2312 | -2.27641 | 0.339549 | -6.70424 | 2.02E-11 | 1.49E-09 | 17.96678 | 4.179142 | 18.45473 | 3.067175 | WNT5B | 12 |
| 163 | ENSG00000111432 | 517.2885 | 5.131637 | 0.276885 | 18.53346 | 1.11E-76 | 2.13E-73 | 0.769946 | 37.87559 | 1.281825 | 30.98467 | FZD10 | 12 |
| 164 | ENSG00000111783 | 2031.205 | 1.751802 | 0.1333 | 13.14178 | 1.90E-39 | 8.91E-37 | 32.09066 | 109.4862 | 33.76066 | 101.8054 | RFX4 | 12 |
| 165 | ENSG00000111885 | 145.3177 | -1.04682 | 0.340809 | -3.07156 | 0.002129 | 0.0288 | 4.082225 | 1.563953 | 4.860032 | 2.512472 | MAN1A1 | 6 |
| 166 | ENSG00000111912 | 122.3991 | -1.14214 | 0.368043 | -3.10327 | 0.001914 | 0.026387 | 4.546941 | 3.00356 | 7.570227 | 2.2671 | NCOA7 | 6 |
| 167 | ENSG00000112175 | 72.10625 | 5.492827 | 0.730774 | 7.516448 | 5.63E-14 | 5.61E-12 | 0.093985 | 3.052138 | 0.089409 | 4.754562 | BMP5 | 6 |
| 168 | ENSG00000112232 | 230.5383 | -1.13145 | 0.305575 | -3.70269 | 0.000213 | 0.004194 | 20.62395 | 8.108664 | 12.22534 | 6.269503 | KHDRBS2 | 6 |
| 169 | ENSG00000112280 | 175.9929 | 1.382176 | 0.336226 | 4.110858 | 3.94E-05 | 0.000945 | 3.232351 | 13.00777 | 5.186607 | 8.036407 | COL9A1 | 6 |
| 170 | ENSG00000112333 | 371.0562 | -5.80709 | 0.970997 | -5.98054 | 2.22E-09 | 1.19E-07 | 36.58512 | 0.727554 | 16.44717 | 0.198997 | NR2E1 | 6 |
| 171 | ENSG00000112414 | 78.57986 | 1.419858 | 0.421489 | 3.368675 | 0.000755 | 0.012175 | 1.743347 | 4.401316 | 1.58237 | 4.04444 | ADGRG6 | 6 |
| 172 | ENSG00000112773 | 1177.847 | 3.147278 | 0.193515 | 16.26373 | 1.79E-59 | 2.64E-56 | 3.80846 | 38.62403 | 5.913238 | 43.19455 | TENT5A | 6 |
| 173 | ENSG00000112902 | 536.3346 | 2.576287 | 0.227714 | 11.31369 | 1.12E-29 | 3.43E-27 | 1.21701 | 9.891339 | 1.979029 | 8.438928 | SEMA5A | 5 |
| 174 | ENSG00000113209 | 132.0909 | -1.91983 | 0.33118 | -5.79693 | 6.75E-09 | 3.36E-07 | 6.896138 | 1.777921 | 7.472605 | 1.8311 | PCDHB5 | 5 |
| 175 | ENSG00000113248 | 144.7308 | -1.27173 | 0.311668 | -4.0804 | 4.50E-05 | 0.00106 | 6.326628 | 2.485951 | 5.869596 | 2.330935 | PCDHB15 | 5 |
| 176 | ENSG00000113361 | 1151.427 | -3.04542 | 0.187375 | -16.2531 | 2.12E-59 | 2.92E-56 | 38.75002 | 6.006119 | 51.2575 | 4.444924 | CDH6 | 5 |
| 177 | ENSG00000113494 | 35.44278 | -2.0552 | 0.619771 | -3.31606 | 0.000913 | 0.014297 | 1.262986 | 0.272984 | 1.603635 | 0.370245 | PRLR | 5 |
| 178 | ENSG00000113739 | 360.2725 | 3.15169 | 0.657762 | 4.791537 | 1.66E-06 | 5.28E-05 | 1.309817 | 17.13666 | 3.430937 | 22.63101 | STC2 | 5 |
| 179 | ENSG00000113805 | 159.8459 | 3.350114 | 0.407441 | 8.222328 | 2.00E-16 | 2.78E-14 | 0.545789 | 11.39983 | 1.700452 | 9.167207 | CNTN3 | 3 |
| 180 | ENSG00000114279 | 310.4241 | 1.306563 | 0.269606 | 4.846188 | 1.26E-06 | 4.11E-05 | 5.173598 | 18.11197 | 9.006244 | 15.32277 | FGF12 | 3 |
| 181 | ENSG00000114541 | 472.018 | -1.52771 | 0.21288 | -7.17639 | 7.16E-13 | 6.41E-11 | 36.53571 | 10.06816 | 30.76123 | 12.09509 | FRMD4B | 3 |
| 182 | ENSG00000114646 | 142.7149 | 1.004992 | 0.334504 | 3.004422 | 0.002661 | 0.034594 | 5.499173 | 9.885201 | 3.763436 | 7.855786 | CSPG5 | 3 |
| 183 | ENSG00000114654 | 163.558 | 2.346372 | 0.310786 | 7.54979 | 4.36E-14 | 4.44E-12 | 2.28669 | 12.08603 | 2.591719 | 11.54387 | EFCC1 | 3 |
| 184 | ENSG00000115252 | 312.4239 | 1.038091 | 0.25687 | 4.041312 | 5.32E-05 | 0.001235 | 8.661639 | 14.18621 | 9.156775 | 20.51541 | PDE1A | 2 |
| 185 | ENSG00000115414 | 8982.883 | -2.20832 | 0.139856 | -15.79 | 3.65E-56 | 4.39E-53 | 296.5112 | 72.73923 | 315.3868 | 53.72217 | FN1 | 2 |
| 186 | ENSG00000115461 | 1302.602 | 2.986273 | 0.207275 | 14.40727 | 4.66E-47 | 3.20E-44 | 3.903665 | 41.78711 | 7.03734 | 40.69477 | IGFBP5 | 2 |
| 187 | ENSG00000115738 | 520.6035 | 1.710769 | 0.215849 | 7.925768 | 2.27E-15 | 2.78E-13 | 21.94756 | 76.87751 | 31.16821 | 88.09803 | ID2 | 2 |
| 188 | ENSG00000115844 | 67.47503 | -7.05888 | 1.11854 | -6.3108 | 2.78E-10 | 1.73E-08 | 8.768549 | 0 | 5.713079 | 0.09204 | DLX2 | 2 |
| 189 | ENSG00000115896 | 146.9949 | -1.26393 | 0.323308 | -3.90936 | 9.25E-05 | 0.002008 | 4.489631 | 2.348613 | 5.59177 | 1.68206 | PLCL1 | 2 |
| 190 | ENSG00000116016 | 327.0433 | 4.156659 | 0.713955 | 5.822019 | 5.81E-09 | 2.92E-07 | 0.587161 | 13.17855 | 1.509858 | 22.10295 | EPAS1 | 2 |
| 191 | ENSG00000116117 | 441.9006 | -1.18068 | 0.223291 | -5.28764 | 1.24E-07 | 4.92E-06 | 13.50043 | 6.759729 | 13.11162 | 4.51923 | PARD3B | 2 |
| 192 | ENSG00000116183 | 63.62657 | -2.29653 | 0.498882 | -4.60334 | 4.16E-06 | 0.000121 | 1.555296 | 0.332145 | 2.753745 | 0.478517 | PAPPA2 | 1 |
| 193 | ENSG00000116194 | 179.1098 | -2.09184 | 0.296407 | -7.05734 | 1.70E-12 | 1.44E-10 | 9.948512 | 2.716036 | 12.19409 | 2.247694 | ANGPTL1 | 1 |
| 194 | ENSG00000116745 | 24.58489 | 2.650119 | 0.800048 | 3.312453 | 0.000925 | 0.014447 | 0.197904 | 2.335649 | 0.423624 | 1.411386 | RPE65 | 1 |
| 195 | ENSG00000117009 | 18.87319 | -3.52401 | 1.009446 | -3.49103 | 0.000481 | 0.008373 | 1.429292 | 0.098938 | 0.406391 | 0.05496 | KMO | 1 |
| 196 | ENSG00000117069 | 428.6427 | -1.9984 | 0.267733 | -7.46413 | 8.39E-14 | 8.23E-12 | 14.4124 | 4.212408 | 25.60546 | 5.256296 | ST6GALNA | 1 |
| 197 | ENSG00000117298 | 480.994 | 1.446109 | 0.254444 | 5.683401 | 1.32E-08 | 6.23E-07 | 5.359246 | 15.84472 | 9.063142 | 21.31866 | ECE1 | 1 |
| 198 | ENSG00000117318 | 157.6517 | 1.852077 | 0.41758 | 4.435263 | 9.20E-06 | 0.00025 | 5.223291 | 32.04444 | 16.74099 | 43.15467 | ID3 | 1 |
| 199 | ENSG00000117394 | 4474.002 | 1.132694 | 0.147288 | 7.69036 | 1.47E-14 | 1.59E-12 | 142.5874 | 351.2783 | 147.1049 | 255.5679 | SLC2A1 | 1 |
| 200 | ENSG00000117643 | 134.1244 | 1.248197 | 0.4155 | 3.004084 | 0.002664 | 0.034594 | 1.500493 | 4.920255 | 3.891197 | 7.110251 | MAN1C1 | 1 |
| 201 | ENSG00000117971 | 101.4246 | -1.08253 | 0.361617 | -2.99359 | 0.002757 | 0.03559 | 6.796983 | 3.141064 | 6.691405 | 2.930023 | CHRNB4 | 15 |
| 202 | ENSG00000118160 | 75.16946 | -1.24972 | 0.432008 | -2.89282 | 0.003818 | 0.045897 | 2.788161 | 1.489973 | 4.199981 | 1.320257 | SLC8A2 | 19 |
| 203 | ENSG00000118257 | 986.3568 | -1.29754 | 0.209028 | -6.2075 | 5.38E-10 | 3.22E-08 | 42.44584 | 14.50828 | 52.4471 | 22.2664 | NRP2 | 2 |
| 204 | ENSG00000118473 | 404.4242 | -1.19896 | 0.203758 | -5.88424 | 4.00E-09 | 2.04E-07 | 17.76175 | 7.809105 | 18.79796 | 7.361362 | SGIP1 | 1 |
| 205 | ENSG00000118495 | 1041.162 | -1.00532 | 0.164038 | -6.12859 | 8.87E-10 | 5.17E-08 | 61.0406 | 29.10736 | 52.76025 | 25.04871 | PLAGL1 | 6 |
| 206 | ENSG00000118503 | 100.1154 | 1.303945 | 0.409125 | 3.187152 | 0.001437 | 0.020823 | 1.904984 | 6.502299 | 3.838504 | 6.892795 | TNFAIP3 | 6 |
| 207 | ENSG00000118508 | 79.32719 | 1.943974 | 0.42991 | 4.521818 | 6.13E-06 | 0.000172 | 3.570071 | 17.36201 | 5.163958 | 14.70693 | RAB32 | 6 |
| 208 | ENSG00000119408 | 519.3927 | -1.26713 | 0.208394 | -6.08045 | 1.20E-09 | 6.78E-08 | 27.18027 | 12.12812 | 24.92273 | 8.637557 | NEK6 | 9 |
| 209 | ENSG00000119514 | 206.1248 | -1.29717 | 0.304679 | -4.25748 | 2.07E-05 | 0.000526 | 10.3501 | 5.319362 | 17.41746 | 5.405276 | GALNT12 | 9 |
| 210 | ENSG00000119866 | 974.6147 | -1.02012 | 0.161023 | -6.33528 | 2.37E-10 | 1.50E-08 | 39.7287 | 21.40635 | 46.11813 | 18.91664 | BCL11A | 2 |
| 211 | ENSG00000120149 | 119.9974 | 5.254659 | 0.545244 | 9.637265 | 5.57E-22 | 1.09E-19 | 0.376727 | 13.35445 | 0.398902 | 14.87365 | MSX2 | 5 |
| 212 | ENSG00000120156 | 31.86974 | 2.265316 | 0.731065 | 3.098653 | 0.001944 | 0.026687 | 0.184746 | 1.860673 | 0.373279 | 0.719618 | TEK | 9 |
| 213 | ENSG00000120162 | 709.7547 | -1.38296 | 0.202457 | -6.8309 | 8.44E-12 | 6.47E-10 | 21.94211 | 7.288009 | 15.73116 | 6.518782 | MOB3B | 9 |
| 214 | ENSG00000120251 | 194.4522 | -1.40313 | 0.351163 | -3.99565 | 6.45E-05 | 0.00148 | 10.3946 | 6.594513 | 15.87897 | 3.120884 | GRIA2 | 4 |
| 215 | ENSG00000120322 | 54.76465 | -1.56949 | 0.527075 | -2.97774 | 0.002904 | 0.037085 | 3.635422 | 1.604506 | 3.45856 | 0.715642 | PCDHB8 | 5 |
| 216 | ENSG00000120907 | 14.32291 | -4.70994 | 1.213318 | -3.88187 | 0.000104 | 0.002227 | 1.354252 | 0.078954 | 1.23736 | 0.024163 | ADRA1A | 8 |
| 217 | ENSG00000121207 | 172.0698 | -1.67509 | 0.371503 | -4.50896 | 6.51E-06 | 0.000182 | 4.148049 | 2.152364 | 9.35614 | 1.877308 | LRAT | 4 |
| 218 | ENSG00000121871 | 101.8998 | -1.17696 | 0.370498 | -3.1767 | 0.00149 | 0.021411 | 3.138227 | 1.647408 | 4.176178 | 1.4321 | SLITRK3 | 3 |
| 219 | ENSG00000121966 | 1060.987 | 2.974718 | 0.200184 | 14.8599 | 6.00E-50 | 5.50E-47 | 13.71683 | 140.0472 | 22.7905 | 133.0592 | CXCR4 | 2 |
| 220 | ENSG00000121989 | 1536.862 | -1.79255 | 0.181349 | -9.88457 | 4.86E-23 | 1.02E-20 | 102.6725 | 28.05663 | 76.72561 | 21.53675 | ACVR2A | 2 |
| 221 | ENSG00000122644 | 1347.769 | 1.08643 | 0.151642 | 7.164461 | 7.81E-13 | 6.93E-11 | 58.81572 | 124.8098 | 54.75251 | 105.4022 | ARL4A | 7 |
| 222 | ENSG00000122735 | 27.93971 | -2.29326 | 0.740926 | -3.09513 | 0.001967 | 0.02691 | 5.828422 | 0.539251 | 5.102311 | 1.508924 | DNAI1 | 9 |
| 223 | ENSG00000123104 | 244.4031 | 1.305408 | 0.255589 | 5.107456 | 3.27E-07 | 1.18E-05 | 1.892559 | 5.1146 | 2.33976 | 4.846081 | ITPR2 | 12 |
| 224 | ENSG00000123360 | 227.0992 | 1.076444 | 0.272661 | 3.947923 | 7.88E-05 | 0.001754 | 4.27893 | 10.77611 | 6.237253 | 10.33854 | PDE1B | 12 |
| 225 | ENSG00000123496 | 46.54278 | 3.245331 | 0.645675 | 5.02626 | 5.00E-07 | 1.75E-05 | 0.642104 | 5.35766 | 1.120117 | 10.4147 | IL13RA2 | X |
| 226 | ENSG00000124126 | 821.7553 | 3.105776 | 0.226382 | 13.71916 | 7.80E-43 | 4.41E-40 | 2.42214 | 25.812 | 4.365642 | 29.9998 | PREX1 | 20 |
| 227 | ENSG00000124140 | 55.57676 | -1.72131 | 0.51071 | -3.37043 | 0.000751 | 0.012118 | 11.29013 | 2.213181 | 7.420191 | 3.140902 | SLC12A5 | 20 |
| 228 | ENSG00000124785 | 148.0186 | -1.07721 | 0.309012 | -3.48597 | 0.00049 | 0.00851 | 17.26756 | 7.77659 | 15.32929 | 6.970897 | NRN1 | 6 |
| 229 | ENSG00000124788 | 369.5859 | -1.31996 | 0.23001 | -5.73873 | 9.54E-09 | 4.60E-07 | 15.56896 | 4.906415 | 12.77844 | 5.86892 | ATXN1 | 6 |
| 230 | ENSG00000125046 | 24.76468 | 3.629063 | 0.844836 | 4.295583 | 1.74E-05 | 0.000454 | 0.163636 | 2.547584 | 0.314341 | 2.988811 | SSUH2 | 3 |
| 231 | ENSG00000125246 | 290.0171 | 2.780101 | 0.257846 | 10.782 | 4.19E-27 | 1.15E-24 | 4.184636 | 24.67618 | 3.969494 | 28.23558 | CLYBL | 13 |
| 232 | ENSG00000125430 | 726.6575 | 1.451814 | 0.183453 | 7.913814 | 2.50E-15 | 3.00E-13 | 8.219542 | 24.8777 | 8.615709 | 19.11539 | HS3ST3B1 | 17 |
| 233 | ENSG00000125462 | 60.17981 | -1.86698 | 0.520114 | -3.58956 | 0.000331 | 0.006089 | 4.28961 | 1.585681 | 8.667781 | 1.764223 | MIR9-1HG | 1 |

| | | | | | | | | | | | | | | |
|---|---|---|---|---|---|---|---|---|---|---|---|---|---|---|
| 234 | ENSG00000125804 | 110.1257 | -1.58249 | 0.421611 | -3.75343 | 0.000174 | 0.003533 | 10.44567 | 3.076631 | 5.302479 | 2.016007 | FAM182A | | 20 |
| 235 | ENSG00000125851 | 144.1497 | -1.10512 | 0.324475 | -3.40587 | 0.00066 | 0.010849 | 7.148046 | 2.582231 | 5.733768 | 3.087976 | PCSK2 | | 20 |
| 236 | ENSG00000126259 | 157.3953 | 1.439059 | 0.312815 | 4.600354 | 4.22E-06 | 0.000122 | 2.997231 | 8.449903 | 3.98051 | 9.512346 | KIRREL2 | | 19 |
| 237 | ENSG00000126500 | 107.0546 | -1.21414 | 0.38302 | -3.16991 | 0.001525 | 0.021852 | 3.960175 | 1.334793 | 4.670502 | 2.168985 | FLRT1 | | 11 |
| 238 | ENSG00000126583 | 119.7834 | 2.198142 | 0.411365 | 5.343536 | 9.12E-08 | 3.72E-06 | 1.244895 | 7.307778 | 2.802745 | 10.19336 | PRKCG | | 19 |
| 239 | ENSG00000127124 | 142.3201 | -1.36501 | 0.331426 | -4.11859 | 3.81E-05 | 0.000918 | 3.021506 | 1.283475 | 4.346182 | 1.426741 | HIVEP3 | | 1 |
| 240 | ENSG00000127241 | 143.0645 | -1.28255 | 0.383228 | -3.34669 | 0.000818 | 0.012976 | 6.995521 | 4.986118 | 14.66235 | 3.465923 | MASP1 | | 3 |
| 241 | ENSG00000127399 | 23.56466 | -2.29886 | 0.771965 | -2.97793 | 0.002902 | 0.037085 | 2.902714 | 0.47409 | 3.044976 | 0.656357 | LRRC61 | | 7 |
| 242 | ENSG00000127920 | 1689.721 | 2.858789 | 0.710683 | 4.022596 | 5.76E-05 | 0.001331 | 8.163997 | 102.5013 | 24.14088 | 118.9728 | GNG11 | | 7 |
| 243 | ENSG00000128045 | 266.5947 | 1.707932 | 0.270396 | 6.31641 | 2.68E-10 | 1.68E-08 | 6.67388 | 28.65364 | 10.60052 | 24.97633 | RASL11B | | 4 |
| 244 | ENSG00000128052 | 85.94846 | -1.53189 | 0.463944 | -3.30189 | 0.00096 | 0.014942 | 2.100716 | 0.612722 | 3.631849 | 1.247303 | KDR | | 4 |
| 245 | ENSG00000128165 | 42.68531 | 2.141623 | 0.633384 | 3.381242 | 0.000722 | 0.01171 | 0.499937 | 1.070846 | 0.363688 | 2.512666 | ADM2 | | 22 |
| 246 | ENSG00000128573 | 220.9522 | -1.43027 | 0.266454 | -5.3678 | 7.97E-08 | 3.29E-06 | 8.087085 | 3.376715 | 9.803393 | 2.952879 | FOXP2 | | 7 |
| 247 | ENSG00000128610 | 1061.729 | -4.82373 | 0.205956 | -23.4212 | 2.60E-121 | 1.55E-117 | 135.6413 | 6.084362 | 158.6297 | 3.913891 | FEZF1 | | 7 |
| 248 | ENSG00000128683 | 81.86537 | -2.78107 | 0.454745 | -6.11566 | 9.62E-10 | 5.57E-08 | 7.767511 | 1.405521 | 7.014173 | 0.767252 | GAD1 | | 2 |
| 249 | ENSG00000128965 | 61.33308 | 1.826018 | 0.581963 | 3.137691 | 0.001703 | 0.024045 | 2.054324 | 3.929155 | 2.851469 | 12.33627 | CHAC1 | | 15 |
| 250 | ENSG00000130226 | 395.4136 | -1.40101 | 0.213919 | -6.54928 | 5.78E-11 | 3.96E-09 | 19.35137 | 6.418136 | 19.44138 | 7.52414 | DPP6 | | 7 |
| 251 | ENSG00000130429 | 70.19439 | 1.291568 | 0.442459 | 2.919065 | 0.003511 | 0.043175 | 4.509468 | 11.06769 | 6.025215 | 13.41392 | ARPC1B | | 7 |
| 252 | ENSG00000130707 | 205.6696 | 2.170362 | 0.343201 | 6.323878 | 2.55E-10 | 1.61E-08 | 4.186644 | 30.98461 | 10.09144 | 29.74628 | ASS1 | | 9 |
| 253 | ENSG00000130940 | 371.5053 | 1.790826 | 0.219853 | 8.145556 | 3.78E-16 | 4.98E-14 | 4.328693 | 16.31588 | 5.177235 | 15.07555 | CASZ1 | | 1 |
| 254 | ENSG00000131019 | 337.472 | 1.402593 | 0.248497 | 5.644302 | 1.66E-08 | 7.75E-07 | 6.476054 | 17.14961 | 8.985303 | 21.44557 | ULBP3 | | 6 |
| 255 | ENSG00000131370 | 210.6153 | -1.15336 | 0.289816 | -3.97963 | 6.90E-05 | 0.001563 | 11.80305 | 4.125718 | 12.39503 | 6.175408 | SH3BP5 | | 3 |
| 256 | ENSG00000131378 | 276.0736 | -1.89079 | 0.266731 | -7.08874 | 1.35E-12 | 1.16E-10 | 18.85653 | 6.333245 | 20.78422 | 3.963134 | RFTN1 | | 3 |
| 257 | ENSG00000132561 | 90.85223 | 1.834838 | 0.428302 | 4.283984 | 1.84E-05 | 0.000476 | 1.558746 | 8.672293 | 2.918296 | 6.627783 | MATN2 | | 8 |
| 258 | ENSG00000132718 | 2549.638 | -1.48624 | 0.137718 | -10.7919 | 3.76E-27 | 1.05E-24 | 83.08963 | 26.28559 | 82.3467 | 29.83316 | SYT11 | | 1 |
| 259 | ENSG00000133101 | 26.01179 | -2.22469 | 0.727235 | -3.0591 | 0.00222 | 0.029816 | 3.0929 | 0.781855 | 3.21973 | 0.520164 | CCNA1 | | 13 |
| 260 | ENSG00000133124 | 3183.014 | -3.18839 | 0.410739 | -7.76257 | 8.32E-15 | 9.26E-13 | 52.18055 | 7.926298 | 65.10682 | 4.387561 | IRS4 | X | |
| 261 | ENSG00000133424 | 722.3077 | 1.088541 | 0.171835 | 6.334814 | 2.38E-10 | 1.50E-08 | 13.77582 | 31.40985 | 15.39942 | 27.73408 | LARGE1 | | 22 |
| 262 | ENSG00000133710 | 59.35173 | -1.49566 | 0.49432 | -3.0257 | 0.002481 | 0.032632 | 3.399683 | 1.371208 | 2.831115 | 0.772482 | SPINK5 | | 5 |
| 263 | ENSG00000133958 | 305.4019 | -1.04786 | 0.233587 | -4.48594 | 7.26E-06 | 0.000201 | 5.852874 | 3.001619 | 5.71399 | 2.348528 | UNC79 | | 14 |
| 264 | ENSG00000134115 | 131.1679 | -1.18545 | 0.362963 | -3.26603 | 0.001091 | 0.016554 | 6.774607 | 3.642508 | 6.266491 | 1.920132 | CNTN6 | | 3 |
| 265 | ENSG00000134198 | 123.3228 | -1.30097 | 0.339756 | -3.82913 | 0.000129 | 0.002711 | 7.386274 | 3.192659 | 7.061537 | 2.426077 | TSPAN2 | | 1 |
| 266 | ENSG00000134245 | 1073.859 | 1.270167 | 0.161925 | 7.844185 | 4.36E-15 | 5.11E-13 | 7.964114 | 21.37029 | 9.00629 | 17.68362 | WNT2B | | 1 |
| 267 | ENSG00000134330 | 63.20188 | -1.54343 | 0.48965 | -3.15211 | 0.001621 | 0.023041 | 5.139069 | 1.254689 | 6.171523 | 2.383975 | IAH1 | | 2 |
| 268 | ENSG00000134363 | 88.50251 | 1.423676 | 0.442239 | 3.219246 | 0.001285 | 0.019028 | 1.76265 | 6.514599 | 3.924655 | 7.864615 | FST | | 5 |
| 269 | ENSG00000134376 | 198.9163 | -1.29533 | 0.270393 | -4.79055 | 1.66E-06 | 5.30E-05 | 7.071334 | 2.87265 | 7.489189 | 2.775797 | CRB1 | | 1 |
| 270 | ENSG00000134504 | 644.4623 | 1.666716 | 0.176239 | 9.457133 | 3.17E-21 | 6.03E-19 | 14.97353 | 47.12957 | 16.02246 | 46.56086 | KCTD1 | | 18 |
| 271 | ENSG00000134548 | 18.11147 | 3.286862 | 0.94555 | 3.476137 | 0.000509 | 0.008798 | 0.243135 | 2.063059 | 0.231994 | 2.355691 | SPX | | 12 |
| 272 | ENSG00000134569 | 669.1939 | 1.026179 | 0.184381 | 5.565545 | 2.61E-08 | 1.20E-06 | 8.280147 | 16.87176 | 10.08027 | 18.67578 | LRP4 | | 11 |
| 273 | ENSG00000134595 | 872.6391 | -1.08552 | 0.209404 | -5.18387 | 2.17E-07 | 8.17E-06 | 83.9915 | 35.07389 | 56.05125 | 28.10173 | SOX3 | X | |
| 274 | ENSG00000134769 | 382.3504 | -2.24237 | 0.264971 | -8.46273 | 2.61E-17 | 3.93E-15 | 20.71875 | 6.06076 | 24.66844 | 3.248174 | DTNA | | 18 |
| 275 | ENSG00000134775 | 867.1332 | -1.34467 | 0.171324 | -7.84868 | 4.20E-15 | 4.96E-13 | 37.92497 | 14.5854 | 33.43002 | 12.2688 | FHOD3 | | 18 |
| 276 | ENSG00000134954 | 137.0995 | 2.051886 | 0.452463 | 4.534925 | 5.76E-06 | 0.000163 | 0.83201 | 5.413076 | 2.825628 | 8.478997 | ETS1 | | 11 |
| 277 | ENSG00000135046 | 177.5792 | 2.026436 | 0.337014 | 6.01291 | 1.82E-09 | 9.93E-08 | 4.561382 | 24.38963 | 8.979654 | 28.08778 | ANXA1 | | 9 |
| 278 | ENSG00000135299 | 1018.574 | -1.04821 | 0.173744 | -6.03308 | 1.61E-09 | 8.82E-08 | 60.54741 | 31.51685 | 57.49475 | 23.16282 | ANKRD6 | | 6 |
| 279 | ENSG00000135363 | 57.11677 | -3.15167 | 0.532341 | -5.92041 | 3.21E-09 | 1.68E-07 | 7.379977 | 0.902682 | 9.408237 | 0.893144 | LMO2 | | 11 |
| 280 | ENSG00000135525 | 208.5589 | -1.71825 | 0.269983 | -6.3643 | 1.96E-10 | 1.26E-08 | 11.10464 | 3.104116 | 11.12015 | 3.312589 | MAP7 | | 6 |
| 281 | ENSG00000135842 | 80.69702 | 2.020391 | 0.427507 | 4.725988 | 2.29E-06 | 7.10E-05 | 0.652568 | 2.616958 | 0.791058 | 2.905323 | NIBAN1 | | 1 |
| 282 | ENSG00000135903 | 1297.407 | 2.405496 | 0.170106 | 14.14113 | 2.12E-45 | 1.41E-42 | 16.96069 | 103.7707 | 23.56907 | 100.5411 | PAX3 | | 2 |
| 283 | ENSG00000136014 | 351.6785 | -1.20744 | 0.24833 | -4.86224 | 1.16E-06 | 3.82E-05 | 18.62179 | 8.645718 | 27.5725 | 10.26756 | USP44 | | 12 |
| 284 | ENSG00000136099 | 320.2374 | 1.784869 | 0.323115 | 5.523947 | 3.31E-08 | 1.49E-06 | 4.661668 | 19.7899 | 10.48366 | 29.20757 | PCDH8 | | 13 |
| 285 | ENSG00000136160 | 1092.883 | 2.328615 | 0.201562 | 11.55285 | 7.14E-31 | 2.25E-28 | 8.175931 | 44.67859 | 12.62056 | 54.38529 | EDNRB | | 13 |
| 286 | ENSG00000136267 | 99.57436 | -1.88514 | 0.388684 | -4.85006 | 1.23E-06 | 4.03E-05 | 6.437405 | 2.042817 | 9.264946 | 1.996057 | DGKB | | 7 |
| 287 | ENSG00000136531 | 239.7121 | -1.48323 | 0.306569 | -4.83817 | 1.31E-06 | 4.27E-05 | 3.500968 | 1.927377 | 5.580656 | 1.190071 | SCN2A | | 2 |
| 288 | ENSG00000136535 | 577.708 | -4.04097 | 0.787115 | -5.1339 | 2.84E-07 | 1.04E-05 | 31.95536 | 2.676248 | 26.53906 | 0.76139 | TBR1 | | 2 |
| 289 | ENSG00000136732 | 161.4565 | 2.146784 | 0.309415 | 6.93821 | 3.97E-12 | 3.13E-10 | 7.671662 | 35.44788 | 8.949835 | 34.61223 | GYPC | | 2 |
| 290 | ENSG00000136750 | 78.55957 | -4.81806 | 1.343975 | -3.58493 | 0.000337 | 0.006186 | 5.916145 | 0.392149 | 5.587307 | 0.033045 | GAD2 | | 10 |
| 291 | ENSG00000136770 | 303.5213 | 1.397149 | 0.26633 | 5.245942 | 1.55E-07 | 6.04E-06 | 7.820779 | 22.62787 | 12.3231 | 27.6868 | DNAJC1 | | 10 |
| 292 | ENSG00000136943 | 110.5648 | 1.085742 | 0.360485 | 3.011894 | 0.002596 | 0.033968 | 4.718658 | 8.486415 | 4.96704 | 10.99988 | CTSV | | 9 |
| 293 | ENSG00000136944 | 178.0778 | 1.65902 | 0.317283 | 5.228842 | 1.71E-07 | 6.54E-06 | 1.331325 | 5.615115 | 1.741794 | 3.677827 | LMX1B | | 9 |
| 294 | ENSG00000136960 | 591.9854 | 1.084251 | 0.210031 | 5.162346 | 2.44E-07 | 9.08E-06 | 14.01314 | 30.45846 | 19.43904 | 36.73581 | ENPP2 | | 8 |
| 295 | ENSG00000137142 | 8719.8 | 1.565299 | 0.141205 | 11.08532 | 1.48E-28 | 4.31E-26 | 143.913 | 359.3536 | 144.5297 | 451.5639 | IGFBPL1 | | 9 |
| 296 | ENSG00000137203 | 129.6337 | -1.40333 | 0.412449 | -3.40243 | 0.000668 | 0.010968 | 12.10084 | 5.774727 | 9.602866 | 2.154886 | TFAP2A | | 6 |
| 297 | ENSG00000137573 | 150.1559 | -1.19495 | 0.319287 | -3.74255 | 0.000182 | 0.003671 | 8.90643 | 3.923315 | 7.44012 | 2.890565 | SULF1 | | 8 |
| 298 | ENSG00000137642 | 674.8709 | -1.37714 | 0.173837 | -7.92204 | 2.34E-15 | 2.85E-13 | 15.76843 | 5.894557 | 17.05216 | 6.117501 | SORL1 | | 11 |
| 299 | ENSG00000137691 | 9.820765 | -6.56813 | 1.80961 | -3.62958 | 0.000284 | 0.00532 | 1.901996 | 0 | 2.730223 | 0 | CFAP300 | | 11 |
| 300 | ENSG00000137871 | 169.1244 | -2.69622 | 0.316277 | -8.52488 | 1.53E-17 | 2.34E-15 | 10.95456 | 1.878877 | 10.65876 | 1.364564 | ZNF280D | | 15 |
| 301 | ENSG00000137936 | 236.5747 | -1.18269 | 0.267891 | -4.41484 | 1.01E-05 | 0.000271 | 12.53893 | 5.564271 | 16.49211 | 6.565091 | BCAR3 | | 1 |
| 302 | ENSG00000137962 | 720.3795 | 3.134631 | 0.223917 | 13.99907 | 1.58E-44 | 9.81E-42 | 1.968446 | 21.44502 | 3.435389 | 23.57878 | ARHGAP29 | | 1 |
| 303 | ENSG00000138061 | 39.9355 | 2.36148 | 0.686602 | 3.439374 | 0.000583 | 0.009826 | 0.563217 | 1.431778 | 0.147764 | 2.477702 | CYP1B1 | | 2 |
| 304 | ENSG00000138083 | 244.6051 | -4.19168 | 0.31323 | -13.3821 | 7.69E-41 | 3.90E-38 | 20.38964 | 1.302847 | 20.47788 | 0.864829 | SIX3 | | 2 |
| 305 | ENSG00000138411 | 775.125 | -1.93116 | 0.20229 | -9.54653 | 1.34E-21 | 2.61E-19 | 22.99112 | 6.433148 | 19.67501 | 4.303832 | HECW2 | | 2 |
| 306 | ENSG00000138448 | 1153.116 | -1.24113 | 0.16798 | -7.38856 | 1.48E-13 | 1.42E-11 | 29.61657 | 12.43938 | 36.82905 | 14.2392 | ITGAV | | 2 |
| 307 | ENSG00000138653 | 364.8528 | -2.38784 | 0.23104 | -10.3352 | 4.88E-25 | 1.18E-22 | 24.20269 | 5.182297 | 24.75712 | 3.798656 | NDST4 | | 4 |
| 308 | ENSG00000138670 | 87.00437 | -2.04872 | 0.410338 | -4.99275 | 5.95E-07 | 2.08E-05 | 6.241645 | 1.5911 | 6.58155 | 1.364098 | RASGEF1B | | 4 |
| 309 | ENSG00000138696 | 850.8394 | 1.869955 | 0.196955 | 9.494312 | 2.22E-21 | 4.27E-19 | 11.37333 | 32.11511 | 8.038485 | 35.46522 | BMPR1B | | 4 |
| 310 | ENSG00000138735 | 446.6156 | 1.043784 | 0.21172 | 4.930028 | 8.22E-07 | 2.78E-05 | 10.83992 | 26.35534 | 13.19644 | 21.02659 | PDE5A | | 4 |
| 311 | ENSG00000138759 | 1569.966 | -1.02453 | 0.139541 | -7.34211 | 2.10E-13 | 2.00E-11 | 21.38024 | 10.58227 | 21.41696 | 9.474183 | FRAS1 | | 4 |

| | | | | | | | | | | | | | |
|---|---|---|---|---|---|---|---|---|---|---|---|---|---|
| 312 | ENSG00000138769 | 76.70414 | -1.57079 | 0.427724 | -3.67243 | 0.00024 | 0.004633 | 2.953877 | 0.832544 | 3.268181 | 1.153403 | CDKL2 | 4 |
| 313 | ENSG00000138772 | 36.02813 | 2.789642 | 0.685463 | 4.069717 | 4.71E-05 | 0.001106 | 0.88665 | 8.702374 | 1.075551 | 4.251571 | ANXA3 | 4 |
| 314 | ENSG00000138823 | 512.7777 | 1.522196 | 0.193569 | 7.86385 | 3.73E-15 | 4.45E-13 | 8.000757 | 24.52756 | 9.509662 | 23.36251 | MTTP | 4 |
| 315 | ENSG00000138944 | 173.1922 | -1.01405 | 0.291929 | -3.4736 | 0.000514 | 0.00884 | 3.406971 | 1.929128 | 4.214913 | 1.671298 | SHISAL1 | 22 |
| 316 | ENSG00000139178 | 65.35785 | 1.565869 | 0.455989 | 3.434007 | 0.000595 | 0.009971 | 1.339777 | 4.63645 | 1.866919 | 4.382172 | C1RL | 12 |
| 317 | ENSG00000139219 | 923.0595 | 1.21596 | 0.197914 | 6.143889 | 8.05E-10 | 4.74E-08 | 12.65164 | 32.63613 | 19.05526 | 37.20634 | COL2A1 | 12 |
| 318 | ENSG00000139289 | 538.1884 | 1.372923 | 0.238413 | 5.758586 | 8.48E-09 | 4.15E-07 | 4.713309 | 11.97323 | 7.006651 | 16.75021 | PHLDA1 | 12 |
| 319 | ENSG00000139318 | 191.374 | -1.15617 | 0.339041 | -3.41013 | 0.000649 | 0.010711 | 8.436739 | 4.929594 | 16.82015 | 5.761455 | DUSP6 | 12 |
| 320 | ENSG00000139514 | 1225.852 | 1.054666 | 0.1728 | 6.103385 | 1.04E-09 | 5.97E-08 | 13.39164 | 22.34409 | 12.38379 | 28.49016 | SLC7A1 | 13 |
| 321 | ENSG00000139722 | 518.7743 | -1.53569 | 0.218092 | -7.04149 | 1.90E-12 | 1.60E-10 | 38.43349 | 13.66612 | 31.74329 | 9.560889 | VPS37B | 12 |
| 322 | ENSG00000139800 | 573.6035 | 1.209469 | 0.184503 | 6.555268 | 5.55E-11 | 3.83E-09 | 8.125713 | 18.39977 | 9.018272 | 19.30464 | ZIC5 | 13 |
| 323 | ENSG00000140015 | 233.5052 | -2.54368 | 0.309205 | -8.22649 | 1.93E-16 | 2.71E-14 | 7.713096 | 0.975791 | 9.426109 | 1.776296 | KCNH5 | 14 |
| 324 | ENSG00000140285 | 40.53446 | -3.93938 | 0.657811 | -5.98862 | 2.12E-09 | 1.14E-07 | 2.221508 | 0.203622 | 2.548523 | 0.100507 | FGF7 | 15 |
| 325 | ENSG00000140416 | 1290.955 | 1.51658 | 0.176014 | 8.616269 | 6.92E-18 | 1.08E-15 | 43.30951 | 152.5389 | 53.44224 | 111.7939 | TPM1 | 15 |
| 326 | ENSG00000140488 | 39.35523 | -2.28723 | 0.684598 | -3.34099 | 0.000835 | 0.013191 | 2.497947 | 0.811133 | 2.293826 | 0.163491 | CELF6 | 15 |
| 327 | ENSG00000140807 | 1127.081 | 1.726619 | 0.177574 | 9.7234 | 2.40E-22 | 4.80E-20 | 2.971855 | 12.13842 | 3.877146 | 9.484645 | NKD1 | 16 |
| 328 | ENSG00000140873 | 156.5295 | 2.874629 | 0.363033 | 7.918372 | 2.41E-15 | 2.91E-13 | 0.864043 | 9.829256 | 1.500777 | 6.671955 | ADAMTS1 | 16 |
| 329 | ENSG00000140937 | 605.7914 | -1.87069 | 0.180337 | -10.3733 | 3.28E-25 | 8.09E-23 | 41.59074 | 11.32904 | 41.49049 | 10.33238 | CDH11 | 16 |
| 330 | ENSG00000141013 | 31.07629 | -2.48062 | 0.684217 | -3.62549 | 0.000288 | 0.005384 | 5.11612 | 0.827187 | 4.221631 | 0.770444 | GAS8 | 16 |
| 331 | ENSG00000141314 | 898.3585 | 1.125574 | 0.164634 | 6.836834 | 8.10E-12 | 6.23E-10 | 16.66449 | 38.06596 | 16.83442 | 31.69463 | RHBDL3 | 17 |
| 332 | ENSG00000141338 | 109.6188 | 2.189122 | 0.381097 | 5.744266 | 9.23E-09 | 4.48E-07 | 1.683979 | 8.808314 | 1.83764 | 6.523135 | ABCA8 | 17 |
| 333 | ENSG00000142178 | 167.8401 | 1.043662 | 0.317786 | 3.28417 | 0.001023 | 0.015774 | 5.971717 | 16.83403 | 8.976214 | 12.60234 | SIK1 | 21 |
| 334 | ENSG00000142609 | 20.79675 | -2.8333 | 0.841616 | -3.36649 | 0.000761 | 0.012251 | 2.00233 | 0.169952 | 1.936588 | 0.330016 | CFAP74 | 1 |
| 335 | ENSG00000142627 | 771.6961 | 1.235328 | 0.185908 | 6.644833 | 3.04E-11 | 2.15E-09 | 13.76216 | 30.32398 | 16.32019 | 36.93252 | EPHA2 | 1 |
| 336 | ENSG00000142700 | 181.9842 | -3.76085 | 0.351799 | -10.6903 | 1.13E-26 | 2.98E-24 | 11.94099 | 1.23875 | 14.465 | 0.658056 | DMRTA2 | 1 |
| 337 | ENSG00000143162 | 257.8069 | 1.168633 | 0.298553 | 3.914325 | 9.07E-05 | 0.001976 | 6.776621 | 21.44282 | 13.34374 | 21.51675 | CREG1 | 1 |
| 338 | ENSG00000143341 | 154.9428 | 1.356316 | 0.317864 | 4.266973 | 1.98E-05 | 0.000506 | 0.786267 | 1.601082 | 0.587573 | 1.694435 | HMCN1 | 1 |
| 339 | ENSG00000143355 | 401.6125 | -2.86542 | 0.61485 | -4.66036 | 3.16E-06 | 9.46E-05 | 27.25236 | 3.965212 | 16.54281 | 1.83944 | LHX9 | 1 |
| 340 | ENSG00000143473 | 127.0021 | 2.59128 | 0.362365 | 7.151013 | 8.61E-13 | 7.60E-11 | 0.689582 | 5.022553 | 1.016104 | 4.678853 | KCNH1 | 1 |
| 341 | ENSG00000143479 | 481.0573 | -1.08014 | 0.218695 | -4.939 | 7.85E-07 | 2.67E-05 | 13.59011 | 7.467541 | 19.7269 | 7.48415 | DYRK3 | 1 |
| 342 | ENSG00000143842 | 307.8616 | 1.520047 | 0.250611 | 6.065364 | 1.32E-09 | 7.32E-08 | 5.179093 | 16.37078 | 7.512468 | 18.17814 | SOX13 | 1 |
| 343 | ENSG00000143869 | 149.3904 | 1.120067 | 0.306041 | 3.659857 | 0.000252 | 0.004838 | 1.085912 | 2.456784 | 1.180549 | 2.239298 | GDF7 | 2 |
| 344 | ENSG00000144290 | 41.21978 | -2.04811 | 0.583808 | -3.50819 | 0.000451 | 0.007915 | 1.762452 | 0.370109 | 2.138362 | 0.510591 | SLC4A10 | 2 |
| 345 | ENSG00000144331 | 103.9426 | -1.25025 | 0.397135 | -3.14817 | 0.001643 | 0.023336 | 5.488844 | 3.48373 | 9.795516 | 2.636532 | ZNF385B | 2 |
| 346 | ENSG00000144355 | 103.4605 | -6.60874 | 0.806178 | -8.19762 | 2.45E-16 | 3.37E-14 | 16.62515 | 0.215874 | 12.60117 | 0.103768 | DLX1 | 2 |
| 347 | ENSG00000144369 | 910.762 | -1.76044 | 0.16563 | -10.6288 | 2.19E-26 | 5.70E-24 | 25.96506 | 7.578049 | 29.79784 | 8.057313 | FAM171B | 2 |
| 348 | ENSG00000144476 | 307.5108 | -1.05036 | 0.24951 | -4.20969 | 2.56E-05 | 0.000636 | 29.61097 | 17.77114 | 34.72829 | 12.01307 | ACKR3 | 2 |
| 349 | ENSG00000144583 | 36.3077 | -2.14464 | 0.630205 | -3.40309 | 0.000666 | 0.010951 | 1.177276 | 0.221893 | 1.596498 | 0.365214 | MARCHF4 | 2 |
| 350 | ENSG00000144596 | 137.9955 | -1.31997 | 0.383318 | -3.44352 | 0.000574 | 0.009745 | 2.474686 | 1.688028 | 5.146212 | 1.226156 | GRIP2 | 3 |
| 351 | ENSG00000144847 | 185.8171 | -2.98253 | 0.301979 | -9.87664 | 5.26E-23 | 1.09E-20 | 14.22447 | 1.797631 | 14.87684 | 1.709418 | IGSF11 | 3 |
| 352 | ENSG00000144857 | 2187.159 | 1.229868 | 0.14434 | 8.520627 | 1.59E-17 | 2.40E-15 | 46.5626 | 100.8812 | 50.92501 | 116.386 | BOC | 3 |
| 353 | ENSG00000145358 | 91.16696 | 1.388331 | 0.448417 | 3.096075 | 0.001961 | 0.026843 | 1.385915 | 6.766518 | 3.437523 | 5.265203 | DDIT4L | 4 |
| 354 | ENSG00000145365 | 88.11496 | 1.162676 | 0.406001 | 2.863727 | 0.004187 | 0.049711 | 1.911835 | 4.147519 | 2.654064 | 5.548415 | TIFA | 4 |
| 355 | ENSG00000145428 | 305.7527 | 1.116893 | 0.226937 | 4.921603 | 8.58E-07 | 2.89E-05 | 14.17269 | 31.17749 | 15.41416 | 29.93365 | RNF175 | 4 |
| 356 | ENSG00000145431 | 266.3835 | 1.317665 | 0.29848 | 4.41459 | 1.01E-05 | 0.000271 | 4.917679 | 12.4699 | 8.133608 | 18.22297 | PDGFC | 4 |
| 357 | ENSG00000145506 | 85.24192 | 2.005853 | 0.512041 | 3.917365 | 8.95E-05 | 0.001959 | 1.122686 | 8.569913 | 4.181932 | 10.94864 | NKD2 | 5 |
| 358 | ENSG00000145526 | 244.4764 | -1.08616 | 0.316542 | -3.43132 | 0.000601 | 0.010061 | 14.14955 | 6.927368 | 25.05703 | 10.47358 | CDH18 | 5 |
| 359 | ENSG00000145555 | 2605.501 | -2.04567 | 0.137859 | -14.8388 | 8.22E-50 | 7.19E-47 | 73.38163 | 19.60906 | 82.10247 | 16.34166 | MYO10 | 5 |
| 360 | ENSG00000145632 | 990.8304 | 1.167695 | 0.183055 | 6.378931 | 1.78E-10 | 1.16E-08 | 36.03697 | 71.12964 | 40.86847 | 92.80558 | PLK2 | 5 |
| 361 | ENSG00000145721 | 672.9993 | -2.20141 | 0.216935 | -10.1478 | 3.39E-24 | 7.68E-22 | 39.98462 | 8.330874 | 29.12857 | 6.097672 | LIX1 | 5 |
| 362 | ENSG00000145920 | 512.857 | -1.74661 | 0.200179 | -8.72526 | 2.66E-18 | 4.22E-16 | 23.23716 | 6.013114 | 19.71413 | 6.175428 | CPLX2 | 5 |
| 363 | ENSG00000145934 | 2792.393 | 1.911152 | 0.131638 | 14.51819 | 9.29E-48 | 6.88E-45 | 18.26276 | 73.35228 | 20.81862 | 66.73034 | TENM2 | 5 |
| 364 | ENSG00000146001 | 87.79306 | -1.45505 | 0.395649 | -3.67763 | 0.000235 | 0.00455 | 5.397119 | 1.892148 | 4.572928 | 1.588608 | PCDHB18F | 5 |
| 365 | ENSG00000146005 | 160.5344 | -1.67364 | 0.31659 | -5.28647 | 1.25E-07 | 4.94E-06 | 7.943168 | 2.996161 | 8.363867 | 1.947863 | PSD2 | 5 |
| 366 | ENSG00000146242 | 1159.425 | -1.13934 | 0.159464 | -7.14483 | 9.01E-13 | 7.92E-11 | 57.47211 | 28.75346 | 71.02456 | 26.78194 | TPBG | 6 |
| 367 | ENSG00000146374 | 200.8917 | -2.48494 | 0.356764 | -6.96521 | 3.28E-12 | 2.67E-10 | 8.96159 | 2.550775 | 11.26199 | 1.019025 | RSPO3 | 6 |
| 368 | ENSG00000146648 | 652.5197 | 2.693873 | 0.186235 | 14.46493 | 2.02E-47 | 1.44E-44 | 2.181581 | 14.27248 | 2.156044 | 12.49725 | EGFR | 7 |
| 369 | ENSG00000146910 | 64.90365 | 2.288395 | 0.534983 | 4.27751 | 1.89E-05 | 0.000487 | 0.894454 | 6.237289 | 2.37421 | 9.562309 | CNPY1 | 7 |
| 370 | ENSG00000146950 | 842.0726 | -1.08069 | 0.167979 | -6.43345 | 1.25E-10 | 8.31E-09 | 19.96882 | 10.4351 | 22.96072 | 8.933406 | SHROOM2 | X |
| 371 | ENSG00000147100 | 1203.181 | -1.39444 | 0.151322 | -9.21503 | 3.11E-20 | 5.65E-18 | 47.21409 | 19.16644 | 53.37687 | 17.30444 | SLC16A2 | X |
| 372 | ENSG00000147119 | 53.93456 | -2.0117 | 0.513554 | -3.9172 | 8.96E-05 | 0.001959 | 4.723342 | 0.854188 | 4.028793 | 1.192133 | CHST7 | X |
| 373 | ENSG00000147256 | 29.72013 | -2.95016 | 0.721838 | -4.08701 | 4.37E-05 | 0.001036 | 2.439339 | 0.260752 | 2.575352 | 0.349283 | ARHGAP3 | X |
| 374 | ENSG00000147257 | 1810.864 | -1.04456 | 0.154653 | -6.75418 | 1.44E-11 | 1.07E-09 | 152.0216 | 62.08628 | 149.0677 | 76.46982 | GPC3 | X |
| 375 | ENSG00000147459 | 393.1218 | 2.242716 | 0.253713 | 8.839587 | 9.61E-19 | 1.59E-16 | 2.752673 | 10.18601 | 2.956507 | 15.45645 | DOCK5 | 8 |
| 376 | ENSG00000147488 | 691.3068 | -1.26891 | 0.243968 | -5.20112 | 1.98E-07 | 7.48E-06 | 54.9244 | 21.03522 | 33.61231 | 14.25411 | ST18 | 8 |
| 377 | ENSG00000147571 | 17.07525 | -3.37343 | 1.030087 | -3.2749 | 0.001057 | 0.016145 | 3.813618 | 0.550435 | 2.668221 | 0.082754 | CRH | 8 |
| 378 | ENSG00000148123 | 96.66936 | -1.62186 | 0.382828 | -4.23652 | 2.27E-05 | 0.000574 | 8.31179 | 2.204287 | 8.127986 | 2.832112 | PLPPR1 | 9 |
| 379 | ENSG00000148541 | 137.1838 | -1.20301 | 0.317357 | -3.7907 | 0.00015 | 0.003095 | 15.88553 | 6.70885 | 15.29159 | 6.205817 | FAM13C | 10 |
| 380 | ENSG00000148926 | 54.15485 | -1.94468 | 0.579393 | -3.35642 | 0.00079 | 0.012614 | 5.562086 | 0.870266 | 8.738647 | 2.606823 | ADM | 11 |
| 381 | ENSG00000149257 | 543.4268 | 1.13341 | 0.204842 | 5.533084 | 3.15E-08 | 1.42E-06 | 17.41197 | 38.91391 | 22.8363 | 44.89732 | SERPINH1 | 11 |
| 382 | ENSG00000149294 | 3695.196 | -1.13666 | 0.134514 | -8.45014 | 2.91E-17 | 4.34E-15 | 181.6876 | 86.85496 | 178.6112 | 69.63078 | NCAM1 | 11 |
| 383 | ENSG00000149571 | 150.0864 | 1.629149 | 0.343975 | 4.736247 | 2.18E-06 | 6.78E-05 | 2.502633 | 10.8391 | 4.607158 | 9.994173 | KIRREL3 | 11 |
| 384 | ENSG00000150275 | 282.0691 | -3.40443 | 0.86713 | -3.92609 | 8.63E-05 | 0.001901 | 8.525221 | 0.712299 | 20.34103 | 1.851264 | PCDH15 | 10 |
| 385 | ENSG00000150672 | 269.6541 | -1.89282 | 0.251028 | -7.54027 | 4.69E-14 | 4.73E-12 | 13.68883 | 4.013779 | 16.97138 | 3.845971 | DLG2 | 11 |
| 386 | ENSG00000150687 | 2081.775 | -1.08613 | 0.144096 | -7.53753 | 4.79E-14 | 4.80E-12 | 86.98137 | 42.80617 | 106.5003 | 43.83331 | PRSS23 | 11 |
| 387 | ENSG00000151023 | 190.3746 | -1.20197 | 0.290772 | -4.13373 | 3.57E-05 | 0.000863 | 19.70338 | 7.194603 | 14.83024 | 7.115508 | ENKUR | 10 |
| 388 | ENSG00000151090 | 70.12362 | 1.667362 | 0.486684 | 3.425966 | 0.000613 | 0.010191 | 0.620126 | 2.350181 | 1.294482 | 3.152883 | THRB | 3 |
| 389 | ENSG00000151468 | 52.66347 | 2.196019 | 0.551573 | 3.981377 | 6.85E-05 | 0.001553 | 1.05339 | 2.978021 | 0.491862 | 3.700937 | CCDC3 | 10 |

| | | | | | | | | | | | | | |
|---|---|---|---|---|---|---|---|---|---|---|---|---|---|
| 390 | ENSG00000151490 | 416.1496 | -1.51109 | 0.226385 | -6.67487 | 2.47E-11 | 1.78E-09 | 20.00601 | 8.488387 | 22.24563 | 5.75335 | PTPRO | 12 |
| 391 | ENSG00000151726 | 194.9387 | 1.348735 | 0.27786 | 4.854005 | 1.21E-06 | 3.96E-05 | 3.566009 | 8.92388 | 4.041124 | 9.500898 | ACSL1 | 4 |
| 392 | ENSG00000151834 | 57.9136 | -2.84972 | 0.558301 | -5.10426 | 3.32E-07 | 1.20E-05 | 3.494765 | 0.408623 | 4.532252 | 0.687416 | GABRA2 | 4 |
| 393 | ENSG00000152092 | 1070.425 | -1.69593 | 0.164387 | -10.3167 | 5.92E-25 | 1.41E-22 | 29.98243 | 10.35316 | 32.60401 | 8.11159 | ASTN1 | 1 |
| 394 | ENSG00000152192 | 165.1015 | 1.83507 | 0.315406 | 5.818124 | 5.95E-09 | 2.98E-07 | 1.793454 | 7.328037 | 1.861207 | 5.152084 | POU4F1 | 13 |
| 395 | ENSG00000152689 | 25.01253 | -2.74643 | 0.818274 | -3.35637 | 0.00079 | 0.012614 | 1.427641 | 0.529871 | 3.176852 | 0.177216 | RASGRP3 | 2 |
| 396 | ENSG00000152760 | 19.93457 | 2.642357 | 0.841806 | 3.138915 | 0.001696 | 0.02398 | 0.579046 | 2.082848 | 0.440715 | 3.941397 | DYNLT5 | 1 |
| 397 | ENSG00000152778 | 474.2671 | -1.05783 | 0.224453 | -4.7129 | 2.44E-06 | 7.53E-05 | 26.43295 | 15.01423 | 26.88573 | 9.605729 | IFIT5 | 10 |
| 398 | ENSG00000152910 | 55.85935 | -1.88555 | 0.504166 | -3.73994 | 0.000184 | 0.003706 | 2.834849 | 0.828527 | 2.391814 | 0.518516 | CNTNAP4 | 16 |
| 399 | ENSG00000153012 | 390.8324 | 1.989238 | 0.233199 | 8.530222 | 1.46E-17 | 2.25E-15 | 3.173367 | 10.27163 | 3.107118 | 13.34664 | LGI2 | 4 |
| 400 | ENSG00000153071 | 117.4943 | 1.131503 | 0.369821 | 3.059599 | 0.002216 | 0.029808 | 3.883294 | 6.263763 | 2.368072 | 6.49961 | DAB2 | 5 |
| 401 | ENSG00000153266 | 119.9185 | -3.12474 | 0.402268 | -7.76781 | 7.99E-15 | 8.94E-13 | 13.50002 | 1.978877 | 12.39204 | 0.926673 | FEZF2 | 3 |
| 402 | ENSG00000153993 | 121.0061 | -1.75558 | 0.360199 | -4.87393 | 1.09E-06 | 3.62E-05 | 4.193778 | 1.010814 | 2.922191 | 0.99647 | SEMA3D | 7 |
| 403 | ENSG00000154188 | 157.3713 | 2.404385 | 0.41728 | 5.762041 | 8.31E-09 | 4.09E-07 | 0.998825 | 8.381301 | 3.101664 | 12.03742 | ANGPT1 | 8 |
| 404 | ENSG00000154229 | 330.5494 | 1.578884 | 0.258977 | 6.09661 | 1.08E-09 | 6.19E-08 | 2.303488 | 9.31814 | 3.168781 | 6.294235 | PRKCA | 17 |
| 405 | ENSG00000154265 | 268.3274 | -1.54703 | 0.284038 | -5.44654 | 5.14E-08 | 2.22E-06 | 12.19822 | 3.344736 | 15.03134 | 5.4651 | ABCA5 | 17 |
| 406 | ENSG00000154654 | 105.9163 | -1.43443 | 0.393448 | -3.64579 | 0.000267 | 0.00506 | 4.019502 | 1.495293 | 6.317824 | 2.137501 | NCAM2 | 21 |
| 407 | ENSG00000154856 | 318.9738 | 2.176589 | 0.314296 | 6.925291 | 4.35E-12 | 3.39E-10 | 3.42336 | 28.19369 | 7.809203 | 19.94437 | APCDD1 | 18 |
| 408 | ENSG00000154874 | 53.84023 | -2.11182 | 0.508794 | -4.15064 | 3.32E-05 | 0.000807 | 1.218716 | 0.272048 | 1.600795 | 0.341888 | CCDC144B | 17 |
| 409 | ENSG00000155875 | 69.21722 | -2.87481 | 0.50558 | -5.68617 | 1.30E-08 | 6.14E-07 | 1.777726 | 0.28718 | 1.36079 | 0.131424 | SAXO1 | 9 |
| 410 | ENSG00000156804 | 256.3444 | -3.07786 | 0.899554 | -3.42154 | 0.000623 | 0.01034 | 15.11683 | 1.042589 | 4.053869 | 1.133604 | FBXO32 | 8 |
| 411 | ENSG00000156869 | 42.50935 | 2.162606 | 0.579743 | 3.730286 | 0.000191 | 0.003824 | 0.329514 | 1.062543 | 0.272814 | 1.495919 | FRRS1 | 1 |
| 412 | ENSG00000157168 | 311.6062 | 1.499003 | 0.26785 | 5.596437 | 2.19E-08 | 1.01E-06 | 3.826919 | 15.07892 | 6.561786 | 12.79375 | NRG1 | 8 |
| 413 | ENSG00000157423 | 505.0767 | -1.30948 | 0.196958 | -6.64852 | 2.96E-11 | 2.10E-09 | 17.21141 | 7.770474 | 20.33879 | 6.696014 | HYDIN | 16 |
| 414 | ENSG00000157542 | 107.2015 | 1.63435 | 0.465866 | 3.508195 | 0.000451 | 0.007915 | 0.665609 | 1.929794 | 0.297392 | 0.891649 | KCNJ6 | 21 |
| 415 | ENSG00000157856 | 22.05871 | -2.59235 | 0.805949 | -3.21651 | 0.001298 | 0.019123 | 3.447735 | 0.568274 | 3.024406 | 0.473826 | DRC1 | 2 |
| 416 | ENSG00000157978 | 69.34099 | 1.42841 | 0.490733 | 2.910766 | 0.003605 | 0.043974 | 0.82656 | 2.696376 | 1.777613 | 3.89349 | LDLRAP1 | 1 |
| 417 | ENSG00000158246 | 65.79897 | 1.751661 | 0.470669 | 3.721645 | 0.000198 | 0.003943 | 1.256894 | 4.298917 | 1.819657 | 5.520068 | TENT5B | 1 |
| 418 | ENSG00000158270 | 29.48704 | 4.033767 | 0.876681 | 4.601178 | 4.20E-06 | 0.000122 | 0 | 1.370364 | 0.141787 | 1.185776 | COLEC12 | 18 |
| 419 | ENSG00000158856 | 66.77433 | -1.50623 | 0.458803 | -3.28296 | 0.001027 | 0.015829 | 4.959534 | 2.197081 | 7.159456 | 1.870302 | DMTN | 8 |
| 420 | ENSG00000158955 | 66.92699 | 5.321267 | 0.696972 | 7.634831 | 2.26E-14 | 2.38E-12 | 0.097833 | 3.101465 | 0.093921 | 4.134765 | WNT9B | 17 |
| 421 | ENSG00000159176 | 81.19018 | -3.98266 | 0.521936 | -7.63055 | 2.34E-14 | 2.45E-12 | 8.718547 | 0.927564 | 12.12622 | 0.485601 | CSRP1 | 1 |
| 422 | ENSG00000159248 | 83.79636 | -3.40804 | 0.484536 | -7.03362 | 2.01E-12 | 1.68E-10 | 7.588026 | 0.686366 | 4.812538 | 0.446217 | GJD2 | 15 |
| 423 | ENSG00000159307 | 1306.079 | -1.10046 | 0.145882 | -7.54348 | 4.58E-14 | 4.64E-12 | 32.57832 | 15.65138 | 36.11013 | 14.85881 | SCUBE1 | 22 |
| 424 | ENSG00000159387 | 45.46301 | 3.532079 | 0.661293 | 5.341172 | 9.23E-08 | 3.76E-06 | 0.14379 | 3.111333 | 0.501583 | 3.995842 | IRX6 | 16 |
| 425 | ENSG00000159409 | 837.9335 | -1.22275 | 0.169444 | -7.21629 | 5.34E-13 | 4.85E-11 | 46.41865 | 18.07632 | 40.26647 | 17.32672 | CELF3 | 1 |
| 426 | ENSG00000159713 | 108.6482 | -1.28763 | 0.363157 | -3.54566 | 0.000392 | 0.007018 | 19.43709 | 7.133876 | 22.6844 | 9.184763 | TPPP3 | 16 |
| 427 | ENSG00000159784 | 713.6759 | -1.16528 | 0.171662 | -6.78822 | 1.14E-11 | 8.60E-10 | 28.8647 | 13.68417 | 31.40913 | 11.95273 | FAM131B | 7 |
| 428 | ENSG00000160190 | 166.1976 | -1.58482 | 0.300379 | -5.27607 | 1.32E-07 | 5.21E-06 | 10.97298 | 3.481305 | 12.77558 | 4.037609 | SLC37A1 | 21 |
| 429 | ENSG00000161082 | 422.2641 | -1.08905 | 0.210768 | -5.16705 | 2.38E-07 | 8.87E-06 | 20.15883 | 8.844624 | 22.96844 | 10.37482 | CELF5 | 19 |
| 430 | ENSG00000161298 | 312.7719 | -1.7657 | 0.248034 | -7.1188 | 1.09E-12 | 9.48E-11 | 17.568 | 5.473457 | 24.04978 | 6.113411 | ZNF382 | 19 |
| 431 | ENSG00000161638 | 424.7359 | 1.01969 | 0.222528 | 4.582292 | 4.60E-06 | 0.000132 | 7.674615 | 17.62749 | 10.98901 | 18.36004 | ITGA5 | 12 |
| 432 | ENSG00000161996 | 141.5041 | -1.02395 | 0.350365 | -2.92253 | 0.003472 | 0.042753 | 11.89726 | 4.603155 | 14.41244 | 7.667753 | WDR90 | 16 |
| 433 | ENSG00000162374 | 737.8374 | -1.5855 | 0.181519 | -8.7346 | 2.45E-18 | 3.92E-16 | 54.13832 | 18.27145 | 48.76124 | 14.52979 | ELAVL4 | 1 |
| 434 | ENSG00000162407 | 685.2432 | -1.00713 | 0.177938 | -5.66003 | 1.51E-08 | 7.09E-07 | 36.0891 | 16.928 | 31.72345 | 15.27119 | PLPP3 | 1 |
| 435 | ENSG00000162493 | 496.5088 | 2.600219 | 0.204363 | 12.72354 | 4.38E-37 | 1.79E-34 | 9.434678 | 52.68234 | 9.682925 | 57.53256 | PDPN | 1 |
| 436 | ENSG00000162551 | 318.0989 | 1.165763 | 0.251801 | 4.6297 | 3.66E-06 | 0.000108 | 8.092714 | 19.47213 | 11.83142 | 23.01881 | ALPL | 1 |
| 437 | ENSG00000162552 | 138.0034 | 2.308142 | 0.441233 | 5.231116 | 1.68E-07 | 6.48E-06 | 0.78363 | 7.772576 | 2.836781 | 8.692905 | WNT4 | 1 |
| 438 | ENSG00000162692 | 399.8578 | -5.30251 | 0.694211 | -7.63819 | 2.20E-14 | 2.33E-12 | 42.67936 | 0.823375 | 17.84304 | 0.649286 | VCAM1 | 1 |
| 439 | ENSG00000162753 | 12.28036 | -5.37946 | 1.277685 | -4.21031 | 2.55E-05 | 0.000635 | 5.048247 | 0.26793 | 4.958805 | 0.027604 | SLC9C2 | 1 |
| 440 | ENSG00000162772 | 438.3731 | 1.341272 | 0.20748 | 6.464601 | 1.02E-10 | 6.79E-09 | 16.56588 | 39.64205 | 18.59609 | 44.96663 | ATF3 | 1 |
| 441 | ENSG00000162804 | 149.0241 | 1.97813 | 0.363098 | 5.447915 | 5.10E-08 | 2.20E-06 | 0.844471 | 2.986543 | 1.274217 | 4.889612 | SNED1 | 2 |
| 442 | ENSG00000162849 | 321.1544 | -1.12107 | 0.235261 | -4.76523 | 1.89E-06 | 5.92E-05 | 4.132443 | 1.548719 | 3.550929 | 1.81221 | KIF26B | 1 |
| 443 | ENSG00000162873 | 1438.486 | -1.92005 | 0.180893 | -10.6143 | 2.56E-26 | 6.56E-24 | 83.15783 | 28.33974 | 113.7931 | 21.3476 | KLHDC8A | 1 |
| 444 | ENSG00000162981 | 114.9198 | -1.05606 | 0.362529 | -2.91303 | 0.003579 | 0.04374 | 3.918045 | 2.461864 | 4.885278 | 1.603867 | LRATD1 | 2 |
| 445 | ENSG00000162992 | 57.94487 | -1.92713 | 0.528287 | -3.64789 | 0.000264 | 0.005028 | 5.869774 | 1.557017 | 3.570578 | 0.874187 | NEUROD1 | 2 |
| 446 | ENSG00000162998 | 133.498 | -1.1248 | 0.38742 | -2.9033 | 0.003693 | 0.044668 | 5.467066 | 4.320762 | 11.23815 | 2.998908 | FRZB | 2 |
| 447 | ENSG00000163017 | 24.22503 | 2.474795 | 0.787741 | 3.141637 | 0.00168 | 0.023793 | 1.279671 | 5.10598 | 0.858416 | 5.754374 | ACTG2 | 2 |
| 448 | ENSG00000163071 | 71.84937 | -1.39482 | 0.47313 | -2.94808 | 0.003198 | 0.040196 | 3.647184 | 1.067624 | 5.032251 | 2.070139 | SPATA18 | 4 |
| 449 | ENSG00000163081 | 22.6118 | 3.744405 | 0.930394 | 4.024536 | 5.71E-05 | 0.001322 | 0.080381 | 2.630581 | 0.458869 | 4.243284 | CCDC140 | 2 |
| 450 | ENSG00000163191 | 23.3346 | 2.844649 | 0.825533 | 3.445833 | 0.000569 | 0.009688 | 1.131753 | 12.97354 | 2.876564 | 14.4376 | S100A11 | 1 |
| 451 | ENSG00000163273 | 108.6142 | 4.227581 | 0.470356 | 8.988054 | 2.52E-19 | 4.29E-17 | 1.798537 | 34.27266 | 1.650954 | 26.90946 | NPPC | 2 |
| 452 | ENSG00000163285 | 30.24427 | -4.3578 | 0.85099 | -5.12085 | 3.04E-07 | 1.10E-05 | 1.020889 | 0.035266 | 0.902991 | 0.052177 | GABRG1 | 4 |
| 453 | ENSG00000163347 | 84.9801 | 3.024085 | 0.484172 | 6.245891 | 4.21E-10 | 2.55E-08 | 0.328163 | 4.484509 | 0.936566 | 5.298025 | CLDN1 | 3 |
| 454 | ENSG00000163359 | 27.9688 | 6.414453 | 1.298715 | 4.939075 | 7.85E-07 | 2.67E-05 | 0.027827 | 2.553648 | 0 | 0.75741 | COL6A3 | 2 |
| 455 | ENSG00000163377 | 124.8939 | 2.066985 | 0.423133 | 4.884952 | 1.03E-06 | 3.45E-05 | 1.471736 | 9.872662 | 4.144553 | 12.39983 | TAFA4 | 3 |
| 456 | ENSG00000163508 | 229.5328 | -3.78042 | 0.880567 | -4.29316 | 1.76E-05 | 0.000458 | 18.14261 | 1.984627 | 16.69047 | 0.480057 | EOMES | 3 |
| 457 | ENSG00000163618 | 1266.112 | -1.48939 | 0.157743 | -9.44185 | 3.66E-21 | 6.91E-19 | 98.5783 | 39.75717 | 114.1987 | 32.67115 | CADPS | 3 |
| 458 | ENSG00000163629 | 5268.972 | 1.299338 | 0.128911 | 10.07937 | 6.82E-24 | 1.51E-21 | 42.97837 | 103.2581 | 49.26809 | 112.5667 | PTPN13 | 4 |
| 459 | ENSG00000163686 | 141.6639 | -1.23368 | 0.366747 | -3.36384 | 0.000769 | 0.012328 | 8.511559 | 5.455029 | 16.73629 | 4.744716 | ABHD6 | 3 |
| 460 | ENSG00000163873 | 136.5419 | -1.53155 | 0.32613 | -4.69612 | 2.65E-06 | 8.14E-05 | 2.870944 | 1.091013 | 3.379697 | 0.973963 | GRIK3 | 1 |
| 461 | ENSG00000164089 | 37.69613 | -3.71526 | 0.711483 | -5.22185 | 1.77E-07 | 6.75E-06 | 9.21808 | 0.511414 | 4.761159 | 0.490079 | ETNPPL | 4 |
| 462 | ENSG00000164171 | 250.1645 | -2.52367 | 0.282888 | -8.92108 | 4.62E-19 | 7.73E-17 | 7.681629 | 1.284783 | 5.693984 | 0.950021 | ITGA2 | 5 |
| 463 | ENSG00000164330 | 867.7336 | 1.049738 | 0.169548 | 6.191394 | 5.96E-10 | 3.55E-08 | 14.46605 | 33.36166 | 17.89291 | 30.38836 | EBF1 | 5 |
| 464 | ENSG00000164418 | 300.9044 | -1.06829 | 0.229788 | -4.64902 | 3.34E-06 | 9.97E-05 | 14.23544 | 6.54643 | 13.36593 | 6.012873 | GRIK2 | 6 |
| 465 | ENSG00000164434 | 1140.499 | -3.10309 | 0.160948 | -19.28 | 7.91E-83 | 1.69E-79 | 412.9019 | 44.23542 | 388.05 | 44.48974 | FABP7 | 6 |
| 466 | ENSG00000164438 | 25.10669 | 2.791883 | 0.800294 | 3.488572 | 0.000486 | 0.008443 | 0.725594 | 4.364991 | 0.345191 | 2.735468 | TLX3 | 5 |
| 467 | ENSG00000164483 | 81.22194 | -2.82013 | 0.541384 | -5.20911 | 1.90E-07 | 7.20E-06 | 12.82312 | 3.110822 | 11.88864 | 0.743393 | SAMD3 | 6 |

| | | | | | | | | | | | | | | |
|---|---|---|---|---|---|---|---|---|---|---|---|---|---|---|
| 468 | ENSG00000164651 | 115.3887 | -3.53814 | 0.47113 | -7.50991 | 5.92E-14 | 5.87E-12 | 5.988339 | 0.989082 | 8.693831 | 0.262113 | SP8 | | 7 |
| 469 | ENSG00000164690 | 45.9755 | -3.03211 | 0.607565 | -4.99059 | 6.02E-07 | 2.09E-05 | 2.516384 | 0.260708 | 1.511888 | 0.212456 | SHH | | 7 |
| 470 | ENSG00000164692 | 2501.557 | 5.724303 | 0.473266 | 12.09532 | 1.12E-33 | 4.06E-31 | 2.255274 | 132.1477 | 4.078609 | 186.369 | COL1A2 | | 7 |
| 471 | ENSG00000164778 | 240.639 | 2.583741 | 0.336799 | 7.671467 | 1.70E-14 | 1.83E-12 | 1.556249 | 10.75478 | 3.207743 | 16.26123 | EN2 | | 7 |
| 472 | ENSG00000164853 | 93.44091 | -2.28228 | 0.422593 | -5.40065 | 6.64E-08 | 2.80E-06 | 10.82069 | 2.177218 | 7.405008 | 1.437569 | UNCX | | 7 |
| 473 | ENSG00000164946 | 59.17141 | 1.510001 | 0.512912 | 2.943977 | 0.00324 | 0.040574 | 0.59522 | 2.628887 | 1.327577 | 2.475781 | FREM1 | | 9 |
| 474 | ENSG00000165061 | 259.1483 | 2.076454 | 0.324383 | 6.401245 | 1.54E-10 | 1.01E-08 | 5.411283 | 26.21378 | 10.89679 | 38.62818 | ZMAT4 | | 8 |
| 475 | ENSG00000165072 | 97.69307 | 1.369761 | 0.409095 | 3.348274 | 0.000813 | 0.012923 | 1.428004 | 5.762361 | 2.609743 | 4.203362 | MAMDC2 | | 9 |
| 476 | ENSG00000165175 | 660.5846 | -1.07777 | 0.177216 | -6.08171 | 1.19E-09 | 6.76E-08 | 26.27602 | 11.08823 | 24.94845 | 11.99173 | MID1IP1 | X | |
| 477 | ENSG00000165186 | 199.053 | -2.69545 | 0.384592 | -7.00859 | 2.41E-12 | 1.99E-10 | 2.8052 | 0.717901 | 3.483933 | 0.228229 | PTCHD1 | X | |
| 478 | ENSG00000165194 | 3801.969 | 1.373754 | 0.125906 | 10.91095 | 1.02E-27 | 2.89E-25 | 26.0138 | 71.58822 | 28.37078 | 62.75145 | PCDH19 | X | |
| 479 | ENSG00000165309 | 50.84914 | -1.96315 | 0.539603 | -3.63813 | 0.000275 | 0.005182 | 6.083285 | 1.002018 | 5.573806 | 1.871746 | ARMC3 | | 10 |
| 480 | ENSG00000165588 | 939.9592 | -1.1143 | 0.187238 | -5.95125 | 2.66E-09 | 1.41E-07 | 63.88305 | 37.08557 | 81.70395 | 27.17448 | OTX2 | | 14 |
| 481 | ENSG00000165655 | 279.3068 | -1.67138 | 0.299722 | -5.57645 | 2.45E-08 | 1.13E-06 | 15.65565 | 6.401474 | 15.53146 | 3.053352 | ZNF503 | | 10 |
| 482 | ENSG00000165757 | 152.1267 | 1.700957 | 0.343178 | 4.95648 | 7.18E-07 | 2.47E-05 | 0.783626 | 2.17572 | 1.002052 | 3.321557 | JCAD | | 10 |
| 483 | ENSG00000166016 | 171.6768 | -1.04788 | 0.295465 | -3.54655 | 0.00039 | 0.007005 | 4.882346 | 2.291913 | 5.930706 | 2.664916 | ABTB2 | | 11 |
| 484 | ENSG00000166073 | 259.7232 | 1.073521 | 0.29011 | 3.700391 | 0.000215 | 0.004223 | 3.757312 | 8.894105 | 6.395027 | 11.35553 | GPR176 | | 15 |
| 485 | ENSG00000166265 | 209.2833 | -1.10282 | 0.321771 | -3.42733 | 0.00061 | 0.010157 | 13.22255 | 6.341241 | 8.996789 | 3.631283 | CYYR1 | | 21 |
| 486 | ENSG00000166426 | 4636.341 | 3.451941 | 0.141519 | 24.39204 | 2.08E-131 | 2.00E-127 | 161.8778 | 1679.612 | 187.3851 | 1951.894 | CRABP1 | | 15 |
| 487 | ENSG00000166450 | 4276.985 | 2.256479 | 0.173412 | 13.01222 | 1.04E-38 | 4.67E-36 | 21.08272 | 115.7764 | 32.76659 | 128.0607 | PRTG | | 15 |
| 488 | ENSG00000166473 | 43.00885 | 2.628238 | 0.64744 | 4.059433 | 4.92E-05 | 0.001152 | 0.209596 | 2.401084 | 0.720926 | 3.437475 | PKD1L2 | | 16 |
| 489 | ENSG00000166770 | 49.80172 | -6.40598 | 0.973908 | -6.5776 | 4.78E-11 | 3.32E-09 | 7.388136 | 0.075552 | 5.767596 | 0.072176 | ZNF667-A! | | 19 |
| 490 | ENSG00000166801 | 133.4844 | 1.412284 | 0.340547 | 4.147101 | 3.37E-05 | 0.000818 | 2.128358 | 6.442825 | 3.193998 | 6.97002 | FAM111A | | 11 |
| 491 | ENSG00000166888 | 54.42238 | 1.920833 | 0.51307 | 3.743802 | 0.000181 | 0.003657 | 0.928946 | 3.671447 | 1.360438 | 4.513945 | STAT6 | | 12 |
| 492 | ENSG00000167114 | 269.8132 | 1.367903 | 0.241673 | 5.660144 | 1.51E-08 | 7.09E-07 | 5.750247 | 13.50162 | 5.544806 | 14.22581 | SLC27A4 | | 9 |
| 493 | ENSG00000167118 | 347.4779 | 1.054557 | 0.265344 | 3.974296 | 7.06E-05 | 0.001593 | 14.33162 | 22.64484 | 8.339141 | 21.99318 | URM1 | | 9 |
| 494 | ENSG00000167123 | 317.5185 | 3.611562 | 0.299145 | 12.07294 | 1.47E-33 | 5.23E-31 | 1.515637 | 18.56328 | 2.372411 | 26.82211 | CERCAM | | 9 |
| 495 | ENSG00000167178 | 229.3048 | -2.9699 | 0.277302 | -10.71 | 9.14E-27 | 2.44E-24 | 15.79182 | 1.828671 | 15.454 | 1.980717 | ISLR2 | | 15 |
| 496 | ENSG00000167553 | 306.1678 | 1.063231 | 0.29014 | 3.664544 | 0.000248 | 0.004764 | 22.23326 | 42.84142 | 33.86105 | 68.12178 | TUBA1C | | 12 |
| 497 | ENSG00000167555 | 133.829 | -1.66187 | 0.365923 | -4.5416 | 5.58E-06 | 0.000158 | 18.2891 | 3.909229 | 18.06206 | 7.161593 | ZNF528 | | 19 |
| 498 | ENSG00000167723 | 64.67559 | 2.179167 | 0.475309 | 4.584741 | 4.55E-06 | 0.000131 | 0.655396 | 2.608938 | 0.778504 | 3.555949 | TRPV3 | | 17 |
| 499 | ENSG00000167785 | 289.4451 | 1.345841 | 0.279863 | 4.808927 | 1.52E-06 | 4.87E-05 | 9.131114 | 15.82568 | 5.657594 | 19.52693 | ZNF558 | | 19 |
| 500 | ENSG00000167912 | 105.2521 | -1.22238 | 0.365846 | -3.34123 | 0.000834 | 0.01319 | 14.67885 | 7.008237 | 14.56811 | 5.012885 | TOX-DT | | 8 |
| 501 | ENSG00000168026 | 51.45093 | -1.54379 | 0.520635 | -2.9652 | 0.003025 | 0.038427 | 5.608675 | 1.31539 | 5.051724 | 2.150419 | TTC21A | | 3 |
| 502 | ENSG00000168264 | 1223.049 | 1.17496 | 0.173433 | 6.774704 | 1.25E-11 | 9.34E-10 | 33.9114 | 67.21262 | 25.4813 | 60.6266 | IRF2BP2 | | 1 |
| 503 | ENSG00000168348 | 30.18699 | -2.40604 | 0.682294 | -3.5264 | 0.000421 | 0.007493 | 2.073685 | 0.434004 | 2.098709 | 0.322488 | INSM2 | | 14 |
| 504 | ENSG00000168505 | 101.6089 | 2.973524 | 0.432017 | 6.882893 | 5.86E-12 | 4.55E-10 | 0.963548 | 12.52769 | 1.951263 | 9.511067 | GBX2 | | 2 |
| 505 | ENSG00000168542 | 856.357 | 5.343422 | 0.228233 | 23.41215 | 3.21E-121 | 1.55E-117 | 0.89901 | 32.58599 | 0.846382 | 34.77203 | COL3A1 | | 2 |
| 506 | ENSG00000168646 | 573.5151 | 1.347135 | 0.190333 | 7.077781 | 1.46E-12 | 1.25E-10 | 15.08665 | 42.48758 | 16.61125 | 34.45338 | AXIN2 | | 17 |
| 507 | ENSG00000168734 | 197.569 | -1.13588 | 0.275087 | -4.12916 | 3.64E-05 | 0.000879 | 33.3153 | 13.07037 | 31.26153 | 14.83325 | PKIG | | 20 |
| 508 | ENSG00000168785 | 578.2503 | -1.09701 | 0.209388 | -5.23911 | 1.61E-07 | 6.24E-06 | 31.64511 | 11.54206 | 23.60981 | 13.03876 | TSPAN5 | | 4 |
| 509 | ENSG00000168843 | 214.2311 | -1.81093 | 0.295092 | -6.13681 | 8.42E-10 | 4.93E-08 | 7.260185 | 2.502251 | 11.40093 | 2.554969 | FSTL5 | | 4 |
| 510 | ENSG00000168952 | 83.66621 | 1.360119 | 0.40857 | 3.328971 | 0.000872 | 0.013706 | 1.552473 | 4.376798 | 1.519573 | 3.170882 | STXBP6 | | 14 |
| 511 | ENSG00000168959 | 27.28428 | -2.36967 | 0.775017 | -3.05757 | 0.002231 | 0.029927 | 1.374827 | 0.109627 | 0.679082 | 0.270563 | GRM5 | | 11 |
| 512 | ENSG00000168994 | 43.63075 | 1.886262 | 0.581624 | 3.243094 | 0.001182 | 0.017695 | 1.097249 | 3.559707 | 1.548772 | 5.667659 | PXDC1 | | 6 |
| 513 | ENSG00000169071 | 119.5142 | 1.093055 | 0.366661 | 2.981105 | 0.002872 | 0.036777 | 3.152477 | 6.745483 | 2.282481 | 4.418659 | ROR2 | | 9 |
| 514 | ENSG00000169184 | 1288.746 | 1.196171 | 0.153575 | 7.788854 | 6.76E-15 | 7.61E-13 | 11.98926 | 25.77893 | 10.46675 | 23.30322 | MN1 | | 22 |
| 515 | ENSG00000169218 | 105.8102 | -1.69657 | 0.380372 | -4.46027 | 8.19E-06 | 0.000224 | 6.589494 | 2.642978 | 8.041225 | 1.690335 | RSPO1 | | 1 |
| 516 | ENSG00000169302 | 82.31844 | -2.44411 | 0.449336 | -5.43937 | 5.35E-08 | 2.30E-06 | 4.292914 | 0.847247 | 3.094127 | 0.467923 | STK32A | | 5 |
| 517 | ENSG00000169432 | 175.5244 | -1.04421 | 0.313917 | -3.3264 | 0.00088 | 0.013822 | 6.20981 | 2.958078 | 4.590968 | 2.073965 | SCN9A | | 2 |
| 518 | ENSG00000169436 | 86.49488 | -1.17499 | 0.389948 | -3.01319 | 0.002585 | 0.033869 | 4.057688 | 1.82461 | 4.731587 | 1.871561 | COL22A1 | | 8 |
| 519 | ENSG00000169744 | 267.3756 | 1.090629 | 0.257779 | 4.230876 | 2.33E-05 | 0.000586 | 9.380692 | 21.10642 | 12.93467 | 23.95519 | LDB2 | | 4 |
| 520 | ENSG00000169836 | 167.1335 | 1.812782 | 0.302672 | 5.989267 | 2.11E-09 | 1.14E-07 | 1.391615 | 5.277283 | 1.757862 | 5.253896 | TACR3 | | 4 |
| 521 | ENSG00000169840 | 36.7197 | 2.209337 | 0.623979 | 3.540724 | 0.000399 | 0.00714 | 0.814554 | 4.65724 | 0.986349 | 3.315182 | GSX1 | | 13 |
| 522 | ENSG00000169851 | 505.0274 | -1.09782 | 0.266818 | -4.11449 | 3.88E-05 | 0.000931 | 13.28262 | 8.411472 | 14.3259 | 4.017935 | PCDH7 | | 4 |
| 523 | ENSG00000170091 | 372.0241 | -2.03538 | 0.268236 | -7.58805 | 3.25E-14 | 3.36E-12 | 29.32344 | 9.895145 | 34.92795 | 5.239237 | NSG2 | | 5 |
| 524 | ENSG00000170214 | 19.43837 | -3.42371 | 0.923973 | -3.70542 | 0.000211 | 0.004162 | 1.703556 | 0.152302 | 1.718926 | 0.151083 | ADRA1B | | 5 |
| 525 | ENSG00000170370 | 174.7915 | -2.51086 | 0.32203 | -7.79697 | 6.34E-15 | 7.18E-13 | 17.30766 | 3.894638 | 20.12512 | 2.403234 | EMX2 | | 10 |
| 526 | ENSG00000170382 | 216.225 | -1.54616 | 0.270892 | -5.70767 | 1.15E-08 | 5.47E-07 | 9.304473 | 2.953141 | 10.57258 | 3.502006 | LRRN2 | | 1 |
| 527 | ENSG00000170396 | 67.75357 | -1.78549 | 0.456223 | -3.91362 | 9.09E-05 | 0.00198 | 2.970224 | 0.67049 | 2.358957 | 0.79486 | ZNF804A | | 2 |
| 528 | ENSG00000170542 | 72.92879 | 1.747936 | 0.524979 | 3.329538 | 0.00087 | 0.013689 | 0.508923 | 2.091766 | 1.366986 | 3.843644 | SERPINB9 | | 6 |
| 529 | ENSG00000170549 | 172.0303 | 2.304689 | 0.305212 | 7.551098 | 4.32E-14 | 4.42E-12 | 3.119289 | 17.08409 | 3.755029 | 15.30257 | IRX1 | | 5 |
| 530 | ENSG00000170561 | 558.7647 | 2.853693 | 0.204984 | 13.92157 | 4.68E-44 | 2.73E-41 | 4.760294 | 37.88029 | 6.293841 | 38.15804 | IRX2 | | 5 |
| 531 | ENSG00000170624 | 30.49049 | -3.98773 | 0.787484 | -5.06388 | 4.11E-07 | 1.45E-05 | 1.140512 | 0.025136 | 0.754614 | 0.083602 | SGCD | | 5 |
| 532 | ENSG00000170629 | 118.573 | -1.10271 | 0.339314 | -3.24983 | 0.001155 | 0.01739 | 6.009354 | 2.685167 | 6.654846 | 2.913733 | DPY19L2P2 | | 7 |
| 533 | ENSG00000170631 | 92.14477 | -1.17231 | 0.378463 | -3.09755 | 0.001951 | 0.026765 | 7.430691 | 3.124551 | 6.974094 | 2.96907 | ZNF16 | | 8 |
| 534 | ENSG00000170775 | 67.91265 | -1.65226 | 0.45293 | -3.64793 | 0.000264 | 0.005028 | 1.916384 | 0.613645 | 2.525882 | 0.723593 | GPR37 | | 7 |
| 535 | ENSG00000170893 | 171.3763 | -3.02484 | 0.365814 | -8.26878 | 1.35E-16 | 1.97E-14 | 17.5135 | 3.056217 | 33.30419 | 2.876711 | TRH | | 3 |
| 536 | ENSG00000170921 | 1426.894 | -1.25614 | 0.1639 | -7.66404 | 1.80E-14 | 1.93E-12 | 37.37678 | 13.35023 | 38.83931 | 16.96581 | TANC2 | | 17 |
| 537 | ENSG00000170959 | 597.6803 | -3.27463 | 0.212608 | -15.4022 | 1.58E-53 | 1.69E-50 | 38.89321 | 4.854614 | 41.909 | 3.25257 | DCDC1 | | 11 |
| 538 | ENSG00000171004 | 805.4468 | -1.02048 | 0.176375 | -5.78582 | 7.22E-09 | 3.58E-07 | 28.68485 | 12.54461 | 30.58267 | 15.16538 | HS6ST2 | X | |
| 539 | ENSG00000171189 | 113.7387 | -1.81775 | 0.356138 | -5.10406 | 3.32E-07 | 1.20E-05 | 6.510842 | 2.112145 | 7.721322 | 1.737777 | GRIK1 | | 21 |
| 540 | ENSG00000171388 | 145.6191 | 1.596032 | 0.322173 | 4.953954 | 7.27E-07 | 2.49E-05 | 2.233434 | 7.786976 | 3.056632 | 7.441273 | APLN | X | |
| 541 | ENSG00000171451 | 701.2451 | 1.893894 | 0.230051 | 8.232482 | 1.83E-16 | 2.60E-14 | 3.102949 | 9.955532 | 4.070888 | 15.27221 | DSEL | | 18 |
| 542 | ENSG00000171502 | 1010.173 | 3.886916 | 0.481216 | 8.077274 | 6.62E-16 | 8.55E-14 | 1.689977 | 39.67133 | 3.342572 | 31.18643 | COL24A1 | | 1 |
| 543 | ENSG00000171587 | 36.76939 | -1.91178 | 0.613568 | -3.11584 | 0.001834 | 0.025452 | 0.88711 | 0.271595 | 0.822528 | 0.16605 | DSCAM | | 21 |
| 544 | ENSG00000171757 | 70.068 | -1.70978 | 0.458365 | -3.73017 | 0.000191 | 0.003824 | 6.647753 | 1.521861 | 4.262016 | 1.648255 | LRRC34 | | 3 |
| 545 | ENSG00000171864 | 54.24119 | 4.023117 | 0.733953 | 5.481433 | 4.22E-08 | 1.85E-06 | 0.15618 | 4.641576 | 0.208005 | 1.093959 | PRND | | 20 |

| | | | | | | | | | | | | | |
|---|---|---|---|---|---|---|---|---|---|---|---|---|---|
| 546 | ENSG00000171951 | 1099.547 | -1.5514 | 0.200435 | -7.7402 | 9.93E-15 | 1.09E-12 | 63.08847 | 28.06615 | 100.7446 | 25.10385 | SCG2 | 2 |
| 547 | ENSG00000172014 | 51.89031 | -1.5943 | 0.547945 | -2.9096 | 0.003619 | 0.044047 | 3.403016 | 0.868414 | 1.737053 | 0.765545 | ANKRD20A | 9 |
| 548 | ENSG00000172031 | 20.50369 | 2.537266 | 0.884337 | 2.869116 | 0.004116 | 0.048994 | 0.19637 | 3.195611 | 0.934222 | 3.06217 | EPHX4 | 1 |
| 549 | ENSG00000172123 | 89.7865 | 1.46891 | 0.409481 | 3.587248 | 0.000334 | 0.006137 | 2.221762 | 7.698939 | 3.757536 | 8.004987 | SLFN12 | 17 |
| 550 | ENSG00000172201 | 1979.199 | -1.44351 | 0.194486 | -7.42217 | 1.15E-13 | 1.12E-11 | 106.2791 | 30.16617 | 66.67871 | 30.53827 | ID4 | 6 |
| 551 | ENSG00000172260 | 121.9986 | -1.488 | 0.348724 | -4.26699 | 1.98E-05 | 0.000506 | 2.494426 | 0.968995 | 3.434577 | 1.036123 | NEGR1 | 1 |
| 552 | ENSG00000172318 | 262.9688 | 1.160049 | 0.25024 | 4.635737 | 3.56E-06 | 0.000106 | 3.151998 | 7.886678 | 4.142906 | 7.629265 | B3GALT1 | 2 |
| 553 | ENSG00000172461 | 1042.109 | -1.2202 | 0.168762 | -7.23034 | 4.82E-13 | 4.42E-11 | 12.80419 | 6.368458 | 16.25421 | 5.519432 | FUT9 | 6 |
| 554 | ENSG00000172572 | 2907.044 | 1.156714 | 0.148224 | 7.803822 | 6.01E-15 | 6.84E-13 | 26.13747 | 57.53396 | 32.27201 | 66.13325 | PDE3A | 12 |
| 555 | ENSG00000172716 | 28.68668 | 4.95452 | 1.038479 | 4.770939 | 1.83E-06 | 5.79E-05 | 0 | 1.513779 | 0.09517 | 2.702203 | SLFN11 | 17 |
| 556 | ENSG00000172771 | 148.5475 | -1.26653 | 0.323789 | -3.9116 | 9.17E-05 | 0.001994 | 9.580999 | 3.389075 | 6.945736 | 3.147488 | EFCAB12 | 3 |
| 557 | ENSG00000172965 | 557.9173 | 1.988579 | 0.222169 | 8.950742 | 3.53E-19 | 5.96E-17 | 14.0471 | 60.13629 | 21.08538 | 71.74136 | MIR4435-2 | 2 |
| 558 | ENSG00000173041 | 199.8938 | -1.31857 | 0.282067 | -4.67466 | 2.94E-06 | 8.97E-05 | 17.3181 | 7.141535 | 15.25894 | 5.420694 | ZNF680 | 7 |
| 559 | ENSG00000173193 | 275.2389 | 1.452682 | 0.338779 | 4.287997 | 1.80E-05 | 0.000468 | 1.651071 | 4.51383 | 3.242072 | 8.132227 | PARP14 | 3 |
| 560 | ENSG00000173208 | 79.81894 | -1.73756 | 0.443847 | -3.91477 | 9.05E-05 | 0.001975 | 1.659607 | 0.550945 | 2.771977 | 0.702868 | ABCD2 | 12 |
| 561 | ENSG00000173218 | 183.9687 | -1.07764 | 0.299316 | -3.60035 | 0.000318 | 0.005875 | 4.707 | 2.284229 | 3.835504 | 1.610447 | VANGL1 | 1 |
| 562 | ENSG00000173376 | 484.4419 | 1.107515 | 0.209028 | 5.298416 | 1.17E-07 | 4.68E-06 | 12.03631 | 30.23951 | 13.57974 | 22.51214 | NDNF | 4 |
| 563 | ENSG00000173421 | 19.36391 | -3.3087 | 0.862027 | -3.83828 | 0.000124 | 0.002623 | 3.029117 | 0.232277 | 3.147938 | 0.347778 | IHO1 | 3 |
| 564 | ENSG00000173482 | 1138.471 | -1.39308 | 0.154496 | -9.01694 | 1.93E-19 | 3.32E-17 | 49.85537 | 20.17486 | 52.21166 | 16.93933 | PTPRM | 18 |
| 565 | ENSG00000173706 | 1025.276 | 1.598208 | 0.204588 | 7.81185 | 5.64E-15 | 6.46E-13 | 7.996727 | 32.42613 | 13.27968 | 28.87888 | HEG1 | 3 |
| 566 | ENSG00000173947 | 55.75422 | -1.54468 | 0.499848 | -3.0903 | 0.002 | 0.027313 | 5.282312 | 2.359426 | 6.419422 | 1.555711 | PIFO | 1 |
| 567 | ENSG00000174080 | 10.23572 | -7.01662 | 1.826841 | -3.84085 | 0.000123 | 0.002602 | 0.742475 | 0 | 2.107506 | 0 | CTSF | 11 |
| 568 | ENSG00000174498 | 1206.387 | 1.184639 | 0.189249 | 6.259685 | 3.86E-10 | 2.34E-08 | 22.07112 | 52.53575 | 18.45928 | 35.70624 | IGDCC3 | 15 |
| 569 | ENSG00000174721 | 1495.014 | 1.04901 | 0.145667 | 7.201427 | 5.96E-13 | 5.38E-11 | 48.11338 | 95.18238 | 43.70355 | 86.01513 | FGFBP3 | 10 |
| 570 | ENSG00000174948 | 20.73315 | -3.71902 | 1.00818 | -3.68885 | 0.000225 | 0.004384 | 0.353396 | 0.086969 | 1.239662 | 0.032191 | GPR149 | 3 |
| 571 | ENSG00000175093 | 350.0363 | 1.363858 | 0.226786 | 6.013861 | 1.81E-09 | 9.90E-08 | 13.64147 | 33.22752 | 11.36787 | 28.32856 | SPSB4 | 3 |
| 572 | ENSG00000175161 | 1058.194 | -2.6364 | 0.198303 | -13.2948 | 2.48E-40 | 1.22E-37 | 61.36153 | 9.633942 | 46.04231 | 6.959431 | CADM2 | 3 |
| 573 | ENSG00000175344 | 818.4739 | 1.072478 | 0.163006 | 6.579385 | 4.72E-11 | 3.29E-09 | 13.9591 | 28.9257 | 13.67811 | 26.4836 | CHRNA7 | 15 |
| 574 | ENSG00000175426 | 189.8629 | -1.16813 | 0.338102 | -3.45497 | 0.00055 | 0.009382 | 5.539207 | 3.633153 | 6.930359 | 1.731667 | PCSK1 | 5 |
| 575 | ENSG00000175445 | 137.8384 | 1.318548 | 0.331697 | 3.975158 | 7.03E-05 | 0.001591 | 3.074615 | 7.236512 | 3.592139 | 8.622887 | LPL | 8 |
| 576 | ENSG00000175785 | 142.7799 | 1.14924 | 0.340176 | 3.378368 | 0.000729 | 0.011823 | 2.365718 | 7.185924 | 3.294277 | 4.830266 | PRIMA1 | 14 |
| 577 | ENSG00000175874 | 23.66059 | -5.08641 | 1.055885 | -4.8172 | 1.46E-06 | 4.69E-05 | 1.43225 | 0 | 1.446333 | 0.056211 | CREG2 | 2 |
| 578 | ENSG00000176165 | 554.2498 | -5.1537 | 0.95676 | -5.38661 | 7.18E-08 | 3.01E-06 | 38.40152 | 1.627422 | 33.76147 | 0.341275 | FOXG1 | 14 |
| 579 | ENSG00000176170 | 46.66173 | 1.945302 | 0.556045 | 3.498462 | 0.000468 | 0.00818 | 1.613453 | 4.845786 | 1.119567 | 4.909583 | SPHK1 | 17 |
| 580 | ENSG00000176204 | 177.6358 | -1.67567 | 0.294331 | -5.69314 | 1.25E-08 | 5.91E-07 | 9.470084 | 2.843225 | 11.1207 | 3.28571 | LRRTM4 | 2 |
| 581 | ENSG00000176293 | 36.26141 | -8.63513 | 1.552707 | -5.56134 | 2.68E-08 | 1.22E-06 | 2.581207 | 0 | 3.206763 | 0 | ZNF135 | 19 |
| 582 | ENSG00000176399 | 329.54 | -3.88753 | 0.266803 | -14.5708 | 4.31E-48 | 3.46E-45 | 13.45536 | 0.838583 | 11.63621 | 0.779935 | DMRTA1 | 9 |
| 583 | ENSG00000176697 | 134.2272 | -1.07198 | 0.348002 | -3.0804 | 0.002067 | 0.028058 | 3.910138 | 1.974346 | 5.973359 | 2.468971 | BDNF | 11 |
| 584 | ENSG00000176771 | 1177.532 | 1.519646 | 0.152182 | 9.985684 | 1.76E-23 | 3.72E-21 | 12.21024 | 37.27812 | 13.47807 | 32.96594 | NCKAP5 | 2 |
| 585 | ENSG00000176842 | 423.7693 | 2.49726 | 0.214097 | 11.66417 | 1.94E-31 | 6.23E-29 | 6.826212 | 39.79065 | 7.058973 | 34.98399 | IRX5 | 16 |
| 586 | ENSG00000176907 | 108.7119 | -2.28336 | 0.397106 | -5.74998 | 8.93E-09 | 4.34E-07 | 13.94239 | 1.812344 | 10.87882 | 2.977635 | TCIM | 8 |
| 587 | ENSG00000176945 | 93.9991 | -1.32287 | 0.45433 | -2.9117 | 0.003595 | 0.043898 | 6.244195 | 1.256079 | 5.035018 | 3.064936 | MUC20 | 3 |
| 588 | ENSG00000177182 | 125.735 | -2.07131 | 0.435955 | -4.75119 | 2.02E-06 | 6.33E-05 | 11.02747 | 3.932711 | 10.85774 | 1.241675 | CLVS1 | 8 |
| 589 | ENSG00000177283 | 46.24198 | -2.2157 | 0.581411 | -3.81091 | 0.000138 | 0.002884 | 2.790884 | 0.39236 | 1.54631 | 0.492259 | FZD8 | 10 |
| 590 | ENSG00000177469 | 194.8729 | 1.557735 | 0.314503 | 4.953005 | 7.31E-07 | 2.50E-05 | 2.492189 | 10.74139 | 3.973704 | 7.45674 | CAVIN1 | 17 |
| 591 | ENSG00000177508 | 342.3458 | 2.258942 | 0.277183 | 8.149627 | 3.65E-16 | 4.88E-14 | 3.880395 | 28.62777 | 7.404778 | 22.94237 | IRX3 | 16 |
| 592 | ENSG00000177570 | 417.8084 | -1.66107 | 0.229105 | -7.25024 | 4.16E-13 | 3.85E-11 | 15.11459 | 6.083978 | 19.31554 | 4.382717 | SAMD12 | 8 |
| 593 | ENSG00000177606 | 1392.236 | 1.846471 | 0.165338 | 11.16788 | 5.86E-29 | 1.76E-26 | 21.89871 | 67.1799 | 22.24537 | 83.566 | JUN | 1 |
| 594 | ENSG00000177675 | 10.04279 | -4.70881 | 1.318678 | -3.57086 | 0.000356 | 0.00646 | 0.821662 | 0.026473 | 1.642822 | 0.058841 | CD163L1 | 12 |
| 595 | ENSG00000178031 | 546.9414 | -3.24983 | 0.257714 | -12.6102 | 1.85E-36 | 7.44E-34 | 33.44766 | 4.25566 | 28.34003 | 2.110421 | ADAMTSL1 | 9 |
| 596 | ENSG00000178033 | 265.562 | 3.403858 | 0.290047 | 11.73554 | 8.38E-32 | 2.88E-29 | 0.427796 | 4.790234 | 0.616229 | 5.710608 | CALHM5 | 6 |
| 597 | ENSG00000178187 | 19.51839 | -4.47312 | 1.041728 | -4.29394 | 1.76E-05 | 0.000457 | 2.142029 | 0.110047 | 2.276161 | 0.081936 | ZNF454 | 5 |
| 598 | ENSG00000178343 | 68.93731 | -2.26201 | 0.504292 | -4.48553 | 7.27E-06 | 0.000201 | 3.823568 | 1.271638 | 8.184724 | 1.111507 | SHISA3 | 4 |
| 599 | ENSG00000178403 | 266.118 | -2.90516 | 0.762498 | -3.81005 | 0.000139 | 0.002891 | 31.78972 | 4.705404 | 17.88628 | 1.710287 | NEUROG2 | 4 |
| 600 | ENSG00000178404 | 50.46419 | -1.89176 | 0.578647 | -3.26928 | 0.001078 | 0.016391 | 5.036564 | 2.12518 | 5.60508 | 0.687863 | CEP295NL | 17 |
| 601 | ENSG00000178538 | 207.1205 | -2.34835 | 0.284617 | -8.25093 | 1.57E-16 | 2.24E-14 | 15.62905 | 3.254398 | 15.15523 | 2.559736 | CA8 | 8 |
| 602 | ENSG00000178597 | 72.67866 | -2.38507 | 0.453867 | -5.255 | 1.48E-07 | 5.78E-06 | 3.340556 | 0.600173 | 2.673431 | 0.502627 | PSAPL1 | 4 |
| 603 | ENSG00000178878 | 231.0508 | 1.480102 | 0.303428 | 4.877936 | 1.07E-06 | 3.56E-05 | 3.145528 | 7.656194 | 4.280139 | 11.91457 | APOLD1 | 12 |
| 604 | ENSG00000179455 | 351.4724 | -1.41188 | 0.219525 | -6.43152 | 1.26E-10 | 8.38E-09 | 31.47001 | 11.68985 | 29.34697 | 10.13405 | MKRN3 | 15 |
| 605 | ENSG00000179520 | 44.72722 | -2.40729 | 0.79067 | -3.04463 | 0.00233 | 0.030965 | 0.686641 | 0.311274 | 5.121344 | 0.711604 | SLC17A8 | 12 |
| 606 | ENSG00000179673 | 28.35434 | 2.346437 | 0.716538 | 3.274685 | 0.001058 | 0.016145 | 0.823529 | 5.787841 | 1.567344 | 5.79214 | RPRML | 17 |
| 607 | ENSG00000179833 | 633.093 | 1.00891 | 0.180338 | 5.594548 | 2.21E-08 | 1.02E-06 | 10.43463 | 19.05876 | 9.194671 | 18.58428 | SERTAD2 | 2 |
| 608 | ENSG00000179915 | 478.736 | -1.33927 | 0.19994 | -6.69835 | 2.11E-11 | 1.55E-09 | 24.50783 | 10.13681 | 23.94953 | 8.139041 | NRXN1 | 2 |
| 609 | ENSG00000180447 | 431.2601 | 1.258017 | 0.206864 | 6.081373 | 1.19E-09 | 6.76E-08 | 8.81514 | 23.56166 | 10.26248 | 19.96613 | GAS1 | 9 |
| 610 | ENSG00000180613 | 36.59187 | -5.49008 | 1.021177 | -5.37623 | 7.61E-08 | 3.16E-06 | 7.12081 | 0.248681 | 5.460886 | 0 | GSX2 | 4 |
| 611 | ENSG00000180628 | 859.4637 | 1.739849 | 0.171411 | 10.15016 | 3.31E-24 | 7.58E-22 | 13.99329 | 40.48106 | 12.13715 | 42.36485 | PCGF5 | 10 |
| 612 | ENSG00000180730 | 95.61492 | -2.77807 | 0.442569 | -6.27714 | 3.45E-10 | 2.13E-08 | 5.705433 | 0.446601 | 4.379119 | 0.922603 | SHISA2 | 13 |
| 613 | ENSG00000180828 | 38.67683 | -2.39035 | 0.747435 | -3.19807 | 0.001383 | 0.020157 | 2.629976 | 0.799077 | 2.023652 | 0.084744 | BHLHE22 | 8 |
| 614 | ENSG00000181007 | 361.274 | -1.40601 | 0.224221 | -6.27062 | 3.60E-10 | 2.20E-08 | 13.80242 | 4.386481 | 13.39625 | 5.343858 | ZFP82 | 19 |
| 615 | ENSG00000181215 | 23.52648 | -2.28482 | 0.765526 | -2.98464 | 0.002839 | 0.036477 | 0.289114 | 0.07141 | 0.393004 | 0.062125 | C4orf50 | 4 |
| 616 | ENSG00000181234 | 204.6831 | 1.777088 | 0.298261 | 5.958168 | 2.55E-09 | 1.36E-07 | 1.537941 | 6.684198 | 2.543409 | 6.62236 | TMEM132C | 12 |
| 617 | ENSG00000181744 | 497.2818 | -1.00769 | 0.206023 | -4.89116 | 1.00E-06 | 3.35E-05 | 28.22664 | 14.03069 | 24.18514 | 10.92385 | DIPK2A | 3 |
| 618 | ENSG00000181965 | 139.7883 | -3.10885 | 0.436012 | -7.13018 | 1.00E-12 | 8.77E-11 | 19.10028 | 3.43367 | 18.79154 | 0.895663 | NEUROG1 | 5 |
| 619 | ENSG00000182021 | 108.8812 | -1.4662 | 0.445455 | -3.29146 | 0.000997 | 0.015407 | 11.75836 | 4.325073 | 6.185659 | 1.99015 | | 9 |
| 620 | ENSG00000182050 | 86.55034 | -2.25077 | 0.423239 | -5.31797 | 1.05E-07 | 4.26E-06 | 2.621401 | 0.426399 | 2.427753 | 0.592091 | MGAT4C | 12 |
| 621 | ENSG00000182158 | 2047.365 | 1.238644 | 0.141778 | 8.736508 | 2.40E-18 | 3.89E-16 | 18.79684 | 46.74306 | 22.77735 | 46.59091 | CREB3L2 | 7 |
| 622 | ENSG00000182165 | 130.5966 | -1.43893 | 0.351618 | -4.09231 | 4.27E-05 | 0.001015 | 13.77067 | 5.376066 | 20.72021 | 6.633422 | TP53TG1 | 7 |
| 623 | ENSG00000182168 | 619.0543 | 1.245092 | 0.183042 | 6.80222 | 1.03E-11 | 7.87E-10 | 5.456868 | 11.70154 | 5.515316 | 13.02321 | UNC5C | 4 |

| | | | | | | | | | | | | | | |
|---|---|---|---|---|---|---|---|---|---|---|---|---|---|---|
| 624 | ENSG00000182240 | 266.5627 | 1.411351 | 0.2568 | 5.495909 | 3.89E-08 | 1.72E-06 | 3.848161 | 10.36321 | 5.029934 | 12.08066 | BACE2 | | 21 |
| 625 | ENSG00000182463 | 244.8927 | 1.042533 | 0.249098 | 4.185233 | 2.85E-05 | 0.000698 | 3.795829 | 7.892568 | 3.826656 | 7.076936 | TSHZ2 | | 20 |
| 626 | ENSG00000182601 | 103.6718 | -1.91886 | 0.376631 | -5.09478 | 3.49E-07 | 1.25E-05 | 7.055505 | 1.558038 | 5.548482 | 1.607993 | HS3ST4 | | 16 |
| 627 | ENSG00000182667 | 93.36492 | -1.94264 | 0.415092 | -4.68002 | 2.87E-06 | 8.75E-05 | 8.696999 | 3.07211 | 14.67319 | 2.733791 | NTM | | 11 |
| 628 | ENSG00000183023 | 711.7626 | -1.08734 | 0.170781 | -6.36685 | 1.93E-10 | 1.25E-08 | 25.39028 | 12.02326 | 25.18877 | 10.6936 | SLC8A1 | | 2 |
| 629 | ENSG00000183148 | 31.77916 | -2.93635 | 0.886181 | -3.31348 | 0.000921 | 0.014411 | 3.812347 | 0.421945 | 0.543943 | 0.130451 | ANKRD20A | | 9 |
| 630 | ENSG00000183454 | 567.4085 | -1.92132 | 0.24901 | -7.71581 | 1.20E-14 | 1.31E-12 | 12.04486 | 2.168609 | 7.285285 | 2.691448 | GRIN2A | | 16 |
| 631 | ENSG00000183579 | 813.6515 | 1.14408 | 0.16675 | 6.861044 | 6.84E-12 | 5.28E-10 | 8.044142 | 18.85472 | 8.573086 | 16.17779 | ZNRF3 | | 22 |
| 632 | ENSG00000183690 | 19.05646 | -2.93101 | 0.928979 | -3.15509 | 0.001605 | 0.022857 | 0.813818 | 0.228762 | 1.622151 | 0.084914 | EFHC2 | X | |
| 633 | ENSG00000183762 | 285.6511 | 1.193438 | 0.268734 | 4.440956 | 8.96E-06 | 0.000244 | 2.951518 | 7.261881 | 4.536321 | 8.973646 | KREMEN1 | | 22 |
| 634 | ENSG00000183960 | 554.9085 | 1.265778 | 0.199456 | 6.346144 | 2.21E-10 | 1.41E-08 | 7.528895 | 19.2836 | 7.185831 | 14.54594 | KCNH8 | | 3 |
| 635 | ENSG00000184156 | 1009.213 | -1.93197 | 0.205121 | -9.4187 | 4.57E-21 | 8.53E-19 | 20.31653 | 5.357363 | 15.28838 | 3.610354 | KCNQ3 | | 8 |
| 636 | ENSG00000184185 | 58.37964 | 1.394898 | 0.476416 | 2.927897 | 0.003413 | 0.042156 | 0.786323 | 1.879389 | 0.739091 | 1.931795 | KCNJ12 | | 17 |
| 637 | ENSG00000184221 | 14.50065 | 3.429658 | 1.075989 | 3.187448 | 0.001435 | 0.020817 | 0.138371 | 2.624923 | 0.21916 | 1.175504 | OLIG1 | | 21 |
| 638 | ENSG00000184304 | 138.0745 | -1.13759 | 0.371909 | -3.05877 | 0.002222 | 0.029828 | 8.044947 | 2.540567 | 9.42655 | 5.00087 | PRKD1 | | 14 |
| 639 | ENSG00000184305 | 277.3364 | -2.49055 | 0.248312 | -10.0299 | 1.13E-23 | 2.41E-21 | 20.16308 | 3.607942 | 19.4707 | 3.130658 | CCSER1 | | 4 |
| 640 | ENSG00000184347 | 223.599 | -1.65037 | 0.349697 | -4.71942 | 2.37E-06 | 7.32E-05 | 3.815828 | 1.752628 | 8.874797 | 2.071276 | SLIT3 | | 5 |
| 641 | ENSG00000184588 | 1071.021 | 1.065132 | 0.198715 | 5.360098 | 8.32E-08 | 3.41E-06 | 24.3315 | 64.04613 | 39.46756 | 62.72494 | PDE4B | | 1 |
| 642 | ENSG00000184672 | 143.8836 | 1.550184 | 0.31515 | 4.918883 | 8.70E-07 | 2.93E-05 | 5.236292 | 15.3745 | 5.753459 | 15.2488 | RALYL | | 8 |
| 643 | ENSG00000184903 | 41.12018 | 1.789773 | 0.559544 | 3.198629 | 0.001381 | 0.020133 | 4.12978 | 14.69306 | 4.215131 | 12.85075 | IMMP2L | | 7 |
| 644 | ENSG00000184985 | 799.5996 | -1.9883 | 0.204634 | -9.71637 | 2.57E-22 | 5.09E-20 | 40.95867 | 11.4217 | 36.36174 | 7.28873 | SORCS2 | | 4 |
| 645 | ENSG00000185201 | 35.76711 | 2.223815 | 0.63753 | 3.488175 | 0.000486 | 0.008448 | 2.71409 | 11.66089 | 2.527817 | 11.4685 | IFITM2 | | 11 |
| 646 | ENSG00000185432 | 33.00057 | 1.892039 | 0.640516 | 2.95393 | 0.003138 | 0.039597 | 0.441586 | 1.558634 | 0.503088 | 1.781216 | METTL7A | | 12 |
| 647 | ENSG00000185519 | 40.07543 | 2.164403 | 0.588705 | 3.67655 | 0.000236 | 0.004564 | 1.030042 | 4.748852 | 1.130797 | 4.480863 | FAM131C | | 1 |
| 648 | ENSG00000185666 | 163.1459 | -1.03944 | 0.314198 | -3.30823 | 0.000939 | 0.014631 | 5.379259 | 2.961519 | 5.01323 | 1.909381 | SYN3 | | 22 |
| 649 | ENSG00000185745 | 433.2238 | -2.59177 | 0.236876 | -10.9415 | 7.30E-28 | 2.10E-25 | 16.17637 | 3.255296 | 23.72256 | 3.041936 | IFIT1 | | 10 |
| 650 | ENSG00000185985 | 865.0567 | -1.50399 | 0.161678 | -9.3024 | 1.37E-20 | 2.54E-18 | 18.16857 | 6.087832 | 17.4104 | 5.864164 | SLITRK2 | X | |
| 651 | ENSG00000186369 | 59.17079 | -4.3393 | 0.623815 | -6.95606 | 3.50E-12 | 2.84E-10 | 5.674949 | 0.236024 | 5.023065 | 0.261868 | LINC00643 | | 14 |
| 652 | ENSG00000186409 | 290.6773 | -1.02932 | 0.25286 | -4.07069 | 4.69E-05 | 0.001103 | 15.52059 | 8.054937 | 21.64623 | 9.212227 | CCDC30 | | 1 |
| 653 | ENSG00000186472 | 599.4344 | -1.34628 | 0.186148 | -7.23231 | 4.75E-13 | 4.37E-11 | 7.171112 | 2.959975 | 8.703421 | 2.980071 | PCLO | | 7 |
| 654 | ENSG00000186479 | 70.10824 | -1.5651 | 0.4385 | -3.56922 | 0.000358 | 0.006489 | 2.718772 | 0.982951 | 2.691928 | 0.768825 | RGS7BP | | 5 |
| 655 | ENSG00000186487 | 296.386 | -1.15204 | 0.245769 | -4.68747 | 2.77E-06 | 8.48E-05 | 20.68947 | 9.987171 | 19.0147 | 7.151209 | MYT1L | | 2 |
| 656 | ENSG00000186493 | 63.23 | 2.913015 | 0.550103 | 5.295403 | 1.19E-07 | 4.75E-06 | 4.243752 | 16.24532 | 1.75895 | 22.68688 | C5orf38 | | 5 |
| 657 | ENSG00000186868 | 382.1834 | -1.03765 | 0.219222 | -4.73333 | 2.21E-06 | 6.87E-05 | 16.77459 | 9.31807 | 18.30754 | 7.058302 | MAPT | | 17 |
| 658 | ENSG00000186960 | 134.7596 | -4.74256 | 0.469284 | -10.106 | 5.20E-24 | 1.16E-21 | 20.7651 | 1.178194 | 17.4014 | 0.262316 | LINC01551 | | 14 |
| 659 | ENSG00000186976 | 81.9851 | -1.31055 | 0.403379 | -3.24892 | 0.001158 | 0.017432 | 3.70679 | 1.60265 | 4.343914 | 1.491052 | EFCAB6 | | 22 |
| 660 | ENSG00000187210 | 336.4211 | -1.07006 | 0.255564 | -4.18707 | 2.83E-05 | 0.000695 | 10.77879 | 6.771803 | 17.10074 | 5.874688 | GCNT1 | | 9 |
| 661 | ENSG00000187260 | 80.11262 | -1.41145 | 0.412885 | -3.4185 | 0.00063 | 0.010444 | 9.911119 | 4.029127 | 12.7836 | 4.081277 | WDR86 | | 7 |
| 662 | ENSG00000187391 | 702.8353 | -1.30839 | 0.174684 | -7.49004 | 6.89E-14 | 6.80E-12 | 31.29195 | 11.25249 | 28.87603 | 11.85164 | MAGI2 | | 7 |
| 663 | ENSG00000187498 | 786.9047 | 1.259149 | 0.181398 | 6.941377 | 3.88E-12 | 3.08E-10 | 9.126991 | 22.44638 | 11.64314 | 24.75403 | COL4A1 | | 13 |
| 664 | ENSG00000187553 | 86.9579 | 3.849479 | 0.480845 | 8.00565 | 1.19E-15 | 1.51E-13 | 0.408672 | 5.41801 | 0.413832 | 5.850293 | CYP26C1 | | 10 |
| 665 | ENSG00000187678 | 201.2956 | 1.257996 | 0.365825 | 3.438795 | 0.000584 | 0.00983 | 1.922226 | 6.611124 | 4.88969 | 8.895506 | SPRY4 | | 5 |
| 666 | ENSG00000187720 | 356.1225 | -3.64957 | 0.268904 | -13.5721 | 5.87E-42 | 3.05E-39 | 11.27722 | 1.142769 | 11.40949 | 0.617016 | THSD4 | | 15 |
| 667 | ENSG00000188064 | 1348.627 | -3.38462 | 0.161195 | -20.997 | 6.98E-98 | 2.24E-94 | 68.83909 | 7.25425 | 80.9003 | 6.423253 | WNT7B | | 22 |
| 668 | ENSG00000188191 | 98.39064 | -2.27479 | 0.422835 | -5.37984 | 7.46E-08 | 3.11E-06 | 8.77424 | 2.619054 | 15.71155 | 2.177168 | PRKAR1B | | 7 |
| 669 | ENSG00000188517 | 65.64255 | -2.56451 | 0.512039 | -5.00843 | 5.49E-07 | 1.92E-05 | 2.026135 | 0.350198 | 3.319229 | 0.508543 | COL25A1 | | 4 |
| 670 | ENSG00000188596 | 181.0987 | -2.07961 | 0.306294 | -6.78958 | 1.12E-11 | 8.55E-10 | 9.834287 | 1.795548 | 9.8028 | 2.615983 | CFAP54 | | 12 |
| 671 | ENSG00000188761 | 21.68626 | 3.830176 | 0.832933 | 4.598418 | 4.26E-06 | 0.000123 | 0.267349 | 3.235071 | 0.151673 | 2.534387 | BCL2L15 | | 1 |
| 672 | ENSG00000188803 | 110.9043 | -1.14763 | 0.361868 | -3.1714 | 0.001517 | 0.021756 | 3.081293 | 1.496184 | 2.742681 | 1.02924 | SHISA6 | | 17 |
| 673 | ENSG00000188859 | 43.28225 | -1.96297 | 0.562008 | -3.49278 | 0.000478 | 0.008336 | 2.484034 | 0.534175 | 2.046497 | 0.569762 | FAM78B | | 1 |
| 674 | ENSG00000189056 | 928.4992 | -2.54857 | 0.578468 | -4.40572 | 1.05E-05 | 0.000282 | 24.93052 | 5.194469 | 20.09448 | 2.213748 | RELN | | 7 |
| 675 | ENSG00000189108 | 42.00541 | -8.37957 | 1.540488 | -5.43955 | 5.34E-08 | 2.30E-06 | 5.313449 | 0 | 3.849476 | 0 | IL1RAPL2 | X | |
| 676 | ENSG00000189127 | 62.83908 | -2.31772 | 0.477197 | -4.85695 | 1.19E-06 | 3.91E-05 | 3.214854 | 0.692006 | 3.313196 | 0.562154 | ANKRD34B | | 5 |
| 677 | ENSG00000189184 | 2635.109 | 1.450661 | 0.144396 | 10.04642 | 9.53E-24 | 2.06E-21 | 29.8673 | 86.1106 | 37.78929 | 89.64618 | PCDH18 | | 4 |
| 678 | ENSG00000189337 | 570.1326 | 1.202951 | 0.181375 | 6.632402 | 3.30E-11 | 2.32E-09 | 8.925741 | 19.92466 | 9.138312 | 19.67064 | KAZN | | 1 |
| 679 | ENSG00000189410 | 43.61974 | 1.996884 | 0.563075 | 3.546392 | 0.000391 | 0.007005 | 0.663028 | 2.323206 | 0.59113 | 2.44083 | SH2D5 | | 1 |
| 680 | ENSG00000196083 | 221.4993 | 1.838128 | 0.301415 | 6.098325 | 1.07E-09 | 6.14E-08 | 3.454233 | 8.498063 | 2.803809 | 12.63745 | IL1RAP | | 3 |
| 681 | ENSG00000196091 | 277.5752 | 2.734328 | 0.740857 | 3.690762 | 0.000224 | 0.004359 | 1.740284 | 20.56796 | 5.471535 | 24.83562 | MYBPC1 | | 12 |
| 682 | ENSG00000196263 | 51.64685 | -8.16579 | 1.456597 | -5.60607 | 2.07E-08 | 9.60E-07 | 3.846038 | 0.019074 | 3.016782 | 0 | ZNF471 | | 19 |
| 683 | ENSG00000196268 | 391.5624 | -1.17029 | 0.213359 | -5.48505 | 4.13E-08 | 1.82E-06 | 25.06541 | 11.55009 | 30.32916 | 11.83247 | ZNF493 | | 19 |
| 684 | ENSG00000196376 | 1268.778 | 1.05977 | 0.1477 | 7.175144 | 7.22E-13 | 6.44E-11 | 20.20161 | 38.75758 | 19.74764 | 40.49128 | SLC35F1 | | 6 |
| 685 | ENSG00000196458 | 425.2469 | -1.21628 | 0.219637 | -5.53767 | 3.07E-08 | 1.39E-06 | 16.71312 | 7.173274 | 13.98944 | 5.493077 | ZNF605 | | 12 |
| 686 | ENSG00000196459 | 340.0434 | -1.17716 | 0.225515 | -5.2199 | 1.79E-07 | 6.81E-06 | 26.34215 | 10.03498 | 22.54263 | 10.52873 | TRAPPC2 | X | |
| 687 | ENSG00000196517 | 95.91124 | 1.160753 | 0.384158 | 3.021548 | 0.002515 | 0.033016 | 2.500421 | 4.909375 | 2.811042 | 6.341536 | SLC6A9 | | 1 |
| 688 | ENSG00000196569 | 92.23233 | -1.46934 | 0.410208 | -3.58195 | 0.000341 | 0.006245 | 1.298225 | 0.669614 | 1.822188 | 0.413909 | LAMA2 | | 6 |
| 689 | ENSG00000196604 | 81.04655 | -2.07363 | 0.455387 | -4.55356 | 5.27E-06 | 0.00015 | 4.172974 | 0.561296 | 3.500143 | 1.143922 | POTEF | | 2 |
| 690 | ENSG00000196739 | 402.8027 | 1.118598 | 0.231586 | 4.830166 | 1.36E-06 | 4.41E-05 | 4.937422 | 12.72138 | 7.465543 | 12.77684 | COL27A1 | | 9 |
| 691 | ENSG00000196781 | 873.0888 | -1.58079 | 0.173724 | -9.09942 | 9.08E-20 | 1.59E-17 | 57.64091 | 20.34678 | 55.08004 | 15.70591 | TLE1 | | 9 |
| 692 | ENSG00000196782 | 586.8678 | -1.58329 | 0.201994 | -7.83832 | 4.57E-15 | 5.33E-13 | 16.1563 | 5.913023 | 15.32502 | 4.171086 | MAML3 | | 4 |
| 693 | ENSG00000196867 | 163.2618 | -2.6685 | 0.330857 | -8.06541 | 7.30E-16 | 9.37E-14 | 9.02287 | 1.218781 | 10.8126 | 1.749107 | ZFP28 | | 19 |
| 694 | ENSG00000197013 | 190.4641 | -1.36879 | 0.28451 | -4.81105 | 1.50E-06 | 4.83E-05 | 11.7635 | 3.956926 | 12.3932 | 4.936532 | ZNF429 | | 19 |
| 695 | ENSG00000197177 | 56.01172 | -3.12689 | 0.538933 | -5.80201 | 6.55E-09 | 3.28E-07 | 2.595422 | 0.371469 | 3.065889 | 0.254138 | ADGRA1 | | 10 |
| 696 | ENSG00000197406 | 930.0601 | 3.196064 | 0.542997 | 5.885966 | 3.96E-09 | 2.03E-07 | 7.35969 | 73.89618 | 14.18349 | 112.6577 | DIO3 | | 14 |
| 697 | ENSG00000197415 | 447.5463 | -1.09332 | 0.219942 | -4.97096 | 6.66E-07 | 2.30E-05 | 49.4667 | 21.2509 | 37.60892 | 17.77918 | VEPH1 | | 3 |
| 698 | ENSG00000197496 | 113.6074 | 1.33367 | 0.360315 | 3.701404 | 0.000214 | 0.004211 | 2.693689 | 5.120802 | 2.230202 | 6.631137 | SLC2A10 | | 20 |
| 699 | ENSG00000197550 | 19.57142 | -3.6255 | 0.993944 | -3.64758 | 0.000265 | 0.005029 | 1.81057 | 0.151602 | 0.769603 | 0.056271 | | | 9 |
| 700 | ENSG00000197557 | 203.6797 | -1.00807 | 0.272981 | -3.69282 | 0.000222 | 0.004329 | 5.834152 | 2.652313 | 4.904103 | 2.442376 | TTC30A | | 2 |
| 701 | ENSG00000197565 | 3449.672 | -1.53478 | 0.126726 | -12.111 | 9.23E-34 | 3.42E-31 | 106.1915 | 38.79214 | 119.214 | 35.33019 | COL4A6 | X | |

| | | | | | | | | | | | | | | |
|---|---|---|---|---|---|---|---|---|---|---|---|---|---|---|
| 702 | ENSG00000197584 | 21.20034 | -2.5377 | 0.831299 | -3.05269 | 0.002268 | 0.030313 | 2.978855 | 0.299832 | 1.547502 | 0.434475 | KCNMB2 | | 3 |
| 703 | ENSG00000197705 | 139.0906 | -1.22454 | 0.320913 | -3.81582 | 0.000136 | 0.002836 | 5.010713 | 2.172391 | 6.104172 | 2.341915 | KLHL14 | | 18 |
| 704 | ENSG00000197712 | 111.4461 | 1.261146 | 0.355463 | 3.547896 | 0.000388 | 0.006985 | 3.455392 | 7.897349 | 3.997079 | 9.125482 | FAM114A1 | | 4 |
| 705 | ENSG00000197928 | 214.3189 | -1.71357 | 0.273643 | -6.26205 | 3.80E-10 | 2.32E-08 | 24.4699 | 7.861408 | 23.07564 | 6.060094 | ZNF677 | | 19 |
| 706 | ENSG00000198046 | 36.26378 | -7.14051 | 1.503469 | -4.74935 | 2.04E-06 | 6.38E-05 | 2.341595 | 0 | 2.548349 | 0.028229 | ZNF667 | | 19 |
| 707 | ENSG00000198049 | 9.892528 | -6.7295 | 1.8418 | -3.65376 | 0.000258 | 0.004929 | 1.210462 | 0 | 0.417866 | 0 | AVPR1B | | 1 |
| 708 | ENSG00000198189 | 72.16239 | 1.570255 | 0.466211 | 3.368122 | 0.000757 | 0.012189 | 2.095093 | 9.144844 | 4.335896 | 9.047976 | HSD17B11 | | 4 |
| 709 | ENSG00000198205 | 22.33772 | -7.99004 | 1.61611 | -4.944 | 7.65E-07 | 2.61E-05 | 1.121858 | 0 | 0.904809 | 0 | ZXDA | X | |
| 710 | ENSG00000198300 | 2532.351 | -10.4196 | 0.685271 | -15.2051 | 3.27E-52 | 3.15E-49 | 75.18594 | 0.114498 | 110.908 | 0.037761 | PEG3 | | 19 |
| 711 | ENSG00000198429 | 553.3128 | -1.17361 | 0.192051 | -6.1109 | 9.91E-10 | 5.73E-08 | 48.50219 | 21.46026 | 43.88342 | 17.68519 | ZNF69 | | 19 |
| 712 | ENSG00000198453 | 142.2506 | -1.11092 | 0.338849 | -3.2785 | 0.001044 | 0.016017 | 7.039634 | 3.019886 | 9.626195 | 4.264013 | ZNF568 | | 19 |
| 713 | ENSG00000198467 | 238.1797 | 2.277656 | 0.347093 | 6.562095 | 5.31E-11 | 3.67E-09 | 5.393398 | 44.8795 | 14.58794 | 46.91797 | TPM2 | | 9 |
| 714 | ENSG00000198682 | 360.4185 | -1.28252 | 0.257909 | -4.97278 | 6.60E-07 | 2.28E-05 | 11.67911 | 6.654308 | 19.05693 | 5.39067 | PAPSS2 | | 10 |
| 715 | ENSG00000198718 | 418.8851 | -1.17055 | 0.213356 | -5.48637 | 4.10E-08 | 1.81E-06 | 18.81249 | 8.421803 | 16.65816 | 6.673131 | TOGARAM | | 14 |
| 716 | ENSG00000198729 | 338.6379 | 3.203995 | 0.248606 | 12.88785 | 5.27E-38 | 2.20E-35 | 3.490925 | 30.30819 | 3.82784 | 33.80233 | PPP1R14C | | 6 |
| 717 | ENSG00000198732 | 1616.409 | 1.015133 | 0.178301 | 5.693378 | 1.25E-08 | 5.91E-07 | 34.11015 | 89.15794 | 49.46474 | 71.89779 | SMOC1 | | 14 |
| 718 | ENSG00000198739 | 98.53343 | -1.86639 | 0.44286 | -4.2144 | 2.50E-05 | 0.000626 | 4.727139 | 0.850838 | 2.141467 | 0.92251 | LRRTM3 | | 10 |
| 719 | ENSG00000198797 | 124.0806 | -2.44883 | 0.366221 | -6.68675 | 2.28E-11 | 1.66E-09 | 7.764352 | 1.100606 | 7.918497 | 1.654089 | BRINP2 | | 1 |
| 720 | ENSG00000198825 | 877.6494 | -1.74984 | 0.17199 | -10.1741 | 2.59E-24 | 6.00E-22 | 38.36471 | 9.710068 | 36.34108 | 11.38096 | INPP5F | | 10 |
| 721 | ENSG00000198846 | 835.2861 | -1.48722 | 0.183162 | -8.11969 | 4.67E-16 | 6.08E-14 | 38.9288 | 12.71012 | 30.83201 | 11.06694 | TOX | | 8 |
| 722 | ENSG00000198873 | 514.4423 | 1.077152 | 0.249089 | 4.324361 | 1.53E-05 | 0.000402 | 3.873628 | 10.60692 | 7.177348 | 11.50955 | GRK5 | | 10 |
| 723 | ENSG00000198879 | 336.0671 | 1.586349 | 0.260969 | 6.078692 | 1.21E-09 | 6.84E-08 | 3.487541 | 10.04291 | 4.931077 | 13.89511 | SFMBT2 | | 10 |
| 724 | ENSG00000198963 | 216.4058 | -2.00963 | 0.296581 | -6.77598 | 1.24E-11 | 9.29E-10 | 3.711239 | 1.02101 | 5.614438 | 1.176635 | RORB | | 9 |
| 725 | ENSG00000203335 | 39.20598 | -2.39912 | 0.610612 | -3.92904 | 8.53E-05 | 0.001885 | 6.722164 | 1.02014 | 4.739224 | 1.050083 | | | 7 |
| 726 | ENSG00000203952 | 250.8584 | 1.307639 | 0.278225 | 4.699934 | 2.60E-06 | 8.00E-05 | 9.974496 | 23.1003 | 13.37555 | 31.6416 | CCDC160 | X | |
| 727 | ENSG00000204128 | 105.5176 | 1.522309 | 0.401018 | 3.796109 | 0.000147 | 0.003042 | 1.652605 | 5.277356 | 2.924302 | 7.008169 | C2orf72 | | 2 |
| 728 | ENSG00000204131 | 727.6068 | 1.140441 | 0.197222 | 5.782525 | 7.36E-09 | 3.64E-07 | 4.177934 | 10.06955 | 5.959493 | 11.13251 | NHSL2 | X | |
| 729 | ENSG00000204335 | 21.30646 | 3.685296 | 0.991576 | 3.716605 | 0.000202 | 0.004004 | 0 | 2.548997 | 0.370059 | 2.89859 | SP5 | | 2 |
| 730 | ENSG00000204389 | 218.0569 | -4.4018 | 0.865338 | -5.0868 | 3.64E-07 | 1.30E-05 | 19.99453 | 0.414054 | 26.68084 | 1.627556 | HSPA1A | | 6 |
| 731 | ENSG00000204525 | 463.259 | 1.084727 | 0.213813 | 5.073254 | 3.91E-07 | 1.39E-05 | 22.79652 | 46.93565 | 28.57111 | 56.44647 | HLA-C | | 6 |
| 732 | ENSG00000205213 | 1035.486 | 1.229127 | 0.160486 | 7.658784 | 1.88E-14 | 2.00E-12 | 19.25185 | 48.6694 | 20.41123 | 40.06972 | LGR4 | | 11 |
| 733 | ENSG00000205922 | 60.3955 | -2.0554 | 0.520515 | -3.94878 | 7.85E-05 | 0.00175 | 1.865664 | 0.463348 | 1.167175 | 0.244788 | ONECUT3 | | 19 |
| 734 | ENSG00000206144 | 15.73297 | -3.31821 | 1.018585 | -3.25767 | 0.001123 | 0.01697 | 1.344695 | 0.180272 | 1.918995 | 0.134339 | MAPK6P1 | | 8 |
| 735 | ENSG00000206503 | 218.7681 | -1.17552 | 0.287387 | -4.09038 | 4.31E-05 | 0.001022 | 21.03093 | 9.514447 | 30.14524 | 11.91729 | HLA-A | | 6 |
| 736 | ENSG00000206538 | 963.8356 | 3.973353 | 0.191194 | 20.78177 | 6.33E-96 | 1.74E-92 | 1.529397 | 19.52596 | 1.273438 | 22.14051 | VGLL3 | | 3 |
| 737 | ENSG00000206557 | 870.6645 | 1.185656 | 0.195897 | 6.052442 | 1.43E-09 | 7.89E-08 | 8.052363 | 15.59957 | 5.574402 | 14.00088 | TRIM71 | | 3 |
| 738 | ENSG00000207955 | 118.4923 | 3.207232 | 0.409874 | 7.82493 | 5.08E-15 | 5.85E-13 | 2.693131 | 16.78148 | 1.325409 | 18.45004 | MIR219A2 | | 9 |
| 739 | ENSG00000211448 | 85.8159 | -3.75768 | 0.473872 | -7.92973 | 2.20E-15 | 2.71E-13 | 3.607636 | 0.258656 | 3.979847 | 0.27757 | DIO2 | | 14 |
| 740 | ENSG00000213047 | 427.3561 | -1.78094 | 0.267307 | -6.66252 | 2.69E-11 | 1.93E-09 | 21.16501 | 7.554933 | 19.05203 | 3.798024 | DENND1B | | 1 |
| 741 | ENSG00000213468 | 277.5745 | 3.347399 | 0.270535 | 12.37326 | 3.65E-35 | 1.43E-32 | 3.763289 | 38.54366 | 4.444785 | 41.42224 | FIRRE | X | |
| 742 | ENSG00000214548 | 5621.868 | -6.87627 | 0.197102 | -34.8869 | 1.17E-266 | 2.26E-262 | 210.0524 | 1.787713 | 300.6096 | 2.294501 | MEG3 | | 14 |
| 743 | ENSG00000215196 | 174.7815 | -1.34461 | 0.292305 | -4.60003 | 4.22E-06 | 0.000122 | 14.70373 | 6.371765 | 15.1517 | 4.864659 | BASP1-AS1 | | 5 |
| 744 | ENSG00000215218 | 104.8831 | -1.10764 | 0.368525 | -3.00559 | 0.002651 | 0.03454 | 2.560321 | 1.443162 | 2.946263 | 1.007916 | UBE2QL1 | | 5 |
| 745 | ENSG00000215374 | 19.67783 | -6.3748 | 1.625319 | -3.92218 | 8.78E-05 | 0.00193 | 1.621339 | 0 | 1.617975 | 0.035807 | FAM66B | | 8 |
| 746 | ENSG00000219410 | 51.82006 | 1.638997 | 0.537561 | 3.04895 | 0.002296 | 0.030607 | 2.782536 | 6.759906 | 1.390184 | 5.720513 | | | 12 |
| 747 | ENSG00000221866 | 204.3327 | -2.97919 | 0.375081 | -7.9428 | 1.98E-15 | 2.45E-13 | 4.97499 | 0.665479 | 2.846809 | 0.29468 | PLXNA4 | | 7 |
| 748 | ENSG00000221890 | 231.002 | -1.62281 | 0.259199 | -6.26085 | 3.83E-10 | 2.33E-08 | 6.329128 | 2.076841 | 7.252737 | 2.11547 | NPTXR | | 22 |
| 749 | ENSG00000222041 | 159.7085 | 2.013895 | 0.317358 | 6.345814 | 2.21E-10 | 1.41E-08 | 6.752949 | 28.03587 | 8.909992 | 31.70579 | CYTOR | | 2 |
| 750 | ENSG00000223403 | 109.8283 | -4.95505 | 0.542979 | -9.12567 | 7.13E-20 | 1.26E-17 | 8.478957 | 0.614168 | 15.35148 | 0.184553 | MEG9 | | 14 |
| 751 | ENSG00000223486 | 44.2756 | 1.988744 | 0.560721 | 3.546764 | 0.00039 | 0.007005 | 0.811476 | 3.427255 | 0.772 | 2.583103 | | X | |
| 752 | ENSG00000223561 | 68.26278 | 1.681552 | 0.466168 | 3.607178 | 0.00031 | 0.005756 | 1.289534 | 5.329538 | 2.32633 | 5.700097 | LINC03007 | | 7 |
| 753 | ENSG00000223650 | 86.32996 | -1.16426 | 0.400786 | -2.90493 | 0.003673 | 0.044464 | 5.342939 | 2.948608 | 6.794788 | 2.234751 | UHRF2P1 | X | |
| 754 | ENSG00000223691 | 18.54099 | -3.87953 | 1.012022 | -3.83344 | 0.000126 | 0.00267 | 1.685339 | 0.132524 | 2.693787 | 0.148294 | | | 2 |
| 755 | ENSG00000223839 | 70.56588 | -1.78788 | 0.462583 | -3.86499 | 0.000111 | 0.002376 | 3.481912 | 0.756331 | 2.226128 | 0.817437 | FAM95B1 | | 9 |
| 756 | ENSG00000224165 | 32.68663 | -3.13951 | 0.939922 | -3.34018 | 0.000837 | 0.013219 | 0.510609 | 0.350185 | 7.451716 | 0.36162 | DNAJC27-/ | | 2 |
| 757 | ENSG00000224383 | 19.86626 | -2.48984 | 0.826367 | -3.01299 | 0.002587 | 0.033869 | 2.519803 | 0.437791 | 2.415342 | 0.398236 | PRR29 | | 17 |
| 758 | ENSG00000224597 | 41.89627 | -2.41294 | 0.629416 | -3.83361 | 0.000126 | 0.00267 | 9.504626 | 2.416908 | 8.047491 | 0.856434 | SVIL-AS1 | | 10 |
| 759 | ENSG00000224717 | 61.98682 | -2.09964 | 0.475418 | -4.41641 | 1.00E-05 | 0.00027 | 4.112587 | 1.043976 | 4.164902 | 0.808066 | LEMD1-DT | | 1 |
| 760 | ENSG00000225156 | 64.88401 | -3.98862 | 0.573901 | -6.95002 | 3.65E-12 | 2.94E-10 | 4.258776 | 0.423577 | 6.646448 | 0.244953 | | | 2 |
| 761 | ENSG00000225174 | 19.10341 | -4.29464 | 1.132941 | -3.7907 | 0.00015 | 0.003095 | 2.371689 | 0.148707 | 0.912081 | 0 | OSTM1-AS | | 6 |
| 762 | ENSG00000225206 | 409.7669 | -1.11542 | 0.292974 | -3.80724 | 0.000141 | 0.002918 | 20.40036 | 12.56682 | 20.06959 | 5.430856 | MIR137HG | | 1 |
| 763 | ENSG00000225649 | 23.83883 | 2.794065 | 0.811093 | 3.444815 | 0.000571 | 0.009716 | 0.189336 | 1.708543 | 0.413416 | 2.362894 | | | 2 |
| 764 | ENSG00000225746 | 1641.876 | -5.12088 | 0.222318 | -23.034 | 2.13E-117 | 8.19E-114 | 189.9643 | 5.342477 | 292.0586 | 7.578777 | MEG8 | | 14 |
| 765 | ENSG00000225868 | 10.39497 | -6.28414 | 1.814094 | -3.46406 | 0.000532 | 0.009135 | 0.789565 | 0 | 2.173623 | 0 | WDR87BP | | 19 |
| 766 | ENSG00000225968 | 184.6065 | -1.27143 | 0.280489 | -4.53289 | 5.82E-06 | 0.000164 | 7.423546 | 3.094308 | 8.357333 | 3.122655 | ELFN1 | | 7 |
| 767 | ENSG00000226539 | 57.78545 | 1.477797 | 0.486643 | 3.036715 | 0.002392 | 0.031615 | 5.960158 | 17.47185 | 8.274436 | 20.16704 | MLXP1 | | 2 |
| 768 | ENSG00000226686 | 27.31891 | -4.93708 | 0.913698 | -5.40341 | 6.54E-08 | 2.76E-06 | 2.978292 | 0.085065 | 2.852277 | 0.094797 | LINC01535 | | 19 |
| 769 | ENSG00000227115 | 23.8843 | -2.55163 | 0.768053 | -3.3222 | 0.000893 | 0.01402 | 2.23149 | 0.519509 | 2.323841 | 0.236237 | LINC01630 | | 18 |
| 770 | ENSG00000227124 | 27.68273 | -4.88564 | 0.965274 | -5.0614 | 4.16E-07 | 1.47E-05 | 2.854427 | 0.117843 | 2.45476 | 0.06425 | ZNF717 | | 3 |
| 771 | ENSG00000227487 | 207.5004 | -1.00784 | 0.266205 | -3.78597 | 0.000153 | 0.003138 | 5.767132 | 2.702435 | 5.263472 | 2.528036 | NCAM1-AS | | 11 |
| 772 | ENSG00000228061 | 75.97401 | -2.0298 | 0.43599 | -4.6556 | 3.23E-06 | 9.67E-05 | 5.83483 | 1.856142 | 7.518072 | 1.302551 | | | 11 |
| 773 | ENSG00000228623 | 115.4302 | -1.77495 | 0.359966 | -4.93088 | 8.19E-07 | 2.77E-05 | 9.474962 | 2.190611 | 9.386463 | 3.006698 | ZNF883 | | 9 |
| 774 | ENSG00000228999 | 43.31846 | -3.33976 | 0.628612 | -5.31291 | 1.08E-07 | 4.36E-06 | 13.00986 | 1.451754 | 18.53584 | 1.496479 | | | 2 |
| 775 | ENSG00000229618 | 109.7074 | 4.048577 | 0.489816 | 8.265509 | 1.39E-16 | 2.01E-14 | 0.383877 | 7.030544 | 0.913079 | 11.38394 | | | 7 |
| 776 | ENSG00000229847 | 331.8344 | -2.55505 | 0.779394 | -3.27825 | 0.001045 | 0.016018 | 6.872675 | 2.347249 | 12.49538 | 0.814653 | EMX2OS | | 10 |
| 777 | ENSG00000230001 | 32.60731 | -2.29451 | 0.663313 | -3.45916 | 0.000542 | 0.009262 | 12.06267 | 3.236157 | 18.66692 | 2.741433 | | | 9 |
| 778 | ENSG00000230316 | 668.7416 | -5.10084 | 0.254738 | -20.0239 | 3.41E-89 | 8.21E-86 | 187.3038 | 7.620018 | 215.0174 | 3.741624 | FEZF1-AS1 | | 7 |
| 779 | ENSG00000230910 | 65.45104 | -1.7942 | 0.483593 | -3.71015 | 0.000207 | 0.004101 | 7.469046 | 1.523614 | 4.192617 | 1.666339 | ADGRB3-D | | 6 |

| | | | | | | | | | | | | | |
|---|---|---|---|---|---|---|---|---|---|---|---|---|---|
| 780 | ENSG00000231424 | 15.70586 | -4.89452 | 1.191175 | -4.10899 | 3.97E-05 | 0.00095 | 1.250795 | 0.052764 | 0.891106 | 0.019546 | | 1 |
| 781 | ENSG00000231764 | 383.4691 | -7.49675 | 1.695923 | -4.42045 | 9.85E-06 | 0.000265 | 16.34837 | 0.155219 | 13.14306 | 0 | DLX6-AS1 | 7 |
| 782 | ENSG00000231806 | 63.74387 | -3.54098 | 0.531809 | -6.65836 | 2.77E-11 | 1.97E-09 | 3.4766 | 0.215576 | 3.490299 | 0.342507 | PCAT7 | 9 |
| 783 | ENSG00000233058 | 44.94497 | -3.2978 | 0.634492 | -5.19755 | 2.02E-07 | 7.61E-06 | 2.003637 | 0.278936 | 3.684693 | 0.268422 | ATP13A3-[ | 3 |
| 784 | ENSG00000233237 | 2132.188 | 1.289669 | 0.164285 | 7.850198 | 4.15E-15 | 4.93E-13 | 20.56414 | 59.18118 | 29.50609 | 57.12971 | LINC00472 | 6 |
| 785 | ENSG00000233363 | 40.27669 | -1.90928 | 0.587189 | -3.25156 | 0.001148 | 0.017298 | 2.669191 | 0.779536 | 2.261563 | 0.487579 | | 7 |
| 786 | ENSG00000233382 | 49.68109 | -1.60083 | 0.536227 | -2.98536 | 0.002832 | 0.036414 | 6.105641 | 1.253222 | 4.528195 | 2.046662 | NKAPP1 | X |
| 787 | ENSG00000233429 | 23.8251 | 2.194399 | 0.762155 | 2.879202 | 0.003987 | 0.047528 | 2.369788 | 9.184088 | 1.8336 | 9.148711 | HOTAIRM1 | 7 |
| 788 | ENSG00000233587 | 25.66928 | -3.32969 | 0.809195 | -4.11481 | 3.87E-05 | 0.000931 | 3.439672 | 0.397313 | 5.555894 | 0.447157 | LINC01884 | 2 |
| 789 | ENSG00000233639 | 874.7956 | 1.913602 | 0.163222 | 11.72389 | 9.61E-32 | 3.19E-29 | 35.36231 | 138.4035 | 38.55749 | 126.9391 | PANTR1 | 2 |
| 790 | ENSG00000233729 | 56.94987 | 1.574632 | 0.486696 | 3.235348 | 0.001215 | 0.018098 | 1.833597 | 6.109575 | 2.427216 | 5.972618 | NCKAP5-A | 2 |
| 791 | ENSG00000233967 | 453.8135 | 1.334711 | 0.248666 | 5.367491 | 7.98E-08 | 3.29E-06 | 9.32793 | 23.60652 | 14.25064 | 32.64678 | | 6 |
| 792 | ENSG00000234173 | 14.17555 | -3.5562 | 1.189409 | -2.98989 | 0.002791 | 0.035927 | 0.795648 | 0 | 2.000543 | 0.208059 | | 10 |
| 793 | ENSG00000234444 | 15.95943 | -6.72596 | 1.71971 | -3.9111 | 9.19E-05 | 0.001996 | 0.566031 | 0 | 2.002678 | 0 | ZNF736 | 7 |
| 794 | ENSG00000234745 | 583.0207 | 1.531205 | 0.212045 | 7.221143 | 5.16E-13 | 4.70E-11 | 28.35697 | 67.04809 | 29.78295 | 92.24135 | HLA-B | 6 |
| 795 | ENSG00000234840 | 14.58578 | -5.74964 | 1.629051 | -3.52944 | 0.000416 | 0.007428 | 2.124364 | 0.054631 | 1.570384 | 0 | LINC01239 | 9 |
| 796 | ENSG00000235597 | 36.18474 | 2.09437 | 0.622166 | 3.366254 | 0.000762 | 0.012251 | 0.78352 | 4.159933 | 1.157438 | 3.716371 | LINC01102 | 2 |
| 797 | ENSG00000235770 | 62.81216 | -7.23645 | 1.110403 | -6.51696 | 7.17E-11 | 4.90E-09 | 4.467521 | 0.038627 | 5.156912 | 0.024727 | LINC00607 | 2 |
| 798 | ENSG00000236333 | 71.67401 | 1.876723 | 0.466836 | 4.020087 | 5.82E-05 | 0.001344 | 0.541473 | 2.859543 | 1.066241 | 2.658588 | TRHDE-AS | 12 |
| 799 | ENSG00000236502 | 8.922646 | -6.58571 | 1.851001 | -3.55792 | 0.000374 | 0.006736 | 2.908377 | 0 | 5.325547 | 0 | SIX3-AS1 | 2 |
| 800 | ENSG00000236857 | 8.794101 | -5.16955 | 1.803889 | -2.86578 | 0.00416 | 0.049419 | 6.496513 | 0 | 5.828148 | 0.292767 | RAP1BP1 | 9 |
| 801 | ENSG00000237065 | 19.09513 | -4.16872 | 1.084993 | -3.84217 | 0.000122 | 0.002591 | 6.141536 | 0.687457 | 5.677951 | 0 | NANOGP4 | 7 |
| 802 | ENSG00000237149 | 84.04627 | -2.72102 | 0.426192 | -6.3845 | 1.72E-10 | 1.12E-08 | 12.30089 | 2.107432 | 13.09534 | 1.574999 | ZNF503-AS | 10 |
| 803 | ENSG00000237238 | 26.45373 | -2.44267 | 0.745506 | -3.27653 | 0.001051 | 0.01609 | 4.672598 | 0.51072 | 3.871068 | 0.98508 | BMS1P10 | 9 |
| 804 | ENSG00000237440 | 465.5026 | -1.96192 | 0.258667 | -7.58473 | 3.33E-14 | 3.43E-12 | 22.82024 | 4.270882 | 26.32162 | 7.695667 | ZNF737 | 19 |
| 805 | ENSG00000237499 | 19.82198 | 2.503634 | 0.859414 | 2.913189 | 0.003578 | 0.04374 | 0.652766 | 5.158564 | 0.871494 | 3.219057 | WAKMAR2 | 6 |
| 806 | ENSG00000237807 | 36.66219 | -2.0597 | 0.65603 | -3.13964 | 0.001692 | 0.023938 | 2.169309 | 0.816144 | 2.49535 | 0.285562 | LINC02984 | 8 |
| 807 | ENSG00000237863 | 36.45657 | -1.80308 | 0.600559 | -3.00234 | 0.002679 | 0.034745 | 4.908227 | 1.496234 | 5.572456 | 1.366963 | IRS4-AS1 | X |
| 808 | ENSG00000237877 | 101.6051 | -1.60322 | 0.407595 | -3.93336 | 8.38E-05 | 0.001853 | 19.83042 | 5.342028 | 11.33161 | 4.481582 | LINC01473 | 2 |
| 809 | ENSG00000239519 | 67.42408 | -2.66355 | 0.493362 | -5.39877 | 6.71E-08 | 2.82E-06 | 6.05348 | 1.000234 | 4.42292 | 0.6021 | CADM2-AS | 3 |
| 810 | ENSG00000239922 | 362.6477 | -2.05922 | 0.304149 | -6.77042 | 1.28E-11 | 9.58E-10 | 42.7662 | 11.13903 | 82.09852 | 17.03403 | | 3 |
| 811 | ENSG00000240086 | 81.30031 | 1.445868 | 0.409804 | 3.528191 | 0.000418 | 0.007456 | 2.556267 | 7.799827 | 2.862931 | 6.310909 | | 3 |
| 812 | ENSG00000240225 | 191.8564 | -3.24381 | 0.302379 | -10.7276 | 7.55E-27 | 2.05E-24 | 17.01474 | 1.960639 | 17.87052 | 1.566718 | ZNF542P | 19 |
| 813 | ENSG00000240240 | 50.34122 | -2.25732 | 0.579947 | -3.89228 | 9.93E-05 | 0.00214 | 3.260843 | 0.538547 | 1.480473 | 0.413165 | | 9 |
| 814 | ENSG00000241131 | 27.60512 | -2.71584 | 0.844348 | -3.2165 | 0.001298 | 0.019123 | 1.952023 | 0.888094 | 9.13753 | 0.711667 | LINC02032 | 3 |
| 815 | ENSG00000242288 | 8.077932 | 6.383534 | 1.90814 | 3.345422 | 0.000822 | 0.013014 | 0 | 0.708691 | 0 | 0.2389 | BMS1P4-A | 10 |
| 816 | ENSG00000243244 | 532.4355 | -1.11028 | 0.190492 | -5.82848 | 5.59E-09 | 2.82E-07 | 16.56416 | 6.803504 | 16.1948 | 7.60716 | STON1 | 2 |
| 817 | ENSG00000243915 | 49.31753 | -2.50935 | 0.579748 | -4.32834 | 1.50E-05 | 0.000396 | 5.256264 | 1.079054 | 3.736795 | 0.465273 | THAP12P2 | 3 |
| 818 | ENSG00000244405 | 603.6792 | -1.12593 | 0.211837 | -5.31507 | 1.07E-07 | 4.32E-06 | 27.56348 | 14.12446 | 40.58011 | 15.47148 | ETV5 | 3 |
| 819 | ENSG00000245532 | 4503.979 | 1.788374 | 0.387395 | 4.61641 | 3.90E-06 | 0.000114 | 11.62994 | 40.83227 | 17.88615 | 55.70444 | NEAT1 | 11 |
| 820 | ENSG00000245680 | 217.4007 | -1.16246 | 0.280056 | -4.15082 | 3.31E-05 | 0.000807 | 8.345046 | 4.516855 | 11.91302 | 4.086302 | ZNF585B | 19 |
| 821 | ENSG00000245694 | 1859.638 | 2.071621 | 0.135722 | 15.26375 | 1.33E-52 | 1.35E-49 | 28.38977 | 115.9518 | 28.28547 | 110.9913 | CRNDE | 16 |
| 822 | ENSG00000246214 | 40.85208 | -2.37972 | 0.588238 | -4.0455 | 5.22E-05 | 0.001218 | 4.616776 | 0.766003 | 4.449243 | 0.883649 | RETREG1-A | 5 |
| 823 | ENSG00000248445 | 127.106 | -1.32498 | 0.332569 | -3.98406 | 6.77E-05 | 0.001543 | 55.26447 | 22.82836 | 53.10766 | 18.52152 | SEMA6A-A | 5 |
| 824 | ENSG00000248491 | 44.35696 | 2.623784 | 0.584892 | 4.485932 | 7.26E-06 | 0.000201 | 1.24981 | 6.827292 | 1.37819 | 8.545092 | | 4 |
| 825 | ENSG00000248905 | 55.55775 | 1.677473 | 0.489206 | 3.428968 | 0.000606 | 0.010114 | 0.476612 | 1.373031 | 0.417379 | 1.339364 | FMN1 | 15 |
| 826 | ENSG00000249860 | 19.66972 | -2.90261 | 0.886308 | -3.27495 | 0.001057 | 0.016145 | 2.107346 | 0.238894 | 3.161771 | 0.416801 | NA | NA |
| 827 | ENSG00000250033 | 47.57932 | 3.464706 | 0.570251 | 6.075751 | 1.23E-09 | 6.92E-08 | 0.344035 | 3.256144 | 0.327278 | 3.75058 | SLC7A11-A | 4 |
| 828 | ENSG00000250062 | 35.21249 | 1.935214 | 0.627104 | 3.085954 | 0.002029 | 0.027636 | 3.379752 | 9.69444 | 2.788124 | 12.66549 | MAPK10-A | 4 |
| 829 | ENSG00000250208 | 463.2205 | 3.408757 | 0.571741 | 5.962064 | 2.49E-09 | 1.33E-07 | 1.218717 | 21.06604 | 2.950228 | 20.51548 | FZD10-AS1 | 12 |
| 830 | ENSG00000250337 | 358.2985 | -1.65139 | 0.217546 | -7.591 | 3.17E-14 | 3.30E-12 | 47.03744 | 15.10374 | 52.39396 | 15.00591 | PURPL | 5 |
| 831 | ENSG00000250420 | 22.08716 | -3.61593 | 0.891303 | -4.0569 | 4.97E-05 | 0.001163 | 1.676247 | 0.178249 | 1.811967 | 0.099357 | AACSP1 | 5 |
| 832 | ENSG00000250786 | 65.2541 | 2.475971 | 0.483081 | 5.125377 | 2.97E-07 | 1.08E-05 | 1.730347 | 9.795877 | 2.489887 | 12.52188 | SNHG18 | 5 |
| 833 | ENSG00000251632 | 29.96341 | 5.821498 | 1.246889 | 4.668817 | 3.03E-06 | 9.19E-05 | 0.057847 | 1.365041 | 0.055035 | 4.617353 | LINC02172 | 4 |
| 834 | ENSG00000253284 | 705.005 | -1.48709 | 0.180011 | -8.26109 | 1.44E-16 | 2.07E-14 | 19.67878 | 7.543107 | 24.26281 | 7.369022 | | 12 |
| 835 | ENSG00000253357 | 34.8355 | 1.943138 | 0.63249 | 3.072206 | 0.002125 | 0.028758 | 5.125996 | 25.86001 | 8.813922 | 25.17605 | | 5 |
| 836 | ENSG00000253661 | 491.6017 | -1.19349 | 0.204362 | -5.84006 | 5.22E-09 | 2.64E-07 | 32.70044 | 13.27487 | 26.80351 | 11.57059 | ZFHX4-AS1 | 8 |
| 837 | ENSG00000253706 | 38.32374 | -2.1813 | 0.636109 | -3.42914 | 0.000606 | 0.010114 | 3.842522 | 0.933739 | 2.453487 | 0.432575 | | 8 |
| 838 | ENSG00000253767 | 84.3924 | -1.48514 | 0.416021 | -3.56987 | 0.000357 | 0.006479 | 2.134949 | 0.962521 | 2.371086 | 0.590204 | PCDHGA8 | 5 |
| 839 | ENSG00000253925 | 35.0665 | 2.194959 | 0.631882 | 3.473687 | 0.000513 | 0.00884 | 0.723981 | 3.440258 | 0.688787 | 2.740125 | | 5 |
| 840 | ENSG00000253953 | 182.7898 | -3.92165 | 0.342153 | -11.4617 | 2.05E-30 | 6.38E-28 | 7.604172 | 0.484538 | 8.958749 | 0.548085 | PCDHGB4 | 5 |
| 841 | ENSG00000254186 | 61.66787 | 2.973227 | 0.585614 | 5.077111 | 3.83E-07 | 1.36E-05 | 0.856442 | 3.274601 | 0.79708 | 8.925489 | | 5 |
| 842 | ENSG00000254187 | 76.58656 | 1.594454 | 0.420666 | 3.790308 | 0.00015 | 0.003097 | 4.794421 | 13.45393 | 4.689057 | 13.79599 | | 5 |
| 843 | ENSG00000254245 | 73.08497 | -4.2006 | 0.542948 | -7.73665 | 1.02E-14 | 1.12E-12 | 4.055515 | 0.251791 | 4.906636 | 0.209096 | PCDHGA3 | 5 |
| 844 | ENSG00000254579 | 36.1166 | 1.998989 | 0.657276 | 3.041321 | 0.002355 | 0.031242 | 0.293962 | 1.454094 | 0.69715 | 2.26015 | | 11 |
| 845 | ENSG00000254656 | 1830.001 | 2.289689 | 0.584078 | 3.920176 | 8.85E-05 | 0.001942 | 10.8222 | 39.11293 | 15.44248 | 82.05421 | RTL1 | 14 |
| 846 | ENSG00000254685 | 185.349 | -1.17352 | 0.314233 | -3.73455 | 0.000188 | 0.003774 | 10.17347 | 3.260935 | 6.450492 | 3.737137 | FPGT | 1 |
| 847 | ENSG00000255043 | 79.49106 | 1.422018 | 0.410858 | 3.461092 | 0.000538 | 0.00922 | 1.639147 | 4.05049 | 1.486836 | 3.92823 | NAV2-AS5 | 11 |
| 848 | ENSG00000255571 | 294.719 | -1.61149 | 0.27211 | -5.9222 | 3.18E-09 | 1.67E-07 | 15.92484 | 4.709212 | 10.62498 | 3.626117 | MIR9-3HG | 15 |
| 849 | ENSG00000256463 | 502.504 | -4.25392 | 0.719806 | -5.90981 | 3.42E-09 | 1.78E-07 | 34.79095 | 1.881117 | 18.01225 | 0.795087 | SALL3 | 18 |
| 850 | ENSG00000256637 | 10.51946 | -5.45255 | 1.779542 | -3.06402 | 0.002184 | 0.029453 | 3.691561 | 0.101344 | 1.30131 | 0 | LINC01965 | 2 |
| 851 | ENSG00000257056 | 39.70465 | -4.38291 | 0.790675 | -5.54325 | 2.97E-08 | 1.35E-06 | 6.007698 | 0.439681 | 4.402021 | 0.060913 | LINC02282 | 14 |
| 852 | ENSG00000257126 | 23.37712 | -3.81807 | 1.021238 | -3.73867 | 0.000185 | 0.00372 | 2.930413 | 0 | 2.922851 | 0.409378 | FOXG1-AS | 14 |
| 853 | ENSG00000257354 | 1074.268 | 1.133747 | 0.157671 | 7.190586 | 6.45E-13 | 5.80E-11 | 8.849808 | 17.2512 | 8.444729 | 18.84295 | MIRLET7IH | 12 |
| 854 | ENSG00000257522 | 8.456225 | -6.30969 | 1.858097 | -3.39578 | 0.000684 | 0.01119 | 0.805226 | 0 | 0.89746 | 0 | | 14 |
| 855 | ENSG00000257545 | 177.5423 | 1.867065 | 0.292088 | 6.39214 | 1.64E-10 | 1.07E-08 | 5.715703 | 19.35325 | 5.291145 | 18.88524 | | 12 |
| 856 | ENSG00000257711 | 424.5217 | 1.637069 | 0.225071 | 7.273554 | 3.50E-13 | 3.29E-11 | 6.975172 | 19.34705 | 5.124991 | 16.56069 | | 12 |
| 857 | ENSG00000257800 | 29.12356 | -4.32333 | 0.873545 | -4.94918 | 7.45E-07 | 2.55E-05 | 3.740552 | 0.065656 | 3.177486 | 0.24487 | FNBP1P1 | 2 |

| | | | | | | | | | | | | | |
|---|---|---|---|---|---|---|---|---|---|---|---|---|---|
| 858 | ENSG00000257918 | 64.44464 | 1.702682 | 0.461704 | 3.687825 | 0.000226 | 0.004393 | 6.24098 | 20.15025 | 5.544144 | 16.48744 | | 12 |
| 859 | ENSG00000257935 | 61.48598 | -3.67141 | 0.605253 | -6.0659 | 1.31E-09 | 7.32E-08 | 54.37407 | 5.782062 | 36.46518 | 1.309324 | LHX5-AS1 | 12 |
| 860 | ENSG00000257986 | 23.47855 | 4.655772 | 1.080168 | 4.310231 | 1.63E-05 | 0.000426 | 0.289866 | 5.608758 | 0.137922 | 4.63184 | LINC02306 | 14 |
| 861 | ENSG00000258232 | 73.83 | 2.112641 | 0.498548 | 4.237586 | 2.26E-05 | 0.000572 | 4.322991 | 22.1618 | 9.775119 | 35.45782 | | 12 |
| 862 | ENSG00000258377 | 22.72943 | 3.26637 | 0.956819 | 3.413781 | 0.000641 | 0.010611 | 0.420051 | 1.91048 | 0 | 1.921096 | | 14 |
| 863 | ENSG00000258399 | 25.62711 | -8.18716 | 1.593443 | -5.13803 | 2.78E-07 | 1.02E-05 | 3.222806 | 0 | 3.193959 | 0 | MIR493HG | 14 |
| 864 | ENSG00000258498 | 159.3262 | 1.560108 | 0.342113 | 4.560212 | 5.11E-06 | 0.000146 | 3.652857 | 10.91796 | 5.771037 | 15.34661 | DIO3OS | 14 |
| 865 | ENSG00000258548 | 39.87598 | -2.23173 | 0.634014 | -3.52 | 0.000432 | 0.007634 | 3.93661 | 1.196609 | 3.956278 | 0.437447 | LINC00645 | 14 |
| 866 | ENSG00000258636 | 30.76393 | 3.458845 | 0.722181 | 4.789442 | 1.67E-06 | 5.32E-05 | 0.174216 | 1.758569 | 0.142054 | 1.556399 | LRFN5-DT | 14 |
| 867 | ENSG00000259417 | 66.67056 | 2.258166 | 0.472439 | 4.779803 | 1.75E-06 | 5.56E-05 | 0.368475 | 1.986621 | 0.383902 | 1.456756 | CTXND1 | 15 |
| 868 | ENSG00000259439 | 149.4866 | -3.29727 | 0.402906 | -8.18372 | 2.75E-16 | 3.76E-14 | 21.81528 | 3.565855 | 23.7074 | 1.269885 | LINC01833 | 2 |
| 869 | ENSG00000259498 | 52.89567 | 1.734859 | 0.534352 | 3.246661 | 0.001168 | 0.017516 | 1.065297 | 3.528283 | 0.710223 | 2.13845 | TPM1-AS | 15 |
| 870 | ENSG00000259867 | 91.39243 | -2.57966 | 0.422535 | -6.10519 | 1.03E-09 | 5.92E-08 | 7.572115 | 1.790325 | 11.79925 | 1.328081 | DYNLRB2- | 16 |
| 871 | ENSG00000260578 | 34.28314 | 2.496036 | 0.686432 | 3.636246 | 0.000277 | 0.005215 | 0.199128 | 1.84149 | 0.568309 | 2.269185 | | 18 |
| 872 | ENSG00000260645 | 156.1869 | 1.474291 | 0.354458 | 4.159282 | 3.19E-05 | 0.000779 | 3.794114 | 9.939243 | 5.976709 | 15.70973 | | 6 |
| 873 | ENSG00000261087 | 86.82565 | -1.14536 | 0.397758 | -2.87954 | 0.003983 | 0.047519 | 4.977633 | 1.869442 | 5.1229 | 2.447923 | ZNNT1 | 8 |
| 874 | ENSG00000261136 | 138.4882 | 1.194126 | 0.323564 | 3.690542 | 0.000224 | 0.004359 | 3.374573 | 6.516504 | 3.170721 | 7.70153 | | 15 |
| 875 | ENSG00000261586 | 74.63635 | 1.580155 | 0.461383 | 3.424823 | 0.000615 | 0.010225 | 2.842921 | 5.465268 | 1.562741 | 7.037169 | | 12 |
| 876 | ENSG00000261659 | 52.54348 | -1.45113 | 0.499051 | -2.90777 | 0.00364 | 0.044117 | 2.73944 | 1.159992 | 3.607547 | 1.049189 | | 16 |
| 877 | ENSG00000261771 | 34.63531 | -5.09162 | 0.941825 | -5.40612 | 6.44E-08 | 2.72E-06 | 1.396747 | 0.105747 | 1.967526 | 0 | DNAAF4-C | 15 |
| 878 | ENSG00000262655 | 84.42021 | -2.28411 | 0.430745 | -5.30271 | 1.14E-07 | 4.58E-06 | 3.325187 | 0.549485 | 3.830162 | 0.84851 | SPON1 | 11 |
| 879 | ENSG00000263146 | 31.47826 | -5.10114 | 0.97501 | -5.23188 | 1.68E-07 | 6.47E-06 | 14.83775 | 1.229639 | 12.38499 | 0.085424 | LINC01896 | 18 |
| 880 | ENSG00000263424 | 36.33314 | 8.34439 | 1.55015 | 5.382956 | 7.33E-08 | 3.07E-06 | 0 | 2.942651 | 0 | 2.800521 | | 18 |
| 881 | ENSG00000263677 | 16.29751 | 2.882849 | 0.996278 | 2.893618 | 0.003808 | 0.045844 | 0.129194 | 2.121961 | 0.393045 | 1.900909 | | 18 |
| 882 | ENSG00000263711 | 35.70858 | -8.54755 | 1.555159 | -5.49626 | 3.88E-08 | 1.72E-06 | 4.565592 | 0 | 6.078502 | 0 | LINC02864 | 18 |
| 883 | ENSG00000264596 | 44.85652 | -1.82752 | 0.564186 | -3.23922 | 0.001199 | 0.017868 | 6.750366 | 1.343009 | 4.390193 | 1.633067 | | 18 |
| 884 | ENSG00000265413 | 32.0542 | -2.04854 | 0.658488 | -3.11097 | 0.001865 | 0.025801 | 3.159488 | 0.721182 | 2.254399 | 0.536997 | | 18 |
| 885 | ENSG00000266709 | 89.19795 | 1.259834 | 0.406053 | 3.10263 | 0.001918 | 0.026425 | 3.322925 | 10.77276 | 4.821443 | 7.87479 | | 17 |
| 886 | ENSG00000267254 | 122.7126 | -1.44056 | 0.338181 | -4.25974 | 2.05E-05 | 0.000522 | 10.90556 | 3.61996 | 9.599801 | 3.572701 | ZNF790-AS | 19 |
| 887 | ENSG00000267313 | 83.62851 | 1.319364 | 0.416876 | 3.164884 | 0.001551 | 0.0222 | 4.117986 | 13.4231 | 6.386777 | 11.59762 | | 18 |
| 888 | ENSG00000267586 | 42.93573 | 2.000526 | 0.581257 | 3.441724 | 0.000578 | 0.009797 | 1.349202 | 4.953471 | 1.722981 | 6.727411 | LINC00907 | 18 |
| 889 | ENSG00000267605 | 36.93952 | -2.36141 | 0.638996 | -3.6955 | 0.000219 | 0.004297 | 4.637743 | 0.502191 | 3.045112 | 0.899947 | | 19 |
| 890 | ENSG00000267640 | 42.96019 | -8.04357 | 1.533465 | -5.24535 | 1.56E-07 | 6.04E-06 | 3.811033 | 0 | 4.163104 | 0 | | 19 |
| 891 | ENSG00000268119 | 112.3973 | -7.56341 | 0.903806 | -8.36841 | 5.84E-17 | 8.65E-15 | 10.96362 | 0.085948 | 12.70721 | 0.050372 | | 19 |
| 892 | ENSG00000268654 | 11.75438 | -7.01493 | 1.759374 | -3.98717 | 6.69E-05 | 0.001525 | 3.233268 | 0 | 2.126276 | 0 | MIMT1 | 19 |
| 893 | ENSG00000268658 | 29.73914 | -6.98134 | 1.539955 | -4.53347 | 5.80E-06 | 0.000164 | 2.421485 | 0.041249 | 2.472307 | 0 | LINC00664 | 19 |
| 894 | ENSG00000269699 | 42.14386 | -9.14483 | 1.554119 | -5.88425 | 4.00E-09 | 2.04E-07 | 4.592238 | 0 | 9.531423 | 0 | ZIM2 | 19 |
| 895 | ENSG00000269834 | 71.79794 | -2.39814 | 0.448247 | -5.35005 | 8.79E-08 | 3.60E-06 | 6.044759 | 1.104427 | 5.877988 | 1.051221 | ZNF528-AS | 19 |
| 896 | ENSG00000270276 | 67.46057 | -1.49875 | 0.475464 | -3.15218 | 0.001621 | 0.023041 | 6.35073 | 2.298154 | 10.35675 | 3.298241 | H4C15 | 1 |
| 897 | ENSG00000270953 | 82.93278 | -1.68164 | 0.442913 | -3.79677 | 0.000147 | 0.003037 | 45.4343 | 13.3541 | 28.15591 | 8.734503 | | 7 |
| 898 | ENSG00000271848 | 11.87955 | -4.9974 | 1.485676 | -3.36372 | 0.000769 | 0.012328 | 2.26336 | 0 | 1.114853 | 0.073446 | SYNPO2L- | 10 |
| 899 | ENSG00000272078 | 47.82869 | 1.980846 | 0.541501 | 3.658069 | 0.000254 | 0.004867 | 0.243064 | 1.166694 | 0.303473 | 0.895858 | | 1 |
| 900 | ENSG00000272129 | 130.8706 | 1.088422 | 0.34081 | 3.193636 | 0.001405 | 0.020407 | 6.872939 | 13.40097 | 8.501423 | 17.56883 | | 6 |
| 901 | ENSG00000272573 | 29.27249 | 2.402999 | 0.717743 | 3.347993 | 0.000814 | 0.012926 | 4.655977 | 22.63466 | 2.666018 | 14.34822 | MUSTN1 | 3 |
| 902 | ENSG00000272636 | 86.11483 | -1.72487 | 0.434015 | -3.97421 | 7.06E-05 | 0.001593 | 3.013254 | 0.592281 | 3.033319 | 1.136002 | DOC2B | 17 |
| 903 | ENSG00000272865 | 19.7247 | -3.86439 | 1.097391 | -3.52143 | 0.000429 | 0.0076 | 6.81622 | 0.442226 | 1.514061 | 0.186221 | | 1 |
| 904 | ENSG00000273703 | 179.3801 | -3.57694 | 0.341398 | -10.4773 | 1.10E-25 | 2.74E-23 | 177.2826 | 16.32672 | 151.3032 | 10.27107 | H2BC14 | 6 |
| 905 | ENSG00000273706 | 46.9915 | -3.39967 | 0.62226 | -5.46342 | 4.67E-08 | 2.03E-06 | 5.786938 | 0.546125 | 3.83606 | 0.369126 | LHX1 | 17 |
| 906 | ENSG00000275126 | 178.5122 | -3.17436 | 0.321516 | -9.87309 | 5.45E-23 | 1.12E-20 | 255.3206 | 30.28453 | 237.6346 | 22.11069 | H4C13 | 6 |
| 907 | ENSG00000275379 | 598.7114 | -2.51813 | 0.191881 | -13.1234 | 2.42E-39 | 1.11E-36 | 460.7785 | 78.23391 | 418.3374 | 68.35207 | H3C11 | 6 |
| 908 | ENSG00000275805 | 20.25698 | 2.945089 | 0.904461 | 3.256181 | 0.001129 | 0.017032 | 0.163124 | 3.125123 | 0.543214 | 2.097272 | | 18 |
| 909 | ENSG00000275993 | 316.7805 | 1.931429 | 0.249066 | 7.754678 | 8.86E-15 | 9.80E-13 | 3.531216 | 10.68645 | 3.254425 | 13.84479 | | 21 |
| 910 | ENSG00000276203 | 25.05683 | -2.31625 | 0.788072 | -2.93913 | 0.003291 | 0.041027 | 3.692585 | 0.672327 | 2.143268 | 0.47772 | ANKRD20A | 9 |
| 911 | ENSG00000276266 | 15.32913 | -3.30779 | 1.035652 | -3.19392 | 0.001404 | 0.020402 | 3.562815 | 0.22558 | 2.405921 | 0.339302 | | 9 |
| 912 | ENSG00000276368 | 261.9774 | -2.51214 | 0.27625 | -9.09373 | 9.57E-20 | 1.66E-17 | 247.5139 | 37.47893 | 179.6285 | 34.05721 | H2AC14 | 6 |
| 913 | ENSG00000276547 | 72.24414 | -2.9196 | 0.483419 | -6.03947 | 1.55E-09 | 8.50E-08 | 4.392948 | 0.710328 | 5.051207 | 0.492961 | PCDHGB5 | 5 |
| 914 | ENSG00000277268 | 33.16342 | -3.54709 | 0.733817 | -4.83375 | 1.34E-06 | 4.35E-05 | 9.134623 | 0.396148 | 6.57407 | 0.896689 | LHX1-DT | 17 |
| 915 | ENSG00000277534 | 299.8221 | 1.380569 | 0.236599 | 5.835058 | 5.38E-09 | 2.72E-07 | 6.299698 | 18.50055 | 7.760975 | 16.40024 | | 18 |
| 916 | ENSG00000278905 | 186.8569 | 1.53947 | 0.292739 | 5.258842 | 1.45E-07 | 5.67E-06 | 3.972639 | 9.771059 | 2.964146 | 9.444492 | | 5 |
| 917 | ENSG00000278909 | 51.79474 | 1.487566 | 0.507595 | 2.930617 | 0.003383 | 0.041896 | 0.935256 | 2.870977 | 0.960911 | 2.212819 | | 16 |
| 918 | ENSG00000279289 | 327.7335 | 1.212014 | 0.226775 | 5.344566 | 9.06E-08 | 3.70E-06 | 5.470702 | 13.90515 | 6.037012 | 11.54147 | | 6 |
| 919 | ENSG00000279358 | 54.37045 | 1.840184 | 0.521657 | 3.527571 | 0.000419 | 0.007467 | 1.618916 | 4.888975 | 0.935125 | 3.852928 | | 11 |
| 920 | ENSG00000279417 | 157.3991 | 1.165399 | 0.302858 | 3.848007 | 0.000119 | 0.002535 | 3.071869 | 6.419847 | 2.651155 | 5.823194 | | 15 |
| 921 | ENSG00000279512 | 124.3656 | 1.453379 | 0.34013 | 4.273016 | 1.93E-05 | 0.000496 | 4.144324 | 11.64874 | 3.881304 | 9.350527 | | 2 |
| 922 | ENSG00000279521 | 287.7639 | 5.112805 | 0.34627 | 14.76536 | 2.45E-49 | 2.05E-46 | 0.602819 | 28.39884 | 1.042737 | 26.0742 | | 14 |
| 923 | ENSG00000279620 | 55.43953 | -2.11978 | 0.616962 | -3.43583 | 0.000591 | 0.009921 | 8.545654 | 2.410084 | 4.667613 | 0.581132 | | 16 |
| 924 | ENSG00000279694 | 59.83742 | 3.095127 | 0.524672 | 5.899165 | 3.65E-09 | 1.88E-07 | 0.557242 | 4.93221 | 0.626527 | 4.706652 | | 15 |
| 925 | ENSG00000279739 | 83.87149 | 1.896337 | 0.409707 | 4.628524 | 3.68E-06 | 0.000108 | 2.829728 | 11.11785 | 3.028939 | 9.689237 | | 5 |
| 926 | ENSG00000279912 | 116.3904 | 1.304974 | 0.344566 | 3.787298 | 0.000152 | 0.003125 | 2.029557 | 4.961857 | 1.930735 | 4.373361 | | 15 |
| 927 | ENSG00000280138 | 467.939 | -1.6111 | 0.201944 | -7.97796 | 1.49E-15 | 1.87E-13 | 4.197826 | 1.440495 | 4.107568 | 1.160223 | | 12 |
| 928 | ENSG00000280255 | 51.91167 | -1.48047 | 0.513337 | -2.88401 | 0.003926 | 0.046937 | 3.657725 | 0.946714 | 3.205073 | 1.372679 | | 7 |
| 929 | ENSG00000280424 | 33.15811 | -2.5819 | 0.661059 | -3.90571 | 9.39E-05 | 0.002034 | 1.400192 | 0.263867 | 1.757117 | 0.239057 | | 22 |
| 930 | ENSG00000280650 | 75.47649 | -3.57142 | 0.510952 | -6.98974 | 2.75E-12 | 2.27E-10 | 1.906779 | 0.107634 | 1.236178 | 0.141415 | KCNIP4-IT | 4 |
| 931 | ENSG00000280707 | 10.39002 | 6.729023 | 1.804251 | 3.729537 | 0.000192 | 0.00383 | 0 | 4.643014 | 0 | 9.065759 | | 6 |
| 932 | ENSG00000281406 | 588.8992 | -2.05106 | 0.198167 | -10.3502 | 4.18E-25 | 1.02E-22 | 16.99733 | 4.716561 | 18.40844 | 3.452057 | BLACAT1 | 1 |
| 933 | ENSG00000281566 | 48.70958 | -4.60563 | 0.731859 | -6.29306 | 3.11E-10 | 1.94E-08 | 17.9269 | 0.675032 | 15.2241 | 0.621449 | | X |
| 934 | ENSG00000281880 | 63.24496 | -2.16315 | 0.488814 | -4.4253 | 9.63E-06 | 0.00026 | 7.909293 | 1.361153 | 5.240103 | 1.435617 | PAUPAR | 11 |
| 935 | ENSG00000283128 | 93.13348 | 2.120513 | 0.453941 | 4.671341 | 2.99E-06 | 9.10E-05 | 0.888248 | 4.028005 | 1.753226 | 6.765031 | | 7 |

| | | | | | | | | | | | | | |
|---|---|---|---|---|---|---|---|---|---|---|---|---|---|
| 936 | ENSG00000283486 | 22.47833 | 2.431103 | 0.799757 | 3.039804 | 0.002367 | 0.031378 | 0.289485 | 1.754376 | 0.484134 | 2.157793 | NA | NA |
| 937 | ENSG00000283638 | 130.7177 | 1.062045 | 0.325367 | 3.264146 | 0.001098 | 0.016639 | 2.483696 | 4.856432 | 2.608248 | 5.249007 | | X |
| 938 | ENSG00000284299 | 12.7581 | 4.531852 | 1.412385 | 3.208651 | 0.001334 | 0.019548 | 0.05925 | 0.408168 | 0 | 0.850063 | NA | NA |
| 939 | ENSG00000284610 | 27.60985 | 2.966672 | 1.013201 | 2.92802 | 0.003411 | 0.042156 | 0.403074 | 0.396541 | 0.46015 | 7.232305 | | 8 |
| 940 | ENSG00000285041 | 18.65481 | -2.61888 | 0.891152 | -2.93876 | 0.003295 | 0.041049 | 1.025504 | 0.240143 | 1.904782 | 0.214298 | | 1 |
| 941 | ENSG00000285283 | 11.61265 | -7.04457 | 1.768943 | -3.98236 | 6.82E-05 | 0.00155 | 1.855652 | 0 | 3.531544 | 0 | | 11 |
| 942 | ENSG00000285633 | 275.025 | 4.783813 | 0.341395 | 14.01256 | 1.31E-44 | 8.38E-42 | 0.299398 | 13.74241 | 0.673201 | 11.92878 | | 14 |
| 943 | ENSG00000285846 | 19.32227 | 6.198785 | 1.651354 | 3.75376 | 0.000174 | 0.003533 | 0 | 1.562841 | 0.079665 | 4.061157 | | 10 |
| 944 | ENSG00000285850 | 27.66708 | -2.55503 | 0.727377 | -3.51267 | 0.000444 | 0.007819 | 6.474145 | 0.855026 | 4.501962 | 0.920129 | | 16 |
| 945 | ENSG00000286125 | 136.5349 | -2.20525 | 0.350926 | -6.28407 | 3.30E-10 | 2.05E-08 | 9.769231 | 1.995973 | 12.57697 | 2.563462 | | 19 |
| 946 | ENSG00000286214 | 1504.482 | -1.71582 | 0.55787 | -3.07566 | 0.0021 | 0.028447 | 41.086 | 12.21211 | 23.74116 | 6.758581 | COPG2IT1 | 7 |
| 947 | ENSG00000286215 | 100.0025 | -2.18366 | 0.414966 | -5.26226 | 1.42E-07 | 5.58E-06 | 9.542134 | 3.048009 | 11.91815 | 1.531889 | | 6 |
| 948 | ENSG00000286329 | 13.51636 | -7.26288 | 1.734256 | -4.18789 | 2.82E-05 | 0.000694 | 2.043875 | 0 | 0.972259 | 0 | | 3 |
| 949 | ENSG00000286449 | 8.886437 | -6.66069 | 1.843246 | -3.61357 | 0.000302 | 0.005627 | 0.486101 | 0 | 0.520235 | 0 | | 19 |
| 950 | ENSG00000286742 | 104.8607 | 1.106126 | 0.37543 | 2.946292 | 0.003216 | 0.040376 | 4.658214 | 8.727507 | 3.176669 | 7.4145 | | 7 |
| 951 | ENSG00000286757 | 139.0532 | 2.225046 | 0.37376 | 5.95314 | 2.63E-09 | 1.40E-07 | 1.581823 | 11.42264 | 3.489421 | 11.27706 | | 13 |
| 952 | ENSG00000287215 | 53.36706 | -1.86952 | 0.518325 | -3.60686 | 0.00031 | 0.005758 | 3.898036 | 0.829 | 2.588471 | 0.860132 | | X |
| 953 | ENSG00000287373 | 16.60901 | -7.56172 | 1.674756 | -4.51512 | 6.33E-06 | 0.000177 | 1.292907 | 0 | 0.8643 | 0 | | 11 |
| 954 | ENSG00000288079 | 515.1055 | 1.206751 | 0.196063 | 6.154924 | 7.51E-10 | 4.43E-08 | 7.324039 | 17.03782 | 8.788353 | 18.31017 | | 3 |
| 955 | ENSG00000288597 | 92.71818 | -1.41725 | 0.411183 | -3.44676 | 0.000567 | 0.009663 | 5.437323 | 2.99674 | 7.999653 | 1.855606 | | X |
| 956 | ENSG00000288658 | 65.80849 | 2.193188 | 0.469803 | 4.668315 | 3.04E-06 | 9.19E-05 | 1.029299 | 4.472005 | 0.856811 | 3.761031 | | 2 |

| | ensembl_gene_id | baseMean | log2FoldCl | lfcSE | stat | pvalue | padj | D6_WT_1 | D6_varA_1 | D6_WT_2 | D6_varA_2 | hgnc_symb | chromosome_name |
|---|---|---|---|---|---|---|---|---|---|---|---|---|---|
| 1 | ENSG00000001036 | 356.5369 | 1.19467 | 0.211522 | 5.64798 | 1.62E-08 | 5.87E-07 | 12.27775 | 25.46014 | 11.41779 | 25.72216 | FUCA2 | 6 |
| 2 | ENSG00000002746 | 271.9148 | -1.05752 | 0.245414 | -4.30913 | 1.64E-05 | 0.000331 | 11.11017 | 4.53479 | 8.503695 | 4.333933 | HECW1 | 7 |
| 3 | ENSG00000003137 | 64.80972 | -1.84777 | 0.526805 | -3.50751 | 0.000452 | 0.006286 | 1.493956 | 0.643646 | 3.69018 | 0.728387 | CYP26B1 | 2 |
| 4 | ENSG00000003402 | 436.1943 | 1.163023 | 0.209613 | 5.548421 | 2.88E-08 | 1.01E-06 | 6.894823 | 12.0064 | 6.120549 | 15.54205 | CFLAR | 2 |
| 5 | ENSG00000004799 | 47.23909 | -1.50019 | 0.52178 | -2.87514 | 0.004039 | 0.038805 | 3.000824 | 0.987787 | 2.505164 | 0.851025 | PDK4 | 7 |
| 6 | ENSG00000004848 | 209.9812 | -5.65909 | 0.444997 | -12.7171 | 4.75E-37 | 2.27E-34 | 19.99704 | 0.282882 | 13.67272 | 0.34678 | ARX | X |
| 7 | ENSG00000005249 | 804.0645 | -1.25267 | 0.17115 | -7.31909 | 2.50E-13 | 1.62E-11 | 40.06481 | 14.40321 | 32.44061 | 14.28085 | PRKAR2B | 7 |
| 8 | ENSG00000005379 | 284.1119 | -1.23087 | 0.237403 | -5.18472 | 2.16E-07 | 6.46E-06 | 17.64448 | 6.731917 | 14.70996 | 6.248056 | TSPOAP1 | 17 |
| 9 | ENSG00000006128 | 52.7113 | 1.513027 | 0.501318 | 3.018102 | 0.002544 | 0.026612 | 4.817509 | 11.58256 | 3.601836 | 10.93891 | TAC1 | 7 |
| 10 | ENSG00000006210 | 109.6805 | -1.64136 | 0.373386 | -4.39586 | 1.10E-05 | 0.000232 | 6.064386 | 2.100914 | 5.365787 | 1.342612 | CX3CL1 | 16 |
| 11 | ENSG00000006327 | 92.15385 | 1.723379 | 0.456467 | 3.775477 | 0.00016 | 0.002527 | 4.153272 | 25.79322 | 9.311275 | 16.43617 | TNFRSF12/ | 16 |
| 12 | ENSG00000006377 | 38.18586 | -7.34961 | 1.474624 | -4.98406 | 6.23E-07 | 1.69E-05 | 6.722587 | 0.045594 | 3.515612 | 0 | DLX6 | 7 |
| 13 | ENSG00000007174 | 114.0905 | -1.73348 | 0.356969 | -4.85612 | 1.20E-06 | 3.05E-05 | 5.021107 | 1.377885 | 6.007192 | 1.778141 | DNAH9 | 17 |
| 14 | ENSG00000007350 | 35.5013 | -2.27784 | 0.769908 | -2.95858 | 0.003091 | 0.031018 | 5.53552 | 0.370886 | 1.596973 | 1.155216 | TKTL1 | X |
| 15 | ENSG00000007372 | 3667.417 | -2.01439 | 0.156038 | -12.9096 | 3.97E-38 | 2.04E-35 | 309.6462 | 73.18579 | 250.4006 | 57.26516 | PAX6 | 11 |
| 16 | ENSG00000008196 | 573.1368 | 4.070451 | 0.215241 | 18.91111 | 9.24E-80 | 2.01E-76 | 1.496303 | 21.83087 | 1.432962 | 24.66092 | TFAP2B | 6 |
| 17 | ENSG00000008394 | 205.8718 | 2.393375 | 0.297424 | 8.047003 | 8.48E-16 | 7.15E-14 | 9.136078 | 56.82709 | 14.27152 | 59.5141 | MGST1 | 12 |
| 18 | ENSG00000009694 | 476.1859 | -1.1167 | 0.196205 | -5.69146 | 1.26E-08 | 4.66E-07 | 5.962463 | 2.5726 | 5.262125 | 2.305061 | TENM1 | X |
| 19 | ENSG00000009709 | 161.7154 | 2.296446 | 0.307024 | 7.479705 | 7.45E-14 | 5.11E-12 | 1.238802 | 5.85013 | 1.260559 | 5.754684 | PAX7 | 1 |
| 20 | ENSG00000009950 | 28.48402 | 3.338407 | 0.814422 | 4.099111 | 4.15E-05 | 0.000763 | 0.614943 | 2.561567 | 0.175305 | 3.913681 | MLXIPL | 7 |
| 21 | ENSG00000010278 | 151.3232 | 1.606356 | 0.350664 | 4.580904 | 4.63E-06 | 0.000106 | 8.977692 | 28.11993 | 6.819998 | 17.00736 | CD9 | 12 |
| 22 | ENSG00000011201 | 3508.452 | -3.21942 | 0.406563 | -7.91862 | 2.40E-15 | 1.95E-13 | 119.5505 | 15.97049 | 131.7287 | 9.394928 | ANOS1 | X |
| 23 | ENSG00000011677 | 172.1296 | -1.49815 | 0.326285 | -4.59154 | 4.40E-06 | 0.000101 | 8.766859 | 3.36186 | 7.292417 | 1.966108 | GABRA3 | X |
| 24 | ENSG00000015592 | 755.4712 | -1.18484 | 0.183585 | -6.45391 | 1.09E-10 | 5.49E-09 | 118.5532 | 42.84619 | 88.27048 | 42.88659 | STMN4 | 8 |
| 25 | ENSG00000016082 | 20.44267 | -4.60249 | 1.151403 | -3.99729 | 6.41E-05 | 0.001132 | 3.49546 | 0.087188 | 1.380157 | 0.105758 | ISL1 | 5 |
| 26 | ENSG00000018408 | 649.6212 | 1.289339 | 0.174607 | 7.384241 | 1.53E-13 | 1.01E-11 | 10.04643 | 21.23132 | 9.784934 | 24.63817 | WWTR1 | 3 |
| 27 | ENSG00000018625 | 372.5785 | 2.115016 | 0.238958 | 8.850992 | 8.67E-19 | 9.80E-17 | 3.012597 | 15.24383 | 3.605826 | 12.04308 | ATP1A2 | 1 |
| 28 | ENSG00000019549 | 55.35991 | 2.068053 | 0.51249 | 4.035302 | 5.45E-05 | 0.000979 | 1.241099 | 5.121576 | 0.984602 | 3.603408 | SNAI2 | 8 |
| 29 | ENSG00000019582 | 46.48945 | 1.723948 | 0.579452 | 2.975134 | 0.002929 | 0.029718 | 0.754421 | 4.520132 | 1.62912 | 2.890173 | CD74 | 5 |
| 30 | ENSG00000021645 | 785.8734 | -1.69067 | 0.223439 | -7.56659 | 3.83E-14 | 2.74E-12 | 41.52313 | 11.85301 | 27.71474 | 8.279786 | NRXN3 | 14 |
| 31 | ENSG00000021826 | 877.7859 | 1.141037 | 0.160193 | 7.122883 | 1.06E-12 | 6.42E-11 | 11.92711 | 26.22245 | 12.6804 | 25.05024 | CPS1 | 2 |
| 32 | ENSG00000024422 | 88.20052 | 2.61758 | 0.44538 | 5.877184 | 4.17E-09 | 1.67E-07 | 0.625946 | 5.816713 | 1.111489 | 4.264013 | EHD2 | 19 |
| 33 | ENSG00000026508 | 186.0958 | 1.927911 | 0.313937 | 6.141076 | 8.20E-10 | 3.72E-08 | 4.896267 | 15.87202 | 3.159968 | 12.66081 | CD44 | 11 |
| 34 | ENSG00000026559 | 147.2066 | 2.413416 | 0.323798 | 7.453454 | 9.09E-14 | 6.17E-12 | 3.456172 | 16.03084 | 3.350206 | 18.24556 | KCNG1 | 20 |
| 35 | ENSG00000028116 | 66.42699 | 1.577234 | 0.497759 | 3.168672 | 0.001531 | 0.017768 | 1.575957 | 7.981697 | 3.0155 | 5.00014 | VRK2 | 2 |
| 36 | ENSG00000033122 | 437.6271 | -1.57786 | 0.207716 | -7.59623 | 3.05E-14 | 2.22E-12 | 7.783687 | 2.694155 | 7.7583 | 2.211197 | LRRC7 | 1 |
| 37 | ENSG00000033170 | 772.7175 | -1.2313 | 0.181831 | -6.77165 | 1.27E-11 | 6.97E-10 | 52.94536 | 19.22482 | 40.25496 | 18.16636 | FUT8 | 14 |
| 38 | ENSG00000036565 | 76.73563 | -2.17028 | 0.435692 | -4.98123 | 6.32E-07 | 1.70E-05 | 8.357232 | 1.978192 | 7.992872 | 1.388053 | SLC18A1 | 8 |
| 39 | ENSG00000038295 | 13.61985 | 3.030165 | 1.091315 | 2.77662 | 0.005493 | 0.049201 | 0.158584 | 1.387409 | 0.069848 | 0.311635 | TLL1 | 4 |
| 40 | ENSG00000039139 | 129.3326 | -1.26924 | 0.356272 | -3.56256 | 0.000367 | 0.00528 | 2.131694 | 1.028291 | 3.470924 | 1.177903 | DNAH5 | 5 |
| 41 | ENSG00000040731 | 239.5089 | -1.70043 | 0.260529 | -6.52682 | 6.72E-11 | 3.42E-09 | 12.89837 | 4.045236 | 12.10148 | 3.194706 | CDH10 | 5 |
| 42 | ENSG00000042980 | 13.24176 | 3.103982 | 1.11235 | 2.790473 | 0.005263 | 0.047558 | 0.21629 | 0.980294 | 0.031207 | 0.743271 | ADAM28 | 8 |
| 43 | ENSG00000044524 | 3850.314 | -1.12147 | 0.346839 | -3.23341 | 0.001223 | 0.014685 | 152.1462 | 56.53409 | 93.67678 | 49.73264 | EPHA3 | 3 |
| 44 | ENSG00000046889 | 1743.544 | -1.25605 | 0.195433 | -6.42699 | 1.30E-10 | 6.47E-09 | 48.44855 | 16.36995 | 29.48373 | 14.30021 | PREX2 | 8 |
| 45 | ENSG00000047932 | 472.5921 | -1.01113 | 0.190336 | -5.31231 | 1.08E-07 | 3.42E-06 | 16.15302 | 6.970977 | 15.22135 | 7.740788 | GOPC | 6 |
| 46 | ENSG00000048540 | 748.5109 | -1.29324 | 0.193842 | -6.67162 | 2.53E-11 | 1.34E-09 | 50.53411 | 18.2486 | 37.24176 | 15.44666 | LMO3 | 12 |
| 47 | ENSG00000049130 | 466.1083 | 1.215918 | 0.198808 | 6.116036 | 9.59E-10 | 4.30E-08 | 5.510519 | 13.34069 | 6.362461 | 12.73013 | KITLG | 12 |
| 48 | ENSG00000049540 | 131.0923 | 1.186384 | 0.354006 | 3.35131 | 0.000804 | 0.010273 | 3.280373 | 10.02086 | 5.389922 | 8.530298 | ELN | 7 |
| 49 | ENSG00000050030 | 561.1957 | -1.00029 | 0.200018 | -5.00103 | 5.70E-07 | 1.56E-05 | 7.820641 | 3.124527 | 5.680699 | 3.236479 | NEXMIF | X |
| 50 | ENSG00000050767 | 23.77257 | 2.303568 | 0.784336 | 2.936967 | 0.003314 | 0.032882 | 0.579218 | 1.806142 | 0.340516 | 2.467614 | COL23A1 | 5 |
| 51 | ENSG00000052126 | 2376.675 | -1.2665 | 0.125537 | -10.0886 | 6.20E-24 | 1.12E-21 | 148.4546 | 55.95225 | 140.053 | 57.28613 | PLEKHA5 | 12 |
| 52 | ENSG00000053438 | 3365.422 | -2.40437 | 0.130251 | -18.4594 | 4.38E-76 | 6.12E-73 | 642.8234 | 117.0198 | 600.4858 | 104.4058 | NNAT | 20 |
| 53 | ENSG00000059804 | 2020.298 | 2.059531 | 0.160129 | 12.86169 | 7.39E-38 | 3.71E-35 | 27.94733 | 132.7949 | 34.0683 | 111.4476 | SLC2A3 | 12 |
| 54 | ENSG00000060709 | 271.8407 | -1.01319 | 0.239702 | -4.22686 | 2.37E-05 | 0.000459 | 15.79112 | 6.596415 | 12.8012 | 6.755963 | RIMBP2 | 12 |
| 55 | ENSG00000064012 | 23.86457 | 2.447257 | 0.782396 | 3.127899 | 0.001761 | 0.019913 | 0.580193 | 2.860468 | 0.424009 | 2.146361 | CASP8 | 2 |
| 56 | ENSG00000064218 | 44.32497 | -2.70321 | 0.597793 | -4.52198 | 6.13E-06 | 0.000137 | 3.525321 | 0.583367 | 5.350878 | 0.724015 | DMRT3 | 9 |
| 57 | ENSG00000064300 | 162.0348 | 1.27639 | 0.303108 | 4.211007 | 2.54E-05 | 0.000489 | 3.416514 | 8.581641 | 3.378621 | 6.919411 | NGFR | 17 |
| 58 | ENSG00000064651 | 1256.07 | 1.080227 | 0.152779 | 7.070509 | 1.54E-12 | 9.23E-11 | 17.52824 | 38.61249 | 19.94376 | 36.22001 | SLC12A2 | 5 |
| 59 | ENSG00000064692 | 239.1865 | -1.76888 | 0.357347 | -4.95003 | 7.42E-07 | 1.96E-05 | 25.64433 | 6.557834 | 11.81575 | 3.789698 | SNCAIP | 5 |
| 60 | ENSG00000065357 | 78.80925 | 1.200148 | 0.416214 | 2.883487 | 0.003933 | 0.037959 | 6.285503 | 13.11337 | 4.899178 | 10.87084 | DGKA | 12 |
| 61 | ENSG00000065413 | 202.6026 | -1.41751 | 0.290318 | -4.8826 | 1.05E-06 | 2.71E-05 | 7.351736 | 3.185998 | 10.72929 | 3.169387 | ANKRD44 | 2 |
| 62 | ENSG00000065485 | 129.7252 | 1.083165 | 0.384227 | 2.819074 | 0.004816 | 0.044266 | 5.036968 | 13.86002 | 10.66749 | 17.56001 | PDIA5 | 3 |
| 63 | ENSG00000066382 | 238.9301 | -3.58448 | 0.320659 | -11.1785 | 5.20E-29 | 1.32E-26 | 36.38691 | 1.962284 | 22.04302 | 2.642506 | MPPED2 | 11 |
| 64 | ENSG00000066468 | 685.1731 | 1.429931 | 0.173411 | 8.245913 | 1.64E-16 | 1.46E-14 | 14.12516 | 35.46062 | 12.8846 | 33.19085 | FGFR2 | 10 |
| 65 | ENSG00000067182 | 240.1009 | 1.119525 | 0.351164 | 3.188036 | 0.001432 | 0.016792 | 6.117311 | 20.82767 | 15.87357 | 24.2034 | TNFRSF1A | 12 |
| 66 | ENSG00000068724 | 207.432 | 1.261959 | 0.267799 | 4.712328 | 2.45E-06 | 5.89E-05 | 3.486309 | 8.670965 | 3.96838 | 8.214657 | TTC7A | 2 |
| 67 | ENSG00000068781 | 49.3575 | -1.85815 | 0.572623 | -3.24498 | 0.001175 | 0.014239 | 3.544347 | 0.921399 | 1.88013 | 0.460347 | STON1-GT | 2 |
| 68 | ENSG00000069696 | 47.48614 | -1.99494 | 0.553103 | -3.60682 | 0.00031 | 0.004561 | 6.971988 | 2.197245 | 7.271438 | 1.151383 | DRD4 | 11 |
| 69 | ENSG00000069702 | 113.7814 | 1.230865 | 0.410917 | 2.995408 | 0.002741 | 0.028148 | 3.297919 | 4.099138 | 1.606054 | 6.883687 | TGFBR3 | 1 |
| 70 | ENSG00000070193 | 191.1907 | 6.568733 | 0.573884 | 11.4461 | 2.46E-30 | 6.97E-28 | 0.061384 | 8.344809 | 0.127575 | 7.995322 | FGF10 | 5 |
| 71 | ENSG00000070669 | 763.0324 | 1.433417 | 0.164873 | 8.694089 | 3.50E-18 | 3.74E-16 | 30.11777 | 71.05776 | 27.35546 | 75.49813 | ASNS | 7 |
| 72 | ENSG00000070729 | 73.60958 | -2.82148 | 0.469757 | -6.00624 | 1.90E-09 | 8.07E-08 | 3.727149 | 0.397463 | 4.364519 | 0.730841 | CNGB1 | 16 |
| 73 | ENSG00000071909 | 15.63029 | 2.658278 | 0.945666 | 2.81101 | 0.004939 | 0.045147 | 0.127694 | 0.951463 | 0.121926 | 0.545519 | MYO3B | 2 |
| 74 | ENSG00000071967 | 180.6904 | 1.4462 | 0.324149 | 4.461526 | 8.14E-06 | 0.000177 | 2.309847 | 8.627063 | 4.163289 | 8.00328 | CYBRD1 | 2 |
| 75 | ENSG00000072682 | 122.4518 | 1.215183 | 0.342818 | 3.544685 | 0.000393 | 0.00559 | 3.173191 | 8.161623 | 4.380508 | 8.459859 | P4HA2 | 5 |
| 76 | ENSG00000073464 | 973.531 | -1.07864 | 0.159643 | -6.75657 | 1.41E-11 | 7.72E-10 | 24.7635 | 10.23828 | 20.74755 | 10.08012 | CLCN4 | X |
| 77 | ENSG00000073910 | 588.3727 | -1.10095 | 0.202336 | -5.44118 | 5.29E-08 | 1.77E-06 | 12.7596 | 4.792527 | 8.991541 | 4.752158 | FRY | 13 |

| | | | | | | | | | | | | | | |
|---|---|---|---|---|---|---|---|---|---|---|---|---|---|---|
| 78 | ENSG00000074211 | 77.79683 | -2.3058 | 0.442302 | -5.21317 | 1.86E-07 | 5.60E-06 | 3.68882 | 0.859578 | 4.840382 | 0.773283 | PPP2R2C | | 4 |
| 79 | ENSG00000074706 | 19.57867 | -2.49947 | 0.901062 | -2.77392 | 0.005539 | 0.049588 | 0.958068 | 0.266784 | 0.971686 | 0.055748 | IPCEF1 | | 6 |
| 80 | ENSG00000075035 | 85.82384 | 1.876463 | 0.409403 | 4.583412 | 4.57E-06 | 0.000105 | 1.934492 | 5.555134 | 1.629434 | 6.781154 | WSCD2 | | 12 |
| 81 | ENSG00000075223 | 491.2289 | 1.522153 | 0.249111 | 6.110343 | 9.94E-10 | 4.44E-08 | 11.00016 | 24.75636 | 6.073898 | 21.08778 | SEMA3C | | 7 |
| 82 | ENSG00000075426 | 323.0775 | 1.759372 | 0.242556 | 7.253472 | 4.06E-13 | 2.60E-11 | 2.625736 | 9.413133 | 2.582101 | 7.175704 | FOSL2 | | 2 |
| 83 | ENSG00000076356 | 1602.558 | -1.176 | 0.143813 | -8.17732 | 2.90E-16 | 2.51E-14 | 34.56801 | 13.91266 | 29.89763 | 12.98821 | PLXNA2 | | 1 |
| 84 | ENSG00000076706 | 491.0099 | 1.047302 | 0.198198 | 5.284129 | 1.26E-07 | 3.92E-06 | 12.61012 | 24.29181 | 10.793 | 21.27644 | MCAM | | 11 |
| 85 | ENSG00000076770 | 552.7313 | 1.342085 | 0.190362 | 7.050185 | 1.79E-12 | 1.05E-10 | 4.074217 | 8.903947 | 3.25898 | 8.628405 | MBNL3 | X | |
| 86 | ENSG00000077264 | 5834.226 | -1.33689 | 0.136866 | -9.76792 | 1.55E-22 | 2.34E-20 | 145.6882 | 51.72763 | 116.9903 | 46.235 | PAK3 | X | |
| 87 | ENSG00000077327 | 47.46961 | -1.63064 | 0.540776 | -3.01537 | 0.002567 | 0.026781 | 5.558876 | 1.765683 | 4.017699 | 1.152093 | SPAG6 | | 10 |
| 88 | ENSG00000077616 | 238.1323 | -1.14888 | 0.248612 | -4.62117 | 3.82E-06 | 8.88E-05 | 13.47376 | 5.278136 | 13.28424 | 6.142858 | NAALAD2 | | 11 |
| 89 | ENSG00000077943 | 30.54401 | -1.89931 | 0.671756 | -2.82738 | 0.004693 | 0.043504 | 1.06693 | 0.348864 | 1.511189 | 0.298831 | ITGA8 | | 10 |
| 90 | ENSG00000078070 | 273.9067 | -1.41344 | 0.235311 | -6.0067 | 1.89E-09 | 8.07E-08 | 27.40492 | 10.25903 | 29.16349 | 9.804525 | MCCC1 | | 3 |
| 91 | ENSG00000078295 | 309.4172 | -2.49865 | 0.273349 | -9.14087 | 6.20E-20 | 7.77E-18 | 20.78878 | 2.506209 | 13.33807 | 3.231702 | ADCY2 | | 5 |
| 92 | ENSG00000078401 | 99.85905 | 3.55886 | 0.466167 | 7.634298 | 2.27E-14 | 1.68E-12 | 0.939152 | 13.24881 | 0.947438 | 7.642924 | EDN1 | | 6 |
| 93 | ENSG00000078549 | 411.0173 | 1.730405 | 0.205777 | 8.409129 | 4.13E-17 | 3.88E-15 | 4.120714 | 12.19637 | 3.642038 | 12.07155 | ADCYAP1F | | 7 |
| 94 | ENSG00000078725 | 200.9184 | -1.36237 | 0.28366 | -4.80281 | 1.56E-06 | 3.90E-05 | 12.59011 | 4.447125 | 9.596119 | 3.665887 | BRINP1 | | 9 |
| 95 | ENSG00000079102 | 868.2019 | -1.04965 | 0.182072 | -5.76503 | 8.16E-09 | 3.11E-07 | 43.9428 | 22.88453 | 44.274 | 17.28798 | RUNX1T1 | | 8 |
| 96 | ENSG00000079215 | 1447.47 | -1.22862 | 0.202112 | -6.07893 | 1.21E-09 | 5.29E-08 | 84.83543 | 29.40398 | 51.12339 | 25.1231 | SLC1A3 | | 5 |
| 97 | ENSG00000079931 | 253.9081 | -3.49691 | 0.317094 | -11.028 | 2.80E-28 | 6.76E-26 | 24.58477 | 1.850482 | 14.75249 | 1.414903 | MOXD1 | | 6 |
| 98 | ENSG00000080031 | 75.66087 | 1.662083 | 0.508961 | 3.265641 | 0.001092 | 0.013373 | 1.008815 | 6.256034 | 3.251431 | 6.22138 | PTPRH | | 19 |
| 99 | ENSG00000080166 | 195.8519 | -3.78526 | 0.75111 | -5.03956 | 4.67E-07 | 1.31E-05 | 12.87436 | 0.953138 | 6.726664 | 0.391778 | DCT | | 13 |
| 100 | ENSG00000080493 | 77.32706 | -1.43068 | 0.417962 | -3.423 | 0.000619 | 0.008255 | 2.650699 | 1.007363 | 2.499849 | 0.787203 | SLC4A4 | | 4 |
| 101 | ENSG00000080561 | 186.7987 | -1.00801 | 0.330163 | -3.05305 | 0.002265 | 0.024398 | 5.364761 | 3.892251 | 9.356151 | 2.995333 | MID2 | X | |
| 102 | ENSG00000080573 | 125.1892 | 2.341703 | 0.3765 | 6.219656 | 4.98E-10 | 2.32E-08 | 0.563888 | 4.078148 | 0.939742 | 3.151364 | COL5A3 | | 19 |
| 103 | ENSG00000080644 | 182.121 | -1.53461 | 0.288647 | -5.31657 | 1.06E-07 | 3.36E-06 | 14.16031 | 5.40176 | 17.19155 | 4.836634 | CHRNA3 | | 15 |
| 104 | ENSG00000081052 | 40.03221 | 2.498792 | 0.598209 | 4.177124 | 2.95E-05 | 0.000562 | 0.123284 | 0.875704 | 0.184445 | 0.776912 | COL4A4 | | 2 |
| 105 | ENSG00000081138 | 428.307 | -3.30799 | 0.274866 | -12.0349 | 2.33E-33 | 8.43E-31 | 23.78136 | 2.516056 | 16.891 | 1.385626 | CDH7 | | 18 |
| 106 | ENSG00000081665 | 575.5178 | -1.44522 | 0.1804 | -8.01122 | 1.14E-15 | 9.41E-14 | 35.0267 | 11.033 | 32.84311 | 12.53059 | ZNF506 | | 19 |
| 107 | ENSG00000081803 | 328.5269 | -2.50334 | 0.26165 | -9.56753 | 1.09E-21 | 1.56E-19 | 14.16678 | 3.017064 | 15.34783 | 1.867309 | CADPS2 | | 7 |
| 108 | ENSG00000082175 | 20.88697 | 2.887479 | 0.865562 | 3.335958 | 0.00085 | 0.010773 | 0.15045 | 0.920546 | 0.057812 | 0.490697 | PGR | | 11 |
| 109 | ENSG00000082438 | 105.5328 | 3.141536 | 0.447585 | 7.018866 | 2.24E-12 | 1.31E-10 | 0.505125 | 7.028324 | 0.807743 | 3.77716 | COBLL1 | | 2 |
| 110 | ENSG00000082497 | 122.6812 | 1.593114 | 0.339349 | 4.694613 | 2.67E-06 | 6.39E-05 | 1.640481 | 4.272529 | 1.770219 | 5.495853 | SERTAD4 | | 1 |
| 111 | ENSG00000082512 | 139.0194 | 1.024297 | 0.310647 | 3.297297 | 0.000976 | 0.012097 | 3.45537 | 6.563694 | 3.403305 | 6.610313 | TRAF5 | | 1 |
| 112 | ENSG00000082684 | 381.169 | 1.239125 | 0.212529 | 5.830373 | 5.53E-09 | 2.16E-07 | 9.917935 | 18.9367 | 8.58434 | 22.32299 | SEMA5B | | 3 |
| 113 | ENSG00000083067 | 1028.861 | 1.073838 | 0.173446 | 6.191182 | 5.97E-10 | 2.74E-08 | 15.90347 | 37.57923 | 18.72663 | 31.16325 | TRPM3 | | 9 |
| 114 | ENSG00000084710 | 978.2594 | -1.11486 | 0.154917 | -7.1965 | 6.18E-13 | 3.85E-11 | 19.65925 | 8.847596 | 21.32917 | 9.041651 | EFR3B | | 2 |
| 115 | ENSG00000085276 | 383.8055 | -1.23111 | 0.242659 | -5.07342 | 3.91E-07 | 1.11E-05 | 12.63417 | 6.652923 | 18.46089 | 5.858014 | MECOM | | 3 |
| 116 | ENSG00000085563 | 45.80025 | -1.82566 | 0.559988 | -3.26017 | 0.001113 | 0.013582 | 3.290757 | 0.670788 | 1.926029 | 0.709342 | ABCB1 | | 7 |
| 117 | ENSG00000085999 | 189.2196 | 1.3982 | 0.273734 | 5.107879 | 3.26E-07 | 9.41E-06 | 7.061219 | 16.7189 | 6.300292 | 16.5102 | RAD54L | | 1 |
| 118 | ENSG00000086696 | 111.0038 | 4.097279 | 0.441681 | 9.276553 | 1.75E-20 | 2.28E-18 | 0.630404 | 14.32323 | 1.043545 | 12.32151 | HSD17B2 | | 16 |
| 119 | ENSG00000087258 | 1652.293 | -1.01128 | 0.148247 | -6.82159 | 9.00E-12 | 5.04E-10 | 95.66628 | 37.53926 | 82.35058 | 45.9498 | GNAO1 | | 16 |
| 120 | ENSG00000087495 | 158.9158 | -2.5726 | 0.325767 | -7.89706 | 2.86E-15 | 2.28E-13 | 15.24799 | 2.060637 | 11.11868 | 2.118591 | PHACTR3 | | 20 |
| 121 | ENSG00000087510 | 145.3677 | -3.64856 | 0.381554 | -9.56239 | 1.15E-21 | 1.63E-19 | 12.43401 | 1.073402 | 9.830209 | 0.584822 | TFAP2C | | 20 |
| 122 | ENSG00000089116 | 293.6419 | -3.15227 | 0.260434 | -12.1039 | 1.01E-33 | 3.79E-31 | 23.83175 | 2.816173 | 22.54216 | 2.086597 | LHX5 | | 12 |
| 123 | ENSG00000091129 | 2776.738 | -2.09938 | 0.157622 | -13.3191 | 1.79E-40 | 1.00E-37 | 128.1951 | 21.7245 | 102.7158 | 29.32445 | NRCAM | | 7 |
| 124 | ENSG00000091409 | 1241.051 | -1.18053 | 0.157336 | -7.50324 | 6.23E-14 | 4.32E-12 | 54.39759 | 22.67695 | 47.23373 | 19.56821 | ITGA6 | | 2 |
| 125 | ENSG00000091986 | 715.1053 | 1.739657 | 0.501033 | 3.472137 | 0.000516 | 0.007031 | 6.493654 | 27.68826 | 12.88874 | 33.81499 | CCDC80 | | 3 |
| 126 | ENSG00000092421 | 4449.974 | -1.49619 | 0.122441 | -12.2197 | 2.44E-34 | 1.02E-31 | 228.8639 | 79.00766 | 223.6138 | 72.2971 | SEMA6A | | 5 |
| 127 | ENSG00000092758 | 44.99898 | 1.836228 | 0.551883 | 3.327205 | 0.000877 | 0.011052 | 1.518022 | 6.172338 | 1.731415 | 4.858811 | COL9A3 | | 20 |
| 128 | ENSG00000095596 | 134.0227 | 4.185755 | 0.427933 | 9.781326 | 1.35E-22 | 2.07E-20 | 1.155224 | 21.82886 | 1.869222 | 32.06422 | CYP26A1 | | 10 |
| 129 | ENSG00000096060 | 742.8126 | 1.713189 | 0.199742 | 8.577002 | 9.74E-18 | 1.00E-15 | 8.526289 | 26.8698 | 11.38754 | 35.14247 | FKBP5 | | 6 |
| 130 | ENSG00000096696 | 182.6836 | 2.693428 | 0.319349 | 8.434115 | 3.34E-17 | 3.20E-15 | 0.622435 | 4.113236 | 0.544975 | 2.990138 | DSP | | 6 |
| 131 | ENSG00000099250 | 1700.15 | -1.42205 | 0.172504 | -8.24362 | 1.67E-16 | 1.48E-14 | 91.22798 | 37.25891 | 90.0225 | 26.50925 | NRP1 | | 10 |
| 132 | ENSG00000099337 | 23.82155 | 2.215216 | 0.74595 | 2.969659 | 0.002981 | 0.030126 | 0.17541 | 0.802703 | 0.212372 | 0.908304 | KCNK6 | | 19 |
| 133 | ENSG00000099860 | 38.01502 | 1.918949 | 0.592845 | 3.236848 | 0.001209 | 0.014579 | 2.023443 | 6.521523 | 1.782414 | 7.051092 | GADD45B | | 19 |
| 134 | ENSG00000099994 | 47.92373 | -2.55914 | 0.584708 | -4.37678 | 1.20E-05 | 0.000251 | 2.432174 | 0.634885 | 3.34137 | 0.278831 | SUSD2 | | 22 |
| 135 | ENSG00000100003 | 164.5225 | 1.03613 | 0.313386 | 3.306242 | 0.000946 | 0.011801 | 6.131554 | 13.70584 | 5.853422 | 9.478544 | SEC14L2 | | 22 |
| 136 | ENSG00000100095 | 430.8613 | -1.08059 | 0.203031 | -5.32231 | 1.02E-07 | 3.26E-06 | 12.16522 | 5.162752 | 10.27944 | 4.840969 | SEZ6L | | 22 |
| 137 | ENSG00000100311 | 233.2738 | 1.47888 | 0.257574 | 5.741567 | 9.38E-09 | 3.53E-07 | 5.026705 | 11.74149 | 4.106436 | 12.17121 | PDGFB | | 22 |
| 138 | ENSG00000100345 | 3564.167 | 1.069516 | 0.168293 | 6.35509 | 2.08E-10 | 1.01E-08 | 41.26818 | 105.9079 | 53.96384 | 82.68786 | MYH9 | | 22 |
| 139 | ENSG00000100504 | 162.7403 | 1.021404 | 0.313496 | 3.258107 | 0.001122 | 0.013664 | 6.246177 | 11.57295 | 4.584294 | 8.998657 | PYGL | | 14 |
| 140 | ENSG00000100889 | 87.12199 | 1.266939 | 0.401236 | 3.15759 | 0.001591 | 0.018319 | 3.194334 | 6.004576 | 3.277423 | 8.760956 | PCK2 | | 14 |
| 141 | ENSG00000100918 | 88.88167 | -1.32637 | 0.396813 | -3.34257 | 0.00083 | 0.010562 | 9.141313 | 3.99889 | 12.51637 | 4.172 | REC8 | | 14 |
| 142 | ENSG00000100968 | 177.9906 | 1.046311 | 0.279889 | 3.7383 | 0.000185 | 0.002887 | 7.355044 | 13.01203 | 6.644184 | 14.27299 | NFATC4 | | 14 |
| 143 | ENSG00000101000 | 28.44296 | 2.770226 | 0.730502 | 3.792223 | 0.000149 | 0.002393 | 0.652639 | 5.275293 | 0.769904 | 3.888867 | PROCR | | 20 |
| 144 | ENSG00000101096 | 42.5101 | 3.552727 | 0.613997 | 5.786231 | 7.20E-09 | 2.76E-07 | 0.133556 | 2.214714 | 0.291336 | 2.500425 | NFATC2 | | 20 |
| 145 | ENSG00000101204 | 331.8581 | 1.278871 | 0.233997 | 5.465331 | 4.62E-08 | 1.56E-06 | 5.813632 | 10.53961 | 4.842902 | 13.9065 | CHRNA4 | | 20 |
| 146 | ENSG00000101335 | 369.3483 | 2.597713 | 0.262017 | 9.914311 | 3.61E-23 | 6.08E-21 | 8.745151 | 65.16623 | 10.30521 | 43.51513 | MYL9 | | 20 |
| 147 | ENSG00000101336 | 9.663803 | 6.623618 | 1.80675 | 3.666039 | 0.000246 | 0.003722 | 0 | 1.030052 | 0 | 1.353172 | HCK | | 20 |
| 148 | ENSG00000101349 | 196.7696 | -1.20279 | 0.270918 | -4.43969 | 9.01E-06 | 0.000194 | 6.901739 | 2.827228 | 6.28068 | 2.570435 | PAK5 | | 20 |
| 149 | ENSG00000101438 | 20.96466 | -2.67597 | 0.919069 | -2.91162 | 0.003596 | 0.035225 | 2.031626 | 0.446995 | 1.378364 | 0.044274 | SLC32A1 | | 20 |
| 150 | ENSG00000101489 | 356.1102 | -1.41073 | 0.256972 | -5.48983 | 4.02E-08 | 1.37E-06 | 33.52925 | 8.090738 | 22.5795 | 11.97045 | CELF4 | | 18 |
| 151 | ENSG00000101542 | 697.0874 | -4.19045 | 0.223057 | -18.7864 | 9.75E-79 | 1.91E-75 | 50.13962 | 2.615203 | 37.95308 | 1.91598 | CDH20 | | 18 |
| 152 | ENSG00000101746 | 239.2662 | -1.68635 | 0.262132 | -6.43321 | 1.25E-10 | 6.25E-09 | 13.24691 | 4.10163 | 11.66938 | 3.217116 | NOL4 | | 18 |
| 153 | ENSG00000101825 | 391.9377 | 1.815749 | 0.215995 | 8.406443 | 4.23E-17 | 3.95E-15 | 1.7836 | 6.102223 | 2.122398 | 6.931222 | MXRA5 | X | |
| 154 | ENSG00000101856 | 3549.456 | 1.067465 | 0.131366 | 8.125885 | 4.44E-16 | 3.78E-14 | 147.8205 | 318.2781 | 155.6807 | 281.9826 | PGRMC1 | X | |
| 155 | ENSG00000102034 | 127.9224 | 1.199113 | 0.35217 | 3.404931 | 0.000662 | 0.008732 | 1.539425 | 4.533695 | 2.506227 | 4.288813 | ELF4 | X | |

| | | | | | | | | | | | | | | |
|---|---|---|---|---|---|---|---|---|---|---|---|---|---|---|
| 156 | ENSG00000102265 | 92.01148 | 1.261505 | 0.421366 | 2.993844 | 0.002755 | 0.028233 | 8.600384 | 25.53897 | 16.75902 | 32.31449 | TIMP1 | X | |
| 157 | ENSG00000102290 | 485.2309 | 1.701177 | 0.200746 | 8.47428 | 2.37E-17 | 2.34E-15 | 5.157601 | 14.30518 | 4.062 | 13.91668 | PCDH11X | X | |
| 158 | ENSG00000102466 | 291.9384 | -1.77141 | 0.309036 | -5.73204 | 9.92E-09 | 3.72E-07 | 9.61797 | 3.974729 | 11.34065 | 1.895475 | FGF14 | | 13 |
| 159 | ENSG00000102760 | 36.61515 | 2.081268 | 0.614409 | 3.387428 | 0.000706 | 0.009203 | 1.852838 | 9.382073 | 2.392363 | 7.611951 | RGCC | | 13 |
| 160 | ENSG00000103056 | 289.259 | -1.11726 | 0.234052 | -4.77358 | 1.81E-06 | 4.48E-05 | 10.45184 | 3.878381 | 9.388402 | 4.767107 | SMPD3 | | 16 |
| 161 | ENSG00000103257 | 650.513 | 1.044408 | 0.180922 | 5.772692 | 7.80E-09 | 2.98E-07 | 12.3373 | 21.9608 | 9.909071 | 21.26331 | SLC7A5 | | 16 |
| 162 | ENSG00000103449 | 684.4587 | -2.69404 | 0.221167 | -12.181 | 3.92E-34 | 1.53E-31 | 34.13621 | 5.349979 | 26.81416 | 3.477004 | SALL1 | | 16 |
| 163 | ENSG00000103485 | 286.2097 | 1.352832 | 0.24623 | 5.494185 | 3.93E-08 | 1.34E-06 | 8.122073 | 23.28393 | 10.73202 | 22.31603 | QPRT | | 16 |
| 164 | ENSG00000103942 | 330.1333 | -1.01073 | 0.243503 | -4.1508 | 3.31E-05 | 0.000621 | 31.02273 | 10.92598 | 21.70071 | 13.77105 | HOMER2 | | 15 |
| 165 | ENSG00000104267 | 269.0145 | -1.1571 | 0.260978 | -4.43369 | 9.26E-06 | 0.000198 | 28.45547 | 15.39238 | 37.17484 | 12.33726 | CA2 | | 8 |
| 166 | ENSG00000104321 | 13.90608 | 5.489603 | 1.429273 | 3.840836 | 0.000123 | 0.00202 | 0.046776 | 2.169195 | 0.021378 | 0.457166 | TRPA1 | | 8 |
| 167 | ENSG00000104327 | 2949.668 | -2.14283 | 0.60619 | -3.53492 | 0.000408 | 0.005771 | 229.6976 | 79.86117 | 319.6381 | 37.0408 | CALB1 | | 8 |
| 168 | ENSG00000104361 | 23.19804 | 2.951289 | 0.797824 | 3.699171 | 0.000216 | 0.003312 | 0.23107 | 1.05266 | 0.178098 | 1.985521 | NIPAL2 | | 8 |
| 169 | ENSG00000104635 | 371.6359 | 1.038033 | 0.209936 | 4.944519 | 7.63E-07 | 2.01E-05 | 6.573132 | 13.14184 | 6.535198 | 12.25857 | SLC39A14 | | 8 |
| 170 | ENSG00000105048 | 115.9563 | 2.000661 | 0.401867 | 4.978415 | 6.41E-07 | 1.72E-05 | 5.897969 | 38.02728 | 13.48942 | 37.16089 | TNNT1 | | 19 |
| 171 | ENSG00000105281 | 420.892 | 1.471585 | 0.202975 | 7.250081 | 4.17E-13 | 2.66E-11 | 9.942614 | 25.49081 | 10.67857 | 28.60842 | SLC1A5 | | 19 |
| 172 | ENSG00000105642 | 187.7047 | -1.25407 | 0.302892 | -4.1403 | 3.47E-05 | 0.000648 | 15.24232 | 5.210725 | 9.799764 | 4.605661 | KCNN1 | | 19 |
| 173 | ENSG00000105792 | 85.11097 | -1.45293 | 0.422452 | -3.43928 | 0.000583 | 0.00786 | 7.220461 | 3.037228 | 6.622905 | 1.743067 | CFAP69 | | 7 |
| 174 | ENSG00000105810 | 3764.182 | 1.730246 | 0.122185 | 14.16088 | 1.60E-45 | 1.04E-42 | 16.87916 | 53.36536 | 18.16544 | 56.55977 | CDK6 | | 7 |
| 175 | ENSG00000105825 | 2577.116 | 1.859847 | 0.398441 | 4.667813 | 3.04E-06 | 7.19E-05 | 59.27128 | 253.6422 | 100.7419 | 297.5294 | TFPI2 | | 7 |
| 176 | ENSG00000105866 | 626.7221 | -1.13936 | 0.175926 | -6.47634 | 9.40E-11 | 4.75E-09 | 20.15898 | 7.912029 | 17.51353 | 8.220719 | SP4 | | 7 |
| 177 | ENSG00000105877 | 322.2924 | -2.37041 | 0.269872 | -8.78345 | 1.59E-18 | 1.73E-16 | 10.15052 | 2.24428 | 15.69801 | 2.50439 | DNAH11 | | 7 |
| 178 | ENSG00000105880 | 24.83176 | -4.39168 | 0.985397 | -4.45676 | 8.32E-06 | 0.000181 | 5.911054 | 0.340552 | 4.595086 | 0.123439 | DLX5 | | 7 |
| 179 | ENSG00000105939 | 216.7262 | 1.380246 | 0.269892 | 5.11406 | 3.15E-07 | 9.16E-06 | 1.943266 | 4.49017 | 2.192184 | 5.725591 | ZC3HAV1 | | 7 |
| 180 | ENSG00000106018 | 100.4855 | -2.93511 | 0.407781 | -7.19777 | 6.12E-13 | 3.82E-11 | 6.838221 | 0.622625 | 5.564838 | 0.908226 | VIPR2 | | 7 |
| 181 | ENSG00000106069 | 575.6537 | -1.98628 | 0.212793 | -9.33432 | 1.02E-20 | 1.34E-18 | 48.37393 | 10.8495 | 34.95489 | 8.888191 | CHN2 | | 7 |
| 182 | ENSG00000106236 | 361.4617 | -1.70275 | 0.215848 | -7.88867 | 3.05E-15 | 2.43E-13 | 26.75853 | 7.871502 | 25.03123 | 7.126214 | NPTX2 | | 7 |
| 183 | ENSG00000106341 | 32.18704 | -3.36463 | 0.753682 | -4.46426 | 8.03E-06 | 0.000175 | 5.714687 | 0.246336 | 2.785704 | 0.536474 | PPP1R17 | | 7 |
| 184 | ENSG00000106366 | 211.8088 | 2.111668 | 0.380387 | 5.551369 | 2.83E-08 | 9.95E-07 | 2.433616 | 16.91559 | 3.454038 | 7.009133 | SERPINE1 | | 7 |
| 185 | ENSG00000106483 | 35.29559 | 3.709629 | 0.696365 | 5.327135 | 9.98E-08 | 3.18E-06 | 0.228885 | 2.986199 | 0.282525 | 3.371021 | SFRP4 | | 7 |
| 186 | ENSG00000106538 | 102.6681 | 3.336497 | 0.424682 | 7.856464 | 3.95E-15 | 3.12E-13 | 3.40375 | 34.73186 | 5.061467 | 46.29803 | RARRES2 | | 7 |
| 187 | ENSG00000106546 | 173.5599 | 1.630342 | 0.413165 | 3.945979 | 7.95E-05 | 0.001367 | 2.504042 | 11.47606 | 2.862432 | 4.128703 | AHR | | 7 |
| 188 | ENSG00000106689 | 182.2746 | -3.17912 | 0.38116 | -8.34065 | 7.39E-17 | 6.72E-15 | 27.76489 | 3.68873 | 19.3979 | 1.518354 | LHX2 | | 9 |
| 189 | ENSG00000106789 | 43.18352 | 1.637042 | 0.568171 | 2.881249 | 0.003961 | 0.038136 | 0.529951 | 1.489711 | 0.344925 | 1.056613 | CORO2A | | 9 |
| 190 | ENSG00000107242 | 343.1935 | 2.077903 | 0.254727 | 8.157379 | 3.42E-16 | 2.94E-14 | 4.820545 | 26.25952 | 7.41468 | 22.66066 | PIP5K1B | | 9 |
| 191 | ENSG00000107295 | 141.39 | -1.18674 | 0.339512 | -3.49544 | 0.000473 | 0.006522 | 10.07366 | 4.461825 | 7.857334 | 2.925438 | SH3GL2 | | 9 |
| 192 | ENSG00000107438 | 80.13861 | 1.447767 | 0.405433 | 3.570915 | 0.000356 | 0.005145 | 4.353809 | 10.36824 | 3.852611 | 10.76206 | PDLIM1 | | 10 |
| 193 | ENSG00000107719 | 114.1977 | 1.042562 | 0.355324 | 2.93412 | 0.003345 | 0.033124 | 2.228092 | 3.57911 | 1.477925 | 3.597799 | PALD1 | | 10 |
| 194 | ENSG00000107731 | 242.2836 | -1.52898 | 0.267659 | -5.71242 | 1.11E-08 | 4.16E-07 | 7.204525 | 2.096086 | 5.135433 | 1.927244 | UNC5B | | 10 |
| 195 | ENSG00000107796 | 1101.587 | 2.384829 | 0.42105 | 5.664001 | 1.48E-08 | 5.41E-07 | 31.82736 | 221.5005 | 40.47375 | 134.0455 | ACTA2 | | 10 |
| 196 | ENSG00000107798 | 674.4066 | -1.06258 | 0.180313 | -5.89298 | 3.79E-09 | 1.53E-07 | 46.93676 | 22.6135 | 44.95372 | 18.83374 | LIPA | | 10 |
| 197 | ENSG00000107954 | 93.53954 | -1.19244 | 0.3842 | -3.10369 | 0.001911 | 0.021222 | 4.568569 | 2.148181 | 4.388727 | 1.553925 | NEURL1 | | 10 |
| 198 | ENSG00000108018 | 1321.558 | -2.78063 | 0.192545 | -14.4414 | 2.84E-47 | 1.91E-44 | 47.87323 | 8.271052 | 51.86571 | 5.397424 | SORCS1 | | 10 |
| 199 | ENSG00000108375 | 88.52376 | 2.103274 | 0.403008 | 5.218935 | 1.80E-07 | 5.46E-06 | 0.921948 | 3.850944 | 0.814526 | 3.161871 | RNF43 | | 17 |
| 200 | ENSG00000108379 | 384.6403 | 4.081383 | 0.585483 | 6.970962 | 3.15E-12 | 1.83E-10 | 3.323308 | 34.5011 | 1.338895 | 39.18179 | WNT3 | | 17 |
| 201 | ENSG00000108439 | 16.56109 | -4.60179 | 1.310818 | -3.51063 | 0.000447 | 0.006231 | 1.251258 | 0.039452 | 1.970257 | 0.115466 | PNPO | | 17 |
| 202 | ENSG00000108691 | 125.015 | 1.847981 | 0.437663 | 4.222388 | 2.42E-05 | 0.000468 | 10.01387 | 56.80359 | 13.45957 | 22.55196 | CCL2 | | 17 |
| 203 | ENSG00000108821 | 1448.018 | 2.902759 | 0.666443 | 4.355603 | 1.33E-05 | 0.000275 | 5.37782 | 74.21064 | 11.0539 | 41.7789 | COL1A1 | | 17 |
| 204 | ENSG00000109072 | 24.00587 | 2.790156 | 0.822646 | 3.391684 | 0.000695 | 0.009092 | 0.635943 | 5.030036 | 0.614712 | 3.106836 | VTN | | 17 |
| 205 | ENSG00000109099 | 155.9741 | 1.811782 | 0.324444 | 5.584271 | 2.35E-08 | 8.34E-07 | 4.351255 | 17.86309 | 4.988057 | 13.07935 | PMP22 | | 17 |
| 206 | ENSG00000109113 | 167.2704 | 1.100423 | 0.295634 | 3.722249 | 0.000197 | 0.003059 | 9.918439 | 23.00821 | 12.34568 | 22.16819 | RAB34 | | 17 |
| 207 | ENSG00000109339 | 6804.042 | 1.163806 | 0.128253 | 9.074273 | 1.14E-19 | 1.43E-17 | 206.747 | 378.8951 | 197.4853 | 478.1196 | MAPK10 | | 4 |
| 208 | ENSG00000109738 | 74.52962 | 1.380911 | 0.426525 | 3.237583 | 0.001205 | 0.01455 | 1.913689 | 4.594404 | 2.385797 | 6.023013 | GLRB | | 4 |
| 209 | ENSG00000109991 | 177.9054 | 1.798685 | 0.358642 | 5.015271 | 5.30E-07 | 1.47E-05 | 2.911214 | 5.35909 | 2.147507 | 11.33399 | P2RX3 | | 11 |
| 210 | ENSG00000110660 | 31.0327 | 2.724365 | 0.724372 | 3.761003 | 0.000169 | 0.002657 | 0.237863 | 2.78262 | 0.461429 | 1.601866 | SLC35F2 | | 11 |
| 211 | ENSG00000110675 | 57.63192 | 2.605117 | 0.520814 | 5.002009 | 5.67E-07 | 1.56E-05 | 1.236477 | 5.33758 | 0.589726 | 5.060355 | ELMOD1 | | 11 |
| 212 | ENSG00000110693 | 1006.115 | -2.15796 | 0.182313 | -11.8366 | 2.52E-32 | 8.23E-30 | 45.92708 | 9.008634 | 34.01882 | 7.834367 | SOX6 | | 11 |
| 213 | ENSG00000110723 | 81.73707 | 1.298312 | 0.407304 | 3.187575 | 0.001435 | 0.016799 | 0.553547 | 1.523947 | 0.638582 | 1.24594 | EXPH5 | | 11 |
| 214 | ENSG00000110786 | 93.87095 | -1.25661 | 0.372257 | -3.37564 | 0.000736 | 0.009505 | 5.789001 | 2.261351 | 5.619386 | 2.245887 | PTPN5 | | 11 |
| 215 | ENSG00000111057 | 241.0956 | 3.707225 | 0.296381 | 12.50829 | 6.73E-36 | 2.99E-33 | 2.981549 | 41.0151 | 3.381887 | 37.66101 | KRT18 | | 12 |
| 216 | ENSG00000111186 | 106.3451 | -1.32359 | 0.361619 | -3.66017 | 0.000252 | 0.003797 | 12.07942 | 3.728206 | 11.4975 | 5.218029 | WNT5B | | 12 |
| 217 | ENSG00000111432 | 390.99 | 5.079077 | 0.706815 | 7.185865 | 6.68E-13 | 4.13E-11 | 0.396015 | 23.58772 | 1.186738 | 27.5565 | FZD10 | | 12 |
| 218 | ENSG00000111783 | 1636.807 | 1.599188 | 0.141527 | 11.29948 | 1.32E-29 | 3.54E-27 | 28.6788 | 72.2789 | 25.39913 | 82.53094 | RFX4 | | 12 |
| 219 | ENSG00000111799 | 234.7223 | 1.152298 | 0.283207 | 4.068742 | 4.73E-05 | 0.00086 | 2.389582 | 5.889727 | 2.34188 | 4.006506 | COL12A1 | | 6 |
| 220 | ENSG00000111834 | 160.0457 | -1.19331 | 0.298083 | -4.00329 | 6.25E-05 | 0.001107 | 10.34694 | 3.824917 | 8.37693 | 3.894752 | RSPH4A | | 6 |
| 221 | ENSG00000111907 | 88.9802 | 1.255555 | 0.444702 | 2.823364 | 0.004752 | 0.043839 | 4.827122 | 16.60771 | 6.254536 | 8.242007 | TPD52L1 | | 6 |
| 222 | ENSG00000111981 | 100.2743 | 1.340583 | 0.369609 | 3.627028 | 0.000287 | 0.004263 | 2.064623 | 5.36543 | 2.069587 | 4.505748 | ULBP1 | | 6 |
| 223 | ENSG00000112139 | 1021.693 | -1.52241 | 0.153693 | -9.9055 | 3.94E-23 | 6.59E-21 | 30.2365 | 9.172844 | 26.7116 | 9.538179 | MDGA1 | | 6 |
| 224 | ENSG00000112164 | 82.63437 | -2.00383 | 0.412172 | -4.86165 | 1.16E-06 | 2.98E-05 | 2.387549 | 0.56654 | 2.052649 | 0.475143 | GLP1R | | 6 |
| 225 | ENSG00000112175 | 84.96437 | 4.590302 | 0.544704 | 8.427143 | 3.54E-17 | 3.38E-15 | 0.193402 | 4.521603 | 0.19879 | 4.408193 | BMP5 | | 6 |
| 226 | ENSG00000112232 | 192.6687 | -1.0626 | 0.290679 | -3.6556 | 0.000257 | 0.003859 | 16.07358 | 5.785763 | 10.96721 | 6.415935 | KHDRBS2 | | 6 |
| 227 | ENSG00000112280 | 145.3681 | 1.19808 | 0.322147 | 3.71905 | 0.0002 | 0.003091 | 3.313797 | 6.796622 | 4.055862 | 9.253382 | COL9A1 | | 6 |
| 228 | ENSG00000112319 | 34.20393 | 1.823866 | 0.625411 | 2.916268 | 0.003542 | 0.034756 | 0.611203 | 2.221095 | 0.572641 | 1.729252 | EYA4 | | 6 |
| 229 | ENSG00000112320 | 621.2732 | -1.28048 | 0.192392 | -6.65558 | 2.82E-11 | 1.50E-09 | 27.48846 | 8.592034 | 24.75358 | 11.78222 | SOBP | | 6 |
| 230 | ENSG00000112333 | 283.2399 | -5.4481 | 0.417121 | -13.0612 | 5.48E-39 | 2.90E-36 | 28.25671 | 0.554341 | 13.58627 | 0.345098 | NR2E1 | | 6 |
| 231 | ENSG00000112419 | 726.2214 | 1.099926 | 0.185024 | 5.944788 | 2.77E-09 | 1.13E-07 | 11.02732 | 26.45546 | 13.87238 | 24.07037 | PHACTR2 | | 6 |
| 232 | ENSG00000112530 | 45.80611 | -1.9706 | 0.553907 | -3.55764 | 0.000374 | 0.005364 | 9.753362 | 2.966224 | 10.13858 | 1.911366 | PACRG | | 6 |
| 233 | ENSG00000112773 | 984.9853 | 2.750227 | 0.168513 | 16.32053 | 7.05E-60 | 6.89E-57 | 4.746951 | 30.66367 | 5.538184 | 34.91732 | TENT5A | | 6 |

| | | | | | | | | | | | | | |
|---|---|---|---|---|---|---|---|---|---|---|---|---|---|
| 234 | ENSG00000112902 | 557.2917 | 2.266763 | 0.187342 | 12.09962 | 1.06E-33 | 3.91E-31 | 2.063234 | 8.895178 | 1.890939 | 9.048499 | SEMA5A | 5 |
| 235 | ENSG00000113083 | 118.3242 | 1.025493 | 0.347062 | 2.954784 | 0.003129 | 0.031263 | 1.952665 | 4.25009 | 1.960419 | 3.247629 | LOX | 5 |
| 236 | ENSG00000113209 | 171.3845 | -1.43729 | 0.287733 | -4.99522 | 5.88E-07 | 1.60E-05 | 9.104946 | 2.92774 | 7.990763 | 3.02962 | PCDHB5 | 5 |
| 237 | ENSG00000113248 | 176.2528 | -1.5862 | 0.290471 | -5.4608 | 4.74E-08 | 1.60E-06 | 8.475173 | 2.4611 | 6.870615 | 2.354054 | PCDHB15 | 5 |
| 238 | ENSG00000113262 | 15.62921 | -3.52654 | 1.020145 | -3.4569 | 0.000546 | 0.00742 | 0.84964 | 0.054147 | 0.54598 | 0.059029 | GRM6 | 5 |
| 239 | ENSG00000113389 | 65.4346 | 4.24815 | 0.582498 | 7.292987 | 3.03E-13 | 1.96E-11 | 0.094813 | 2.933467 | 0.199683 | 2.25117 | NPR3 | 5 |
| 240 | ENSG00000113494 | 66.07093 | -1.61358 | 0.471451 | -3.42258 | 0.00062 | 0.008257 | 1.915158 | 0.422647 | 1.205345 | 0.539908 | PRLR | 5 |
| 241 | ENSG00000113721 | 164.1058 | 1.514482 | 0.377256 | 4.01447 | 5.96E-05 | 0.001059 | 3.429377 | 8.034942 | 1.602019 | 5.094846 | PDGFRB | 5 |
| 242 | ENSG00000113739 | 437.1408 | 2.589011 | 0.24978 | 10.36517 | 3.57E-25 | 7.05E-23 | 3.909531 | 25.56167 | 6.379588 | 33.25881 | STC2 | 5 |
| 243 | ENSG00000113805 | 207.8785 | 3.105713 | 0.308792 | 10.05763 | 8.50E-24 | 1.51E-21 | 1.918495 | 12.77926 | 1.169193 | 11.4912 | CNTN3 | 3 |
| 244 | ENSG00000114200 | 327.8934 | 1.034709 | 0.218881 | 4.727259 | 2.28E-06 | 5.51E-05 | 13.19967 | 23.90432 | 11.59378 | 24.02285 | BCHE | 3 |
| 245 | ENSG00000114279 | 809.1229 | 2.87002 | 0.194696 | 14.74105 | 3.51E-49 | 2.45E-46 | 6.658817 | 54.23506 | 7.301048 | 42.11501 | FGF12 | 3 |
| 246 | ENSG00000114541 | 460.8501 | -1.70601 | 0.245853 | -6.93914 | 3.95E-12 | 2.26E-10 | 43.98071 | 10.09234 | 25.75148 | 9.993005 | FRMD4B | 3 |
| 247 | ENSG00000114654 | 147.588 | 2.221584 | 0.319102 | 6.961987 | 3.36E-12 | 1.94E-10 | 2.509149 | 9.933681 | 2.175026 | 10.62462 | EFCC1 | 3 |
| 248 | ENSG00000114805 | 1324.457 | -1.24359 | 0.140207 | -8.86963 | 7.34E-19 | 8.39E-17 | 35.61858 | 13.5873 | 34.58927 | 14.42623 | PLCH1 | 3 |
| 249 | ENSG00000114859 | 150.8357 | -1.12695 | 0.300879 | -3.74553 | 0.00018 | 0.002814 | 10.41416 | 4.778203 | 10.86383 | 4.427855 | CLCN2 | 3 |
| 250 | ENSG00000115353 | 36.25097 | -2.73518 | 0.69906 | -3.91265 | 9.13E-05 | 0.001544 | 2.274872 | 0.339264 | 1.072896 | 0.134945 | TACR1 | 2 |
| 251 | ENSG00000115380 | 144.578 | 1.061432 | 0.363481 | 2.920182 | 0.003498 | 0.034468 | 7.33154 | 16.5971 | 5.810004 | 9.092022 | EFEMP1 | 2 |
| 252 | ENSG00000115414 | 10881.71 | -2.49105 | 0.336089 | -7.41187 | 1.25E-13 | 8.31E-12 | 381.898 | 75.21589 | 358.564 | 48.63594 | FN1 | 2 |
| 253 | ENSG00000115461 | 1346.477 | 2.041629 | 0.203418 | 10.03663 | 1.05E-23 | 1.84E-21 | 8.012539 | 43.45619 | 11.07555 | 30.72838 | IGFBP5 | 2 |
| 254 | ENSG00000115738 | 703.4973 | 1.557756 | 0.175957 | 8.853053 | 8.52E-19 | 9.68E-17 | 34.40814 | 103.3157 | 40.33324 | 104.8894 | ID2 | 2 |
| 255 | ENSG00000115844 | 59.45719 | -5.94587 | 0.875067 | -6.79476 | 1.08E-11 | 5.99E-10 | 7.83264 | 0.12722 | 3.757562 | 0.046201 | DLX2 | 2 |
| 256 | ENSG00000116016 | 182.7821 | 2.293243 | 0.377026 | 6.082459 | 1.18E-09 | 5.19E-08 | 1.267964 | 7.313466 | 2.929869 | 11.80391 | EPAS1 | 2 |
| 257 | ENSG00000116106 | 2776.5 | -1.16231 | 0.128875 | -9.01888 | 1.90E-19 | 2.34E-17 | 137.4671 | 59.91963 | 134.3179 | 54.63366 | EPHA4 | 2 |
| 258 | ENSG00000116147 | 24.03792 | -4.45061 | 0.988571 | -4.50206 | 6.73E-06 | 0.00015 | 0.229993 | 0.02238 | 0.674075 | 0.01627 | TNR | 1 |
| 259 | ENSG00000116194 | 224.6257 | -2.40576 | 0.273803 | -8.78649 | 1.54E-18 | 1.70E-16 | 13.50045 | 2.752698 | 14.00536 | 2.134091 | ANGPTL1 | 1 |
| 260 | ENSG00000116254 | 82.44067 | -1.45849 | 0.411701 | -3.54261 | 0.000396 | 0.005626 | 1.88149 | 0.624147 | 2.331529 | 0.843687 | CHD5 | 1 |
| 261 | ENSG00000116745 | 56.29127 | 2.348023 | 0.574601 | 4.086356 | 4.38E-05 | 0.000805 | 1.170213 | 5.437045 | 0.538459 | 2.637654 | RPE65 | 1 |
| 262 | ENSG00000116774 | 88.92358 | 2.144915 | 0.46614 | 4.601438 | 4.20E-06 | 9.68E-05 | 1.278295 | 11.11253 | 3.317941 | 8.273422 | OLFML3 | 1 |
| 263 | ENSG00000117069 | 439.2372 | -2.20008 | 0.254454 | -8.6463 | 5.32E-18 | 5.56E-16 | 19.39843 | 3.987427 | 28.73668 | 6.029118 | ST6GALNA | 1 |
| 264 | ENSG00000117266 | 115.9915 | -1.21317 | 0.346317 | -3.50307 | 0.00046 | 0.006374 | 9.341215 | 3.643539 | 7.475324 | 3.167643 | CDK18 | 1 |
| 265 | ENSG00000117298 | 603.1246 | 1.739443 | 0.184184 | 9.444048 | 3.59E-21 | 4.87E-19 | 6.95667 | 23.99654 | 7.819077 | 22.63912 | ECE1 | 1 |
| 266 | ENSG00000117318 | 286.149 | 1.424822 | 0.274256 | 5.195216 | 2.04E-07 | 6.12E-06 | 18.3068 | 61.43001 | 31.28587 | 65.04148 | ID3 | 1 |
| 267 | ENSG00000117394 | 4247.616 | 1.20827 | 0.140257 | 8.614689 | 7.01E-18 | 7.29E-16 | 121.9164 | 306.8675 | 138.4341 | 261.0288 | SLC2A1 | 1 |
| 268 | ENSG00000117643 | 141.8105 | 1.24994 | 0.319074 | 3.9174 | 8.95E-05 | 0.001519 | 2.466605 | 6.390983 | 3.181748 | 6.379395 | MAN1C1 | 1 |
| 269 | ENSG00000117971 | 153.6771 | -1.51515 | 0.313178 | -4.83797 | 1.31E-06 | 3.31E-05 | 10.9505 | 4.324665 | 12.28983 | 3.344899 | CHRNB4 | 15 |
| 270 | ENSG00000118257 | 1712.803 | -1.45586 | 0.19349 | -7.52424 | 5.30E-14 | 3.73E-12 | 69.35527 | 29.9692 | 73.2622 | 18.91062 | NRP2 | 2 |
| 271 | ENSG00000118407 | 185.1373 | -1.39534 | 0.354228 | -3.93911 | 8.18E-05 | 0.001404 | 8.20884 | 2.553785 | 4.019801 | 1.789439 | FILIP1 | 6 |
| 272 | ENSG00000118473 | 641.808 | -1.36244 | 0.176727 | -7.7093 | 1.27E-14 | 9.51E-13 | 29.23057 | 9.775234 | 24.92457 | 10.09666 | SGIP1 | 1 |
| 273 | ENSG00000118508 | 68.82219 | 2.025994 | 0.450135 | 4.500861 | 6.77E-06 | 0.00015 | 3.807832 | 14.57613 | 3.376058 | 12.97916 | RAB32 | 6 |
| 274 | ENSG00000118523 | 2108.337 | 1.668212 | 0.175946 | 9.481413 | 2.51E-21 | 3.43E-19 | 51.28876 | 183.7556 | 52.89213 | 127.9611 | CCN2 | 6 |
| 275 | ENSG00000118733 | 187.4363 | 1.113548 | 0.297199 | 3.746816 | 0.000179 | 0.002804 | 5.112215 | 11.98339 | 4.936386 | 8.465752 | OLFM3 | 1 |
| 276 | ENSG00000118898 | 90.26697 | 1.256581 | 0.442929 | 2.836981 | 0.004554 | 0.042548 | 1.649287 | 4.675156 | 1.483251 | 2.357432 | PPL | 16 |
| 277 | ENSG00000119638 | 833.9085 | -1.01604 | 0.16904 | -6.01067 | 1.85E-09 | 7.89E-08 | 31.29589 | 16.07404 | 32.65478 | 13.78234 | NEK9 | 14 |
| 278 | ENSG00000119699 | 152.4113 | 1.834968 | 0.344571 | 5.325364 | 1.01E-07 | 3.21E-06 | 1.994845 | 9.941586 | 3.283133 | 7.777582 | TGFB3 | 14 |
| 279 | ENSG00000120149 | 169.5353 | 5.708729 | 0.502531 | 11.35995 | 6.62E-30 | 1.82E-27 | 0.184733 | 18.1204 | 0.572766 | 19.08758 | MSX2 | 5 |
| 280 | ENSG00000120156 | 75.78064 | 1.869184 | 0.452253 | 4.133052 | 3.58E-05 | 0.000667 | 0.705975 | 3.559595 | 1.144258 | 2.782791 | TEK | 9 |
| 281 | ENSG00000120162 | 581.507 | -1.64911 | 0.219449 | -7.51475 | 5.70E-14 | 3.98E-12 | 20.23507 | 4.803785 | 12.92621 | 5.160276 | MOB3B | 9 |
| 282 | ENSG00000120251 | 425.208 | -1.28075 | 0.201815 | -6.34617 | 2.21E-10 | 1.06E-08 | 27.56682 | 10.66615 | 30.22313 | 11.85428 | GRIA2 | 4 |
| 283 | ENSG00000120322 | 70.46834 | -1.3578 | 0.449472 | -3.02087 | 0.00252 | 0.026432 | 4.697485 | 1.981335 | 4.057648 | 1.222013 | PCDHB8 | 5 |
| 284 | ENSG00000120708 | 44.43612 | 2.763672 | 0.65239 | 4.236229 | 2.27E-05 | 0.000444 | 0.237388 | 4.666489 | 0.887406 | 2.961195 | TGFBI | 5 |
| 285 | ENSG00000120937 | 42.27079 | 4.100026 | 0.742993 | 5.518255 | 3.42E-08 | 1.18E-06 | 0.778254 | 23.71178 | 1.38863 | 11.53097 | NPPB | 1 |
| 286 | ENSG00000121039 | 198.2313 | 1.480778 | 0.349953 | 4.231367 | 2.32E-05 | 0.000451 | 6.310003 | 20.31398 | 5.195547 | 9.86445 | RDH10 | 8 |
| 287 | ENSG00000121207 | 160.9876 | -1.18142 | 0.30666 | -3.85253 | 0.000117 | 0.001937 | 4.571245 | 2.3275 | 5.604462 | 1.90778 | LRAT | 4 |
| 288 | ENSG00000121743 | 19.68029 | 4.173186 | 1.038927 | 4.016824 | 5.90E-05 | 0.001051 | 0.024164 | 0.771764 | 0.063838 | 0.75926 | GJA3 | 13 |
| 289 | ENSG00000121871 | 170.9905 | -1.3718 | 0.293403 | -4.67548 | 2.93E-06 | 6.97E-05 | 7.915897 | 2.717601 | 6.40033 | 2.488941 | SLITRK3 | 3 |
| 290 | ENSG00000121966 | 1041.954 | 3.033918 | 0.191249 | 15.86368 | 1.13E-56 | 1.05E-53 | 13.81974 | 137.7462 | 20.26187 | 127.0406 | CXCR4 | 2 |
| 291 | ENSG00000121989 | 1635.356 | -1.65468 | 0.16834 | -9.82942 | 8.41E-23 | 1.32E-20 | 104.4457 | 26.66687 | 73.60235 | 26.61445 | ACVR2A | 2 |
| 292 | ENSG00000122584 | 34.82953 | -4.03099 | 0.742745 | -5.42715 | 5.73E-08 | 1.90E-06 | 3.16529 | 0.231208 | 3.918936 | 0.178429 | NXPH1 | 7 |
| 293 | ENSG00000122735 | 56.97085 | -2.75283 | 0.512894 | -5.36725 | 7.99E-08 | 2.61E-06 | 10.1127 | 1.374233 | 14.18022 | 2.082674 | DNAI1 | 9 |
| 294 | ENSG00000122861 | 111.3229 | 2.060412 | 0.422708 | 4.874312 | 1.09E-06 | 2.82E-05 | 2.538557 | 11.99833 | 2.071999 | 5.935206 | PLAU | 10 |
| 295 | ENSG00000123104 | 517.5691 | 2.142213 | 0.521245 | 4.109802 | 3.96E-05 | 0.000732 | 2.151695 | 14.70317 | 4.35864 | 12.44832 | ITPR2 | 12 |
| 296 | ENSG00000123146 | 20.23097 | 2.344219 | 0.838393 | 2.796087 | 0.005173 | 0.046848 | 0.213728 | 1.774098 | 0.376817 | 1.067404 | ADGRE5 | 19 |
| 297 | ENSG00000123243 | 54.55283 | 1.42922 | 0.494037 | 2.892939 | 0.003817 | 0.03706 | 0.746363 | 2.061923 | 0.672279 | 1.527314 | ITIH5 | 10 |
| 298 | ENSG00000123496 | 46.2751 | 3.797968 | 0.69632 | 5.454344 | 4.92E-08 | 1.65E-06 | 0.109868 | 7.046212 | 1.069339 | 8.979065 | IL13RA2 | X |
| 299 | ENSG00000124126 | 1070.152 | 3.207319 | 0.163542 | 19.61159 | 1.23E-85 | 3.44E-82 | 3.874098 | 34.2657 | 4.31316 | 37.37494 | PREX1 | 20 |
| 300 | ENSG00000124145 | 118.8735 | 1.096366 | 0.341959 | 3.206129 | 0.001345 | 0.015886 | 3.14383 | 7.396232 | 3.756444 | 6.550277 | SDC4 | 20 |
| 301 | ENSG00000124479 | 15.75841 | -3.10159 | 0.984702 | -3.14977 | 0.001634 | 0.018695 | 1.873707 | 0.26466 | 2.425864 | 0.207882 | NDP | X |
| 302 | ENSG00000124813 | 29.99707 | 2.271079 | 0.728362 | 3.118064 | 0.00182 | 0.020446 | 0.379241 | 1.712955 | 0.213269 | 0.854171 | RUNX2 | 6 |
| 303 | ENSG00000125246 | 357.5291 | 2.096046 | 0.233454 | 8.978397 | 2.75E-19 | 3.26E-17 | 6.223671 | 21.23163 | 5.921228 | 27.95096 | CLYBL | 13 |
| 304 | ENSG00000125285 | 1626.802 | -1.02924 | 0.134686 | -7.6418 | 2.14E-14 | 1.59E-12 | 88.42991 | 38.96941 | 87.64412 | 42.5786 | SOX21 | 13 |
| 305 | ENSG00000125378 | 50.49506 | 3.095972 | 0.619283 | 4.999281 | 5.75E-07 | 1.58E-05 | 0.657974 | 9.450684 | 1.541818 | 6.709334 | BMP4 | 14 |
| 306 | ENSG00000125462 | 203.5629 | -2.57041 | 0.331347 | -7.75745 | 8.67E-15 | 6.64E-13 | 24.97884 | 4.404568 | 17.75644 | 2.383404 | MIR9-1HG | 1 |
| 307 | ENSG00000125492 | 24.72094 | 3.898001 | 0.889353 | 4.382964 | 1.17E-05 | 0.000245 | 0.144586 | 3.04823 | 0.255331 | 2.643464 | BARHL1 | 9 |
| 308 | ENSG00000125675 | 537.7523 | -1.15206 | 0.193292 | -5.96019 | 2.52E-09 | 1.04E-07 | 32.12004 | 13.66794 | 27.76939 | 11.72787 | GRIA3 | X |
| 309 | ENSG00000125804 | 112.2748 | -2.17908 | 0.421243 | -5.17298 | 2.30E-07 | 6.87E-06 | 11.50827 | 1.612927 | 5.223017 | 1.861116 | FAM182A | 20 |
| 310 | ENSG00000125851 | 235.756 | -1.77198 | 0.316738 | -5.59448 | 2.21E-08 | 7.88E-07 | 15.76733 | 2.662818 | 9.067433 | 4.249406 | PCSK2 | 20 |
| 311 | ENSG00000125968 | 460.2626 | 1.715493 | 0.209598 | 8.184701 | 2.73E-16 | 2.38E-14 | 25.72521 | 88.55009 | 33.35966 | 95.45334 | ID1 | 20 |

| | | | | | | | | | | | | | | |
|---|---|---|---|---|---|---|---|---|---|---|---|---|---|---|
| 312 | ENSG00000126259 | 165.2977 | 1.487581 | 0.323337 | 4.600714 | 4.21E-06 | 9.70E-05 | 2.758686 | 10.25167 | 4.264033 | 8.405867 | KIRREL2 | | 19 |
| 313 | ENSG00000126583 | 154.37 | 1.741009 | 0.366557 | 4.74963 | 2.04E-06 | 4.99E-05 | 2.527448 | 8.781435 | 4.691504 | 13.938 | PRKCG | | 19 |
| 314 | ENSG00000126950 | 433.0452 | -1.07151 | 0.199761 | -5.36395 | 8.14E-08 | 2.64E-06 | 38.40897 | 15.36141 | 33.00819 | 16.72304 | TMEM35A | X | |
| 315 | ENSG00000127152 | 116.1171 | -1.08928 | 0.381353 | -2.85635 | 0.004285 | 0.040687 | 2.720731 | 1.073018 | 1.55507 | 0.810588 | BCL11B | | 14 |
| 316 | ENSG00000127241 | 213.9461 | -1.20181 | 0.276723 | -4.343 | 1.41E-05 | 0.000289 | 11.27294 | 5.610285 | 14.73259 | 5.053616 | MASP1 | | 3 |
| 317 | ENSG00000127399 | 25.50158 | -3.75849 | 0.847976 | -4.43231 | 9.32E-06 | 0.000199 | 3.594727 | 0.195582 | 3.242336 | 0.283934 | LRRC61 | | 7 |
| 318 | ENSG00000127586 | 164.645 | 1.052124 | 0.321576 | 3.271779 | 0.001069 | 0.013119 | 4.863146 | 9.577183 | 6.969974 | 13.80416 | CHTF18 | | 16 |
| 319 | ENSG00000127920 | 1538.036 | 1.947658 | 0.197011 | 9.88605 | 4.79E-23 | 7.93E-21 | 18.82897 | 82.21649 | 30.29773 | 97.75774 | GNG11 | | 7 |
| 320 | ENSG00000128045 | 255.2852 | 1.390961 | 0.243535 | 5.711555 | 1.12E-08 | 4.17E-07 | 9.905141 | 24.28268 | 10.44012 | 26.17261 | RASL11B | | 4 |
| 321 | ENSG00000128165 | 58.55106 | 2.226904 | 0.507004 | 4.39228 | 1.12E-05 | 0.000236 | 0.393374 | 2.439823 | 0.708458 | 2.479379 | ADM2 | | 22 |
| 322 | ENSG00000128573 | 355.2321 | -1.88549 | 0.227467 | -8.2891 | 1.14E-16 | 1.03E-14 | 17.24381 | 3.938965 | 13.43365 | 3.886683 | FOXP2 | | 7 |
| 323 | ENSG00000128606 | 332.9851 | 1.139281 | 0.226662 | 5.026352 | 5.00E-07 | 1.39E-05 | 13.85253 | 31.09927 | 13.66429 | 26.05226 | LRRC17 | | 7 |
| 324 | ENSG00000128610 | 1296.416 | -4.48185 | 0.195397 | -22.9372 | 1.98E-116 | 9.67E-113 | 165.027 | 8.899755 | 194.3206 | 6.15263 | FEZF1 | | 7 |
| 325 | ENSG00000128656 | 879.7553 | -1.18455 | 0.155824 | -7.60186 | 2.92E-14 | 2.13E-12 | 126.9692 | 52.13915 | 125.9582 | 52.96426 | CHN1 | | 2 |
| 326 | ENSG00000128965 | 62.14392 | 1.869881 | 0.469161 | 3.985587 | 6.73E-05 | 0.00118 | 2.586717 | 7.495237 | 2.071373 | 8.574901 | CHAC1 | | 15 |
| 327 | ENSG00000130176 | 125.1999 | 1.476461 | 0.367489 | 4.017705 | 5.88E-05 | 0.001048 | 8.563457 | 20.19632 | 5.147353 | 15.13758 | CNN1 | | 19 |
| 328 | ENSG00000130226 | 581.2861 | -1.55411 | 0.209341 | -7.42382 | 1.14E-13 | 7.67E-12 | 33.76243 | 10.25451 | 24.6251 | 8.436531 | DPP6 | | 7 |
| 329 | ENSG00000130635 | 566.6709 | 1.117335 | 0.180212 | 6.200125 | 5.64E-10 | 2.60E-08 | 5.780873 | 11.40048 | 5.254702 | 11.1764 | COL5A1 | | 9 |
| 330 | ENSG00000130830 | 62.55598 | 2.652202 | 0.508716 | 5.213527 | 1.85E-07 | 5.60E-06 | 1.553722 | 11.11163 | 2.097743 | 10.44887 | MPP1 | X | |
| 331 | ENSG00000130940 | 411.364 | 1.827262 | 0.242969 | 7.520541 | 5.45E-14 | 3.82E-12 | 5.631173 | 19.32211 | 7.994781 | 26.66795 | CASZ1 | | 1 |
| 332 | ENSG00000131019 | 424.7771 | 1.659388 | 0.234457 | 7.077574 | 1.47E-12 | 8.80E-11 | 10.85921 | 26.94855 | 6.832149 | 25.13401 | ULBP3 | | 6 |
| 333 | ENSG00000131378 | 428.5079 | -1.88927 | 0.236668 | -7.98277 | 1.43E-15 | 1.18E-13 | 25.822 | 8.551904 | 30.90553 | 5.854896 | RFTN1 | | 3 |
| 334 | ENSG00000131620 | 11.89075 | -3.66929 | 1.148535 | -3.19476 | 0.001399 | 0.016436 | 1.130958 | 0.118556 | 1.255674 | 0.061689 | ANO1 | | 11 |
| 335 | ENSG00000131721 | 26.52815 | -2.10079 | 0.709463 | -2.9611 | 0.003065 | 0.030802 | 1.614506 | 0.431676 | 1.952241 | 0.352783 | RHOXF2 | X | |
| 336 | ENSG00000131771 | 31.33104 | -2.36659 | 0.672292 | -3.52018 | 0.000431 | 0.006036 | 4.597762 | 0.721205 | 3.738818 | 0.810056 | PPP1R1B | | 17 |
| 337 | ENSG00000131914 | 371.7984 | 1.094507 | 0.224845 | 4.86782 | 1.13E-06 | 2.90E-05 | 8.782335 | 13.93381 | 6.817328 | 17.52707 | LIN28A | | 1 |
| 338 | ENSG00000132561 | 305.8067 | 3.266991 | 0.288582 | 11.32084 | 1.03E-29 | 2.81E-27 | 2.412514 | 26.70573 | 2.493903 | 17.74224 | MATN2 | | 8 |
| 339 | ENSG00000132718 | 2978.044 | -1.36406 | 0.128891 | -10.5831 | 3.57E-26 | 7.35E-24 | 100.8801 | 34.57585 | 86.84719 | 34.22307 | SYT11 | | 1 |
| 340 | ENSG00000132849 | 782.7347 | 1.456641 | 0.171904 | 8.473581 | 2.38E-17 | 2.34E-15 | 13.18203 | 31.50344 | 13.65287 | 38.25945 | PATJ | | 1 |
| 341 | ENSG00000133101 | 41.92835 | -2.16866 | 0.566608 | -3.82744 | 0.000129 | 0.002119 | 5.062576 | 1.01066 | 4.978311 | 1.10047 | CCNA1 | | 13 |
| 342 | ENSG00000133124 | 4216.114 | -3.04756 | 0.17271 | -17.6455 | 1.10E-69 | 1.35E-66 | 64.96174 | 9.823631 | 87.44079 | 7.573964 | IRS4 | X | |
| 343 | ENSG00000133142 | 579.4313 | -1.00433 | 0.181144 | -5.54436 | 2.95E-08 | 1.03E-06 | 89.67841 | 39.08228 | 91.42713 | 46.37814 | TCEAL4 | X | |
| 344 | ENSG00000133710 | 68.98848 | -1.665 | 0.43759 | -3.80492 | 0.000142 | 0.002283 | 3.851372 | 1.042824 | 3.614939 | 1.184735 | SPINK5 | | 5 |
| 345 | ENSG00000133816 | 36.80245 | -1.82394 | 0.608967 | -2.99513 | 0.002743 | 0.028158 | 1.477154 | 0.587478 | 2.250456 | 0.403051 | MICAL2 | | 11 |
| 346 | ENSG00000134222 | 427.4925 | 1.28434 | 0.200636 | 6.401332 | 1.54E-10 | 7.53E-09 | 22.52716 | 50.02071 | 20.15534 | 47.95364 | PSRC1 | | 1 |
| 347 | ENSG00000134245 | 2195.21 | 2.204798 | 0.158301 | 13.92789 | 4.29E-44 | 2.54E-41 | 11.72225 | 51.0213 | 9.630933 | 41.59823 | WNT2B | | 1 |
| 348 | ENSG00000134330 | 67.02062 | -1.22232 | 0.439579 | -2.78065 | 0.005425 | 0.048728 | 5.778353 | 1.969835 | 5.302689 | 2.536274 | IAH1 | | 2 |
| 349 | ENSG00000134376 | 195.851 | -1.05472 | 0.312356 | -3.37665 | 0.000734 | 0.009483 | 10.2418 | 3.27631 | 6.066076 | 4.149567 | CRB1 | | 1 |
| 350 | ENSG00000134504 | 632.7485 | 1.466875 | 0.215752 | 6.798881 | 1.05E-11 | 5.84E-10 | 12.95701 | 38.70941 | 19.62429 | 46.58693 | KCTD1 | | 18 |
| 351 | ENSG00000134508 | 99.08039 | 1.369417 | 0.374683 | 3.654862 | 0.000257 | 0.003865 | 2.577988 | 6.698151 | 2.346526 | 5.273544 | CABLES1 | | 18 |
| 352 | ENSG00000134531 | 113.4914 | 1.738076 | 0.439766 | 3.952273 | 7.74E-05 | 0.001338 | 1.602723 | 7.408745 | 1.757198 | 3.15244 | EMP1 | | 12 |
| 353 | ENSG00000134548 | 35.4564 | 3.092173 | 0.737326 | 4.193764 | 2.74E-05 | 0.000525 | 0.295026 | 4.867544 | 0.560183 | 1.974183 | SPX | | 12 |
| 354 | ENSG00000134569 | 989.8122 | 1.373649 | 0.155808 | 8.816283 | 1.18E-18 | 1.32E-16 | 12.29138 | 29.01441 | 10.98413 | 27.85724 | LRP4 | | 11 |
| 355 | ENSG00000134595 | 624.6205 | -1.68985 | 0.222894 | -7.58138 | 3.42E-14 | 2.46E-12 | 69.1095 | 17.37025 | 43.25746 | 15.36903 | SOX3 | X | |
| 356 | ENSG00000134709 | 203.602 | 1.069201 | 0.280338 | 3.813972 | 0.000137 | 0.002216 | 3.008831 | 5.487546 | 3.500993 | 7.521534 | HOOK1 | | 1 |
| 357 | ENSG00000134769 | 469.1126 | -1.41524 | 0.215018 | -6.58195 | 4.64E-11 | 2.42E-09 | 25.03064 | 9.687268 | 22.68036 | 7.129354 | DTNA | | 18 |
| 358 | ENSG00000134817 | 13.22246 | 5.586164 | 1.686175 | 3.312921 | 0.000923 | 0.011565 | 0 | 1.308613 | 0.031452 | 0.4245 | APLNR | | 11 |
| 359 | ENSG00000134954 | 154.8867 | 2.608666 | 0.356997 | 7.30725 | 2.73E-13 | 1.77E-11 | 0.962381 | 8.786068 | 1.727678 | 6.811712 | ETS1 | | 11 |
| 360 | ENSG00000135046 | 553.3145 | 3.052272 | 0.257686 | 11.84495 | 2.29E-32 | 7.58E-30 | 9.253353 | 108.6195 | 13.4816 | 69.74415 | ANXA1 | | 9 |
| 361 | ENSG00000135074 | 759.8524 | 1.546046 | 0.19557 | 7.905327 | 2.67E-15 | 2.14E-13 | 6.024128 | 20.44794 | 8.582356 | 20.00166 | ADAM19 | | 5 |
| 362 | ENSG00000135269 | 94.41473 | 1.831 | 0.384835 | 4.757884 | 1.96E-06 | 4.81E-05 | 2.398777 | 7.516286 | 2.362172 | 8.500902 | TES | | 7 |
| 363 | ENSG00000135299 | 1092.142 | -1.28675 | 0.160295 | -8.02739 | 9.96E-16 | 8.32E-14 | 71.76404 | 23.82166 | 57.87102 | 26.35935 | ANKRD6 | | 6 |
| 364 | ENSG00000135318 | 39.01828 | 1.937268 | 0.585868 | 3.306662 | 0.000944 | 0.011797 | 0.651577 | 2.616476 | 0.606079 | 1.904478 | NT5E | | 6 |
| 365 | ENSG00000135363 | 65.55142 | -4.07938 | 0.571017 | -7.14407 | 9.06E-13 | 5.55E-11 | 8.567852 | 0.663099 | 11.98857 | 0.481214 | LMO2 | | 11 |
| 366 | ENSG00000135525 | 257.5366 | -1.40569 | 0.247008 | -5.69088 | 1.26E-08 | 4.66E-07 | 12.70076 | 5.00528 | 13.58395 | 4.359667 | MAP7 | | 6 |
| 367 | ENSG00000135842 | 183.6671 | 2.947244 | 0.304801 | 9.669417 | 4.07E-22 | 5.89E-20 | 0.795748 | 5.604912 | 0.69866 | 5.251169 | NIBAN1 | | 1 |
| 368 | ENSG00000135903 | 780.0341 | 2.484371 | 0.233957 | 10.61894 | 2.43E-26 | 5.06E-24 | 8.280419 | 59.85174 | 15.47826 | 66.54808 | PAX3 | | 2 |
| 369 | ENSG00000135905 | 253.88 | 1.348251 | 0.245353 | 5.49515 | 3.90E-08 | 1.34E-06 | 3.436863 | 7.790947 | 3.630391 | 9.262796 | DOCK10 | | 2 |
| 370 | ENSG00000136010 | 230.607 | 1.113588 | 0.301312 | 3.695793 | 0.000219 | 0.003351 | 2.609015 | 7.61401 | 3.526654 | 4.892103 | ALDH1L2 | | 12 |
| 371 | ENSG00000136014 | 378.1458 | -1.28003 | 0.21643 | -5.9143 | 3.33E-09 | 1.35E-07 | 22.605 | 8.405235 | 25.53392 | 10.40547 | USP44 | | 12 |
| 372 | ENSG00000136158 | 759.9012 | 1.055625 | 0.228484 | 4.620119 | 3.84E-06 | 8.92E-05 | 18.55827 | 51.18165 | 32.58536 | 49.63898 | SPRY2 | | 13 |
| 373 | ENSG00000136160 | 1176.291 | 1.664499 | 0.171868 | 9.68476 | 3.50E-22 | 5.15E-20 | 17.17379 | 51.12573 | 13.98073 | 41.80728 | EDNRB | | 13 |
| 374 | ENSG00000136267 | 111.275 | -1.60495 | 0.376566 | -4.26207 | 2.03E-05 | 0.0004 | 9.959392 | 3.075753 | 7.017124 | 2.179163 | DGKB | | 7 |
| 375 | ENSG00000136531 | 577.1606 | -1.19469 | 0.181173 | -6.5942 | 4.28E-11 | 2.24E-09 | 11.05467 | 4.703533 | 10.76902 | 4.293799 | SCN2A | | 2 |
| 376 | ENSG00000136535 | 943.8136 | -4.4724 | 0.573068 | -7.8043 | 5.98E-15 | 4.62E-13 | 53.67955 | 3.043467 | 50.40273 | 1.356291 | TBR1 | | 2 |
| 377 | ENSG00000136732 | 143.0089 | 2.442569 | 0.342861 | 7.124085 | 1.05E-12 | 6.38E-11 | 5.349844 | 26.1896 | 6.672567 | 35.95473 | GYPC | | 2 |
| 378 | ENSG00000136750 | 77.53499 | -3.22557 | 0.530705 | -6.0779 | 1.22E-09 | 5.30E-08 | 8.659809 | 0.515167 | 4.394617 | 0.841618 | GAD2 | | 10 |
| 379 | ENSG00000136770 | 380.2244 | 1.297795 | 0.207187 | 6.263867 | 3.76E-10 | 1.76E-08 | 13.36477 | 28.94247 | 12.7507 | 31.72675 | DNAJC1 | | 10 |
| 380 | ENSG00000136826 | 76.55105 | 1.567094 | 0.433292 | 3.616716 | 0.000298 | 0.004423 | 1.868594 | 6.447527 | 2.134271 | 4.774783 | KLF4 | | 9 |
| 381 | ENSG00000136943 | 328.7019 | 3.014442 | 0.260358 | 11.57807 | 5.32E-31 | 1.58E-28 | 4.560879 | 45.0017 | 6.322132 | 37.97756 | CTSV | | 9 |
| 382 | ENSG00000136944 | 238.5436 | 1.757521 | 0.26143 | 6.722709 | 1.78E-11 | 9.64E-10 | 1.706031 | 5.998493 | 2.168061 | 6.420842 | LMX1B | | 9 |
| 383 | ENSG00000136960 | 416.9514 | 1.117061 | 0.206207 | 5.417189 | 6.05E-08 | 2.00E-06 | 12.60866 | 22.01736 | 10.42331 | 25.11492 | ENPP2 | | 8 |
| 384 | ENSG00000137142 | 9690.476 | 1.581961 | 0.112515 | 14.05994 | 6.69E-45 | 4.22E-42 | 158.6486 | 419.5934 | 156.9125 | 473.6164 | IGFBPL1 | | 9 |
| 385 | ENSG00000137193 | 178.1727 | 1.197236 | 0.282194 | 4.242601 | 2.21E-05 | 0.000432 | 7.32306 | 14.64333 | 6.380492 | 15.03691 | PIM1 | | 6 |
| 386 | ENSG00000137203 | 163.636 | -2.70699 | 0.331856 | -8.15712 | 3.43E-16 | 2.94E-14 | 14.83057 | 2.21056 | 20.1342 | 2.920554 | TFAP2A | | 6 |
| 387 | ENSG00000137261 | 52.66906 | -1.47851 | 0.509971 | -2.8992 | 0.003741 | 0.03645 | 1.600803 | 0.469231 | 1.023922 | 0.415798 | KIAA0319 | | 6 |
| 388 | ENSG00000137273 | 50.71397 | 2.904181 | 0.576259 | 5.039717 | 4.66E-07 | 1.31E-05 | 0.54431 | 5.095748 | 0.624179 | 3.145929 | FOXF2 | | 6 |
| 389 | ENSG00000137331 | 237.0232 | 2.245256 | 0.297386 | 7.549963 | 4.35E-14 | 3.10E-12 | 8.430945 | 48.33841 | 9.436893 | 31.45534 | IER3 | | 6 |

| | | | | | | | | | | | | | |
|---|---|---|---|---|---|---|---|---|---|---|---|---|---|
| 390 | ENSG00000137440 | 11.46984 | 6.901374 | 1.75434 | 3.933887 | 8.36E-05 | 0.001431 | 0 | 2.293128 | 0 | 1.940906 | FGFBP1 | 4 |
| 391 | ENSG00000137502 | 528.0262 | -1.10308 | 0.226952 | -4.8604 | 1.17E-06 | 3.00E-05 | 18.07424 | 7.785647 | 12.82589 | 5.718058 | RAB30 | 11 |
| 392 | ENSG00000137573 | 262.8328 | -1.69128 | 0.30135 | -5.61234 | 2.00E-08 | 7.14E-07 | 18.23043 | 5.091653 | 11.0187 | 3.389673 | SULF1 | 8 |
| 393 | ENSG00000137727 | 36.20683 | 1.816129 | 0.647572 | 2.804522 | 0.005039 | 0.045852 | 0.340921 | 1.344339 | 0.262656 | 0.641519 | ARHGAP2( | 11 |
| 394 | ENSG00000137871 | 186.6134 | -1.75075 | 0.344415 | -5.08326 | 3.71E-07 | 1.06E-05 | 15.59257 | 2.653472 | 14.00754 | 6.015849 | ZNF280D | 15 |
| 395 | ENSG00000137962 | 735.1888 | 1.995737 | 0.189844 | 10.5125 | 7.57E-26 | 1.53E-23 | 4.61516 | 17.49329 | 5.769666 | 21.82951 | ARHGAP2! | 1 |
| 396 | ENSG00000138061 | 111.9187 | 3.034337 | 0.405601 | 7.481086 | 7.37E-14 | 5.08E-12 | 0.386306 | 4.527848 | 0.680386 | 3.790945 | CYP1B1 | 2 |
| 397 | ENSG00000138083 | 300.7471 | -3.06157 | 0.280971 | -10.8964 | 1.20E-27 | 2.73E-25 | 25.10453 | 3.377151 | 21.96612 | 1.901884 | SIX3 | 2 |
| 398 | ENSG00000138119 | 331.3752 | 1.202495 | 0.264297 | 4.549784 | 5.37E-06 | 0.000121 | 3.622464 | 11.12147 | 5.186409 | 8.006877 | MYOF | 10 |
| 399 | ENSG00000138166 | 55.06446 | 1.453748 | 0.508466 | 2.859085 | 0.004249 | 0.040447 | 1.775297 | 4.238825 | 1.091414 | 3.109225 | DUSP5 | 10 |
| 400 | ENSG00000138411 | 851.9723 | -1.8312 | 0.177726 | -10.3035 | 6.80E-25 | 1.33E-22 | 25.53768 | 6.758438 | 21.32978 | 5.636216 | HECW2 | 2 |
| 401 | ENSG00000138653 | 478.0662 | -1.3795 | 0.205976 | -6.69739 | 2.12E-11 | 1.13E-09 | 28.90654 | 10.67817 | 24.67366 | 8.696553 | NDST4 | 4 |
| 402 | ENSG00000138670 | 154.2214 | -2.13605 | 0.357319 | -5.97798 | 2.26E-09 | 9.46E-08 | 12.65774 | 3.625774 | 11.92343 | 1.825574 | RASGEF1B | 4 |
| 403 | ENSG00000138696 | 546.8409 | 1.300226 | 0.216496 | 6.005764 | 1.90E-09 | 8.08E-08 | 9.472026 | 16.10304 | 6.629659 | 21.336 | BMPR1B | 4 |
| 404 | ENSG00000138735 | 398.3393 | 1.122827 | 0.207659 | 5.407071 | 6.41E-08 | 2.11E-06 | 10.24319 | 18.66565 | 8.466483 | 19.76217 | PDE5A | 4 |
| 405 | ENSG00000138759 | 2937.767 | -1.08791 | 0.165716 | -6.56488 | 5.21E-11 | 2.69E-09 | 41.84202 | 20.27317 | 36.82749 | 14.52655 | FRAS1 | 4 |
| 406 | ENSG00000138772 | 75.29116 | 2.678277 | 0.499772 | 5.359 | 8.37E-08 | 2.71E-06 | 2.710822 | 16.77578 | 1.748049 | 9.509789 | ANXA3 | 4 |
| 407 | ENSG00000138798 | 605.8147 | -1.01793 | 0.176958 | -5.7524 | 8.80E-09 | 3.33E-07 | 35.31756 | 17.13437 | 35.83498 | 16.01715 | EGF | 4 |
| 408 | ENSG00000138821 | 82.59485 | 1.311411 | 0.40718 | 3.220718 | 0.001279 | 0.015237 | 2.30681 | 4.608988 | 1.60169 | 4.534121 | SLC39A8 | 4 |
| 409 | ENSG00000139132 | 1092.553 | -1.05838 | 0.153042 | -6.91561 | 4.66E-12 | 2.65E-10 | 45.73166 | 21.47518 | 51.16615 | 22.52291 | FGD4 | 12 |
| 410 | ENSG00000139219 | 945.2706 | 1.088749 | 0.173801 | 6.264338 | 3.74E-10 | 1.76E-08 | 14.67107 | 31.28969 | 18.63403 | 35.87322 | COL2A1 | 12 |
| 411 | ENSG00000139364 | 567.4585 | -1.05557 | 0.200128 | -5.27448 | 1.33E-07 | 4.12E-06 | 11.15987 | 5.325769 | 9.696866 | 4.123178 | TMEM132I | 12 |
| 412 | ENSG00000139514 | 1348.528 | 1.447618 | 0.16671 | 8.683438 | 3.84E-18 | 4.08E-16 | 13.03989 | 30.14181 | 9.589486 | 27.93092 | SLC7A1 | 13 |
| 413 | ENSG00000139722 | 511.3147 | -1.53439 | 0.195346 | -7.85473 | 4.01E-15 | 3.15E-13 | 38.25287 | 11.39209 | 31.16061 | 11.19869 | VPS37B | 12 |
| 414 | ENSG00000139800 | 817.9754 | 1.117416 | 0.182784 | 6.113324 | 9.76E-10 | 4.37E-08 | 10.95763 | 26.74043 | 14.3446 | 25.20391 | ZIC5 | 13 |
| 415 | ENSG00000140015 | 196.0354 | -2.44665 | 0.390345 | -6.26793 | 3.66E-10 | 1.72E-08 | 8.788416 | 1.681683 | 4.827409 | 0.691213 | KCNH5 | 14 |
| 416 | ENSG00000140285 | 21.98424 | -2.54209 | 0.844266 | -3.01101 | 0.002604 | 0.027068 | 1.77871 | 0.319022 | 0.974093 | 0.110962 | FGF7 | 15 |
| 417 | ENSG00000140416 | 2857.101 | 2.40556 | 0.153622 | 15.65891 | 2.89E-55 | 2.46E-52 | 61.13882 | 355.7575 | 66.43386 | 281.7284 | TPM1 | 15 |
| 418 | ENSG00000140488 | 84.2762 | -1.51474 | 0.403415 | -3.75479 | 0.000173 | 0.00272 | 5.296238 | 1.3655 | 4.550839 | 1.894855 | CELF6 | 15 |
| 419 | ENSG00000140682 | 127.8431 | 1.016738 | 0.333255 | 3.050937 | 0.002281 | 0.02451 | 6.786117 | 15.13028 | 7.866877 | 12.85045 | TGFB1I1 | 16 |
| 420 | ENSG00000140743 | 158.4059 | 1.004735 | 0.297687 | 3.375138 | 0.000738 | 0.009516 | 5.939331 | 11.39201 | 5.382506 | 10.00325 | CDR2 | 16 |
| 421 | ENSG00000140807 | 1076.908 | 1.730892 | 0.157722 | 10.97434 | 5.08E-28 | 1.18E-25 | 3.102795 | 10.62629 | 3.47208 | 10.00098 | NKD1 | 16 |
| 422 | ENSG00000140873 | 486.8794 | 4.006681 | 0.285914 | 14.0136 | 1.29E-44 | 7.86E-42 | 2.114637 | 28.53765 | 1.118125 | 18.85142 | ADAMTS1ℓ | 16 |
| 423 | ENSG00000140937 | 778.8067 | -1.56876 | 0.211958 | -7.40128 | 1.35E-13 | 8.97E-12 | 55.7812 | 18.52728 | 42.6084 | 12.65302 | CDH11 | 16 |
| 424 | ENSG00000141668 | 38.29405 | 2.119657 | 0.608095 | 3.48573 | 0.000491 | 0.006716 | 0.732164 | 3.853556 | 1.101871 | 3.509416 | CBLN2 | 18 |
| 425 | ENSG00000142178 | 286.7873 | 1.710246 | 0.253479 | 6.747103 | 1.51E-11 | 8.22E-10 | 8.447824 | 32.78188 | 11.68558 | 29.77902 | SIK1 | 21 |
| 426 | ENSG00000142227 | 58.42786 | 1.354896 | 0.475574 | 2.848969 | 0.004386 | 0.041333 | 7.567935 | 22.11598 | 10.28026 | 21.08708 | EMP3 | 19 |
| 427 | ENSG00000142627 | 864.7197 | 1.569114 | 0.187909 | 8.350383 | 6.80E-17 | 6.25E-15 | 15.66463 | 44.81978 | 12.89744 | 34.8383 | EPHA2 | 1 |
| 428 | ENSG00000142700 | 291.1007 | -4.5111 | 0.302263 | -14.9244 | 2.29E-50 | 1.72E-47 | 21.70928 | 1.046878 | 21.16913 | 0.719727 | DMRTA2 | 1 |
| 429 | ENSG00000142798 | 730.0692 | 1.574671 | 0.211059 | 7.460821 | 8.60E-14 | 5.86E-12 | 2.95588 | 11.10097 | 4.506151 | 9.902116 | HSPG2 | 1 |
| 430 | ENSG00000142871 | 593.144 | 1.583189 | 0.23637 | 6.69793 | 2.11E-11 | 1.13E-09 | 19.63175 | 57.17249 | 14.13616 | 37.61198 | CCN1 | 1 |
| 431 | ENSG00000143028 | 19.37627 | 2.384096 | 0.84432 | 2.823689 | 0.004747 | 0.043839 | 0.173613 | 0.974267 | 0.21417 | 0.943882 | SYPL2 | 1 |
| 432 | ENSG00000143341 | 142.4231 | 1.370659 | 0.380205 | 3.605055 | 0.000312 | 0.004585 | 0.470376 | 1.354253 | 0.943279 | 2.06763 | HMCN1 | 1 |
| 433 | ENSG00000143355 | 518.3378 | -1.91333 | 0.224798 | -8.5113 | 1.72E-17 | 1.73E-15 | 22.78241 | 7.135626 | 24.59841 | 4.70341 | LHX9 | 1 |
| 434 | ENSG00000143473 | 152.6514 | 1.91913 | 0.321702 | 5.965554 | 2.44E-09 | 1.01E-07 | 2.012946 | 7.176184 | 1.762116 | 6.227204 | KCNH1 | 1 |
| 435 | ENSG00000143479 | 584.3417 | -1.05294 | 0.224529 | -4.68954 | 2.74E-06 | 6.54E-05 | 17.5562 | 8.766223 | 26.26676 | 11.29697 | DYRK3 | 1 |
| 436 | ENSG00000143603 | 372.9297 | -1.16632 | 0.226563 | -5.14791 | 2.63E-07 | 7.77E-06 | 5.191641 | 2.277883 | 4.434305 | 1.759671 | KCNN3 | 1 |
| 437 | ENSG00000143842 | 193.263 | 1.107378 | 0.295847 | 3.743078 | 0.000182 | 0.002839 | 6.125679 | 9.019994 | 4.233017 | 11.95181 | SOX13 | 1 |
| 438 | ENSG00000143869 | 168.4532 | 1.467942 | 0.294623 | 4.98244 | 6.28E-07 | 1.69E-05 | 1.171121 | 2.590416 | 0.972147 | 3.005366 | GDF7 | 2 |
| 439 | ENSG00000143882 | 56.85539 | 1.949634 | 0.560249 | 3.479946 | 0.000502 | 0.006839 | 0.420274 | 3.604483 | 1.333697 | 2.86453 | ATP6V1C2 | 2 |
| 440 | ENSG00000144285 | 357.6754 | -1.3968 | 0.272038 | -5.13459 | 2.83E-07 | 8.29E-06 | 11.3835 | 5.761789 | 13.95716 | 3.314846 | SCN1A | 2 |
| 441 | ENSG00000144290 | 94.57105 | -2.32487 | 0.388232 | -5.98834 | 2.12E-09 | 8.95E-08 | 4.397747 | 0.867508 | 4.793709 | 0.864515 | SLC4A10 | 2 |
| 442 | ENSG00000144355 | 85.39491 | -8.30445 | 1.431311 | -5.80199 | 6.55E-09 | 2.52E-07 | 15.26357 | 0.056757 | 8.425753 | 0 | DLX1 | 2 |
| 443 | ENSG00000144369 | 1259.957 | -1.41758 | 0.162232 | -8.738 | 2.37E-18 | 2.56E-16 | 40.9055 | 12.83687 | 31.35449 | 12.65668 | FAM171B | 2 |
| 444 | ENSG00000144460 | 118.1472 | 1.254004 | 0.350301 | 3.579794 | 0.000344 | 0.004999 | 1.450214 | 3.693808 | 1.439185 | 2.795383 | NYAP2 | 2 |
| 445 | ENSG00000144476 | 317.3614 | -2.23443 | 0.24431 | -9.14589 | 5.91E-20 | 7.46E-18 | 35.93132 | 8.66819 | 43.18003 | 7.15695 | ACKR3 | 2 |
| 446 | ENSG00000144481 | 32.7039 | -3.01408 | 0.787279 | -3.82848 | 0.000129 | 0.002112 | 4.421548 | 0.210625 | 1.875254 | 0.719844 | TRPM8 | 2 |
| 447 | ENSG00000144583 | 83.20439 | -1.544 | 0.407664 | -3.78744 | 0.000152 | 0.002428 | 2.603472 | 1.024178 | 3.132879 | 0.831883 | MARCHF4 | 2 |
| 448 | ENSG00000144591 | 117.7962 | 1.198654 | 0.339399 | 3.5317 | 0.000413 | 0.005825 | 4.863266 | 11.48172 | 5.663136 | 11.33837 | GMPPA | 2 |
| 449 | ENSG00000144642 | 74.46559 | 1.429624 | 0.444922 | 3.213198 | 0.001313 | 0.015603 | 1.765897 | 4.404006 | 1.225453 | 3.055736 | RBMS3 | 3 |
| 450 | ENSG00000144655 | 124.7124 | 1.182266 | 0.380271 | 3.10901 | 0.001877 | 0.020951 | 2.756023 | 7.825223 | 2.900674 | 4.229526 | CSRNP1 | 3 |
| 451 | ENSG00000144749 | 1185.848 | -1.03556 | 0.180139 | -5.7487 | 8.99E-09 | 3.39E-07 | 46.79211 | 20.90881 | 35.28555 | 16.7504 | LRIG1 | 3 |
| 452 | ENSG00000144810 | 97.92126 | 1.764218 | 0.402266 | 4.385703 | 1.16E-05 | 0.000242 | 0.994032 | 4.605461 | 1.56047 | 3.529414 | COL8A1 | 3 |
| 453 | ENSG00000144847 | 217.0394 | -3.15929 | 0.342662 | -9.21985 | 2.98E-20 | 3.83E-18 | 23.71027 | 2.558097 | 14.43074 | 1.447947 | IGSF11 | 3 |
| 454 | ENSG00000144857 | 1370.574 | 1.066363 | 0.155824 | 6.843362 | 7.74E-12 | 4.35E-10 | 28.66063 | 54.37378 | 32.4867 | 66.9092 | BOC | 3 |
| 455 | ENSG00000145246 | 508.8193 | 1.308674 | 0.192277 | 6.806205 | 1.00E-11 | 5.57E-10 | 6.161036 | 15.81751 | 6.971594 | 14.92014 | ATP10D | 4 |
| 456 | ENSG00000145358 | 297.9294 | 3.194851 | 0.272643 | 11.71808 | 1.03E-31 | 3.25E-29 | 2.697722 | 26.76056 | 2.738332 | 20.13678 | DDIT4L | 4 |
| 457 | ENSG00000145431 | 300.5141 | 1.352407 | 0.239304 | 5.651419 | 1.59E-08 | 5.77E-07 | 7.286053 | 19.9463 | 7.81397 | 16.42873 | PDGFC | 4 |
| 458 | ENSG00000145506 | 108.4769 | 2.329057 | 0.383756 | 6.069107 | 1.29E-09 | 5.58E-08 | 3.226265 | 14.84244 | 2.407028 | 11.40969 | NKD2 | 5 |
| 459 | ENSG00000145526 | 363.7179 | -1.30824 | 0.239007 | -5.47364 | 4.41E-08 | 1.49E-06 | 23.14954 | 10.25967 | 33.20614 | 11.28619 | CDH18 | 5 |
| 460 | ENSG00000145536 | 83.5064 | 1.304067 | 0.430284 | 3.030708 | 0.00244 | 0.025801 | 1.251514 | 4.036032 | 1.53717 | 2.434383 | ADAMTS1( | 5 |
| 461 | ENSG00000145555 | 2854.829 | -2.03632 | 0.123872 | -16.4389 | 1.01E-60 | 1.04E-57 | 82.80469 | 18.80049 | 84.80705 | 19.82001 | MYO10 | 5 |
| 462 | ENSG00000145632 | 1366.97 | 1.383549 | 0.155396 | 8.903358 | 5.42E-19 | 6.23E-17 | 48.23206 | 110.6417 | 38.73777 | 103.0258 | PLK2 | 5 |
| 463 | ENSG00000145721 | 687.1489 | -2.46755 | 0.243055 | -10.1522 | 3.24E-24 | 6.03E-22 | 45.87107 | 6.912227 | 26.31789 | 5.331632 | LIX1 | 5 |
| 464 | ENSG00000145808 | 16.01807 | 3.8244 | 1.240187 | 3.083729 | 0.002044 | 0.022507 | 0.248397 | 1.555569 | 0.021335 | 0.596657 | ADAMTS1! | 5 |
| 465 | ENSG00000145824 | 38.60261 | 2.444385 | 0.628076 | 3.891864 | 9.95E-05 | 0.001671 | 0.91601 | 5.725083 | 0.882877 | 3.470934 | CXCL14 | 5 |
| 466 | ENSG00000145920 | 743.2275 | -1.02728 | 0.192241 | -5.34371 | 9.11E-08 | 2.93E-06 | 31.47164 | 11.3667 | 22.86641 | 13.82434 | CPLX2 | 5 |
| 467 | ENSG00000145934 | 2277.968 | 1.44852 | 0.129493 | 11.18606 | 4.77E-29 | 1.23E-26 | 22.12541 | 51.95136 | 20.16109 | 57.10157 | TENM2 | 5 |

| | | | | | | | | | | | | | | |
|---|---|---|---|---|---|---|---|---|---|---|---|---|---|---|
| 468 | ENSG00000146001 | 90.99146 | -1.43566 | 0.407636 | -3.52192 | 0.000428 | 0.006005 | 6.069609 | 2.208567 | 4.498556 | 1.443332 | PCDHB18F | | 5 |
| 469 | ENSG00000146005 | 253.353 | -1.07571 | 0.29199 | -3.68406 | 0.00023 | 0.003493 | 14.51323 | 4.169788 | 9.432431 | 6.607832 | PSD2 | | 5 |
| 470 | ENSG00000146072 | 316.861 | 1.009486 | 0.222047 | 4.546264 | 5.46E-06 | 0.000122 | 7.060895 | 13.18628 | 6.50631 | 12.57513 | TNFRSF21 | | 6 |
| 471 | ENSG00000146250 | 303.1319 | 1.068882 | 0.265119 | 4.031706 | 5.54E-05 | 0.000992 | 9.501144 | 23.04666 | 9.729752 | 14.89799 | PRSS35 | | 6 |
| 472 | ENSG00000146374 | 306.6788 | -2.79381 | 0.247551 | -11.2858 | 1.54E-29 | 4.07E-27 | 14.30791 | 2.120339 | 16.82011 | 2.125728 | RSPO3 | | 6 |
| 473 | ENSG00000146411 | 200.206 | -1.04269 | 0.303215 | -3.43879 | 0.000584 | 0.007864 | 6.649179 | 2.826684 | 4.306864 | 2.16288 | SLC2A12 | | 6 |
| 474 | ENSG00000146648 | 622.2207 | 2.061384 | 0.243069 | 8.480659 | 2.24E-17 | 2.22E-15 | 2.386412 | 10.93951 | 4.128482 | 14.81447 | EGFR | | 7 |
| 475 | ENSG00000146674 | 126.5075 | 1.375709 | 0.380774 | 3.612931 | 0.000303 | 0.004469 | 3.954703 | 13.14299 | 4.386487 | 7.272213 | IGFBP3 | | 7 |
| 476 | ENSG00000146910 | 82.25002 | 4.476151 | 1.2156 | 3.682257 | 0.000231 | 0.003515 | 0.086265 | 6.983177 | 0.794596 | 12.5599 | CNPY1 | | 7 |
| 477 | ENSG00000147100 | 1604.764 | -1.30952 | 0.144379 | -9.07002 | 1.19E-19 | 1.47E-17 | 64.98648 | 26.27058 | 64.58554 | 23.03437 | SLC16A2 | X | |
| 478 | ENSG00000147145 | 230.2833 | -1.41365 | 0.259886 | -5.43952 | 5.34E-08 | 1.78E-06 | 11.03787 | 3.986549 | 9.665686 | 3.343595 | LPAR4 | X | |
| 479 | ENSG00000147246 | 73.46689 | 1.676802 | 0.507743 | 3.30246 | 0.000958 | 0.011922 | 1.264306 | 3.304843 | 0.503937 | 1.927034 | HTR2C | X | |
| 480 | ENSG00000147256 | 40.07136 | -1.77611 | 0.640852 | -2.77148 | 0.00558 | 0.049892 | 2.509215 | 1.272147 | 3.671838 | 0.417131 | ARHGAP3( | X | |
| 481 | ENSG00000147459 | 333.4638 | 1.382721 | 0.235759 | 5.864967 | 4.49E-09 | 1.79E-07 | 3.09177 | 8.410396 | 4.1574 | 9.486085 | DOCK5 | | 8 |
| 482 | ENSG00000147488 | 647.9326 | -1.27503 | 0.204862 | -6.22384 | 4.85E-10 | 2.26E-08 | 46.68892 | 14.34345 | 31.62203 | 16.15945 | ST18 | | 8 |
| 483 | ENSG00000147509 | 295.82 | -1.18125 | 0.225853 | -5.23015 | 1.69E-07 | 5.14E-06 | 41.42983 | 16.63285 | 40.82939 | 17.64236 | RGS20 | | 8 |
| 484 | ENSG00000147571 | 20.81639 | -3.30254 | 0.879773 | -3.75385 | 0.000174 | 0.002728 | 3.454082 | 0.360362 | 4.381851 | 0.392103 | CRH | | 8 |
| 485 | ENSG00000147862 | 645.522 | 1.068079 | 0.190147 | 5.617129 | 1.94E-08 | 6.95E-07 | 9.422735 | 21.95 | 12.08473 | 20.93798 | NFIB | | 9 |
| 486 | ENSG00000148053 | 8762.549 | -1.38706 | 0.14104 | -9.83447 | 8.00E-23 | 1.27E-20 | 214.8208 | 66.95029 | 157.9445 | 67.40618 | NTRK2 | | 9 |
| 487 | ENSG00000148123 | 198.6363 | -1.15443 | 0.275405 | -4.19176 | 2.77E-05 | 0.000529 | 17.13842 | 6.420981 | 13.32927 | 6.469916 | PLPPR1 | | 9 |
| 488 | ENSG00000148175 | 58.05188 | 1.410166 | 0.486384 | 2.899283 | 0.00374 | 0.03645 | 1.159317 | 3.486364 | 1.204386 | 2.427343 | STOM | | 9 |
| 489 | ENSG00000148488 | 43.30208 | -2.42056 | 0.631424 | -3.8335 | 0.000126 | 0.002076 | 1.111985 | 0.379184 | 1.728712 | 0.13595 | ST8SIA6 | | 10 |
| 490 | ENSG00000148600 | 64.54575 | -1.48123 | 0.462685 | -3.20138 | 0.001368 | 0.01612 | 2.483447 | 0.830817 | 1.855008 | 0.632049 | CDHR1 | | 10 |
| 491 | ENSG00000148677 | 76.28917 | 1.902552 | 0.457902 | 4.15493 | 3.25E-05 | 0.000612 | 2.098475 | 9.955383 | 2.402629 | 5.883094 | ANKRD1 | | 10 |
| 492 | ENSG00000149218 | 187.8335 | 1.478895 | 0.287608 | 5.142049 | 2.72E-07 | 7.99E-06 | 2.561624 | 7.049053 | 2.304237 | 5.716755 | ENDOD1 | | 11 |
| 493 | ENSG00000149257 | 720.7492 | 1.59718 | 0.217484 | 7.343902 | 2.07E-13 | 1.35E-11 | 16.17469 | 57.02993 | 26.22525 | 64.80627 | SERPINH1 | | 11 |
| 494 | ENSG00000149571 | 191.3042 | 1.704173 | 0.27905 | 6.107057 | 1.01E-09 | 4.52E-08 | 4.037655 | 11.76547 | 3.47756 | 11.31532 | KIRREL3 | | 11 |
| 495 | ENSG00000149591 | 1923.792 | 1.570158 | 0.191824 | 8.185418 | 2.71E-16 | 2.38E-14 | 68.16197 | 250.3224 | 80.53519 | 165.5525 | TAGLN | | 11 |
| 496 | ENSG00000149596 | 46.15177 | 2.012069 | 0.585187 | 3.438338 | 0.000585 | 0.007867 | 0.418255 | 1.383671 | 0.190931 | 0.926998 | JPH2 | | 20 |
| 497 | ENSG00000149633 | 39.29012 | -1.90344 | 0.598874 | -3.17837 | 0.001481 | 0.017259 | 2.372353 | 0.778917 | 2.385635 | 0.40966 | KIAA1755 | | 20 |
| 498 | ENSG00000150275 | 519.9012 | -4.35151 | 0.258955 | -16.8041 | 2.28E-63 | 2.62E-60 | 24.00264 | 1.396077 | 21.22692 | 0.704354 | PCDH15 | | 10 |
| 499 | ENSG00000150551 | 413.1513 | 1.425784 | 0.223944 | 6.366707 | 1.93E-10 | 9.37E-09 | 10.05484 | 24.90427 | 7.78718 | 20.1895 | LYPD1 | | 2 |
| 500 | ENSG00000150672 | 569.2486 | -2.37213 | 0.219164 | -10.8235 | 2.66E-27 | 5.79E-25 | 41.27447 | 8.069008 | 33.69901 | 5.580328 | DLG2 | | 11 |
| 501 | ENSG00000150687 | 2129.573 | -1.10468 | 0.180928 | -6.10561 | 1.02E-09 | 4.55E-08 | 93.3428 | 51.18176 | 102.2584 | 34.43727 | PRSS23 | | 11 |
| 502 | ENSG00000150907 | 189.3291 | 1.402274 | 0.28177 | 4.976659 | 6.47E-07 | 1.74E-05 | 1.909289 | 5.374074 | 2.139647 | 4.734999 | FOXO1 | | 13 |
| 503 | ENSG00000151012 | 749.8015 | 1.852524 | 0.221951 | 8.34655 | 7.03E-17 | 6.42E-15 | 4.802658 | 12.09278 | 2.697905 | 13.38132 | SLC7A11 | | 4 |
| 504 | ENSG00000151067 | 457.6565 | -1.03216 | 0.192964 | -5.34898 | 8.84E-08 | 2.86E-06 | 16.58992 | 7.015451 | 15.62038 | 7.880818 | CACNA1C | | 12 |
| 505 | ENSG00000151090 | 62.9612 | 1.762225 | 0.476611 | 3.697407 | 0.000218 | 0.003333 | 1.120476 | 3.71176 | 1.002777 | 3.071956 | THRB | | 3 |
| 506 | ENSG00000151388 | 461.5027 | -1.02534 | 0.201778 | -5.08151 | 3.74E-07 | 1.07E-05 | 12.91348 | 6.31394 | 12.07141 | 5.246487 | ADAMTS1; | | 5 |
| 507 | ENSG00000151468 | 67.12607 | 2.324697 | 0.478671 | 4.856565 | 1.19E-06 | 3.05E-05 | 0.815847 | 3.435635 | 1.009688 | 5.301747 | CCDC3 | | 10 |
| 508 | ENSG00000151490 | 646.597 | -1.22745 | 0.176363 | -6.9598 | 3.41E-12 | 1.97E-10 | 30.8003 | 12.76873 | 29.39291 | 11.48059 | PTPRO | | 12 |
| 509 | ENSG00000151693 | 2227.547 | -1.18219 | 0.128845 | -9.17528 | 4.50E-20 | 5.76E-18 | 91.02771 | 35.79536 | 82.39111 | 36.33879 | ASAP2 | | 2 |
| 510 | ENSG00000151726 | 256.5396 | 1.737939 | 0.246052 | 7.063294 | 1.63E-12 | 9.63E-11 | 4.149721 | 12.17746 | 3.661929 | 12.38471 | ACSL1 | | 4 |
| 511 | ENSG00000151789 | 272.9901 | -1.43486 | 0.298754 | -4.80282 | 1.56E-06 | 3.90E-05 | 9.02708 | 1.976108 | 5.310366 | 3.088712 | ZNF385D | | 3 |
| 512 | ENSG00000151834 | 244.9934 | -3.70054 | 0.314995 | -11.7479 | 7.24E-32 | 2.32E-29 | 14.3392 | 0.832945 | 10.02794 | 0.945995 | GABRA2 | | 4 |
| 513 | ENSG00000152092 | 1257.883 | -1.60967 | 0.149755 | -10.7487 | 6.01E-27 | 1.26E-24 | 35.60061 | 11.6922 | 36.2645 | 10.52669 | ASTN1 | | 1 |
| 514 | ENSG00000152104 | 652.3148 | 1.081939 | 0.176469 | 6.131036 | 8.73E-10 | 3.94E-08 | 4.039572 | 8.715881 | 4.388625 | 8.131565 | PTPN14 | | 1 |
| 515 | ENSG00000152192 | 251.758 | 2.437418 | 0.289756 | 8.411953 | 4.03E-17 | 3.81E-15 | 1.481994 | 8.888959 | 2.438322 | 11.35199 | POU4F1 | | 13 |
| 516 | ENSG00000152467 | 11.3577 | -7.2922 | 1.76092 | -4.14113 | 3.46E-05 | 0.000646 | 2.790764 | 0 | 2.949357 | 0 | ZSCAN1 | | 19 |
| 517 | ENSG00000152583 | 191.8961 | 1.12615 | 0.371895 | 3.028142 | 0.002461 | 0.025955 | 2.850655 | 10.5363 | 7.618786 | 11.1965 | SPARCL1 | | 4 |
| 518 | ENSG00000152661 | 1949.919 | 1.28625 | 0.191635 | 6.711977 | 1.92E-11 | 1.03E-09 | 37.11232 | 116.5685 | 49.21632 | 82.03477 | GJA1 | | 6 |
| 519 | ENSG00000152778 | 463.4038 | -1.05125 | 0.208659 | -5.03813 | 4.70E-07 | 1.32E-05 | 26.28802 | 9.777899 | 24.73841 | 13.60775 | IFIT5 | | 10 |
| 520 | ENSG00000152785 | 18.12395 | -3.54382 | 0.96991 | -3.65376 | 0.000258 | 0.003878 | 0.637129 | 0.064031 | 0.597156 | 0.034904 | BMP3 | | 4 |
| 521 | ENSG00000152910 | 110.9352 | -1.59114 | 0.377609 | -4.21374 | 2.51E-05 | 0.000484 | 6.214406 | 1.3498 | 4.135434 | 1.927926 | CNTNAP4 | | 16 |
| 522 | ENSG00000153012 | 414.2573 | 2.790271 | 0.243724 | 11.4485 | 2.39E-30 | 6.88E-28 | 2.055559 | 14.6083 | 1.800948 | 10.49496 | LGI2 | | 4 |
| 523 | ENSG00000153071 | 159.8138 | 2.25434 | 0.320321 | 7.037763 | 1.95E-12 | 1.15E-10 | 2.103501 | 11.43022 | 2.578989 | 9.547476 | DAB2 | | 5 |
| 524 | ENSG00000153179 | 150.6999 | 1.099672 | 0.368658 | 2.982909 | 0.002855 | 0.029069 | 2.171736 | 7.584706 | 4.541889 | 6.037299 | RASSF3 | | 12 |
| 525 | ENSG00000153233 | 32.38877 | 2.355597 | 0.692969 | 3.399283 | 0.000676 | 0.008885 | 0.373903 | 2.863678 | 0.577102 | 1.829655 | PTPRR | | 12 |
| 526 | ENSG00000153250 | 347.7764 | 1.393342 | 0.220331 | 6.32385 | 2.55E-10 | 1.22E-08 | 7.625858 | 20.72807 | 8.49562 | 19.27339 | RBMS1 | | 2 |
| 527 | ENSG00000153266 | 79.05801 | -1.81369 | 0.436224 | -4.1577 | 3.21E-05 | 0.000605 | 7.793097 | 2.483867 | 6.981945 | 1.453583 | FEZF2 | | 3 |
| 528 | ENSG00000153822 | 13.13741 | -4.71998 | 1.395987 | -3.3811 | 0.000722 | 0.009355 | 0.953859 | 0.054426 | 0.853623 | 0 | KCNJ16 | | 17 |
| 529 | ENSG00000153930 | 128.1936 | -1.16044 | 0.361802 | -3.2074 | 0.001339 | 0.015854 | 3.654206 | 2.233119 | 5.000118 | 1.431501 | ANKFN1 | | 17 |
| 530 | ENSG00000153976 | 81.0693 | 1.661244 | 0.416206 | 3.991399 | 6.57E-05 | 0.001157 | 1.975864 | 5.257629 | 1.407104 | 4.755213 | HS3ST3A1 | | 17 |
| 531 | ENSG00000153993 | 122.6913 | -1.6039 | 0.344461 | -4.65627 | 3.22E-06 | 7.57E-05 | 3.70378 | 1.358065 | 3.898429 | 0.999655 | SEMA3D | | 7 |
| 532 | ENSG00000154118 | 346.3759 | -1.31517 | 0.215572 | -6.10082 | 1.06E-09 | 4.67E-08 | 14.28469 | 5.092464 | 12.68324 | 5.131719 | JPH3 | | 16 |
| 533 | ENSG00000154188 | 168.617 | 1.962504 | 0.393229 | 4.990737 | 6.01E-07 | 1.64E-05 | 1.786275 | 13.31179 | 3.718223 | 7.189363 | ANGPT1 | | 8 |
| 534 | ENSG00000154265 | 261.199 | -1.42367 | 0.266274 | -5.34664 | 8.96E-08 | 2.89E-06 | 13.8682 | 3.79794 | 9.324744 | 4.345828 | ABCA5 | | 17 |
| 535 | ENSG00000154447 | 582.3099 | -1.12002 | 0.191681 | -5.84316 | 5.12E-09 | 2.01E-07 | 23.36467 | 9.407199 | 18.39353 | 8.696287 | SH3RF1 | | 4 |
| 536 | ENSG00000154678 | 301.0028 | 1.551382 | 0.3002 | 5.167835 | 2.37E-07 | 7.03E-06 | 4.009272 | 13.1467 | 3.31844 | 6.975154 | PDE1C | | 7 |
| 537 | ENSG00000154760 | 133.5564 | 1.274137 | 0.332415 | 3.832971 | 0.000127 | 0.002079 | 2.828542 | 5.072856 | 2.515693 | 7.180521 | SLFN13 | | 17 |
| 538 | ENSG00000154856 | 616.2957 | 3.342303 | 0.201846 | 16.55871 | 1.39E-61 | 1.50E-58 | 5.450517 | 43.47646 | 4.607202 | 52.64301 | APCDD1 | | 18 |
| 539 | ENSG00000155011 | 626.8672 | 2.529787 | 0.570246 | 4.436307 | 9.15E-06 | 0.000196 | 3.874392 | 37.28542 | 8.318508 | 29.3864 | DKK2 | | 4 |
| 540 | ENSG00000155760 | 549.4811 | -1.12925 | 0.207194 | -5.4502 | 5.03E-08 | 1.69E-06 | 21.84589 | 8.553556 | 15.7345 | 7.609524 | FZD7 | | 2 |
| 541 | ENSG00000155875 | 77.53078 | -2.66002 | 0.459625 | -5.78738 | 7.15E-09 | 2.75E-07 | 1.772836 | 0.345096 | 1.848154 | 0.186197 | SAXO1 | | 9 |
| 542 | ENSG00000155974 | 536.4419 | -1.61577 | 0.211301 | -7.64676 | 2.06E-14 | 1.53E-12 | 65.57117 | 18.07749 | 46.09192 | 16.1204 | GRIP1 | | 12 |
| 543 | ENSG00000156113 | 209.3576 | -1.53349 | 0.274734 | -5.58171 | 2.38E-08 | 8.44E-07 | 10.15775 | 3.559179 | 9.382293 | 2.768882 | KCNMA1 | | 10 |
| 544 | ENSG00000156140 | 215.8046 | -1.09351 | 0.265394 | -4.12034 | 3.78E-05 | 0.000703 | 5.488991 | 2.318722 | 6.213677 | 2.885646 | ADAMTS3 | | 4 |
| 545 | ENSG00000156253 | 20.74971 | -3.40562 | 0.894807 | -3.80598 | 0.000141 | 0.002275 | 2.039163 | 0.099597 | 2.076772 | 0.296083 | RWDD2B | | 21 |

| | | | | | | | | | | | | | |
|---|---|---|---|---|---|---|---|---|---|---|---|---|---|
| 546 | ENSG00000156398 | 218.8104 | 1.184938 | 0.290163 | 4.083699 | 4.43E-05 | 0.000812 | 4.222317 | 8.243007 | 5.386423 | 12.57579 | SFXN2 | 10 |
| 547 | ENSG00000156475 | 460.7506 | -1.39425 | 0.205782 | -6.77537 | 1.24E-11 | 6.82E-10 | 19.78434 | 8.18382 | 22.49701 | 7.010572 | PPP2R2B | 5 |
| 548 | ENSG00000156535 | 90.78107 | 1.281768 | 0.399691 | 3.206901 | 0.001342 | 0.015872 | 0.661101 | 1.866842 | 1.035451 | 2.026756 | CD109 | 6 |
| 549 | ENSG00000156675 | 420.195 | 1.330359 | 0.218861 | 6.078558 | 1.21E-09 | 5.29E-08 | 4.676169 | 12.97969 | 5.127997 | 10.26715 | RAB11FIP1 | 8 |
| 550 | ENSG00000157087 | 120.707 | -1.32384 | 0.346397 | -3.82175 | 0.000133 | 0.002157 | 4.922258 | 1.845461 | 3.900773 | 1.459205 | ATP2B2 | 3 |
| 551 | ENSG00000157168 | 449.6689 | 1.432901 | 0.197643 | 7.249949 | 4.17E-13 | 2.66E-11 | 7.156678 | 18.66838 | 7.053651 | 17.54784 | NRG1 | 8 |
| 552 | ENSG00000157227 | 186.4414 | 1.534315 | 0.322844 | 4.752499 | 2.01E-06 | 4.93E-05 | 2.413603 | 9.176911 | 4.490124 | 9.889839 | MMP14 | 14 |
| 553 | ENSG00000157502 | 88.39683 | 2.204197 | 0.427644 | 5.154285 | 2.55E-07 | 7.53E-06 | 1.002743 | 4.674917 | 0.797888 | 3.099767 | PWWP3B | X |
| 554 | ENSG00000157978 | 86.1062 | 1.571339 | 0.401062 | 3.917946 | 8.93E-05 | 0.001519 | 1.990973 | 4.791999 | 1.59285 | 5.156539 | LDLRAP1 | 1 |
| 555 | ENSG00000158104 | 38.50459 | 3.811297 | 0.77248 | 4.933844 | 8.06E-07 | 2.12E-05 | 0.102286 | 8.120512 | 0.952701 | 5.386508 | HPD | 12 |
| 556 | ENSG00000158246 | 86.01743 | 2.02183 | 0.412712 | 4.898883 | 9.64E-07 | 2.51E-05 | 1.910414 | 7.277366 | 1.536639 | 5.87447 | TENT5B | 1 |
| 557 | ENSG00000158270 | 103.3273 | 4.161381 | 0.463462 | 8.978908 | 2.73E-19 | 3.26E-17 | 0.214697 | 3.869364 | 0.265912 | 4.350245 | COLEC12 | 18 |
| 558 | ENSG00000158560 | 78.58954 | -1.65239 | 0.490956 | -3.36565 | 0.000764 | 0.009826 | 6.468825 | 2.817737 | 5.952662 | 0.982381 | DYNC1I1 | 7 |
| 559 | ENSG00000158955 | 106.5928 | 4.5996 | 0.483531 | 9.512521 | 1.86E-21 | 2.58E-19 | 0.254007 | 6.870537 | 0.26609 | 4.977018 | WNT9B | 17 |
| 560 | ENSG00000159176 | 98.2339 | -3.39543 | 0.459506 | -7.3893 | 1.48E-13 | 9.78E-12 | 9.687913 | 1.33972 | 9.662475 | 0.496745 | CSRP1 | 1 |
| 561 | ENSG00000159212 | 214.3345 | 3.018655 | 0.298075 | 10.12715 | 4.19E-24 | 7.72E-22 | 1.158464 | 10.80274 | 1.449977 | 9.214465 | CLIC6 | 21 |
| 562 | ENSG00000159231 | 18.09457 | 2.962032 | 0.9336 | 3.172697 | 0.00151 | 0.017558 | 0.318979 | 4.252291 | 0.849044 | 4.486561 | CBR3 | 21 |
| 563 | ENSG00000159248 | 126.2021 | -3.42812 | 0.39183 | -8.74899 | 2.15E-18 | 2.34E-16 | 9.105263 | 1.049289 | 9.530232 | 0.571697 | GJD2 | 15 |
| 564 | ENSG00000159251 | 45.38389 | 2.507062 | 0.662019 | 3.786994 | 0.000152 | 0.00243 | 1.584235 | 10.90356 | 1.12727 | 3.505956 | ACTC1 | 15 |
| 565 | ENSG00000159263 | 49.94868 | 2.843988 | 0.58017 | 4.901987 | 9.49E-07 | 2.48E-05 | 0.422058 | 3.738862 | 0.443551 | 2.11027 | SIM2 | 21 |
| 566 | ENSG00000159307 | 1450.688 | -1.71485 | 0.141335 | -12.1332 | 7.04E-34 | 2.70E-31 | 40.87343 | 11.20914 | 36.89591 | 11.15006 | SCUBE1 | 22 |
| 567 | ENSG00000159387 | 36.49986 | 2.676543 | 0.659097 | 4.060927 | 4.89E-05 | 0.000887 | 0.394331 | 1.883816 | 0.478145 | 3.434917 | IRX6 | 16 |
| 568 | ENSG00000159409 | 1141.962 | -1.12151 | 0.148588 | -7.54783 | 4.43E-14 | 3.14E-12 | 63.05015 | 25.50079 | 55.79502 | 26.05173 | CELF3 | 1 |
| 569 | ENSG00000159713 | 115.2394 | -1.0315 | 0.340737 | -3.02725 | 0.002468 | 0.025987 | 20.54245 | 10.33897 | 21.83266 | 9.221282 | TPPP3 | 16 |
| 570 | ENSG00000159788 | 730.3777 | -1.24887 | 0.239829 | -5.20735 | 1.92E-07 | 5.76E-06 | 36.34456 | 21.1847 | 65.64884 | 19.31223 | RGS12 | 4 |
| 571 | ENSG00000160188 | 32.47995 | -1.79112 | 0.637128 | -2.81125 | 0.004935 | 0.045135 | 4.011745 | 1.351665 | 4.952151 | 1.096694 | RSPH1 | 21 |
| 572 | ENSG00000160190 | 218.442 | -1.41442 | 0.263571 | -5.36639 | 8.03E-08 | 2.61E-06 | 15.71434 | 5.198591 | 12.92751 | 4.917406 | SLC37A1 | 21 |
| 573 | ENSG00000160691 | 464.0877 | 1.068889 | 0.192014 | 5.566715 | 2.60E-08 | 9.14E-07 | 13.78308 | 26.94395 | 13.24455 | 26.56852 | SHC1 | 1 |
| 574 | ENSG00000160963 | 469.2459 | 1.654491 | 0.209092 | 7.912751 | 2.52E-15 | 2.03E-13 | 9.14617 | 30.64062 | 9.537714 | 24.83265 | COL26A1 | 7 |
| 575 | ENSG00000161298 | 429.3518 | -1.47702 | 0.200787 | -7.35613 | 1.89E-13 | 1.24E-11 | 21.27753 | 6.615115 | 18.66256 | 6.923896 | ZNF382 | 19 |
| 576 | ENSG00000162374 | 1045.626 | -1.96781 | 0.170638 | -11.5321 | 9.09E-31 | 2.65E-28 | 80.1344 | 16.33156 | 61.04678 | 17.72103 | ELAVL4 | 1 |
| 577 | ENSG00000162407 | 623.0231 | -1.52048 | 0.18024 | -8.43589 | 3.29E-17 | 3.17E-15 | 36.21426 | 11.74736 | 32.23913 | 10.74043 | PLPP3 | 1 |
| 578 | ENSG00000162493 | 365.3876 | 2.508795 | 0.231109 | 10.85547 | 1.88E-27 | 4.13E-25 | 6.665266 | 36.21419 | 8.035505 | 42.72066 | PDPN | 1 |
| 579 | ENSG00000162520 | 88.94408 | 2.357175 | 0.41855 | 5.631759 | 1.78E-08 | 6.42E-07 | 0.847649 | 5.228386 | 1.071767 | 4.070169 | SYNC | 1 |
| 580 | ENSG00000162551 | 257.6132 | 1.489857 | 0.246092 | 6.054069 | 1.41E-09 | 6.11E-08 | 6.019087 | 16.61266 | 6.93406 | 17.83808 | ALPL | 1 |
| 581 | ENSG00000162552 | 186.7657 | 2.794493 | 0.336481 | 8.305046 | 9.98E-17 | 9.03E-15 | 2.229277 | 9.130188 | 1.250211 | 13.35464 | WNT4 | 1 |
| 582 | ENSG00000162599 | 3516.314 | 2.017616 | 0.593543 | 3.399274 | 0.000676 | 0.008885 | 5.747004 | 39.53757 | 12.80467 | 31.58618 | NFIA | 1 |
| 583 | ENSG00000162643 | 85.15157 | -1.41242 | 0.416688 | -3.38963 | 0.0007 | 0.009154 | 4.015062 | 1.82875 | 6.386986 | 1.862436 | DNAI3 | 1 |
| 584 | ENSG00000162692 | 341.685 | -6.37332 | 1.04688 | -6.08792 | 1.14E-09 | 5.03E-08 | 41.64799 | 0.420021 | 11.76063 | 0.181993 | VCAM1 | 1 |
| 585 | ENSG00000162772 | 698.958 | 2.377146 | 0.214862 | 11.06358 | 1.88E-28 | 4.61E-26 | 22.55576 | 106.177 | 15.89765 | 80.71809 | ATF3 | 1 |
| 586 | ENSG00000162804 | 114.9591 | 1.493711 | 0.342629 | 4.359555 | 1.30E-05 | 0.000271 | 1.121543 | 3.059835 | 1.123164 | 2.912928 | SNED1 | 2 |
| 587 | ENSG00000162849 | 462.0668 | -1.20777 | 0.214971 | -5.61829 | 1.93E-08 | 6.92E-07 | 6.212087 | 2.240638 | 4.457741 | 2.107444 | KIF26B | 1 |
| 588 | ENSG00000162873 | 1449.108 | -2.01962 | 0.163941 | -12.3192 | 7.14E-35 | 3.04E-32 | 94.09602 | 25.98961 | 113.6081 | 22.34563 | KLHDC8A | 1 |
| 589 | ENSG00000162981 | 173.9023 | -1.2018 | 0.307705 | -3.9057 | 9.40E-05 | 0.001585 | 6.518353 | 3.26015 | 6.727797 | 2.162882 | LRATD1 | 2 |
| 590 | ENSG00000162992 | 55.38672 | -2.28387 | 0.510501 | -4.47377 | 7.69E-06 | 0.000168 | 5.074326 | 1.069561 | 4.557794 | 0.775675 | NEUROD1 | 2 |
| 591 | ENSG00000163017 | 99.44401 | 2.819001 | 0.473094 | 5.958644 | 2.54E-09 | 1.05E-07 | 1.954324 | 23.01172 | 3.10161 | 11.29962 | ACTG2 | 2 |
| 592 | ENSG00000163071 | 60.66036 | -1.54519 | 0.476349 | -3.24382 | 0.001179 | 0.014288 | 4.296136 | 1.56164 | 3.72254 | 1.03669 | SPATA18 | 4 |
| 593 | ENSG00000163081 | 23.64421 | 2.867587 | 0.869724 | 3.297124 | 0.000977 | 0.012097 | 0.165172 | 4.225294 | 1.060328 | 3.643204 | CCDC140 | 2 |
| 594 | ENSG00000163082 | 134.3797 | 1.642802 | 0.322243 | 5.098023 | 3.43E-07 | 9.90E-06 | 1.520502 | 4.24462 | 1.45068 | 4.519568 | SGPP2 | 2 |
| 595 | ENSG00000163132 | 16.27537 | 2.976364 | 0.976653 | 3.047516 | 0.002307 | 0.024723 | 0.28337 | 1.382809 | 0.187641 | 2.108407 | MSX1 | 4 |
| 596 | ENSG00000163191 | 73.78882 | 2.907202 | 0.550533 | 5.280706 | 1.29E-07 | 3.99E-06 | 1.887203 | 43.04396 | 9.824663 | 41.41633 | S100A11 | 1 |
| 597 | ENSG00000163251 | 236.0536 | 2.126807 | 0.325967 | 6.524616 | 6.82E-11 | 3.46E-09 | 1.767257 | 7.142817 | 1.04235 | 4.321568 | FZD5 | 2 |
| 598 | ENSG00000163273 | 25.47896 | 3.751085 | 0.878299 | 4.270853 | 1.95E-05 | 0.000387 | 0.148264 | 6.664797 | 0.78879 | 5.559272 | NPPC | 2 |
| 599 | ENSG00000163285 | 87.05538 | -5.29711 | 0.642185 | -8.24856 | 1.60E-16 | 1.43E-14 | 3.816058 | 0.057428 | 1.83392 | 0.078266 | GABRG1 | 4 |
| 600 | ENSG00000163347 | 141.4265 | 2.327213 | 0.399621 | 5.823555 | 5.76E-09 | 2.24E-07 | 2.397192 | 10.05123 | 1.123949 | 6.182223 | CLDN1 | 3 |
| 601 | ENSG00000163359 | 26.17055 | 2.619475 | 0.838472 | 3.124105 | 0.001783 | 0.020112 | 0.146868 | 1.938674 | 0.40585 | 1.304835 | COL6A3 | 2 |
| 602 | ENSG00000163377 | 91.04596 | 1.744656 | 0.437259 | 3.989985 | 6.61E-05 | 0.001163 | 1.513054 | 8.407056 | 3.150882 | 6.428891 | TAFA4 | 3 |
| 603 | ENSG00000163453 | 60.09296 | 2.595515 | 0.506744 | 5.121949 | 3.02E-07 | 8.83E-06 | 1.874331 | 12.25768 | 2.062427 | 10.23266 | IGFBP7 | 4 |
| 604 | ENSG00000163508 | 282.0954 | -5.28674 | 0.353338 | -14.9623 | 1.30E-50 | 1.01E-47 | 24.65285 | 0.676492 | 26.19091 | 0.5569 | EOMES | 3 |
| 605 | ENSG00000163513 | 67.73364 | 1.60795 | 0.443571 | 3.625009 | 0.000289 | 0.004293 | 1.047498 | 3.031048 | 0.90609 | 2.57637 | TGFBR2 | 3 |
| 606 | ENSG00000163618 | 1907.001 | -1.29954 | 0.153118 | -8.48722 | 2.12E-17 | 2.11E-15 | 149.7494 | 66.84478 | 171.6168 | 56.39376 | CADPS | 3 |
| 607 | ENSG00000163659 | 539.2659 | 1.084089 | 0.22558 | 4.805786 | 1.54E-06 | 3.85E-05 | 10.98171 | 29.14876 | 13.985 | 20.77106 | TIPARP | 3 |
| 608 | ENSG00000163686 | 167.407 | -1.44175 | 0.30359 | -4.74902 | 2.04E-06 | 5.00E-05 | 13.89953 | 5.544048 | 13.72031 | 4.049894 | ABHD6 | 3 |
| 609 | ENSG00000163762 | 119.0794 | 3.725698 | 0.47124 | 7.906159 | 2.65E-15 | 2.14E-13 | 0.604133 | 16.25785 | 1.236124 | 9.36839 | TM4SF18 | 3 |
| 610 | ENSG00000163873 | 214.695 | -1.53969 | 0.316849 | -4.85938 | 1.18E-06 | 3.01E-05 | 5.36715 | 1.911291 | 3.873414 | 1.091429 | GRIK3 | 1 |
| 611 | ENSG00000164023 | 69.0076 | 1.367711 | 0.469898 | 2.910657 | 0.003607 | 0.035298 | 0.649296 | 2.164948 | 1.279398 | 2.515654 | SGMS2 | 4 |
| 612 | ENSG00000164089 | 32.10493 | -2.4988 | 0.663216 | -3.76771 | 0.000165 | 0.002598 | 5.896379 | 1.018228 | 5.608083 | 0.855403 | ETNPPL | 4 |
| 613 | ENSG00000164128 | 87.63971 | 2.211666 | 0.418268 | 5.287679 | 1.24E-07 | 3.85E-06 | 1.868262 | 8.454041 | 1.584627 | 6.509176 | NPY1R | 4 |
| 614 | ENSG00000164129 | 37.50367 | 2.972822 | 0.665301 | 4.468389 | 7.88E-06 | 0.000173 | 0.611 | 5.688168 | 0.633047 | 3.486917 | NPY5R | 4 |
| 615 | ENSG00000164142 | 106.7747 | 2.366883 | 0.381343 | 6.206698 | 5.41E-10 | 2.51E-08 | 0.369205 | 1.782545 | 0.487409 | 2.440928 | FHIP1A | 4 |
| 616 | ENSG00000164161 | 92.98089 | -1.33783 | 0.412881 | -3.24022 | 0.001194 | 0.014452 | 1.697297 | 0.774925 | 3.007551 | 0.980523 | HHIP | 4 |
| 617 | ENSG00000164171 | 211.2187 | -1.58001 | 0.276869 | -5.70672 | 1.15E-08 | 4.28E-07 | 5.3753 | 1.87995 | 5.039448 | 1.400746 | ITGA2 | 5 |
| 618 | ENSG00000164294 | 505.6416 | 1.129617 | 0.204604 | 5.521001 | 3.37E-08 | 1.16E-06 | 11.11684 | 27.05574 | 14.55747 | 26.08201 | GPX8 | 5 |
| 619 | ENSG00000164330 | 1323.559 | 1.576871 | 0.144788 | 10.89087 | 1.27E-27 | 2.86E-25 | 18.89841 | 48.227 | 16.74118 | 52.13554 | EBF1 | 5 |
| 620 | ENSG00000164379 | 28.5714 | 5.763432 | 1.216061 | 4.739426 | 2.14E-06 | 5.22E-05 | 0 | 2.946085 | 0.087657 | 1.773933 | FOXQ1 | 6 |
| 621 | ENSG00000164418 | 477.4238 | -2.03127 | 0.197857 | -10.2663 | 9.99E-25 | 1.90E-22 | 25.42653 | 5.530372 | 22.14123 | 5.445242 | GRIK2 | 6 |
| 622 | ENSG00000164434 | 1153.537 | -3.09819 | 0.165487 | -18.7216 | 3.30E-78 | 5.86E-75 | 405.5776 | 48.85948 | 403.0762 | 40.0834 | FABP7 | 6 |
| 623 | ENSG00000164438 | 20.53401 | 2.780452 | 0.848967 | 3.2751 | 0.001056 | 0.01299 | 0.465797 | 3.058283 | 0.411762 | 2.615793 | TLX3 | 5 |

| | | | | | | | | | | | | | |
|---|---|---|---|---|---|---|---|---|---|---|---|---|---|
| 624 | ENSG00000164483 | 191.7326 | -4.65003 | 0.362891 | -12.8138 | 1.37E-37 | 6.71E-35 | 29.95276 | 0.975374 | 27.76646 | 1.236094 | SAMD3 | 6 |
| 625 | ENSG00000164512 | 12.03639 | -5.90519 | 1.740016 | -3.39376 | 0.000689 | 0.009035 | 1.378351 | 0.073736 | 3.987873 | 0 | ANKRD55 | 5 |
| 626 | ENSG00000164588 | 62.15178 | -1.8664 | 0.540368 | -3.45395 | 0.000552 | 0.007497 | 3.15543 | 0.493498 | 1.202962 | 0.62873 | HCN1 | 5 |
| 627 | ENSG00000164651 | 124.955 | -3.94523 | 0.438025 | -9.00685 | 2.12E-19 | 2.58E-17 | 6.874465 | 0.690945 | 9.770961 | 0.300613 | SP8 | 7 |
| 628 | ENSG00000164690 | 81.10127 | -4.06807 | 0.631509 | -6.44182 | 1.18E-10 | 5.93E-09 | 6.484534 | 0.169895 | 1.512499 | 0.2778 | SHH | 7 |
| 629 | ENSG00000164692 | 1804.913 | 5.268453 | 0.220418 | 23.90215 | 2.91E-126 | 1.90E-122 | 2.942227 | 88.00169 | 3.334261 | 143.58 | COL1A2 | 7 |
| 630 | ENSG00000164756 | 25.81083 | -2.1108 | 0.745236 | -2.83239 | 0.00462 | 0.043 | 0.827213 | 0.242689 | 1.864472 | 0.346893 | SLC30A8 | 8 |
| 631 | ENSG00000164778 | 271.1679 | 3.427852 | 0.906376 | 3.781933 | 0.000156 | 0.002476 | 0.648005 | 12.98512 | 2.485934 | 19.26827 | EN2 | 7 |
| 632 | ENSG00000164853 | 129.8747 | -1.82701 | 0.333283 | -5.48187 | 4.21E-08 | 1.43E-06 | 11.83857 | 2.741743 | 11.88427 | 3.594706 | UNCX | 7 |
| 633 | ENSG00000164946 | 89.83918 | 1.2791 | 0.455479 | 2.808254 | 0.004981 | 0.04545 | 1.827383 | 2.506737 | 0.694906 | 3.084934 | FREM1 | 9 |
| 634 | ENSG00000165023 | 156.5269 | -1.15056 | 0.309291 | -3.71999 | 0.000199 | 0.003082 | 7.030011 | 2.56945 | 5.069672 | 2.562896 | DIRAS2 | 9 |
| 635 | ENSG00000165061 | 416.7231 | 2.441458 | 0.208831 | 11.69106 | 1.42E-31 | 4.40E-29 | 10.18462 | 48.75456 | 9.420235 | 51.82833 | ZMAT4 | 8 |
| 636 | ENSG00000165072 | 348.6312 | 2.524794 | 0.282974 | 8.922353 | 4.56E-19 | 5.31E-17 | 3.20861 | 24.91012 | 4.20278 | 15.23912 | MAMDC2 | 9 |
| 637 | ENSG00000165124 | 22.08546 | 2.972215 | 0.837571 | 3.548611 | 0.000387 | 0.005523 | 0.040156 | 0.402951 | 0.053002 | 0.289751 | SVEP1 | 9 |
| 638 | ENSG00000165186 | 219.3332 | -2.81477 | 0.350263 | -8.03616 | 9.27E-16 | 7.78E-14 | 4.072749 | 0.738562 | 3.626109 | 0.290331 | PTCHD1 | X |
| 639 | ENSG00000165244 | 157.5177 | 1.041485 | 0.307783 | 3.383828 | 0.000715 | 0.009288 | 3.523288 | 6.960386 | 2.950863 | 5.571112 | ZNF367 | 9 |
| 640 | ENSG00000165655 | 365.5935 | -1.37919 | 0.219864 | -6.27295 | 3.54E-10 | 1.67E-08 | 18.49758 | 7.59011 | 20.02282 | 6.375992 | ZNF503 | 10 |
| 641 | ENSG00000165757 | 131.1631 | 1.214525 | 0.324166 | 3.746618 | 0.000179 | 0.002804 | 1.031522 | 2.06961 | 0.849649 | 2.043388 | JCAD | 10 |
| 642 | ENSG00000165966 | 52.9175 | -2.19412 | 0.538449 | -4.07489 | 4.60E-05 | 0.000841 | 3.684369 | 0.687086 | 2.067049 | 0.492818 | PDZRN4 | 12 |
| 643 | ENSG00000166016 | 202.8492 | -1.01689 | 0.267126 | -3.80677 | 0.000141 | 0.00227 | 6.82796 | 2.810262 | 5.763835 | 3.063516 | ABTB2 | 11 |
| 644 | ENSG00000166111 | 116.0914 | -1.23905 | 0.351175 | -3.52828 | 0.000418 | 0.005885 | 2.801742 | 1.182409 | 3.720682 | 1.434573 | SVOP | 12 |
| 645 | ENSG00000166147 | 578.3571 | 1.486503 | 0.226792 | 6.554489 | 5.58E-11 | 2.88E-09 | 6.059076 | 17.80326 | 5.172272 | 11.75149 | FBN1 | 15 |
| 646 | ENSG00000166342 | 100.7776 | 1.707111 | 0.46051 | 3.707003 | 0.00021 | 0.003227 | 2.125086 | 4.082558 | 0.681249 | 4.092462 | NETO1 | 18 |
| 647 | ENSG00000166426 | 3062.742 | 2.879694 | 0.139604 | 20.62766 | 1.55E-94 | 6.06E-91 | 172.1977 | 1011.232 | 152.6691 | 1249.134 | CRABP1 | 15 |
| 648 | ENSG00000166450 | 2796.182 | 2.156186 | 0.440864 | 4.890816 | 1.00E-06 | 2.60E-05 | 13.93841 | 74.72873 | 25.09397 | 90.42462 | PRTG | 15 |
| 649 | ENSG00000166596 | 23.36295 | -2.83528 | 0.785202 | -3.6109 | 0.000305 | 0.004496 | 5.365947 | 0.569476 | 3.404849 | 0.58454 | CFAP52 | 17 |
| 650 | ENSG00000166710 | 717.485 | 1.213942 | 0.175446 | 6.91919 | 4.54E-12 | 2.59E-10 | 70.24071 | 153.2621 | 62.15753 | 136.1219 | B2M | 15 |
| 651 | ENSG00000166770 | 87.14576 | -7.34965 | 0.824511 | -8.91395 | 4.92E-19 | 5.70E-17 | 6.55395 | 0.096854 | 10.37841 | 0.019083 | ZNF667-A[illegible] | 19 |
| 652 | ENSG00000166801 | 128.1803 | 1.350879 | 0.325232 | 4.153588 | 3.27E-05 | 0.000615 | 2.482725 | 5.861032 | 2.602122 | 6.401084 | FAM111A | 11 |
| 653 | ENSG00000166831 | 179.1787 | 1.001096 | 0.28421 | 3.522386 | 0.000428 | 0.005999 | 8.162614 | 13.50072 | 7.916907 | 16.95961 | RBPMS2 | 15 |
| 654 | ENSG00000166888 | 74.48891 | 2.18725 | 0.44721 | 4.890875 | 1.00E-06 | 2.60E-05 | 1.12542 | 4.066101 | 1.226133 | 6.105946 | STAT6 | 12 |
| 655 | ENSG00000166922 | 76.57503 | -1.26579 | 0.420452 | -3.01056 | 0.002608 | 0.027087 | 11.84569 | 5.775994 | 14.76278 | 4.673673 | SCG5 | 15 |
| 656 | ENSG00000166923 | 41.45404 | 3.48215 | 0.651557 | 5.344349 | 9.07E-08 | 2.92E-06 | 0.217857 | 2.176598 | 0.175943 | 1.96507 | GREM1 | 15 |
| 657 | ENSG00000167114 | 278.0135 | 1.019373 | 0.233269 | 4.369947 | 1.24E-05 | 0.000259 | 6.478412 | 12.10904 | 6.809626 | 13.37325 | SLC27A4 | 9 |
| 658 | ENSG00000167123 | 192.3975 | 2.842097 | 0.345037 | 8.23709 | 1.76E-16 | 1.55E-14 | 1.712019 | 12.44839 | 2.8929 | 19.37729 | CERCAM | 9 |
| 659 | ENSG00000167178 | 384.203 | -2.21188 | 0.215279 | -10.2745 | 9.18E-25 | 1.76E-22 | 25.11914 | 5.265984 | 24.50938 | 4.831187 | ISLR2 | 15 |
| 660 | ENSG00000167553 | 410.9474 | 1.495074 | 0.226573 | 6.598654 | 4.15E-11 | 2.19E-09 | 23.2618 | 78.03681 | 31.61672 | 68.77111 | TUBA1C | 12 |
| 661 | ENSG00000167601 | 344.438 | 1.179419 | 0.235423 | 5.00979 | 5.45E-07 | 1.50E-05 | 4.988918 | 12.88503 | 5.752646 | 10.08683 | AXL | 19 |
| 662 | ENSG00000167785 | 300.1587 | 1.36954 | 0.230966 | 5.929623 | 3.04E-09 | 1.24E-07 | 6.368256 | 16.21584 | 7.30974 | 17.23241 | ZNF558 | 19 |
| 663 | ENSG00000167912 | 98.15393 | -1.66882 | 0.375409 | -4.44535 | 8.77E-06 | 0.000189 | 15.24541 | 3.849211 | 14.47007 | 5.032483 | TOX-DT | 8 |
| 664 | ENSG00000168077 | 136.2434 | 1.098168 | 0.3235 | 3.394643 | 0.000687 | 0.009012 | 2.428664 | 5.774222 | 3.076473 | 5.386504 | SCARA3 | 8 |
| 665 | ENSG00000168079 | 14.85387 | 4.632341 | 1.390299 | 3.331902 | 0.000863 | 0.01091 | 0.089307 | 1.584084 | 0 | 0.527546 | SCARA5 | 8 |
| 666 | ENSG00000168264 | 1170.422 | 1.036189 | 0.157397 | 6.583288 | 4.60E-11 | 2.41E-09 | 26.88682 | 45.20429 | 25.85941 | 57.17182 | IRF2BP2 | 1 |
| 667 | ENSG00000168505 | 60.46801 | 2.817256 | 0.547017 | 5.15022 | 2.60E-07 | 7.69E-06 | 0.633649 | 7.5061 | 1.644419 | 6.873483 | GBX2 | 2 |
| 668 | ENSG00000168542 | 560.8772 | 4.608042 | 0.254659 | 18.09496 | 3.49E-73 | 4.55E-70 | 0.961463 | 17.62988 | 0.907112 | 25.53589 | COL3A1 | 2 |
| 669 | ENSG00000168646 | 666.3849 | 1.6041 | 0.172564 | 9.295661 | 1.46E-20 | 1.92E-18 | 14.6191 | 41.84903 | 14.16381 | 40.7478 | AXIN2 | 17 |
| 670 | ENSG00000168672 | 1500.396 | 1.541833 | 0.161725 | 9.533671 | 1.52E-21 | 2.12E-19 | 20.24837 | 58.63152 | 18.25295 | 46.92061 | LRATD2 | 8 |
| 671 | ENSG00000168675 | 978.0663 | 1.039939 | 0.177365 | 5.863278 | 4.54E-09 | 1.81E-07 | 15.07789 | 35.24863 | 18.15868 | 29.23585 | LDLRAD4 | 18 |
| 672 | ENSG00000168702 | 328.2892 | -2.01543 | 0.653751 | -3.08287 | 0.00205 | 0.02256 | 5.766319 | 1.579614 | 3.920388 | 0.666167 | LRP1B | 2 |
| 673 | ENSG00000168843 | 425.8857 | -2.08028 | 0.245569 | -8.47125 | 2.43E-17 | 2.37E-15 | 22.07746 | 5.433045 | 17.99799 | 3.421115 | FSTL5 | 4 |
| 674 | ENSG00000168874 | 172.7984 | 1.375937 | 0.311356 | 4.419178 | 9.91E-06 | 0.00021 | 2.628215 | 8.548338 | 3.599723 | 6.732413 | ATOH8 | 2 |
| 675 | ENSG00000168959 | 34.77208 | -1.77312 | 0.633635 | -2.79834 | 0.005137 | 0.046566 | 0.967326 | 0.350375 | 1.386065 | 0.300355 | GRM5 | 11 |
| 676 | ENSG00000168994 | 58.34601 | 1.645268 | 0.480357 | 3.425094 | 0.000615 | 0.008211 | 2.333936 | 5.431268 | 1.688226 | 6.409837 | PXDC1 | 6 |
| 677 | ENSG00000169085 | 83.56949 | -1.24962 | 0.424408 | -2.94437 | 0.003236 | 0.03217 | 5.959557 | 2.376319 | 4.098899 | 1.585122 | VXN | 8 |
| 678 | ENSG00000169184 | 1326.206 | 1.264775 | 0.141375 | 8.946214 | 3.68E-19 | 4.31E-17 | 11.29446 | 25.34703 | 10.93309 | 25.07797 | MN1 | 22 |
| 679 | ENSG00000169218 | 150.3335 | -1.67837 | 0.309811 | -5.41741 | 6.05E-08 | 2.00E-06 | 9.57487 | 3.128278 | 10.11493 | 2.672794 | RSPO1 | 1 |
| 680 | ENSG00000169242 | 33.83962 | -2.16145 | 0.657316 | -3.2883 | 0.001008 | 0.012435 | 6.79949 | 1.441005 | 4.373443 | 0.90223 | EFNA1 | 1 |
| 681 | ENSG00000169427 | 55.6002 | -2.35129 | 0.527834 | -4.45461 | 8.40E-06 | 0.000182 | 3.477873 | 0.477772 | 2.249686 | 0.590376 | KCNK9 | 8 |
| 682 | ENSG00000169436 | 95.65154 | -1.30919 | 0.418644 | -3.12721 | 0.001765 | 0.019935 | 5.928985 | 2.589486 | 4.366769 | 1.389323 | COL22A1 | 8 |
| 683 | ENSG00000169439 | 2533.86 | 1.202008 | 0.163206 | 7.364956 | 1.77E-13 | 1.17E-11 | 80.82434 | 219.4141 | 103.917 | 181.9154 | SDC2 | 8 |
| 684 | ENSG00000169744 | 274.2431 | 1.38506 | 0.254296 | 5.446649 | 5.13E-08 | 1.72E-06 | 10.76465 | 20.36017 | 8.321189 | 26.86547 | LDB2 | 4 |
| 685 | ENSG00000169836 | 117.8694 | 1.469605 | 0.351987 | 4.17517 | 2.98E-05 | 0.000564 | 1.338269 | 3.895593 | 1.302528 | 2.990745 | TACR3 | 4 |
| 686 | ENSG00000169855 | 2155.151 | -1.17871 | 0.163285 | -7.21874 | 5.25E-13 | 3.30E-11 | 89.23341 | 33.81148 | 64.46131 | 30.08561 | ROBO1 | 3 |
| 687 | ENSG00000169856 | 553.2019 | -1.01917 | 0.190616 | -5.34672 | 8.96E-08 | 2.89E-06 | 32.5619 | 12.70372 | 27.09271 | 15.13838 | ONECUT1 | 15 |
| 688 | ENSG00000169884 | 26.73713 | 3.765067 | 0.836424 | 4.501385 | 6.75E-06 | 0.00015 | 0.240032 | 2.528722 | 0.158849 | 2.550586 | WNT10B | 12 |
| 689 | ENSG00000169908 | 25.77338 | 3.953671 | 0.880722 | 4.489127 | 7.15E-06 | 0.000158 | 0.183332 | 4.098747 | 0.324101 | 3.403992 | TM4SF1 | 3 |
| 690 | ENSG00000170091 | 820.9819 | -1.71783 | 0.198285 | -8.66342 | 4.58E-18 | 4.84E-16 | 74.57123 | 22.44118 | 60.35592 | 16.14623 | NSG2 | 5 |
| 691 | ENSG00000170275 | 599.4393 | 1.04974 | 0.254984 | 4.116883 | 3.84E-05 | 0.000712 | 11.44292 | 30.39633 | 22.04469 | 35.19371 | CRTAP | 3 |
| 692 | ENSG00000170370 | 257.0835 | -3.12169 | 0.340416 | -9.17022 | 4.72E-20 | 5.99E-18 | 27.60959 | 4.279448 | 26.10975 | 1.624619 | EMX2 | 10 |
| 693 | ENSG00000170382 | 277.5067 | -1.09935 | 0.235382 | -4.67049 | 3.00E-06 | 7.11E-05 | 12.23724 | 5.559735 | 11.68172 | 4.968488 | LRRN2 | 1 |
| 694 | ENSG00000170396 | 113.818 | -1.51095 | 0.376358 | -4.01467 | 5.95E-05 | 0.001059 | 4.935131 | 1.704129 | 3.581304 | 1.095515 | ZNF804A | 2 |
| 695 | ENSG00000170421 | 103.6705 | 4.493963 | 0.522675 | 8.598011 | 8.11E-18 | 8.39E-16 | 1.141346 | 17.40481 | 0.486111 | 14.43074 | KRT8 | 12 |
| 696 | ENSG00000170525 | 588.9462 | 1.01162 | 0.176456 | 5.732985 | 9.87E-09 | 3.71E-07 | 12.48069 | 22.26187 | 12.08468 | 24.55817 | PFKFB3 | 10 |
| 697 | ENSG00000170542 | 69.0754 | 2.012708 | 0.484774 | 4.151847 | 3.30E-05 | 0.000619 | 1.016499 | 3.360478 | 0.515584 | 2.40707 | SERPINB9 | 6 |
| 698 | ENSG00000170549 | 155.1894 | 2.413722 | 0.316889 | 7.61692 | 2.60E-14 | 1.91E-12 | 3.010104 | 13.58685 | 2.772353 | 15.52226 | IRX1 | 5 |
| 699 | ENSG00000170561 | 378.7731 | 3.070128 | 0.244298 | 12.56712 | 3.20E-36 | 1.46E-33 | 2.825883 | 22.75366 | 3.660946 | 29.11177 | IRX2 | 5 |
| 700 | ENSG00000170624 | 41.7685 | -3.12842 | 0.73284 | -4.2689 | 1.96E-05 | 0.00039 | 1.608702 | 0.352094 | 1.67337 | 0.04462 | SGCD | 5 |
| 701 | ENSG00000170775 | 123.2045 | -1.36187 | 0.369955 | -3.68117 | 0.000232 | 0.003527 | 3.255374 | 1.72145 | 4.310545 | 1.049316 | GPR37 | 7 |

| | | | | | | | | | | | | | | |
|---|---|---|---|---|---|---|---|---|---|---|---|---|---|---|
| 702 | ENSG00000170893 | 314.9816 | -3.23236 | 0.328797 | -9.83086 | 8.29E-23 | 1.31E-20 | 31.54442 | 5.638222 | 61.68665 | 3.7249 | TRH | | 3 |
| 703 | ENSG00000170921 | 1996.533 | -1.01097 | 0.154582 | -6.54006 | 6.15E-11 | 3.16E-09 | 58.03221 | 30.00258 | 56.27842 | 23.4827 | TANC2 | | 17 |
| 704 | ENSG00000170959 | 773.1465 | -3.56452 | 0.18736 | -19.0249 | 1.06E-80 | 2.59E-77 | 52.5313 | 3.574701 | 43.29103 | 4.081955 | DCDC1 | | 11 |
| 705 | ENSG00000171004 | 831.9974 | -1.00235 | 0.17145 | -5.8463 | 5.03E-09 | 1.99E-07 | 33.43617 | 14.20441 | 26.52886 | 14.01176 | HS6ST2 | X | |
| 706 | ENSG00000171189 | 169.9627 | -2.8782 | 0.326585 | -8.81302 | 1.22E-18 | 1.35E-16 | 14.14445 | 1.776436 | 10.50122 | 1.373501 | GRIK1 | | 21 |
| 707 | ENSG00000171246 | 356.6082 | -1.26409 | 0.293164 | -4.3119 | 1.62E-05 | 0.000327 | 16.21279 | 3.772665 | 8.738436 | 6.062969 | NPTX1 | | 17 |
| 708 | ENSG00000171345 | 23.48227 | 2.669709 | 0.81863 | 3.26119 | 0.001109 | 0.013559 | 0.968478 | 4.8971 | 0.463865 | 3.302852 | KRT19 | | 17 |
| 709 | ENSG00000171388 | 216.8692 | 1.62651 | 0.290608 | 5.596923 | 2.18E-08 | 7.79E-07 | 3.264796 | 12.72645 | 4.405282 | 9.657018 | APLN | X | |
| 710 | ENSG00000171451 | 1061.06 | 2.113287 | 0.17138 | 12.33098 | 6.17E-35 | 2.68E-32 | 4.724933 | 21.46138 | 4.802703 | 17.40141 | DSEL | | 18 |
| 711 | ENSG00000171502 | 428.1819 | 2.022548 | 0.240254 | 8.418371 | 3.82E-17 | 3.62E-15 | 2.795556 | 14.19067 | 4.484908 | 13.66109 | COL24A1 | | 1 |
| 712 | ENSG00000171587 | 87.49418 | -1.20947 | 0.423443 | -2.85628 | 0.004286 | 0.040687 | 1.990639 | 0.914539 | 1.524943 | 0.514216 | DSCAM | | 21 |
| 713 | ENSG00000171621 | 59.89368 | 1.293507 | 0.466881 | 2.770528 | 0.005597 | 0.049993 | 2.13898 | 4.805282 | 1.773856 | 4.177174 | SPSB1 | | 1 |
| 714 | ENSG00000171757 | 70.30859 | -1.44546 | 0.476257 | -3.03503 | 0.002405 | 0.025536 | 3.226698 | 1.592646 | 6.522972 | 1.789331 | LRRC34 | | 3 |
| 715 | ENSG00000171766 | 309.6736 | -1.0952 | 0.256785 | -4.26506 | 2.00E-05 | 0.000395 | 20.68382 | 10.68238 | 19.44908 | 6.962758 | GATM | | 15 |
| 716 | ENSG00000171823 | 121.7114 | -1.00491 | 0.329062 | -3.05387 | 0.002259 | 0.024365 | 7.629618 | 3.488958 | 7.750458 | 3.756522 | FBXL14 | | 12 |
| 717 | ENSG00000171864 | 65.31755 | 3.358856 | 0.56142 | 5.982782 | 2.19E-09 | 9.22E-08 | 0.482081 | 4.057442 | 0.198204 | 2.402346 | PRND | | 20 |
| 718 | ENSG00000171951 | 1428.897 | -1.49329 | 0.153069 | -9.75567 | 1.74E-22 | 2.62E-20 | 98.13336 | 37.0179 | 109.8887 | 32.72972 | SCG2 | | 2 |
| 719 | ENSG00000172137 | 55.95255 | -1.76092 | 0.517715 | -3.40134 | 0.000671 | 0.008836 | 10.81078 | 3.321865 | 7.640179 | 1.859593 | CALB2 | | 16 |
| 720 | ENSG00000172164 | 99.53832 | 1.164357 | 0.376464 | 3.092877 | 0.001982 | 0.021887 | 1.329554 | 3.514605 | 1.685819 | 2.86282 | SNTB1 | | 8 |
| 721 | ENSG00000172216 | 43.04017 | 1.956263 | 0.559962 | 3.493561 | 0.000477 | 0.006549 | 1.054646 | 4.35345 | 1.330545 | 4.420531 | CEBPB | | 20 |
| 722 | ENSG00000172238 | 43.93255 | 5.789756 | 0.976428 | 5.929527 | 3.04E-09 | 1.24E-07 | 0.061045 | 4.144689 | 0.107736 | 4.82624 | ATOH1 | | 4 |
| 723 | ENSG00000172318 | 429.7565 | 1.433292 | 0.220604 | 6.497125 | 8.19E-11 | 4.15E-09 | 5.670399 | 14.71821 | 4.688406 | 11.58683 | B3GALT1 | | 2 |
| 724 | ENSG00000172461 | 1215.787 | -1.55124 | 0.165785 | -9.35694 | 8.21E-21 | 1.09E-18 | 18.97594 | 5.976294 | 15.42466 | 5.074625 | FUT9 | | 6 |
| 725 | ENSG00000172575 | 244.3014 | -1.80457 | 0.250902 | -7.19234 | 6.37E-13 | 3.95E-11 | 9.315211 | 2.384388 | 8.274209 | 2.364938 | RASGRP1 | | 15 |
| 726 | ENSG00000172716 | 21.14544 | 3.608926 | 1.030582 | 3.501832 | 0.000462 | 0.006399 | 0.38268 | 1.297777 | 0.068004 | 1.813845 | SLFN11 | | 17 |
| 727 | ENSG00000172965 | 701.7418 | 2.089383 | 0.204382 | 10.22295 | 1.57E-24 | 2.94E-22 | 16.76623 | 86.83433 | 24.16207 | 78.11808 | MIR4435-2 | | 2 |
| 728 | ENSG00000173041 | 213.6947 | -1.31437 | 0.276782 | -4.74877 | 2.05E-06 | 5.00E-05 | 13.4369 | 5.156068 | 17.09136 | 6.487505 | ZNF680 | | 7 |
| 729 | ENSG00000173157 | 84.12813 | 2.030421 | 0.443004 | 4.583304 | 4.58E-06 | 0.000105 | 1.116444 | 3.299437 | 0.53079 | 2.950705 | ADAMTS2( | | 12 |
| 730 | ENSG00000173208 | 121.1884 | -1.97568 | 0.362824 | -5.44529 | 5.17E-08 | 1.73E-06 | 3.833432 | 0.990055 | 3.077758 | 0.657765 | ABCD2 | | 12 |
| 731 | ENSG00000173482 | 1419.92 | -1.33736 | 0.178034 | -7.51185 | 5.83E-14 | 4.06E-12 | 68.59361 | 29.28133 | 65.32273 | 20.62351 | PTPRM | | 18 |
| 732 | ENSG00000173530 | 307.957 | 1.171309 | 0.237666 | 4.928379 | 8.29E-07 | 2.18E-05 | 5.656849 | 14.22373 | 6.753857 | 12.18463 | TNFRSF10I | | 8 |
| 733 | ENSG00000173706 | 1697.245 | 1.875543 | 0.418589 | 4.480633 | 7.44E-06 | 0.000164 | 11.17776 | 57.15657 | 18.93893 | 47.34242 | HEG1 | | 3 |
| 734 | ENSG00000173805 | 218.4343 | -1.36852 | 0.316195 | -4.3281 | 1.50E-05 | 0.000306 | 9.976056 | 5.180721 | 11.84655 | 2.792656 | HAP1 | | 17 |
| 735 | ENSG00000173947 | 77.33854 | -1.52473 | 0.441948 | -3.45001 | 0.000561 | 0.007591 | 7.207162 | 1.823988 | 8.363646 | 3.418164 | PIFO | | 1 |
| 736 | ENSG00000174080 | 10.12049 | -6.87597 | 1.800818 | -3.81825 | 0.000134 | 0.002186 | 0.961967 | 0 | 1.573642 | 0 | CTSF | | 11 |
| 737 | ENSG00000174680 | 34.37894 | -2.81916 | 0.663601 | -4.24828 | 2.15E-05 | 0.000423 | 2.987876 | 0.259569 | 1.617789 | 0.354329 | GRIK1-AS1 | | 21 |
| 738 | ENSG00000174721 | 1189.471 | 1.32125 | 0.156604 | 8.436885 | 3.26E-17 | 3.15E-15 | 29.75845 | 69.11277 | 33.81766 | 81.35733 | FGFBP3 | | 10 |
| 739 | ENSG00000174807 | 19.03507 | 2.654496 | 0.883469 | 3.004627 | 0.002659 | 0.027438 | 0.260352 | 1.787203 | 0.229627 | 1.106359 | CD248 | | 11 |
| 740 | ENSG00000174844 | 18.71181 | -2.75387 | 0.983851 | -2.79907 | 0.005125 | 0.046481 | 1.0866 | 0.272574 | 0.723811 | 0.043856 | DNAH12 | | 3 |
| 741 | ENSG00000174948 | 31.86614 | -2.99534 | 0.755815 | -3.96306 | 7.40E-05 | 0.001287 | 1.11385 | 0.230188 | 1.200996 | 0.038607 | GPR149 | | 3 |
| 742 | ENSG00000175093 | 283.1074 | 1.545275 | 0.240046 | 6.437407 | 1.22E-10 | 6.09E-09 | 10.11946 | 24.25262 | 7.974991 | 25.58343 | SPSB4 | | 3 |
| 743 | ENSG00000175161 | 1385.067 | -2.21166 | 0.199901 | -11.0638 | 1.88E-28 | 4.61E-26 | 74.26716 | 13.92728 | 48.18586 | 10.89058 | CADM2 | | 3 |
| 744 | ENSG00000175356 | 108.2963 | -1.25328 | 0.439055 | -2.8545 | 0.00431 | 0.040836 | 14.91745 | 4.637949 | 5.89496 | 3.508874 | SCUBE2 | | 11 |
| 745 | ENSG00000175426 | 311.9289 | -1.2618 | 0.250103 | -5.04511 | 4.53E-07 | 1.28E-05 | 11.59487 | 4.60064 | 8.921266 | 3.447687 | PCSK1 | | 5 |
| 746 | ENSG00000175445 | 180.0512 | 1.698581 | 0.290676 | 5.843563 | 5.11E-09 | 2.01E-07 | 2.980863 | 7.977754 | 2.84959 | 9.92375 | LPL | | 8 |
| 747 | ENSG00000176134 | 12.7717 | -4.03119 | 1.238134 | -3.25586 | 0.00113 | 0.013764 | 2.02681 | 0.159152 | 1.394576 | 0 | | | 9 |
| 748 | ENSG00000176165 | 642.4046 | -4.79588 | 0.62206 | -7.70967 | 1.26E-14 | 9.51E-13 | 51.15791 | 1.90303 | 31.58755 | 0.879896 | FOXG1 | | 14 |
| 749 | ENSG00000176170 | 45.77126 | 2.775628 | 0.579636 | 4.788573 | 1.68E-06 | 4.17E-05 | 0.993653 | 4.743861 | 0.553219 | 5.00584 | SPHK1 | | 17 |
| 750 | ENSG00000176204 | 236.0432 | -1.56119 | 0.288483 | -5.41173 | 6.24E-08 | 2.05E-06 | 13.31277 | 5.37389 | 13.61097 | 3.250396 | LRRTM4 | | 2 |
| 751 | ENSG00000176293 | 49.14112 | -9.02192 | 1.52409 | -5.91955 | 3.23E-09 | 1.32E-07 | 4.200693 | 0 | 3.343835 | 0 | ZNF135 | | 19 |
| 752 | ENSG00000176399 | 310.0264 | -2.96717 | 0.249168 | -11.9083 | 1.07E-32 | 3.67E-30 | 12.02707 | 1.4359 | 10.21433 | 1.240871 | DMRTA1 | | 9 |
| 753 | ENSG00000176720 | 121.4063 | 1.161346 | 0.359638 | 3.229212 | 0.001241 | 0.014828 | 2.874513 | 8.299799 | 3.913774 | 6.043019 | BOK | | 2 |
| 754 | ENSG00000176771 | 995.06 | 1.349348 | 0.153255 | 8.804569 | 1.31E-18 | 1.45E-16 | 11.87023 | 28.89855 | 12.66931 | 30.21863 | NCKAP5 | | 2 |
| 755 | ENSG00000176842 | 334.0799 | 2.632828 | 0.266441 | 9.881478 | 5.01E-23 | 8.16E-21 | 3.569346 | 26.95106 | 6.177254 | 30.52552 | IRX5 | | 16 |
| 756 | ENSG00000176896 | 74.52374 | -1.42264 | 0.420469 | -3.38346 | 0.000716 | 0.009294 | 7.096911 | 2.173663 | 5.85685 | 2.384173 | TCEANC | X | |
| 757 | ENSG00000176907 | 122.5009 | -2.02271 | 0.35761 | -5.6562 | 1.55E-08 | 5.65E-07 | 15.99538 | 3.137772 | 10.9127 | 3.094609 | TCIM | | 8 |
| 758 | ENSG00000176928 | 105.348 | 1.1185 | 0.354059 | 3.159081 | 0.001583 | 0.018247 | 1.407254 | 2.897834 | 1.498965 | 3.071875 | GCNT4 | | 5 |
| 759 | ENSG00000177103 | 24.39884 | -2.353 | 0.829848 | -2.83546 | 0.004576 | 0.042691 | 1.748991 | 0.261219 | 0.527702 | 0.139251 | DSCAML1 | | 11 |
| 760 | ENSG00000177182 | 142.3166 | -1.78775 | 0.388293 | -4.60411 | 4.14E-06 | 9.57E-05 | 15.41414 | 4.918106 | 10.75947 | 2.231323 | CLVS1 | | 8 |
| 761 | ENSG00000177283 | 50.85692 | -2.23263 | 0.582314 | -3.83406 | 0.000126 | 0.002073 | 3.250258 | 0.319683 | 1.529025 | 0.643293 | FZD8 | | 10 |
| 762 | ENSG00000177469 | 460.6024 | 2.47109 | 0.249038 | 9.922524 | 3.32E-23 | 5.65E-21 | 3.827831 | 28.11933 | 5.28245 | 19.60288 | CAVIN1 | | 17 |
| 763 | ENSG00000177570 | 530.4961 | -1.1813 | 0.202809 | -5.82469 | 5.72E-09 | 2.22E-07 | 19.44973 | 8.176718 | 16.14768 | 6.569362 | SAMD12 | | 8 |
| 764 | ENSG00000177606 | 1368.968 | 1.278157 | 0.152825 | 8.363516 | 6.09E-17 | 5.67E-15 | 32.20602 | 65.8009 | 25.60144 | 66.3598 | JUN | | 1 |
| 765 | ENSG00000177807 | 88.29913 | -1.61555 | 0.44342 | -3.6434 | 0.000269 | 0.004016 | 4.846184 | 1.452695 | 2.716718 | 0.863528 | KCNJ10 | | 1 |
| 766 | ENSG00000177822 | 237.2665 | 1.172355 | 0.25119 | 4.667199 | 3.05E-06 | 7.20E-05 | 10.64282 | 24.1519 | 11.15324 | 22.22297 | TENM3-AS | | 4 |
| 767 | ENSG00000177873 | 20.69833 | -2.74953 | 0.846498 | -3.24812 | 0.001162 | 0.014109 | 0.867382 | 0.101766 | 1.139758 | 0.189303 | ZNF619 | | 3 |
| 768 | ENSG00000177932 | 46.92735 | -2.85122 | 0.587923 | -4.84965 | 1.24E-06 | 3.14E-05 | 1.636906 | 0.281488 | 1.516152 | 0.126364 | ZNF354C | | 5 |
| 769 | ENSG00000178031 | 718.7948 | -2.81823 | 0.223678 | -12.5995 | 2.12E-36 | 9.89E-34 | 34.61251 | 5.667466 | 32.9894 | 3.326522 | ADAMTSL' | | 9 |
| 770 | ENSG00000178033 | 221.7885 | 2.648828 | 0.294734 | 8.987194 | 2.54E-19 | 3.06E-17 | 0.868223 | 3.889265 | 0.520472 | 4.283652 | CALHM5 | | 6 |
| 771 | ENSG00000178171 | 83.74845 | -1.15337 | 0.392883 | -2.93566 | 0.003328 | 0.033004 | 3.019698 | 1.352739 | 2.918388 | 1.169719 | AMER3 | | 2 |
| 772 | ENSG00000178187 | 25.07329 | -3.66485 | 0.868857 | -4.21801 | 2.46E-05 | 0.000477 | 2.490505 | 0.31425 | 2.498957 | 0.048906 | ZNF454 | | 5 |
| 773 | ENSG00000178343 | 46.38827 | -3.37682 | 0.60093 | -5.61933 | 1.92E-08 | 6.89E-07 | 4.104649 | 0.360877 | 4.692521 | 0.442278 | SHISA3 | | 4 |
| 774 | ENSG00000178403 | 363.1338 | -3.89421 | 0.295878 | -13.1615 | 1.46E-39 | 7.94E-37 | 42.34861 | 2.970862 | 29.56078 | 1.543499 | NEUROG2 | | 4 |
| 775 | ENSG00000178522 | 17.25745 | -5.12198 | 1.388803 | -3.68806 | 0.000226 | 0.003447 | 3.547745 | 0.107013 | 0.722738 | 0 | AMBN | | 4 |
| 776 | ENSG00000178538 | 230.6612 | -2.66002 | 0.280449 | -9.48483 | 2.43E-21 | 3.34E-19 | 16.95214 | 2.604879 | 13.99955 | 2.006834 | CA8 | | 8 |
| 777 | ENSG00000178597 | 70.6193 | -1.94897 | 0.460018 | -4.23671 | 2.27E-05 | 0.000443 | 2.86464 | 0.85025 | 2.643276 | 0.486593 | PSAPL1 | | 4 |
| 778 | ENSG00000178878 | 352.6908 | 1.790691 | 0.247146 | 7.245467 | 4.31E-13 | 2.74E-11 | 3.402907 | 13.7774 | 5.278799 | 14.6141 | APOLD1 | | 12 |
| 779 | ENSG00000179455 | 389.4485 | -1.37418 | 0.22518 | -6.10259 | 1.04E-09 | 4.63E-08 | 40.03137 | 11.85287 | 29.03455 | 13.26697 | MKRN3 | | 15 |

| | | | | | | | | | | | | | | |
|---|---|---|---|---|---|---|---|---|---|---|---|---|---|---|
| 780 | ENSG00000179604 | 921.7003 | -1.08701 | 0.153381 | -7.08698 | 1.37E-12 | 8.30E-11 | 53.2459 | 23.12494 | 52.56918 | 23.93112 | CDC42EP4 | | 17 |
| 781 | ENSG00000179915 | 955.6513 | -1.44735 | 0.174985 | -8.27127 | 1.33E-16 | 1.19E-14 | 43.34919 | 17.70739 | 50.06623 | 14.59541 | NRXN1 | | 2 |
| 782 | ENSG00000180071 | 89.30085 | 1.932173 | 0.40797 | 4.73607 | 2.18E-06 | 5.30E-05 | 1.415902 | 5.142544 | 1.083319 | 3.844932 | ANKRD18A | | 9 |
| 783 | ENSG00000180613 | 20.00939 | -3.57215 | 1.011567 | -3.5313 | 0.000414 | 0.00583 | 5.465858 | 0.393647 | 2.172044 | 0.218549 | GSX2 | | 4 |
| 784 | ENSG00000180616 | 266.1913 | -1.3396 | 0.242202 | -5.5309 | 3.19E-08 | 1.10E-06 | 5.148039 | 2.023239 | 5.909445 | 2.110098 | SSTR2 | | 17 |
| 785 | ENSG00000180628 | 886.0695 | 1.649948 | 0.160166 | 10.30151 | 6.94E-25 | 1.34E-22 | 13.45539 | 41.29531 | 13.94088 | 39.89901 | PCGF5 | | 10 |
| 786 | ENSG00000180730 | 143.8113 | -2.22299 | 0.377996 | -5.881 | 4.08E-09 | 1.63E-07 | 9.538938 | 1.507635 | 4.731232 | 1.359922 | SHISA2 | | 13 |
| 787 | ENSG00000180828 | 41.63264 | -2.94049 | 0.605335 | -4.85762 | 1.19E-06 | 3.03E-05 | 2.784966 | 0.279148 | 2.48971 | 0.371819 | BHLHE22 | | 8 |
| 788 | ENSG00000181007 | 424.0077 | -1.56105 | 0.200562 | -7.78339 | 7.06E-15 | 5.44E-13 | 16.32648 | 4.941511 | 14.81539 | 5.018229 | ZFP82 | | 19 |
| 789 | ENSG00000181215 | 54.53926 | -2.38138 | 0.528669 | -4.50448 | 6.65E-06 | 0.000148 | 0.798006 | 0.186927 | 0.783624 | 0.097324 | C4orf50 | | 4 |
| 790 | ENSG00000181234 | 545.1554 | 2.57242 | 0.23585 | 10.907 | 1.07E-27 | 2.46E-25 | 2.654126 | 21.32088 | 4.181464 | 17.20475 | TMEM132C | | 12 |
| 791 | ENSG00000181965 | 180.2695 | -3.55344 | 0.339451 | -10.4682 | 1.21E-25 | 2.41E-23 | 21.62787 | 1.696227 | 28.25726 | 2.344086 | NEUROG1 | | 5 |
| 792 | ENSG00000182050 | 264.4469 | -2.09113 | 0.687894 | -3.0399 | 0.002367 | 0.025208 | 8.203015 | 2.309474 | 6.348889 | 0.889375 | MGAT4C | | 12 |
| 793 | ENSG00000182103 | 120.037 | -1.21893 | 0.351827 | -3.46458 | 0.000531 | 0.007221 | 4.111106 | 2.110499 | 5.720556 | 1.884634 | FAM181B | | 11 |
| 794 | ENSG00000182158 | 1760.696 | 1.074725 | 0.13559 | 7.926307 | 2.26E-15 | 1.84E-13 | 19.72202 | 38.53767 | 18.24209 | 36.90349 | CREB3L2 | | 7 |
| 795 | ENSG00000182168 | 910.8442 | 1.148389 | 0.180048 | 6.378228 | 1.79E-10 | 8.71E-09 | 9.668698 | 21.67935 | 8.759132 | 16.79058 | UNC5C | | 4 |
| 796 | ENSG00000182199 | 915.4808 | 1.008991 | 0.16389 | 6.156514 | 7.44E-10 | 3.39E-08 | 43.48578 | 70.60957 | 38.56445 | 85.55541 | SHMT2 | | 12 |
| 797 | ENSG00000182240 | 199.5562 | 1.463699 | 0.283107 | 5.170132 | 2.34E-07 | 6.96E-06 | 2.82958 | 8.76213 | 3.237256 | 7.064911 | BACE2 | | 21 |
| 798 | ENSG00000182601 | 107.0352 | -1.22455 | 0.371711 | -3.29437 | 0.000986 | 0.012208 | 4.608978 | 2.43656 | 6.507055 | 2.059171 | HS3ST4 | | 16 |
| 799 | ENSG00000182667 | 176.351 | -1.88887 | 0.319543 | -5.91117 | 3.40E-09 | 1.38E-07 | 25.31186 | 4.45244 | 17.28533 | 6.497758 | NTM | | 11 |
| 800 | ENSG00000182718 | 1298.764 | 1.55081 | 0.208483 | 7.438542 | 1.02E-13 | 6.89E-12 | 47.76777 | 188.4515 | 68.28322 | 132.2665 | ANXA2 | | 15 |
| 801 | ENSG00000183023 | 967.3907 | -1.34788 | 0.174288 | -7.73361 | 1.05E-14 | 7.92E-13 | 35.3201 | 11.33508 | 26.34695 | 11.49616 | SLC8A1 | | 2 |
| 802 | ENSG00000183036 | 35.88918 | 2.355269 | 0.654533 | 3.598399 | 0.00032 | 0.004693 | 6.225227 | 23.4281 | 2.612171 | 18.71138 | PCP4 | | 21 |
| 803 | ENSG00000183117 | 127.7538 | -1.31656 | 0.330733 | -3.98075 | 6.87E-05 | 0.001203 | 2.894924 | 0.978526 | 2.392098 | 1.024212 | CSMD1 | | 8 |
| 804 | ENSG00000183454 | 589.6543 | -1.31769 | 0.249028 | -5.29134 | 1.21E-07 | 3.78E-06 | 11.45084 | 5.302876 | 10.21855 | 2.879825 | GRIN2A | | 16 |
| 805 | ENSG00000183579 | 877.2046 | 1.451567 | 0.16412 | 8.844566 | 9.19E-19 | 1.03E-16 | 7.192817 | 20.13467 | 8.086602 | 19.36818 | ZNRF3 | | 22 |
| 806 | ENSG00000183690 | 23.28926 | -2.57647 | 0.779799 | -3.30401 | 0.000953 | 0.011863 | 1.275664 | 0.248627 | 1.581136 | 0.203215 | EFHC2 | X | |
| 807 | ENSG00000183696 | 138.591 | 1.047273 | 0.341522 | 3.066491 | 0.002166 | 0.023607 | 6.533374 | 16.57385 | 7.699475 | 11.0842 | UPP1 | | 7 |
| 808 | ENSG00000183762 | 417.9267 | 2.309659 | 0.214172 | 10.78411 | 4.09E-27 | 8.70E-25 | 3.142728 | 15.68789 | 3.180351 | 13.88348 | KREMEN1 | | 22 |
| 809 | ENSG00000183873 | 15.71517 | -2.59629 | 0.914052 | -2.84042 | 0.004505 | 0.042173 | 0.643755 | 0.082782 | 0.534412 | 0.101392 | SCN5A | | 3 |
| 810 | ENSG00000184156 | 1019.739 | -1.96757 | 0.186021 | -10.5772 | 3.80E-26 | 7.75E-24 | 20.6765 | 4.417947 | 14.4892 | 4.04458 | KCNQ3 | | 8 |
| 811 | ENSG00000184185 | 42.14332 | 1.702804 | 0.600469 | 2.83579 | 0.004571 | 0.042666 | 0.49586 | 0.857416 | 0.37928 | 1.841226 | KCNJ12 | | 17 |
| 812 | ENSG00000184271 | 128.5235 | -1.41367 | 0.374162 | -3.77823 | 0.000158 | 0.002506 | 10.13863 | 2.656615 | 5.215492 | 2.739204 | POU6F1 | | 12 |
| 813 | ENSG00000184305 | 425.5924 | -1.60589 | 0.204598 | -7.849 | 4.19E-15 | 3.25E-13 | 31.08879 | 8.416109 | 26.72606 | 9.527469 | CCSER1 | | 4 |
| 814 | ENSG00000184347 | 424.3869 | -1.57128 | 0.239743 | -6.55401 | 5.60E-11 | 2.88E-09 | 11.14335 | 4.727126 | 16.58786 | 4.105927 | SLIT3 | | 5 |
| 815 | ENSG00000184349 | 468.5746 | 1.212992 | 0.243154 | 4.988568 | 6.08E-07 | 1.66E-05 | 8.581778 | 19.16419 | 12.73221 | 27.76123 | EFNA5 | | 5 |
| 816 | ENSG00000184515 | 17.7172 | 2.88343 | 0.944172 | 3.053926 | 0.002259 | 0.024365 | 1.354588 | 5.861195 | 0.4021 | 6.169894 | BEX5 | X | |
| 817 | ENSG00000184672 | 158.9242 | 1.055667 | 0.311724 | 3.386542 | 0.000708 | 0.009221 | 8.941612 | 14.32143 | 5.904677 | 14.69731 | RALYL | | 8 |
| 818 | ENSG00000184697 | 154.3893 | 1.529931 | 0.342991 | 4.460555 | 8.17E-06 | 0.000178 | 6.106033 | 24.55046 | 9.099796 | 16.9619 | CLDN6 | | 16 |
| 819 | ENSG00000184985 | 971.7101 | -1.70657 | 0.16956 | -10.0647 | 7.91E-24 | 1.42E-21 | 45.29697 | 11.01575 | 35.77802 | 12.46122 | SORCS2 | | 4 |
| 820 | ENSG00000185046 | 402.142 | -1.26321 | 0.236776 | -5.33504 | 9.55E-08 | 3.06E-06 | 45.47937 | 16.78911 | 31.7604 | 13.38818 | ANKS1B | | 12 |
| 821 | ENSG00000185149 | 29.76621 | -2.22665 | 0.705113 | -3.15786 | 0.001589 | 0.018313 | 1.467415 | 0.457528 | 1.757838 | 0.186992 | NPY2R | | 4 |
| 822 | ENSG00000185201 | 47.12791 | 1.861613 | 0.649731 | 2.865205 | 0.004167 | 0.03979 | 1.792979 | 18.29298 | 7.575008 | 11.7355 | IFITM2 | | 11 |
| 823 | ENSG00000185559 | 9585.926 | 1.526351 | 0.453045 | 3.369096 | 0.000754 | 0.009715 | 228.076 | 734.4548 | 401.4202 | 988.693 | DLK1 | | 14 |
| 824 | ENSG00000185565 | 2307.393 | -1.34294 | 0.148954 | -9.01583 | 1.95E-19 | 2.39E-17 | 68.1494 | 27.82202 | 66.60471 | 22.22714 | LSAMP | | 3 |
| 825 | ENSG00000185585 | 148.2429 | 1.279388 | 0.33755 | 3.790223 | 0.000151 | 0.002403 | 1.784108 | 4.056122 | 1.254017 | 2.849566 | OLFML2A | | 9 |
| 826 | ENSG00000185664 | 124.4034 | 1.745198 | 0.376252 | 4.638371 | 3.51E-06 | 8.21E-05 | 4.373215 | 14.1749 | 6.779078 | 21.47246 | PMEL | | 12 |
| 827 | ENSG00000185668 | 171.9434 | 2.076479 | 0.300432 | 6.911635 | 4.79E-12 | 2.72E-10 | 2.339652 | 10.46433 | 2.879918 | 10.39021 | POU3F1 | | 1 |
| 828 | ENSG00000185745 | 526.7333 | -3.27727 | 0.208188 | -15.7419 | 7.81E-56 | 6.94E-53 | 24.4595 | 2.652048 | 26.33144 | 2.287513 | IFIT1 | | 10 |
| 829 | ENSG00000185885 | 139.5159 | 1.987037 | 0.344789 | 5.76305 | 8.26E-09 | 3.14E-07 | 11.41606 | 57.6799 | 18.52481 | 55.26062 | IFITM1 | | 11 |
| 830 | ENSG00000185920 | 1110.6 | -1.9147 | 0.164769 | -11.6205 | 3.24E-31 | 9.90E-29 | 57.45157 | 12.83996 | 44.68518 | 12.71566 | PTCH1 | | 9 |
| 831 | ENSG00000185985 | 1064.701 | -2.31253 | 0.189801 | -12.1839 | 3.79E-34 | 1.51E-31 | 26.97713 | 5.456796 | 22.35289 | 3.874695 | SLITRK2 | X | |
| 832 | ENSG00000186007 | 38.11508 | -1.61023 | 0.580928 | -2.77183 | 0.005574 | 0.049862 | 16.5503 | 4.924872 | 13.66395 | 4.386416 | LEMD1 | | 1 |
| 833 | ENSG00000186369 | 75.90494 | -3.40917 | 0.506246 | -6.73422 | 1.65E-11 | 8.95E-10 | 7.619653 | 0.758989 | 4.927593 | 0.373803 | LINC00643 | | 14 |
| 834 | ENSG00000186446 | 140.5446 | -1.30522 | 0.321238 | -4.06308 | 4.84E-05 | 0.000879 | 7.119074 | 2.835272 | 9.016915 | 3.35008 | ZNF501 | | 3 |
| 835 | ENSG00000186479 | 84.33801 | -1.30257 | 0.413929 | -3.14684 | 0.00165 | 0.018862 | 3.397119 | 1.618442 | 3.488922 | 1.018342 | RGS7BP | | 5 |
| 836 | ENSG00000186493 | 45.2065 | 3.060462 | 0.600488 | 5.096621 | 3.46E-07 | 9.96E-06 | 1.320762 | 13.95123 | 2.181537 | 14.15648 | C5orf38 | | 5 |
| 837 | ENSG00000186815 | 100.3825 | 1.4849 | 0.394599 | 3.763057 | 0.000168 | 0.00264 | 1.564025 | 5.847158 | 2.604994 | 5.076438 | TPCN1 | | 12 |
| 838 | ENSG00000186897 | 126.1716 | 1.632364 | 0.332211 | 4.913642 | 8.94E-07 | 2.34E-05 | 3.744196 | 10.04983 | 3.53674 | 11.28145 | C1QL4 | | 12 |
| 839 | ENSG00000186960 | 145.891 | -4.3454 | 0.435366 | -9.98103 | 1.85E-23 | 3.17E-21 | 27.57021 | 0.955477 | 15.90479 | 1.075726 | LINC01551 | | 14 |
| 840 | ENSG00000187244 | 149.2256 | 1.326498 | 0.345242 | 3.842231 | 0.000122 | 0.002012 | 4.143647 | 13.4768 | 5.083271 | 8.418819 | BCAM | | 19 |
| 841 | ENSG00000187260 | 124.1953 | -1.25808 | 0.328942 | -3.82462 | 0.000131 | 0.002141 | 17.32555 | 6.266793 | 16.87845 | 7.25979 | WDR86 | | 7 |
| 842 | ENSG00000187391 | 1138.168 | -1.24224 | 0.147073 | -8.44642 | 3.00E-17 | 2.92E-15 | 49.7405 | 18.40968 | 45.53084 | 19.63075 | MAGI2 | | 7 |
| 843 | ENSG00000187398 | 97.78874 | 1.11926 | 0.4011 | 2.79048 | 0.005263 | 0.047558 | 2.140358 | 3.678662 | 1.181827 | 3.051052 | LUZP2 | | 11 |
| 844 | ENSG00000187498 | 834.166 | 1.246887 | 0.18797 | 6.633439 | 3.28E-11 | 1.73E-09 | 10.77438 | 19.98187 | 10.8679 | 28.74816 | COL4A1 | | 13 |
| 845 | ENSG00000187553 | 38.9429 | 2.528707 | 0.660056 | 3.831047 | 0.000128 | 0.002091 | 0.32621 | 1.186345 | 0.415268 | 2.913751 | CYP26C1 | | 10 |
| 846 | ENSG00000187678 | 377.5933 | 1.828752 | 0.244347 | 7.484252 | 7.20E-14 | 4.97E-12 | 3.843867 | 15.69772 | 6.012191 | 17.63795 | SPRY4 | | 5 |
| 847 | ENSG00000187720 | 407.8997 | -2.32901 | 0.626878 | -3.71525 | 0.000203 | 0.003131 | 12.57085 | 3.174159 | 11.29423 | 1.289331 | THSD4 | | 15 |
| 848 | ENSG00000187840 | 213.7517 | 1.030861 | 0.264825 | 3.892613 | 9.92E-05 | 0.001668 | 30.11254 | 54.16477 | 24.44384 | 50.90642 | EIF4EBP1 | | 8 |
| 849 | ENSG00000188064 | 1506.154 | -3.42797 | 0.171091 | -20.036 | 2.68E-89 | 8.72E-86 | 78.95619 | 8.349731 | 91.25359 | 6.54656 | WNT7B | | 22 |
| 850 | ENSG00000188191 | 106.8404 | -1.96634 | 0.391141 | -5.02719 | 4.98E-07 | 1.39E-05 | 14.61162 | 2.3697 | 9.732594 | 3.507912 | PRKAR1B | | 7 |
| 851 | ENSG00000188321 | 319.6704 | -1.04972 | 0.220231 | -4.76644 | 1.88E-06 | 4.62E-05 | 21.89818 | 9.39575 | 19.66461 | 9.554006 | ZNF559 | | 19 |
| 852 | ENSG00000188483 | 196.6341 | 1.325434 | 0.273749 | 4.841789 | 1.29E-06 | 3.26E-05 | 4.530161 | 11.28357 | 5.283458 | 12.00475 | IER5L | | 9 |
| 853 | ENSG00000188596 | 190.5858 | -1.50477 | 0.336085 | -4.47734 | 7.56E-06 | 0.000166 | 9.27219 | 1.907573 | 8.676723 | 4.18285 | CFAP54 | | 12 |
| 854 | ENSG00000188707 | 31.28558 | -2.1536 | 0.683494 | -3.15087 | 0.001628 | 0.018636 | 1.861263 | 0.244772 | 0.8993 | 0.333534 | ZBED10P | | 7 |
| 855 | ENSG00000188738 | 806.6226 | -1.23256 | 0.16605 | -7.4228 | 1.15E-13 | 7.71E-12 | 10.13672 | 3.56137 | 8.790994 | 4.050127 | FSIP2 | | 2 |
| 856 | ENSG00000188803 | 143.9578 | -2.17892 | 0.351718 | -6.19508 | 5.83E-10 | 2.68E-08 | 5.034156 | 1.324088 | 4.772038 | 0.740932 | SHISA6 | | 17 |
| 857 | ENSG00000188859 | 52.34509 | -1.50743 | 0.503466 | -2.9941 | 0.002753 | 0.028224 | 2.366433 | 0.706516 | 2.691522 | 0.987573 | FAM78B | | 1 |

| | | | | | | | | | | | | | | |
|---|---|---|---|---|---|---|---|---|---|---|---|---|---|---|
| 858 | ENSG00000189056 | 1429.992 | -2.1273 | 0.183273 | -11.6073 | 3.78E-31 | 1.14E-28 | 31.01373 | 8.266931 | 33.7914 | 5.700193 | RELN | | 7 |
| 859 | ENSG00000189108 | 60.55303 | -8.90418 | 1.512725 | -5.88619 | 3.95E-09 | 1.59E-07 | 7.861876 | 0 | 5.299602 | 0 | IL1RAPL2 | X | |
| 860 | ENSG00000189127 | 87.4764 | -2.66211 | 0.442166 | -6.02061 | 1.74E-09 | 7.45E-08 | 5.779987 | 0.939853 | 4.157779 | 0.526816 | ANKRD34B | | 5 |
| 861 | ENSG00000189157 | 32.03402 | -1.79682 | 0.632537 | -2.84065 | 0.004502 | 0.042162 | 2.099985 | 0.518663 | 2.192301 | 0.656101 | FAM47E | | 4 |
| 862 | ENSG00000189171 | 122.9309 | 1.226741 | 0.377947 | 3.245805 | 0.001171 | 0.014207 | 29.78906 | 78.43062 | 26.15901 | 44.55501 | S100A13 | | 1 |
| 863 | ENSG00000189369 | 704.0953 | -1.14665 | 0.166235 | -6.89775 | 5.28E-12 | 2.99E-10 | 42.16243 | 17.32349 | 40.1708 | 17.79414 | GSPT2 | X | |
| 864 | ENSG00000196074 | 44.48221 | -2.82087 | 0.670393 | -4.20778 | 2.58E-05 | 0.000496 | 4.19018 | 0.357387 | 1.048538 | 0.311694 | SYCP2 | | 20 |
| 865 | ENSG00000196083 | 332.9441 | 2.787363 | 0.243624 | 11.44123 | 2.60E-30 | 7.27E-28 | 2.790288 | 20.11771 | 2.796983 | 16.24041 | IL1RAP | | 3 |
| 866 | ENSG00000196090 | 106.8334 | -1.79274 | 0.472627 | -3.79313 | 0.000149 | 0.002386 | 2.073076 | 0.287886 | 2.340076 | 1.022917 | PTPRT | | 20 |
| 867 | ENSG00000196154 | 31.7276 | 2.506842 | 0.734563 | 3.412697 | 0.000643 | 0.008522 | 3.022549 | 23.76244 | 3.013167 | 8.332517 | S100A4 | | 1 |
| 868 | ENSG00000196159 | 5535.715 | -1.35206 | 0.42154 | -3.20742 | 0.001339 | 0.015854 | 74.01982 | 23.83346 | 42.33068 | 18.97774 | FAT4 | | 4 |
| 869 | ENSG00000196187 | 127.7985 | 1.027554 | 0.33357 | 3.080475 | 0.002067 | 0.022716 | 4.09014 | 9.129252 | 4.391544 | 7.173387 | TMEM63A | | 1 |
| 870 | ENSG00000196263 | 57.64312 | -7.87317 | 1.414389 | -5.56648 | 2.60E-08 | 9.14E-07 | 2.21088 | 0.016659 | 3.452487 | 0 | ZNF471 | | 19 |
| 871 | ENSG00000196268 | 507.4604 | -1.37154 | 0.189538 | -7.23626 | 4.61E-13 | 2.91E-11 | 32.08934 | 12.11082 | 35.0617 | 12.42357 | ZNF493 | | 19 |
| 872 | ENSG00000196277 | 132.9259 | -2.37653 | 0.35318 | -6.72894 | 1.71E-11 | 9.26E-10 | 10.00532 | 1.736909 | 7.063869 | 1.366928 | GRM7 | | 3 |
| 873 | ENSG00000196376 | 1379.218 | 1.074558 | 0.149561 | 7.18476 | 6.73E-13 | 4.15E-11 | 23.36581 | 41.28243 | 19.00644 | 42.87373 | SLC35F1 | | 6 |
| 874 | ENSG00000196458 | 498.3741 | -1.3373 | 0.192611 | -6.94298 | 3.84E-12 | 2.20E-10 | 18.49076 | 7.475122 | 20.41883 | 7.074936 | ZNF605 | | 12 |
| 875 | ENSG00000196459 | 403.3704 | -1.00653 | 0.20201 | -4.98256 | 6.27E-07 | 1.69E-05 | 26.99236 | 11.96624 | 27.33789 | 13.60707 | TRAPPC2 | X | |
| 876 | ENSG00000196562 | 969.9474 | 1.309783 | 0.16004 | 8.184108 | 2.74E-16 | 2.38E-14 | 22.19379 | 51.29568 | 19.84099 | 46.88877 | SULF2 | | 20 |
| 877 | ENSG00000196604 | 113.1786 | -2.24032 | 0.373844 | -5.99266 | 2.06E-09 | 8.74E-08 | 5.330891 | 0.777367 | 5.100452 | 1.320882 | POTEF | | 2 |
| 878 | ENSG00000196639 | 22.10502 | 4.67544 | 1.013021 | 4.615344 | 3.92E-06 | 9.11E-05 | 0.08239 | 1.272512 | 0.024189 | 1.231883 | HRH1 | | 3 |
| 879 | ENSG00000196659 | 175.3083 | -1.00949 | 0.282759 | -3.57014 | 0.000357 | 0.005156 | 7.669115 | 3.317671 | 6.578644 | 3.357757 | TTC30B | | 2 |
| 880 | ENSG00000196781 | 1023.154 | -1.53245 | 0.159157 | -9.62853 | 6.06E-22 | 8.71E-20 | 71.96774 | 23.02411 | 62.99309 | 20.94886 | TLE1 | | 9 |
| 881 | ENSG00000196782 | 708.1864 | -1.57093 | 0.192867 | -8.14515 | 3.79E-16 | 3.23E-14 | 20.18678 | 6.455563 | 16.31931 | 5.113649 | MAML3 | | 4 |
| 882 | ENSG00000196867 | 207.2423 | -2.02705 | 0.307337 | -6.59553 | 4.24E-11 | 2.23E-09 | 12.18464 | 3.468479 | 11.33941 | 2.000538 | ZFP28 | | 19 |
| 883 | ENSG00000197013 | 227.2758 | -1.43373 | 0.275926 | -5.19607 | 2.04E-07 | 6.10E-06 | 13.40927 | 5.162723 | 11.71293 | 3.665906 | ZNF429 | | 19 |
| 884 | ENSG00000197177 | 64.82407 | -2.02674 | 0.466341 | -4.34606 | 1.39E-05 | 0.000285 | 3.017431 | 0.768229 | 2.684044 | 0.545117 | ADGRA1 | | 10 |
| 885 | ENSG00000197321 | 1340.069 | -1.38296 | 0.165557 | -8.35339 | 6.63E-17 | 6.15E-15 | 56.32877 | 17.21481 | 41.38091 | 18.09126 | SVIL | | 10 |
| 886 | ENSG00000197406 | 918.3665 | 2.424592 | 0.568348 | 4.266032 | 1.99E-05 | 0.000395 | 10.34588 | 77.97814 | 23.05651 | 92.52284 | DIO3 | | 14 |
| 887 | ENSG00000197415 | 375.4332 | -1.41095 | 0.239741 | -5.8853 | 3.97E-09 | 1.60E-07 | 45.58562 | 13.50332 | 30.14397 | 13.28279 | VEPH1 | | 3 |
| 888 | ENSG00000197550 | 26.37264 | -2.00731 | 0.70655 | -2.841 | 0.004497 | 0.042136 | 1.545479 | 0.401614 | 1.46694 | 0.303009 | | | 9 |
| 889 | ENSG00000197584 | 22.6539 | -2.48772 | 0.858122 | -2.89903 | 0.003743 | 0.036451 | 3.586239 | 0.639215 | 1.502114 | 0.251906 | KCNMB2 | | 3 |
| 890 | ENSG00000197705 | 199.8988 | -1.01472 | 0.27781 | -3.65256 | 0.00026 | 0.003893 | 7.729623 | 4.046235 | 7.632174 | 3.103948 | KLHL14 | | 18 |
| 891 | ENSG00000197712 | 157.2843 | 1.422857 | 0.311259 | 4.571294 | 4.85E-06 | 0.00011 | 6.570426 | 13.39159 | 4.597113 | 14.576 | FAM114A1 | | 4 |
| 892 | ENSG00000197747 | 85.75067 | 2.088161 | 0.431061 | 4.84423 | 1.27E-06 | 3.22E-05 | 7.330596 | 40.03062 | 9.318074 | 26.76785 | S100A10 | | 1 |
| 893 | ENSG00000197921 | 349.9222 | -1.88638 | 0.248737 | -7.58385 | 3.35E-14 | 2.42E-12 | 64.66177 | 11.84123 | 46.08957 | 16.48553 | HES5 | | 1 |
| 894 | ENSG00000197928 | 265.9038 | -2.28003 | 0.281923 | -8.08741 | 6.09E-16 | 5.16E-14 | 30.50858 | 4.169915 | 29.45242 | 7.614808 | ZNF677 | | 19 |
| 895 | ENSG00000198046 | 37.67906 | -6.29326 | 1.167092 | -5.39225 | 6.96E-08 | 2.28E-06 | 2.093349 | 0.030996 | 3.284819 | 0.033779 | ZNF667 | | 19 |
| 896 | ENSG00000198049 | 16.4259 | -4.38825 | 1.21716 | -3.60532 | 0.000312 | 0.004584 | 1.079892 | 0 | 0.597163 | 0.063984 | AVPR1B | | 1 |
| 897 | ENSG00000198189 | 84.56406 | 1.246728 | 0.408005 | 3.05567 | 0.002246 | 0.024246 | 3.521791 | 9.074109 | 5.094586 | 10.28341 | HSD17B11 | | 4 |
| 898 | ENSG00000198205 | 27.3513 | -8.27947 | 1.586967 | -5.21717 | 1.82E-07 | 5.50E-06 | 1.003921 | 0 | 1.458814 | 0 | ZXDA | X | |
| 899 | ENSG00000198300 | 3019.13 | -9.82226 | 0.99949 | -9.82727 | 8.59E-23 | 1.33E-20 | 170.134 | 0.055465 | 124.305 | 0.310381 | PEG3 | | 19 |
| 900 | ENSG00000198400 | 26.63588 | -2.95546 | 0.74147 | -3.98595 | 6.72E-05 | 0.00118 | 2.461106 | 0.254983 | 2.859758 | 0.396967 | NTRK1 | | 1 |
| 901 | ENSG00000198429 | 526.4257 | -1.30538 | 0.184234 | -7.08545 | 1.39E-12 | 8.36E-11 | 46.30243 | 16.76567 | 42.56175 | 17.17895 | ZNF69 | | 19 |
| 902 | ENSG00000198453 | 192.9491 | -1.69827 | 0.285618 | -5.94594 | 2.75E-09 | 1.13E-07 | 13.28024 | 3.064474 | 11.99945 | 4.3287 | ZNF568 | | 19 |
| 903 | ENSG00000198467 | 280.5269 | 2.468793 | 0.262188 | 9.416107 | 4.68E-21 | 6.27E-19 | 8.676679 | 53.69113 | 12.64728 | 58.37788 | TPM2 | | 9 |
| 904 | ENSG00000198626 | 107.6625 | 1.267631 | 0.350863 | 3.612897 | 0.000303 | 0.004469 | 0.559438 | 1.317827 | 0.55573 | 1.21866 | RYR2 | | 1 |
| 905 | ENSG00000198673 | 118.4437 | -1.1578 | 0.336552 | -3.4402 | 0.000581 | 0.007844 | 9.494643 | 3.802495 | 8.939057 | 3.999961 | TAFA2 | | 12 |
| 906 | ENSG00000198682 | 469.4044 | -1.15261 | 0.25013 | -4.60805 | 4.06E-06 | 9.41E-05 | 14.24328 | 8.076799 | 24.45719 | 8.435273 | PAPSS2 | | 10 |
| 907 | ENSG00000198718 | 517.5677 | -1.22875 | 0.252536 | -4.86565 | 1.14E-06 | 2.93E-05 | 23.06595 | 12.11828 | 23.58045 | 6.671183 | TOGARAM | | 14 |
| 908 | ENSG00000198729 | 477.5907 | 3.306957 | 0.216846 | 15.25023 | 1.64E-52 | 1.34E-49 | 5.04879 | 46.80606 | 4.562482 | 42.81646 | PPP1R14C | | 6 |
| 909 | ENSG00000198739 | 123.3105 | -1.42639 | 0.354948 | -4.01858 | 5.86E-05 | 0.001046 | 3.951239 | 1.321235 | 2.75035 | 1.018858 | LRRTM3 | | 10 |
| 910 | ENSG00000198796 | 218.8236 | 2.546892 | 0.350578 | 7.264834 | 3.73E-13 | 2.40E-11 | 2.235297 | 9.939024 | 0.967512 | 7.025403 | ALPK2 | | 18 |
| 911 | ENSG00000198797 | 199.9197 | -3.038 | 0.312046 | -9.73576 | 2.12E-22 | 3.17E-20 | 11.41112 | 1.686069 | 12.40955 | 1.053325 | BRINP2 | | 1 |
| 912 | ENSG00000198821 | 27.74408 | 3.05066 | 0.839665 | 3.633187 | 0.00028 | 0.004172 | 0.181681 | 3.883693 | 1.07631 | 4.239241 | CD247 | | 1 |
| 913 | ENSG00000198825 | 1036.27 | -1.52651 | 0.154479 | -9.88165 | 5.00E-23 | 8.16E-21 | 45.80922 | 13.41231 | 40.30794 | 14.81354 | INPP5F | | 10 |
| 914 | ENSG00000198846 | 729.6025 | -2.16175 | 0.196308 | -11.012 | 3.35E-28 | 7.98E-26 | 37.92349 | 7.864487 | 29.40619 | 6.282744 | TOX | | 8 |
| 915 | ENSG00000198879 | 406.0398 | 1.246588 | 0.208616 | 5.975514 | 2.29E-09 | 9.56E-08 | 5.596061 | 12.50785 | 6.407523 | 14.48389 | SFMBT2 | | 10 |
| 916 | ENSG00000198959 | 23.0502 | 2.545303 | 0.814522 | 3.124903 | 0.001779 | 0.020069 | 0.297377 | 1.104631 | 0.111898 | 1.008188 | TGM2 | | 20 |
| 917 | ENSG00000198960 | 353.6756 | -1.16118 | 0.231405 | -5.01797 | 5.22E-07 | 1.45E-05 | 45.90737 | 21.79 | 44.078 | 16.13669 | ARMCX6 | X | |
| 918 | ENSG00000198963 | 302.5853 | -2.4126 | 0.238529 | -10.1145 | 4.77E-24 | 8.71E-22 | 6.11653 | 1.171514 | 6.527691 | 1.067842 | RORB | | 9 |
| 919 | ENSG00000203335 | 40.2765 | -1.7699 | 0.625062 | -2.83156 | 0.004632 | 0.043071 | 7.022263 | 2.08699 | 3.899403 | 0.930851 | | | 7 |
| 920 | ENSG00000203727 | 997.73 | 1.078838 | 0.177044 | 6.093626 | 1.10E-09 | 4.86E-08 | 12.30098 | 26.38854 | 11.2363 | 20.40671 | SAMD5 | | 6 |
| 921 | ENSG00000204128 | 90.72505 | 1.520999 | 0.391248 | 3.887555 | 0.000101 | 0.001695 | 2.031608 | 4.557941 | 1.461883 | 4.878483 | C2orf72 | | 2 |
| 922 | ENSG00000204335 | 61.74568 | 5.260175 | 0.735648 | 7.150393 | 8.65E-13 | 5.32E-11 | 0.266542 | 7.594473 | 0.117638 | 6.005555 | SP5 | | 2 |
| 923 | ENSG00000204389 | 283.0113 | -5.23714 | 0.354575 | -14.7702 | 2.28E-49 | 1.65E-46 | 28.71172 | 0.677341 | 31.99949 | 0.859108 | HSPA1A | | 6 |
| 924 | ENSG00000204525 | 400.0546 | 1.052428 | 0.206543 | 5.095434 | 3.48E-07 | 1.00E-05 | 22.19068 | 46.00626 | 22.50063 | 41.45404 | HLA-C | | 6 |
| 925 | ENSG00000204682 | 635.8795 | 1.05485 | 0.175206 | 6.020611 | 1.74E-09 | 7.45E-08 | 12.29761 | 25.30382 | 12.80008 | 23.93478 | MIR1915H | | 10 |
| 926 | ENSG00000204706 | 66.22559 | 1.775232 | 0.476747 | 3.723639 | 0.000196 | 0.003045 | 1.957166 | 9.456392 | 2.941552 | 6.343728 | MAMDC2- | | 9 |
| 927 | ENSG00000204802 | 73.84723 | 1.199556 | 0.427509 | 2.805918 | 0.005017 | 0.045739 | 2.98094 | 4.915206 | 2.534743 | 7.129884 | FAM88C | | 9 |
| 928 | ENSG00000204969 | 96.55932 | -1.63022 | 0.386954 | -4.21295 | 2.52E-05 | 0.000486 | 4.927329 | 1.809435 | 5.804739 | 1.457182 | PCDHA2 | | 5 |
| 929 | ENSG00000205363 | 255.7658 | -1.52246 | 0.288687 | -5.27376 | 1.34E-07 | 4.13E-06 | 21.18085 | 6.348314 | 13.06531 | 4.83389 | INSYN1 | | 15 |
| 930 | ENSG00000205856 | 22.10605 | -2.69754 | 0.808488 | -3.33653 | 0.000848 | 0.010764 | 4.112535 | 0.541576 | 3.313135 | 0.535573 | C22orf42 | | 22 |
| 931 | ENSG00000206144 | 15.68561 | -3.06022 | 0.984271 | -3.10912 | 0.001876 | 0.020951 | 1.44519 | 0.147122 | 1.719016 | 0.213674 | MAPK6P1 | | 8 |
| 932 | ENSG00000206190 | 28.77857 | 2.013473 | 0.688032 | 2.926422 | 0.003429 | 0.033863 | 0.389559 | 1.341525 | 0.519547 | 2.176858 | ATP10A | | 15 |
| 933 | ENSG00000206503 | 239.7324 | -1.08246 | 0.251553 | -4.3031 | 1.68E-05 | 0.00034 | 25.43433 | 12.12704 | 29.50095 | 12.41105 | HLA-A | | 6 |
| 934 | ENSG00000206538 | 670.2994 | 3.795152 | 0.205501 | 18.46784 | 3.75E-76 | 5.64E-73 | 1.001239 | 15.54011 | 1.303906 | 14.56102 | VGLL3 | | 3 |
| 935 | ENSG00000206557 | 487.7951 | 1.214685 | 0.229081 | 5.302429 | 1.14E-07 | 3.58E-06 | 4.144637 | 6.465297 | 3.36056 | 10.02615 | TRIM71 | | 3 |

| | | | | | | | | | | | | | | |
|---|---|---|---|---|---|---|---|---|---|---|---|---|---|---|
| 936 | ENSG00000207955 | 59.19425 | 2.743218 | 0.570687 | 4.806871 | 1.53E-06 | 3.84E-05 | 0.667657 | 6.113897 | 1.939484 | 10.7132 | MIR219A2 | | 9 |
| 937 | ENSG00000211448 | 106.5559 | -2.94334 | 0.458492 | -6.4196 | 1.37E-10 | 6.73E-09 | 3.340835 | 0.793236 | 6.671077 | 0.419193 | DIO2 | | 14 |
| 938 | ENSG00000213047 | 484.4401 | -1.5841 | 0.212203 | -7.46503 | 8.33E-14 | 5.69E-12 | 24.29511 | 7.678967 | 19.60011 | 6.086126 | DENND1B | | 1 |
| 939 | ENSG00000213468 | 271.0367 | 2.919924 | 0.268809 | 10.86246 | 1.74E-27 | 3.87E-25 | 5.144832 | 34.29928 | 5.789686 | 44.16692 | FIRRE | X | |
| 940 | ENSG00000213853 | 327.1838 | 1.026829 | 0.232799 | 4.410791 | 1.03E-05 | 0.000218 | 6.544561 | 13.21963 | 5.818149 | 10.49534 | EMP2 | | 16 |
| 941 | ENSG00000214132 | 15.91601 | -3.10523 | 1.063393 | -2.92012 | 0.003499 | 0.034468 | 1.450547 | 0.486159 | 3.849052 | 0.088185 | | | 5 |
| 942 | ENSG00000214548 | 5770.764 | -7.32712 | 0.207254 | -35.3533 | 8.91E-274 | 1.74E-269 | 228.6027 | 1.787613 | 311.0682 | 1.407752 | MEG3 | | 14 |
| 943 | ENSG00000214919 | 37.20081 | -1.77484 | 0.601933 | -2.94856 | 0.003193 | 0.031839 | 4.211825 | 0.980453 | 5.068933 | 1.601074 | | | 3 |
| 944 | ENSG00000215190 | 130.7417 | -3.18821 | 0.345758 | -9.22093 | 2.95E-20 | 3.81E-18 | 37.50225 | 3.143124 | 36.59754 | 4.545811 | LINC0068C | | 6 |
| 945 | ENSG00000215196 | 289.6294 | -1.12592 | 0.246589 | -4.56598 | 4.97E-06 | 0.000113 | 26.23535 | 9.000775 | 19.21757 | 10.64302 | BASP1-AS1 | | 5 |
| 946 | ENSG00000215218 | 193.8826 | -1.40076 | 0.279342 | -5.01452 | 5.32E-07 | 1.47E-05 | 5.567761 | 2.11869 | 5.18303 | 1.714334 | UBE2QL1 | | 5 |
| 947 | ENSG00000215374 | 22.01608 | -4.76141 | 1.093108 | -4.35584 | 1.33E-05 | 0.000275 | 1.762818 | 0.04823 | 2.116475 | 0.085573 | FAM66B | | 8 |
| 948 | ENSG00000215808 | 26.66517 | 5.765701 | 1.062584 | 5.426115 | 5.76E-08 | 1.91E-06 | 0.044986 | 1.328692 | 0.019806 | 1.941628 | LINC0113S | | 1 |
| 949 | ENSG00000216895 | 49.3755 | 1.833361 | 0.523976 | 3.498941 | 0.000467 | 0.006455 | 4.636782 | 12.02012 | 4.655039 | 19.74347 | RBM33-DT | | 7 |
| 950 | ENSG00000220323 | 181.6341 | -1.05264 | 0.287977 | -3.65531 | 0.000257 | 0.003861 | 8.344704 | 3.029735 | 7.384696 | 4.150327 | H2BC19P | | 1 |
| 951 | ENSG00000221866 | 225.6765 | -1.68043 | 0.266856 | -6.29714 | 3.03E-10 | 1.44E-08 | 4.477316 | 1.104972 | 4.073728 | 1.435988 | PLXNA4 | | 7 |
| 952 | ENSG00000221890 | 293.1408 | -1.30176 | 0.233489 | -5.57527 | 2.47E-08 | 8.74E-07 | 8.792818 | 2.930396 | 7.344581 | 3.249528 | NPTXR | | 22 |
| 953 | ENSG00000222041 | 214.098 | 2.965181 | 0.302922 | 9.788582 | 1.26E-22 | 1.94E-20 | 5.013227 | 42.10127 | 4.926879 | 31.05465 | CYTOR | | 2 |
| 954 | ENSG00000223403 | 98.91242 | -5.42283 | 0.575145 | -9.42862 | 4.16E-21 | 5.60E-19 | 11.07302 | 0.240625 | 11.69321 | 0.259908 | MEG9 | | 14 |
| 955 | ENSG00000223691 | 26.72192 | -2.5848 | 0.731625 | -3.53296 | 0.000411 | 0.005806 | 2.840847 | 0.540953 | 2.875 | 0.353499 | | | 2 |
| 956 | ENSG00000224165 | 38.12329 | -3.83759 | 0.669984 | -5.72789 | 1.02E-08 | 3.81E-07 | 11.11154 | 0.458521 | 4.456518 | 0.48736 | DNAJC27-/ | | 2 |
| 957 | ENSG00000224597 | 45.08835 | -2.97656 | 0.601936 | -4.94497 | 7.62E-07 | 2.01E-05 | 11.91986 | 1.301844 | 10.77531 | 1.418827 | SVIL-AS1 | | 10 |
| 958 | ENSG00000224668 | 17.37272 | 4.432545 | 1.185952 | 3.737542 | 0.000186 | 0.002891 | 0 | 0.849882 | 0.110869 | 1.531724 | IPO8P1 | | 1 |
| 959 | ENSG00000224717 | 80.69763 | -1.70874 | 0.408617 | -4.18177 | 2.89E-05 | 0.000551 | 4.730148 | 1.454173 | 5.373425 | 1.468623 | LEMD1-DT | | 1 |
| 960 | ENSG00000225156 | 64.1087 | -2.38673 | 0.47071 | -5.07049 | 3.97E-07 | 1.12E-05 | 5.068686 | 0.882787 | 4.469817 | 0.836409 | | | 2 |
| 961 | ENSG00000225345 | 33.25354 | 2.865982 | 0.686309 | 4.175937 | 2.97E-05 | 0.000564 | 0.841864 | 3.598272 | 0.563835 | 6.0319 | SNX18P3 | | 9 |
| 962 | ENSG00000225465 | 74.90596 | -1.76048 | 0.428614 | -4.10737 | 4.00E-05 | 0.000738 | 3.52365 | 0.89838 | 2.557354 | 0.791569 | RFPL1S | | 22 |
| 963 | ENSG00000225649 | 29.18106 | 2.032828 | 0.711778 | 2.855985 | 0.00429 | 0.040705 | 0.806136 | 2.711579 | 0.387646 | 1.78455 | | | 2 |
| 964 | ENSG00000225683 | 46.715 | -2.05912 | 0.556144 | -3.70249 | 0.000213 | 0.003279 | 3.743407 | 0.729657 | 2.143011 | 0.593911 | PACRG-AS | | 6 |
| 965 | ENSG00000225746 | 1450.062 | -5.11996 | 0.200431 | -25.5447 | 6.28E-144 | 6.14E-140 | 186.097 | 5.721292 | 239.4246 | 5.849246 | MEG8 | | 14 |
| 966 | ENSG00000225792 | 213.9352 | -1.08235 | 0.268539 | -4.0305 | 5.57E-05 | 0.000996 | 48.73132 | 23.17323 | 44.97456 | 18.51091 | SNX10-AS | | 7 |
| 967 | ENSG00000225868 | 7.988685 | -6.01283 | 1.872845 | -3.21053 | 0.001325 | 0.015739 | 1.005029 | 0 | 1.411488 | 0 | WDR87BP | | 19 |
| 968 | ENSG00000225930 | 249.0904 | 1.24543 | 0.248653 | 5.0087 | 5.48E-07 | 1.51E-05 | 7.330533 | 16.42952 | 8.358544 | 18.74328 | LINC0224S | | 15 |
| 969 | ENSG00000226440 | 8.054449 | 4.357736 | 1.531324 | 2.845731 | 0.004431 | 0.041655 | 0.107219 | 1.823696 | 0.094855 | 2.0876 | LAMA4-AS | | 6 |
| 970 | ENSG00000226686 | 24.11355 | -4.31338 | 0.942492 | -4.57657 | 4.73E-06 | 0.000108 | 2.374814 | 0.104033 | 1.450216 | 0.075576 | LINC01535 | | 19 |
| 971 | ENSG00000226702 | 36.49472 | -2.01409 | 0.63612 | -3.1662 | 0.001544 | 0.017901 | 10.26814 | 1.295565 | 9.122509 | 3.284578 | MIR217HG | | 2 |
| 972 | ENSG00000226887 | 59.58453 | 1.841198 | 0.48562 | 3.791435 | 0.00015 | 0.002397 | 0.864875 | 3.643027 | 1.3133 | 3.782752 | ERVMER34 | | 4 |
| 973 | ENSG00000227082 | 21.08699 | 2.826637 | 0.840387 | 3.363493 | 0.00077 | 0.009888 | 0.175696 | 1.138812 | 0.150935 | 1.041602 | LINC02798 | | 1 |
| 974 | ENSG00000227124 | 29.04908 | -3.76835 | 0.800266 | -4.70888 | 2.49E-06 | 5.98E-05 | 2.245818 | 0.169261 | 2.997986 | 0.194783 | ZNF717 | | 3 |
| 975 | ENSG00000227157 | 8.599485 | 6.30418 | 1.915765 | 3.290686 | 0.000999 | 0.012345 | 0 | 2.526695 | 0 | 0.492401 | | | 2 |
| 976 | ENSG00000228010 | 18.1798 | -2.52336 | 0.892421 | -2.82755 | 0.004691 | 0.043504 | 4.601505 | 1.272258 | 8.018441 | 0.765961 | | | 7 |
| 977 | ENSG00000228061 | 81.79445 | -1.17525 | 0.412042 | -2.85225 | 0.004341 | 0.041012 | 5.693658 | 3.068182 | 7.369822 | 2.415613 | | | 11 |
| 978 | ENSG00000228412 | 113.2625 | -1.05185 | 0.352326 | -2.98543 | 0.002832 | 0.028883 | 11.12499 | 4.543267 | 8.120215 | 4.210575 | LNC-LBCS | | 6 |
| 979 | ENSG00000228623 | 196.5636 | -1.28601 | 0.277261 | -4.63826 | 3.51E-06 | 8.21E-05 | 17.29736 | 6.064824 | 13.7505 | 5.936158 | ZNF883 | | 9 |
| 980 | ENSG00000228999 | 57.15538 | -2.5394 | 0.506399 | -5.01463 | 5.31E-07 | 1.47E-05 | 19.47988 | 2.706267 | 20.02872 | 3.719773 | | | 2 |
| 981 | ENSG00000229186 | 19.24249 | -7.76916 | 1.642782 | -4.72927 | 2.25E-06 | 5.46E-05 | 2.740603 | 0 | 1.687492 | 0 | ADAM1A | | 12 |
| 982 | ENSG00000229563 | 11.97298 | -4.07187 | 1.302878 | -3.12529 | 0.001776 | 0.020054 | 1.884201 | 0.082858 | 0.613983 | 0.051842 | LINC01204 | X | |
| 983 | ENSG00000229618 | 74.10162 | 4.743142 | 0.615624 | 7.704604 | 1.31E-14 | 9.83E-13 | 0.108773 | 6.136267 | 0.321473 | 7.132254 | | | 7 |
| 984 | ENSG00000229743 | 221.7652 | 1.191009 | 0.279232 | 4.265309 | 2.00E-05 | 0.000395 | 11.03746 | 21.73428 | 7.646129 | 18.19611 | LINC0115S | | 2 |
| 985 | ENSG00000229847 | 431.6488 | -2.13026 | 0.265846 | -8.01314 | 1.12E-15 | 9.30E-14 | 10.13991 | 3.215135 | 17.1649 | 2.670658 | EMX2OS | | 10 |
| 986 | ENSG00000230316 | 786.1363 | -3.99094 | 0.461326 | -8.65101 | 5.10E-18 | 5.37E-16 | 209.7186 | 17.05682 | 232.4075 | 9.071375 | FEZF1-AS1 | | 7 |
| 987 | ENSG00000231419 | 24.39676 | -2.75708 | 0.771592 | -3.57324 | 0.000353 | 0.005111 | 1.246387 | 0.16275 | 1.02572 | 0.15368 | LINC0068S | | 7 |
| 988 | ENSG00000231424 | 13.76116 | -3.7008 | 1.18323 | -3.12771 | 0.001762 | 0.019913 | 2.139558 | 0.047737 | 0.770394 | 0.169022 | | | 1 |
| 989 | ENSG00000231431 | 19.46994 | 2.394759 | 0.853841 | 2.804689 | 0.005037 | 0.045849 | 0.24443 | 1.535159 | 0.485307 | 2.136175 | FAR2P4 | | 2 |
| 990 | ENSG00000231764 | 244.6252 | -6.17616 | 1.315819 | -4.69378 | 2.68E-06 | 6.41E-05 | 14.83921 | 0.327136 | 10.66855 | 0.027321 | DLX6-AS1 | | 7 |
| 991 | ENSG00000231806 | 74.62579 | -2.99397 | 0.474986 | -6.30329 | 2.91E-10 | 1.39E-08 | 4.744056 | 0.401488 | 3.675411 | 0.616071 | PCAT7 | | 9 |
| 992 | ENSG00000232928 | 84.79133 | -1.10988 | 0.389853 | -2.84692 | 0.004414 | 0.04156 | 7.676539 | 3.024273 | 7.026204 | 3.41351 | DDX3P1 | X | |
| 993 | ENSG00000233058 | 71.39828 | -2.87059 | 0.481227 | -5.96514 | 2.44E-09 | 1.01E-07 | 5.761546 | 0.763009 | 4.802706 | 0.622591 | ATP13A3-[ | | 3 |
| 994 | ENSG00000233117 | 24.1385 | 4.398568 | 0.989726 | 4.444227 | 8.82E-06 | 0.00019 | 0.304416 | 3.698505 | 0.072067 | 3.23431 | LINC00702 | | 10 |
| 995 | ENSG00000233237 | 2960.89 | 1.520489 | 0.136914 | 11.10541 | 1.18E-28 | 2.96E-26 | 29.16506 | 87.78835 | 31.88412 | 77.57023 | LINC00472 | | 6 |
| 996 | ENSG00000233429 | 14.18878 | 2.961457 | 1.046958 | 2.82863 | 0.004675 | 0.043405 | 0.746527 | 4.296599 | 0.801311 | 7.138 | HOTAIRM1 | | 7 |
| 997 | ENSG00000233587 | 22.01342 | -2.4957 | 0.802286 | -3.11074 | 0.001866 | 0.020864 | 2.908625 | 0.487427 | 4.171963 | 0.707314 | LINC01884 | | 2 |
| 998 | ENSG00000233639 | 807.8141 | 1.634212 | 0.167898 | 9.733334 | 2.17E-22 | 3.22E-20 | 35.40413 | 107.2197 | 40.43336 | 115.3677 | PANTR1 | | 2 |
| 999 | ENSG00000233930 | 141.1093 | -1.02812 | 0.334233 | -3.07606 | 0.002098 | 0.023019 | 12.48312 | 5.519981 | 8.699601 | 4.244609 | KRTAP5-AS | | 11 |
| 1000 | ENSG00000234173 | 15.20269 | -5.9694 | 1.673741 | -3.5665 | 0.000362 | 0.005213 | 1.285481 | 0.045594 | 1.855927 | 0 | | | 10 |
| 1001 | ENSG00000234444 | 14.30697 | -6.7752 | 1.7172 | -3.9455 | 7.96E-05 | 0.001368 | 0.778234 | 0 | 1.81269 | 0 | ZNF736 | | 7 |
| 1002 | ENSG00000234745 | 538.3828 | 1.433251 | 0.184781 | 7.756471 | 8.73E-15 | 6.67E-13 | 26.61611 | 69.78259 | 27.17015 | 67.41093 | HLA-B | | 6 |
| 1003 | ENSG00000235288 | 77.06909 | 1.629318 | 0.489321 | 3.329757 | 0.000869 | 0.010973 | 20.52374 | 38.76494 | 6.86327 | 40.30722 | | | 3 |
| 1004 | ENSG00000235597 | 45.21625 | 2.016587 | 0.545978 | 3.693533 | 0.000221 | 0.003379 | 1.386057 | 4.714899 | 1.261612 | 5.416203 | LINC01102 | | 2 |
| 1005 | ENSG00000235669 | 11.74503 | -3.92228 | 1.304265 | -3.00727 | 0.002636 | 0.027258 | 4.995422 | 0 | 3.673081 | 0.556321 | CHN2-AS1 | | 7 |
| 1006 | ENSG00000236107 | 84.23647 | -1.16676 | 0.401166 | -2.90843 | 0.003632 | 0.035533 | 4.136487 | 2.117128 | 4.91688 | 1.689399 | SCN1A-AS | | 2 |
| 1007 | ENSG00000236668 | 17.57639 | -4.54683 | 1.15109 | -3.95002 | 7.81E-05 | 0.001348 | 2.337102 | 0.049299 | 1.505112 | 0.1074 | LHX2-AS1 | | 9 |
| 1008 | ENSG00000237065 | 26.92117 | -3.53248 | 0.823312 | -4.29057 | 1.78E-05 | 0.000357 | 6.652237 | 0.987873 | 9.422244 | 0.306697 | NANOGP4 | | 7 |
| 1009 | ENSG00000237149 | 91.5996 | -1.22439 | 0.393343 | -3.11278 | 0.001853 | 0.02076 | 13.17563 | 5.211201 | 9.839274 | 4.052762 | ZNF503-AS | | 10 |
| 1010 | ENSG00000237412 | 23.46844 | 4.880173 | 1.194326 | 4.086131 | 4.39E-05 | 0.000805 | 0 | 4.330233 | 0.159891 | 1.078264 | PRSS56 | | 2 |
| 1011 | ENSG00000237440 | 623.1377 | -2.18255 | 0.201754 | -10.8179 | 2.83E-27 | 6.09E-25 | 34.03846 | 5.787255 | 34.58539 | 8.588035 | ZNF737 | | 19 |
| 1012 | ENSG00000237654 | 21.35428 | -2.94555 | 0.827814 | -3.55823 | 0.000373 | 0.005356 | 0.969974 | 0.183 | 1.749714 | 0.149607 | | | 11 |
| 1013 | ENSG00000237732 | 15.14637 | 2.72344 | 0.974191 | 2.795592 | 0.00518 | 0.046898 | 0.794533 | 5.022599 | 1.015937 | 6.228378 | CT75 | | 2 |

| | | | | | | | | | | | | | | |
|---|---|---|---|---|---|---|---|---|---|---|---|---|---|---|
| 1014 | ENSG00000237807 | 36.83258 | -2.26772 | 0.622078 | -3.6454 | 0.000267 | 0.003988 | 1.912986 | 0.503489 | 2.751239 | 0.410649 | LINC02984 | | 8 |
| 1015 | ENSG00000237863 | 44.10587 | -3.74624 | 0.642788 | -5.82812 | 5.61E-09 | 2.18E-07 | 7.319675 | 0.603705 | 7.187417 | 0.41071 | IRS4-AS1 | X | |
| 1016 | ENSG00000237877 | 95.25555 | -1.22367 | 0.383411 | -3.19154 | 0.001415 | 0.01661 | 14.80639 | 4.768188 | 10.37057 | 5.400449 | LINC01473 | | 2 |
| 1017 | ENSG00000238194 | 21.27619 | -2.30596 | 0.822159 | -2.80476 | 0.005035 | 0.045849 | 2.071327 | 0.522974 | 1.802091 | 0.207158 | PHACTR3-. | | 20 |
| 1018 | ENSG00000239519 | 67.99328 | -2.0268 | 0.457457 | -4.43058 | 9.40E-06 | 0.0002 | 5.677486 | 1.204191 | 4.128456 | 1.058106 | CADM2-AS | | 3 |
| 1019 | ENSG00000239922 | 402.375 | -1.46393 | 0.245646 | -5.95952 | 2.53E-09 | 1.05E-07 | 44.56432 | 18.63807 | 69.39045 | 20.5372 | | | 3 |
| 1020 | ENSG00000240225 | 186.5401 | -3.00175 | 0.310336 | -9.67255 | 3.94E-22 | 5.76E-20 | 16.26412 | 1.583489 | 17.02895 | 2.39829 | ZNF542P | | 19 |
| 1021 | ENSG00000240476 | 11.0457 | -3.85856 | 1.350416 | -2.85731 | 0.004272 | 0.040614 | 4.354996 | 0.394195 | 1.932847 | 0 | LINC00973 | | 3 |
| 1022 | ENSG00000241231 | 176.2401 | -1.62472 | 0.30469 | -5.33236 | 9.69E-08 | 3.10E-06 | 9.223362 | 3.506283 | 12.6835 | 3.217559 | | | 3 |
| 1023 | ENSG00000241743 | 1399.107 | 2.54111 | 0.627246 | 4.051217 | 5.10E-05 | 0.000919 | 0.114779 | 1.155997 | 0.211663 | 0.633697 | XACT | X | |
| 1024 | ENSG00000242808 | 706.1327 | -1.36919 | 0.174387 | -7.85146 | 4.11E-15 | 3.22E-13 | 80.1935 | 28.06878 | 87.03606 | 33.23107 | SOX2-OT | | 3 |
| 1025 | ENSG00000243244 | 512.2664 | -1.27433 | 0.194026 | -6.56782 | 5.11E-11 | 2.65E-09 | 17.7936 | 6.507208 | 14.72138 | 6.156538 | STON1 | | 2 |
| 1026 | ENSG00000243854 | 13.96875 | -3.41178 | 1.080893 | -3.15645 | 0.001597 | 0.01837 | 41.02004 | 2.888247 | 43.2479 | 4.639272 | RN7SL67P | | 12 |
| 1027 | ENSG00000243915 | 54.80116 | -1.98976 | 0.516021 | -3.85597 | 0.000115 | 0.001915 | 5.87908 | 1.112169 | 3.510649 | 1.110582 | THAP12P2 | | 3 |
| 1028 | ENSG00000244227 | 35.30379 | -1.76463 | 0.634102 | -2.78289 | 0.005388 | 0.04846 | 3.349006 | 1.560784 | 5.851418 | 1.026579 | LRRC77P | | 3 |
| 1029 | ENSG00000245526 | 2617.886 | -1.30945 | 0.1329 | -9.85293 | 6.66E-23 | 1.07E-20 | 151.0058 | 57.664 | 137.4534 | 52.07469 | LINC00461 | | 5 |
| 1030 | ENSG00000245532 | 5233.59 | 2.143232 | 0.115929 | 18.48743 | 2.61E-76 | 4.25E-73 | 14.28197 | 56.17077 | 13.14836 | 58.25389 | NEAT1 | | 11 |
| 1031 | ENSG00000245680 | 296.4958 | -1.69116 | 0.234932 | -7.19851 | 6.09E-13 | 3.82E-11 | 15.50075 | 4.211039 | 16.44075 | 5.187868 | ZNF585B | | 19 |
| 1032 | ENSG00000245694 | 1504.542 | 1.657127 | 0.147649 | 11.2234 | 3.13E-29 | 8.16E-27 | 26.37661 | 76.41179 | 29.11357 | 89.37408 | CRNDE | | 16 |
| 1033 | ENSG00000246214 | 58.4317 | -2.38832 | 0.495155 | -4.82337 | 1.41E-06 | 3.56E-05 | 5.898335 | 1.214656 | 7.008607 | 1.114636 | RETREG1-/ | | 5 |
| 1034 | ENSG00000247828 | 2922.9 | -1.13507 | 0.132422 | -8.57162 | 1.02E-17 | 1.04E-15 | 86.96517 | 33.40229 | 72.03094 | 34.9124 | TMEM161I | | 5 |
| 1035 | ENSG00000248445 | 155.3495 | -1.62017 | 0.307692 | -5.26556 | 1.40E-07 | 4.30E-06 | 62.04852 | 21.74327 | 66.92926 | 17.86384 | SEMA6A-A | | 5 |
| 1036 | ENSG00000248485 | 53.32422 | -2.05693 | 0.509725 | -4.03537 | 5.45E-05 | 0.000979 | 8.067697 | 1.455952 | 7.95922 | 2.200808 | PCP4L1 | | 1 |
| 1037 | ENSG00000248905 | 209.1115 | 2.298184 | 0.342493 | 6.710169 | 1.94E-11 | 1.04E-09 | 1.69818 | 7.29956 | 0.912179 | 4.573874 | FMN1 | | 15 |
| 1038 | ENSG00000249860 | 19.78652 | -3.43129 | 0.918061 | -3.73754 | 0.000186 | 0.002891 | 2.331661 | 0.195173 | 3.090071 | 0.283342 | NA | NA | |
| 1039 | ENSG00000250033 | 37.13601 | 2.871862 | 0.625249 | 4.593153 | 4.37E-06 | 0.0001 | 0.381259 | 2.26017 | 0.239861 | 2.008704 | SLC7A11-/ | | 4 |
| 1040 | ENSG00000250208 | 337.1721 | 3.64192 | 0.712388 | 5.112267 | 3.18E-07 | 9.22E-06 | 0.555658 | 9.716777 | 1.499137 | 14.74621 | FZD10-AS | | 12 |
| 1041 | ENSG00000250337 | 415.0172 | -1.60038 | 0.206709 | -7.74216 | 9.77E-15 | 7.44E-13 | 53.04854 | 16.33537 | 58.9281 | 18.62434 | PURPL | | 5 |
| 1042 | ENSG00000250358 | 24.62996 | -2.57491 | 0.77817 | -3.30893 | 0.000937 | 0.011709 | 3.91585 | 0.940789 | 6.693927 | 0.755989 | LINC02200 | | 5 |
| 1043 | ENSG00000250420 | 17.14718 | -4.23887 | 1.150537 | -3.68426 | 0.000229 | 0.003493 | 1.44442 | 0.147028 | 1.830891 | 0.039594 | AACSP1 | | 5 |
| 1044 | ENSG00000250486 | 25.73316 | -2.21628 | 0.721512 | -3.07171 | 0.002128 | 0.023286 | 2.550844 | 0.437007 | 2.196172 | 0.528918 | FAM218A | | 4 |
| 1045 | ENSG00000251632 | 21.82937 | 5.352683 | 1.240323 | 4.315557 | 1.59E-05 | 0.000322 | 0.059473 | 2.134834 | 0.052476 | 2.173914 | LINC02172 | | 4 |
| 1046 | ENSG00000253284 | 778.9446 | -1.31106 | 0.164611 | -7.9646 | 1.66E-15 | 1.36E-13 | 22.70243 | 8.396991 | 23.85427 | 9.352315 | | | 12 |
| 1047 | ENSG00000253368 | 121.9723 | 1.00988 | 0.357876 | 2.821868 | 0.004774 | 0.043996 | 5.740902 | 12.88522 | 5.511034 | 8.449207 | TRNP1 | | 1 |
| 1048 | ENSG00000253537 | 40.92099 | -1.60363 | 0.559387 | -2.86677 | 0.004147 | 0.039652 | 1.989921 | 0.594624 | 2.188558 | 0.710071 | PCDHGA7 | | 5 |
| 1049 | ENSG00000253661 | 789.1259 | -1.50551 | 0.176006 | -8.55378 | 1.19E-17 | 1.21E-15 | 56.44932 | 15.67885 | 45.30924 | 18.15195 | ZFHX4-AS | | 8 |
| 1050 | ENSG00000253706 | 23.68841 | -2.29699 | 0.804553 | -2.85499 | 0.004304 | 0.040793 | 3.083024 | 0.544042 | 1.519633 | 0.313619 | | | 8 |
| 1051 | ENSG00000253767 | 90.14233 | -2.15056 | 0.405048 | -5.3094 | 1.10E-07 | 3.46E-06 | 2.272332 | 0.539919 | 3.031427 | 0.591254 | PCDHGA8 | | 5 |
| 1052 | ENSG00000253953 | 241.2194 | -4.00562 | 0.300393 | -13.3346 | 1.46E-40 | 8.38E-38 | 10.86014 | 0.643505 | 10.90324 | 0.635629 | PCDHGB4 | | 5 |
| 1053 | ENSG00000254186 | 62.44153 | 1.985034 | 0.471819 | 4.207197 | 2.59E-05 | 0.000497 | 1.735529 | 6.223144 | 1.578278 | 6.195082 | | | 5 |
| 1054 | ENSG00000254187 | 49.07979 | 1.602254 | 0.523564 | 3.060283 | 0.002211 | 0.023969 | 2.459206 | 7.919155 | 3.510876 | 9.311116 | | | 5 |
| 1055 | ENSG00000254245 | 100.817 | -4.3257 | 0.481475 | -8.98427 | 2.60E-19 | 3.12E-17 | 5.329961 | 0.307644 | 6.126512 | 0.20681 | PCDHGA3 | | 5 |
| 1056 | ENSG00000254305 | 27.43409 | -2.74086 | 0.77791 | -3.52336 | 0.000426 | 0.005986 | 16.96924 | 3.094841 | 11.39444 | 0.838265 | MRPL9P1 | | 8 |
| 1057 | ENSG00000254579 | 40.54295 | 1.866262 | 0.576627 | 3.236513 | 0.00121 | 0.014586 | 0.752209 | 1.99656 | 0.532962 | 2.37706 | | | 11 |
| 1058 | ENSG00000254622 | 23.25754 | 2.207361 | 0.768368 | 2.872792 | 0.004069 | 0.039018 | 0.769805 | 4.420077 | 1.136381 | 3.933873 | NAV2-AS4 | | 11 |
| 1059 | ENSG00000254656 | 1482.853 | 2.008982 | 0.167028 | 12.02781 | 2.54E-33 | 9.02E-31 | 11.58045 | 40.09464 | 13.11124 | 54.15782 | RTL1 | | 14 |
| 1060 | ENSG00000255043 | 95.86086 | 2.431922 | 0.392241 | 6.200077 | 5.64E-10 | 2.60E-08 | 1.176405 | 5.379881 | 1.03684 | 5.891055 | NAV2-AS5 | | 11 |
| 1061 | ENSG00000255408 | 129.9537 | -1.02569 | 0.325985 | -3.14644 | 0.001653 | 0.018877 | 4.362144 | 2.130503 | 4.048599 | 1.757118 | PCDHA3 | | 5 |
| 1062 | ENSG00000255545 | 89.73621 | -1.22977 | 0.388942 | -3.16184 | 0.001568 | 0.018097 | 7.479723 | 2.574406 | 5.550077 | 2.654051 | B3GAT1-D | | 11 |
| 1063 | ENSG00000256463 | 447.511 | -3.60074 | 0.605449 | -5.94722 | 2.73E-09 | 1.12E-07 | 26.64426 | 2.194816 | 15.82383 | 1.088389 | SALL3 | | 18 |
| 1064 | ENSG00000257056 | 51.7741 | -4.31467 | 0.684232 | -6.30585 | 2.87E-10 | 1.37E-08 | 8.563152 | 0.225006 | 4.310055 | 0.391032 | LINC02282 | | 14 |
| 1065 | ENSG00000257126 | 46.1928 | -5.78716 | 0.886514 | -6.528 | 6.67E-11 | 3.40E-09 | 3.822737 | 0.02585 | 2.525478 | 0.084525 | FOXG1-AS | | 14 |
| 1066 | ENSG00000257501 | 93.45168 | 2.400574 | 0.484692 | 4.952787 | 7.32E-07 | 1.94E-05 | 1.12161 | 12.7754 | 3.954477 | 12.40195 | PAFAH1B2 | | 12 |
| 1067 | ENSG00000257522 | 10.96922 | -6.59914 | 1.764136 | -3.74072 | 0.000183 | 0.002864 | 1.119599 | 0 | 0.951984 | 0 | | | 14 |
| 1068 | ENSG00000257545 | 127.6274 | 1.536421 | 0.342009 | 4.49234 | 7.04E-06 | 0.000156 | 5.238112 | 10.97269 | 4.136193 | 14.71577 | | | 12 |
| 1069 | ENSG00000257594 | 91.98373 | 1.48585 | 0.473568 | 3.137568 | 0.001704 | 0.019369 | 0.490552 | 2.838661 | 1.522204 | 2.539184 | GALNT4 | | 12 |
| 1070 | ENSG00000257711 | 315.5033 | 1.537526 | 0.225724 | 6.811526 | 9.66E-12 | 5.38E-10 | 5.290187 | 13.15551 | 4.596281 | 13.91836 | | | 12 |
| 1071 | ENSG00000257800 | 42.42481 | -7.46173 | 1.525678 | -4.89076 | 1.00E-06 | 2.60E-05 | 5.217781 | 0.053599 | 5.212325 | 0 | FNBP1P1 | | 2 |
| 1072 | ENSG00000257935 | 76.10499 | -3.81692 | 0.501985 | -7.60365 | 2.88E-14 | 2.11E-12 | 61.89083 | 3.434571 | 50.24268 | 4.086156 | LHX5-AS1 | | 12 |
| 1073 | ENSG00000257986 | 17.87576 | 3.346045 | 0.99503 | 3.362758 | 0.000772 | 0.009908 | 0.533409 | 3.623828 | 0.131793 | 2.953861 | LINC02306 | | 14 |
| 1074 | ENSG00000258399 | 27.32898 | -5.20322 | 1.089949 | -4.77383 | 1.81E-06 | 4.48E-05 | 2.070305 | 0 | 4.508314 | 0.176037 | MIR493HG | | 14 |
| 1075 | ENSG00000258498 | 139.4875 | 1.513125 | 0.35326 | 4.283323 | 1.84E-05 | 0.000367 | 3.73317 | 10.90828 | 5.819583 | 15.69048 | DIO3OS | | 14 |
| 1076 | ENSG00000258548 | 145.8429 | -2.35546 | 0.33681 | -6.99344 | 2.68E-12 | 1.57E-10 | 12.45084 | 2.13618 | 8.838581 | 1.7758 | LINC00645 | | 14 |
| 1077 | ENSG00000258555 | 11.28328 | -7.00287 | 1.772554 | -3.95072 | 7.79E-05 | 0.001345 | 0.282086 | 0 | 0.532198 | 0 | SPECC1L-/ | | 22 |
| 1078 | ENSG00000258611 | 81.21892 | -1.16981 | 0.407188 | -2.87289 | 0.004067 | 0.039018 | 17.17851 | 5.581245 | 14.27672 | 7.65068 | YBX2P2 | | 15 |
| 1079 | ENSG00000259439 | 153.9591 | -3.01005 | 0.33642 | -8.9473 | 3.64E-19 | 4.29E-17 | 21.63765 | 2.805743 | 20.19293 | 1.915478 | LINC01833 | | 2 |
| 1080 | ENSG00000259498 | 108.1184 | 2.477204 | 0.386789 | 6.404533 | 1.51E-10 | 7.39E-09 | 1.28014 | 7.165429 | 1.157282 | 5.577904 | TPM1-AS | | 15 |
| 1081 | ENSG00000259571 | 18.13932 | 2.755452 | 0.89783 | 3.069014 | 0.002148 | 0.023422 | 0.761344 | 5.23179 | 0.845827 | 5.034122 | BLID | | 11 |
| 1082 | ENSG00000259867 | 103.6712 | -1.29767 | 0.439321 | -2.9538 | 0.003139 | 0.031336 | 5.480617 | 3.618717 | 13.00411 | 3.488666 | DYNLRB2-. | | 16 |
| 1083 | ENSG00000259871 | 18.02219 | 3.155489 | 0.947336 | 3.33091 | 0.000866 | 0.010934 | 0.322233 | 3.528593 | 0.380102 | 2.377152 | | | 16 |
| 1084 | ENSG00000260293 | 103.7953 | -1.08061 | 0.357521 | -3.0225 | 0.002507 | 0.026356 | 2.727302 | 1.149262 | 2.932991 | 1.385805 | | | 16 |
| 1085 | ENSG00000260398 | 111.3225 | -1.27973 | 0.360444 | -3.55043 | 0.000385 | 0.005493 | 5.440651 | 2.133518 | 4.281315 | 1.627763 | | | 8 |
| 1086 | ENSG00000260412 | 63.19936 | -1.53474 | 0.4642 | -3.30621 | 0.000946 | 0.011801 | 1.429375 | 0.577716 | 1.624682 | 0.414799 | | | 9 |
| 1087 | ENSG00000260578 | 67.25763 | 2.644473 | 0.497873 | 5.311547 | 1.09E-07 | 3.43E-06 | 0.450629 | 4.375645 | 0.902741 | 3.689461 | | | 18 |
| 1088 | ENSG00000260641 | 54.62872 | -2.19621 | 0.591131 | -3.71527 | 0.000203 | 0.003131 | 2.667197 | 0.450138 | 0.939427 | 0.283119 | TSPAN5-D | | 4 |
| 1089 | ENSG00000260645 | 141.4802 | 1.157165 | 0.345044 | 3.353678 | 0.000797 | 0.010192 | 4.411763 | 8.13453 | 5.643008 | 13.20925 | | | 6 |
| 1090 | ENSG00000260691 | 63.36185 | -1.65157 | 0.45975 | -3.59231 | 0.000328 | 0.004793 | 6.05522 | 1.86319 | 5.161553 | 1.50614 | ANKRD20/ | | 9 |
| 1091 | ENSG00000261150 | 312.9707 | 1.946993 | 0.275585 | 7.06495 | 1.61E-12 | 9.55E-11 | 0.754218 | 3.848544 | 1.005429 | 2.554386 | EPPK1 | | 8 |

| | | | | | | | | | | | | | |
|---|---|---|---|---|---|---|---|---|---|---|---|---|---|
| 1092 | ENSG00000261210 | 68.97162 | 3.104336 | 0.589492 | 5.266121 | 1.39E-07 | 4.30E-06 | 0.208734 | 7.019088 | 1.399466 | 5.980901 | CLEC19A | 16 |
| 1093 | ENSG00000261716 | 469.3136 | -1.04295 | 0.196425 | -5.30965 | 1.10E-07 | 3.46E-06 | 11.6676 | 5.324492 | 10.40866 | 4.772577 | H2BC20P | 1 |
| 1094 | ENSG00000261934 | 97.22729 | -1.27675 | 0.40059 | -3.18717 | 0.001437 | 0.016813 | 4.748244 | 1.218088 | 2.893218 | 1.755448 | PCDHGA9 | 5 |
| 1095 | ENSG00000262096 | 79.11328 | -1.31813 | 0.454259 | -2.9017 | 0.003711 | 0.036232 | 6.84324 | 2.208407 | 3.503466 | 1.673416 | PCDHB19P | 5 |
| 1096 | ENSG00000262655 | 179.5517 | -2.59362 | 0.324866 | -7.98364 | 1.42E-15 | 1.17E-13 | 10.25167 | 1.352007 | 6.624174 | 1.272666 | SPON1 | 11 |
| 1097 | ENSG00000263146 | 32.33322 | -4.80002 | 0.803221 | -5.97597 | 2.29E-09 | 9.56E-08 | 9.818566 | 0.371837 | 10.17185 | 0.303418 | LINC01896 | 18 |
| 1098 | ENSG00000263424 | 78.77809 | 2.119525 | 0.445653 | 4.755994 | 1.97E-06 | 4.85E-05 | 1.341143 | 4.704905 | 0.764855 | 3.853955 | | 18 |
| 1099 | ENSG00000263711 | 47.32452 | -9.03466 | 1.526607 | -5.91813 | 3.26E-09 | 1.32E-07 | 6.873184 | 0 | 7.900824 | 0 | LINC02864 | 18 |
| 1100 | ENSG00000263958 | 15.17558 | 3.347123 | 1.006714 | 3.3248 | 0.000885 | 0.011127 | 0.265648 | 1.532042 | 0.078066 | 1.714288 | NETO1-DT | 18 |
| 1101 | ENSG00000265107 | 82.74437 | 3.634933 | 0.56563 | 6.426346 | 1.31E-10 | 6.47E-09 | 0.37209 | 8.156867 | 0.577557 | 3.125089 | GJA5 | 1 |
| 1102 | ENSG00000267165 | 19.20654 | 3.531337 | 1.019466 | 3.46391 | 0.000532 | 0.007234 | 0 | 12.96561 | 2.099441 | 10.69121 | CHMP1B-/ | 18 |
| 1103 | ENSG00000267254 | 125.032 | -1.4605 | 0.335587 | -4.35209 | 1.35E-05 | 0.000279 | 11.44919 | 3.376013 | 11.23157 | 4.457595 | ZNF790-A! | 19 |
| 1104 | ENSG00000267313 | 105.4734 | 1.515374 | 0.382946 | 3.957146 | 7.59E-05 | 0.001315 | 6.907109 | 12.9335 | 4.310351 | 16.93824 | | 18 |
| 1105 | ENSG00000267586 | 48.89638 | 1.616388 | 0.549321 | 2.942517 | 0.003256 | 0.032331 | 2.835082 | 5.787267 | 1.256085 | 5.868287 | LINC00907 | 18 |
| 1106 | ENSG00000267605 | 38.29538 | -2.11363 | 0.594595 | -3.55474 | 0.000378 | 0.005416 | 3.276027 | 0.659784 | 3.81985 | 0.892736 | | 19 |
| 1107 | ENSG00000267640 | 68.28171 | -8.71352 | 1.507393 | -5.78052 | 7.45E-09 | 2.85E-07 | 7.756626 | 0 | 4.904057 | 0 | | 19 |
| 1108 | ENSG00000268119 | 144.7209 | -6.21228 | 0.565656 | -10.9824 | 4.64E-28 | 1.09E-25 | 15.51492 | 0.231465 | 14.776 | 0.145156 | | 19 |
| 1109 | ENSG00000268654 | 17.38122 | -7.59355 | 1.66063 | -4.57269 | 4.82E-06 | 0.00011 | 4.862374 | 0 | 3.114315 | 0 | MIMT1 | 19 |
| 1110 | ENSG00000268658 | 33.16496 | -6.2885 | 1.177549 | -5.34033 | 9.28E-08 | 2.98E-06 | 2.627486 | 0.026623 | 1.983375 | 0.029018 | LINC00664 | 19 |
| 1111 | ENSG00000269113 | 49.27768 | 2.837811 | 0.570352 | 4.975541 | 6.51E-07 | 1.74E-05 | 0.23093 | 1.066126 | 0.130679 | 1.358499 | TRABD2B | 1 |
| 1112 | ENSG00000269699 | 47.2034 | -7.82782 | 1.505499 | -5.19948 | 2.00E-07 | 6.00E-06 | 8.607736 | 0.054506 | 5.790392 | 0 | ZIM2 | 19 |
| 1113 | ENSG00000269834 | 102.7777 | -2.30292 | 0.380982 | -6.0447 | 1.50E-09 | 6.46E-08 | 9.202732 | 1.688027 | 7.397544 | 1.475527 | ZNF528-A! | 19 |
| 1114 | ENSG00000270011 | 53.82596 | -1.58115 | 0.494542 | -3.19719 | 0.001388 | 0.016317 | 5.523117 | 1.998709 | 5.586772 | 1.522285 | ZNF559-ZI | 19 |
| 1115 | ENSG00000270953 | 123.7662 | -1.76949 | 0.349872 | -5.05755 | 4.25E-07 | 1.20E-05 | 65.13124 | 15.0601 | 45.49148 | 15.50316 | | 7 |
| 1116 | ENSG00000271848 | 11.4428 | -3.22342 | 1.158504 | -2.7824 | 0.005396 | 0.048511 | 1.257108 | 0.080536 | 0.834855 | 0.13162 | SYNPO2L-. | 10 |
| 1117 | ENSG00000272078 | 44.87032 | 2.123063 | 0.557423 | 3.808712 | 0.00014 | 0.002261 | 0.265843 | 0.85238 | 0.20647 | 1.088715 | | 1 |
| 1118 | ENSG00000272449 | 69.56014 | -1.68757 | 0.435837 | -3.87202 | 0.000108 | 0.001802 | 14.71732 | 4.153612 | 13.18325 | 4.013248 | | 1 |
| 1119 | ENSG00000272636 | 166.9865 | -1.17689 | 0.317047 | -3.71205 | 0.000206 | 0.003165 | 5.263425 | 2.620249 | 4.939232 | 1.665087 | DOC2B | 17 |
| 1120 | ENSG00000273213 | 9.650232 | -4.53534 | 1.605351 | -2.82514 | 0.004726 | 0.043715 | 0.376489 | 0.096115 | 2.37581 | 0 | H3-7 | 1 |
| 1121 | ENSG00000273507 | 86.74092 | 1.306914 | 0.458456 | 2.850688 | 0.004362 | 0.04119 | 1.532511 | 3.819105 | 0.948873 | 1.921859 | | 13 |
| 1122 | ENSG00000273703 | 148.6625 | -3.67308 | 0.367673 | -9.99007 | 1.68E-23 | 2.92E-21 | 120.4654 | 8.52492 | 148.5567 | 11.50927 | H2BC14 | 6 |
| 1123 | ENSG00000274317 | 598.0907 | 1.346551 | 0.211895 | 6.3548 | 2.09E-10 | 1.01E-08 | 10.66378 | 28.36476 | 9.6126 | 20.11601 | LINC02334 | 13 |
| 1124 | ENSG00000274588 | 40.09864 | -1.62362 | 0.576488 | -2.8164 | 0.004857 | 0.044571 | 1.093176 | 0.27086 | 0.714422 | 0.281263 | DGKK | X |
| 1125 | ENSG00000274600 | 8.745057 | -6.63109 | 1.854475 | -3.57572 | 0.000349 | 0.00507 | 0.430278 | 0 | 0.21648 | 0 | RIMBP3B | 22 |
| 1126 | ENSG00000274810 | 37.50603 | 2.099662 | 0.625486 | 3.356847 | 0.000788 | 0.010095 | 0.509636 | 1.918288 | 0.266355 | 1.127995 | NPHP3-AC | 3 |
| 1127 | ENSG00000275126 | 163.8625 | -2.61115 | 0.312692 | -8.35055 | 6.79E-17 | 6.25E-15 | 201.4165 | 30.1758 | 221.2278 | 35.3283 | H4C13 | 6 |
| 1128 | ENSG00000275180 | 90.74746 | 1.084114 | 0.379494 | 2.856733 | 0.00428 | 0.040669 | 24.32314 | 49.88709 | 27.20523 | 53.49637 | | 12 |
| 1129 | ENSG00000275379 | 519.2545 | -2.38533 | 0.195377 | -12.2089 | 2.79E-34 | 1.14E-31 | 355.555 | 68.48999 | 384.2666 | 65.23449 | H3C11 | 6 |
| 1130 | ENSG00000275993 | 495.4162 | 1.570956 | 0.222451 | 7.062042 | 1.64E-12 | 9.69E-11 | 4.907316 | 17.27428 | 7.396866 | 17.35046 | | 21 |
| 1131 | ENSG00000276368 | 206.3224 | -2.84698 | 0.288656 | -9.86286 | 6.03E-23 | 9.75E-21 | 160.3749 | 22.64087 | 181.2888 | 22.23495 | H2AC14 | 6 |
| 1132 | ENSG00000276547 | 70.72169 | -2.32 | 0.453148 | -5.11974 | 3.06E-07 | 8.92E-06 | 4.103321 | 0.853495 | 4.073069 | 0.673272 | PCDHGB5 | 5 |
| 1133 | ENSG00000277268 | 35.88589 | -2.71539 | 0.669913 | -4.05335 | 5.05E-05 | 0.000912 | 11.95085 | 1.189861 | 6.718036 | 1.533523 | LHX1-DT | 17 |
| 1134 | ENSG00000277534 | 274.5677 | 1.389828 | 0.2511 | 5.534968 | 3.11E-08 | 1.08E-06 | 5.407023 | 15.95804 | 7.274542 | 15.52693 | | 18 |
| 1135 | ENSG00000278570 | 21.79792 | -2.52169 | 0.837692 | -3.01028 | 0.00261 | 0.027089 | 3.191748 | 0.951715 | 4.084639 | 0.340782 | NR2E3 | 15 |
| 1136 | ENSG00000278893 | 25.12047 | -2.08597 | 0.742486 | -2.80945 | 0.004963 | 0.045304 | 3.034199 | 0.554286 | 1.709346 | 0.493975 | | 16 |
| 1137 | ENSG00000278909 | 53.35151 | 2.012408 | 0.516414 | 3.896891 | 9.74E-05 | 0.001642 | 0.577267 | 2.940048 | 0.881853 | 2.648326 | | 16 |
| 1138 | ENSG00000278931 | 16.96877 | 2.654824 | 0.930484 | 2.853164 | 0.004329 | 0.040954 | 0.248786 | 1.74925 | 0.439601 | 2.398399 | | 21 |
| 1139 | ENSG00000278975 | 33.72423 | 2.093228 | 0.674401 | 3.10383 | 0.00191 | 0.021222 | 0.503483 | 3.474679 | 1.408686 | 4.354256 | | 16 |
| 1140 | ENSG00000279050 | 31.15 | -2.2805 | 0.723218 | -3.15327 | 0.001615 | 0.018506 | 2.103665 | 0.215955 | 3.027795 | 0.79992 | PWAR1 | 15 |
| 1141 | ENSG00000279289 | 494.3373 | 1.560703 | 0.21035 | 7.419567 | 1.18E-13 | 7.87E-12 | 6.722394 | 22.33389 | 7.736174 | 17.9228 | | 6 |
| 1142 | ENSG00000279358 | 76.47554 | 2.048062 | 0.427255 | 4.793535 | 1.64E-06 | 4.07E-05 | 1.72386 | 5.983845 | 1.468594 | 6.467976 | | 11 |
| 1143 | ENSG00000279364 | 12.79201 | 7.059462 | 1.753307 | 4.026369 | 5.66E-05 | 0.001012 | 0 | 2.519834 | 0 | 0.960318 | | 15 |
| 1144 | ENSG00000279417 | 153.5444 | 1.003765 | 0.307545 | 3.263799 | 0.001099 | 0.013443 | 3.38869 | 5.154807 | 2.612377 | 6.199776 | | 15 |
| 1145 | ENSG00000279512 | 88.19515 | 1.215034 | 0.387199 | 3.138012 | 0.001701 | 0.01936 | 3.128397 | 6.339007 | 3.231449 | 7.640088 | | 2 |
| 1146 | ENSG00000279521 | 187.5096 | 4.337392 | 0.361384 | 12.00217 | 3.46E-33 | 1.21E-30 | 0.788856 | 17.45345 | 0.99418 | 16.76311 | | 14 |
| 1147 | ENSG00000279693 | 38.52519 | -1.83661 | 0.59834 | -3.06951 | 0.002144 | 0.023396 | 2.630633 | 0.451985 | 2.201303 | 0.8335 | | 16 |
| 1148 | ENSG00000279694 | 35.97913 | 2.728722 | 0.653024 | 4.178593 | 2.93E-05 | 0.000558 | 0.52093 | 2.275611 | 0.367564 | 3.276245 | | 15 |
| 1149 | ENSG00000280105 | 15.87206 | -3.33675 | 1.017886 | -3.27812 | 0.001045 | 0.01286 | 0.754938 | 0.112083 | 1.108757 | 0.061079 | | 2 |
| 1150 | ENSG00000280109 | 103.2232 | 1.400495 | 0.37508 | 3.733851 | 0.000189 | 0.002931 | 0.651475 | 2.099425 | 0.849795 | 1.670313 | PLAC4 | 21 |
| 1151 | ENSG00000280138 | 502.3282 | -1.6572 | 0.190028 | -8.72083 | 2.76E-18 | 2.97E-16 | 4.719943 | 1.304062 | 4.221319 | 1.372348 | | 12 |
| 1152 | ENSG00000280255 | 87.88447 | -1.68971 | 0.392727 | -4.3025 | 1.69E-05 | 0.00034 | 5.790863 | 1.852996 | 6.154287 | 1.640424 | | 7 |
| 1153 | ENSG00000280339 | 65.09742 | -1.44781 | 0.452177 | -3.20186 | 0.001365 | 0.016103 | 2.9978 | 1.18413 | 3.840085 | 1.18858 | | 11 |
| 1154 | ENSG00000280424 | 34.76263 | -2.20946 | 0.662012 | -3.3375 | 0.000845 | 0.010734 | 1.80856 | 0.453849 | 1.350208 | 0.182257 | | 22 |
| 1155 | ENSG00000280650 | 81.82332 | -3.66043 | 0.475173 | -7.70336 | 1.33E-14 | 9.89E-13 | 1.825439 | 0.107078 | 1.5845 | 0.148604 | KCNIP4-IT | 4 |
| 1156 | ENSG00000280707 | 15.07236 | 7.354211 | 1.695662 | 4.337075 | 1.44E-05 | 0.000295 | 0 | 7.327608 | 0 | 13.71627 | | 6 |
| 1157 | ENSG00000281406 | 718.458 | -1.49333 | 0.210945 | -7.07926 | 1.45E-12 | 8.72E-11 | 15.12208 | 6.265212 | 22.88453 | 6.518754 | BLACAT1 | 1 |
| 1158 | ENSG00000281566 | 56.10783 | -6.01856 | 0.851852 | -7.06526 | 1.60E-12 | 9.55E-11 | 15.36412 | 0.415534 | 16.38574 | 0.096085 | | X |
| 1159 | ENSG00000281880 | 68.45432 | -2.2681 | 0.503939 | -4.50075 | 6.77E-06 | 0.00015 | 7.56867 | 0.746305 | 5.497162 | 1.890682 | PAUPAR | 11 |
| 1160 | ENSG00000282057 | 27.34561 | 2.114814 | 0.700886 | 3.017345 | 0.00255 | 0.026665 | 0.850197 | 2.988808 | 0.592927 | 2.75444 | | 1 |
| 1161 | ENSG00000282458 | 591.8734 | 1.706759 | 0.178941 | 9.538109 | 1.45E-21 | 2.05E-19 | 23.55386 | 65.51302 | 20.90863 | 71.46755 | WASH5P | 19 |
| 1162 | ENSG00000283128 | 137.7218 | 2.397845 | 0.372858 | 6.430983 | 1.27E-10 | 6.32E-09 | 1.170905 | 5.676996 | 1.792989 | 9.246765 | | 7 |
| 1163 | ENSG00000284620 | 14.16563 | -4.63998 | 1.351964 | -3.43203 | 0.000599 | 0.008035 | 8.409097 | 0.20761 | 0.939983 | 0.112907 | | 8 |
| 1164 | ENSG00000285103 | 133.522 | -1.02009 | 0.333763 | -3.05633 | 0.002241 | 0.024207 | 2.415666 | 1.188087 | 3.279883 | 1.47113 | | 9 |
| 1165 | ENSG00000285278 | 28.64355 | -2.40098 | 0.70124 | -3.4239 | 0.000617 | 0.008239 | 1.433914 | 0.235319 | 1.829033 | 0.352634 | | 6 |
| 1166 | ENSG00000285304 | 8.413951 | 6.453098 | 1.857092 | 3.47484 | 0.000511 | 0.006965 | 0 | 0.487926 | 0 | 0.70896 | | 22 |
| 1167 | ENSG00000285633 | 189.0246 | 4.174985 | 0.351124 | 11.89034 | 1.33E-32 | 4.48E-30 | 0.532148 | 7.877846 | 0.468726 | 9.182071 | | 14 |
| 1168 | ENSG00000286071 | 67.1316 | 1.477295 | 0.447629 | 3.300265 | 0.000966 | 0.011992 | 0.317461 | 0.64744 | 0.257834 | 0.867601 | | 1 |
| 1169 | ENSG00000286125 | 113.8299 | -1.38803 | 0.383901 | -3.61559 | 0.0003 | 0.004432 | 11.55309 | 3.215364 | 6.408596 | 3.253109 | | 19 |

| | | | | | | | | | | | | | | |
|---|---|---|---|---|---|---|---|---|---|---|---|---|---|---|
| 1170 | ENSG00000286214 | 2522.667 | -1.87184 | 0.186298 | -10.0476 | 9.42E-24 | 1.66E-21 | 67.39154 | 15.92621 | 43.9233 | 12.65325 | COPG2IT1 | | 7 |
| 1171 | ENSG00000286215 | 139.7717 | -2.27806 | 0.348375 | -6.53911 | 6.19E-11 | 3.17E-09 | 12.01666 | 2.342254 | 16.43581 | 3.279274 | | | 6 |
| 1172 | ENSG00000286320 | 35.10348 | -2.09081 | 0.627431 | -3.33233 | 0.000861 | 0.0109 | 2.01961 | 0.534686 | 1.880649 | 0.323734 | | | 4 |
| 1173 | ENSG00000286329 | 12.48502 | -5.67229 | 1.713423 | -3.3105 | 0.000931 | 0.011651 | 1.359587 | 0 | 1.363392 | 0.052537 | | | 3 |
| 1174 | ENSG00000286449 | 13.19942 | -7.2254 | 1.724918 | -4.18884 | 2.80E-05 | 0.000535 | 0.937804 | 0 | 0.550789 | 0 | | | 19 |
| 1175 | ENSG00000286635 | 54.74921 | 2.231412 | 0.509798 | 4.377051 | 1.20E-05 | 0.000251 | 0.734392 | 3.794141 | 0.84985 | 3.237252 | | | 16 |
| 1176 | ENSG00000286742 | 107.3512 | 1.777434 | 0.358536 | 4.957474 | 7.14E-07 | 1.89E-05 | 2.725377 | 8.127264 | 2.484707 | 8.735822 | | | 7 |
| 1177 | ENSG00000286757 | 245.5188 | 2.669781 | 0.311444 | 8.572279 | 1.01E-17 | 1.04E-15 | 2.14435 | 20.26067 | 4.491484 | 20.12821 | | | 13 |
| 1178 | ENSG00000287215 | 52.23358 | -1.86702 | 0.542168 | -3.44362 | 0.000574 | 0.007756 | 4.017495 | 0.671377 | 2.068042 | 0.906533 | | X | |
| 1179 | ENSG00000287289 | 8.54403 | 6.47675 | 1.86584 | 3.471225 | 0.000518 | 0.00705 | 0 | 0.854884 | 0 | 0.409892 | | | 8 |
| 1180 | ENSG00000287355 | 23.51343 | -2.57732 | 0.810976 | -3.17804 | 0.001483 | 0.017268 | 4.554211 | 0.363232 | 2.218324 | 0.711737 | | | 9 |
| 1181 | ENSG00000287373 | 18.55172 | -4.19298 | 1.059539 | -3.95736 | 7.58E-05 | 0.001315 | 0.970634 | 0.028021 | 1.29883 | 0.091619 | | | 11 |
| 1182 | ENSG00000287679 | 102.6219 | -1.10145 | 0.376094 | -2.92865 | 0.003404 | 0.033638 | 4.545609 | 1.537991 | 3.003206 | 1.778049 | | | 6 |
| 1183 | ENSG00000288079 | 334.1821 | 1.022764 | 0.220119 | 4.646406 | 3.38E-06 | 7.91E-05 | 5.994915 | 10.03166 | 5.33088 | 11.71094 | | | 3 |
| 1184 | ENSG00000288658 | 71.38126 | 1.828221 | 0.439723 | 4.157664 | 3.22E-05 | 0.000605 | 1.36747 | 4.437188 | 1.127785 | 3.898149 | | | 2 |
